# Design, Optimization and Development of RIPK1 Degraders with Improved Pharmacokinetic and Pharmacodynamic Properties

**DOI:** 10.1101/2025.09.06.674627

**Authors:** Dong Lu, Xin Yu, Hanfeng Lin, Ran Chen, Bin Yang, Min Zhang, Jingjing Chen, Feng Li, Xiaoli Qi, Jin Wang

## Abstract

The pivotal role of receptor-interacting protein kinase 1 (RIPK1) as a scaffold protein in mediating tumor resistance to immune checkpoint inhibitors (ICBs) underscores the significance of pharmacological RIPK1 degradation as a therapeutic strategy to enhance antitumor immunity. In this study, we present the design, synthesis, and evaluation of a novel series of RIPK1 degraders, derived from the optimization of the previously identified compound **LD4172**. Through systematic refinement of the linker, exit vector of the RIPK1 warhead, and the VHL ligand portion, we identified compound **LD5097** (**24b**), which exhibited potent RIPK1 degradation activity across various cancer cell lines, with DC_50_ values of single digit nanomolar range and inducing more than 95% maximum degradation. Remarkably, **LD5097** (**24b**) induced rapid and complete degradation of RIPK1 within 2 hours of treatment and enhanced TNFα-mediated apoptosis in Jurkat cells. Furthermore, proteomic profiling unveiled the high selectivity of **LD5097** (**24b**) in degrading RIPK1. **LD5097** (**24b**) exhibited excellent metabolic stability and pharmacokinetic properties, characterized by low clearance, an extended half-life, and high plasma drug concentrations. Notably, a single administration of **LD5097** (**24b**) effectively reduced RIPK1 protein levels in Jurkat xenograft tumor tissues in mice at both 6- and 24-hour post-administration. These findings underscore **LD5097** (**24b**) as a promising RIPK1 degrader candidate, offering potent activity, favorable pharmacokinetic profiles, and notable pharmacodynamic effects, thereby holding significant promise in cancer immunology therapies.

## INTRODUCTION

Receptor-interacting protein kinase 1 (RIPK1) plays a pivotal role in regulating cell fate and proinflammatory signaling downstream of various innate immune pathways, including those initiated by tumor necrosis factor-α (TNF-α), toll-like receptor (TLR) ligands, and interferons (IFNs).^1–6^ The kinase activity of RIPK1 is crucial in TNF-α signaling, where it is required for inducing apoptosis and necroptosis.^5^ Extensive investigations have revealed the significant contribution of RIPK1 to the pathogenesis of diverse disorders, ranging from ischemia-reperfusion injury^7^ and neurodegenerative diseases^4,8^ to inflammatory ailments^6,9^, infection disease^10,11^, and tumor metastasis.^12–15^

RIPK1 serves as a kinase-independent scaffolding protein, playing a pivotal role in recruiting the NF-κB activation complex, thereby activating the NF-κB pathway and promoting cell survival.^1,7^ Recent studies have established that the genetic ablation of RIPK1 in cancer cells significantly enhances tumor sensitivity to anti-PD-1 immunotherapy.^16–18^ This sensitization is associated with a beneficial remodeling of the tumor microenvironment (TME), characterized by an increased frequency of effector T cells, a decreased prevalence of immunosuppressive myeloid cells, and altered inflammatory cytokine profiles.^16–18^ Critically, pharmacological inhibition of RIPK1’s kinase activity or the expression of a kinase-dead mutant fails to replicate this synergy with anti-PD-1 blockade.^17,18^ These findings suggest that the pharmacological degradation of the entire RIPK1 protein, rather than its kinase inhibition, represents a more promising and potentially efficacious therapeutic strategy to synergize with immune checkpoint inhibitors (ICBs) and enhance antitumor immunity.

To date, a few RIPK1 degraders have been reported^19–22^. Utilizing the PROTAC technology, we previously developed a RIPK1 degrader **LD4172**.^19^ This degrader exhibited potent and highly specific induction of RIPK1 degradation across various cancer cell lines. Importantly, combining **LD4172** with anti-PD1 treatment has revealed substantial synergy, manifesting in a notable delay in tumor development.^19^ This observed synergistic effect closely mirrors the reported phenotype associated with the synergistic interplay between tumor-specific RIPK1 deletion and anti-PD1 treatment.^17,18^ In parallel, the RIPK1 degrader, R1-ICR-5, developed by the Pascal group showed synergistic anti-tumor effect when combined with radiotherapy.^21^ Collectively, these findings indicate that targeting RIPK1 with a degrader is a viable strategy for enhancing the efficacy of both cancer immunotherapies and radiotherapy.

Despite the promising preclinical data for **LD4172**, its suboptimal pharmacokinetic properties, specifically high *in vivo* clearance, and low plasma free drug concentration, pose challenges for achieving optimal RIPK1 degradation *in vivo*. In this study, we undertook an extensive medicinal chemistry optimization of **LD4172**, focusing on linker, exit vector, and VHL ligand refinement. These efforts led to the identification of the lead compound **LD5097** (**24b**), characterized by a significantly longer plasma half-life and higher plasma concentration. Importantly, **LD5097** (**24b**) demonstrated highly potent RIPK1 degradation *in vivo*, positioning it as a promising candidate for further development in cancer immunotherapies.

## RESULTS AND DISCUSSION

### Identification of the Metabolic Soft Spots of LD4172

First, we performed a metabolite identification study of **LD4172** in mouse liver S9 fraction. The results revealed that the major metabolic part and cleavage of the exit vector of the RIPK1 binder (Figure 1). Based on these findings, we conducted a comprehensive investigation into the structural modification of **LD4172**, with a specific focus on modifying the exit vector of the RIPK1 binder, linker, and the VHL ligand separately (Figure 2). This approach is aimed at developing more metabolically stable compounds.

**Figure 1.**
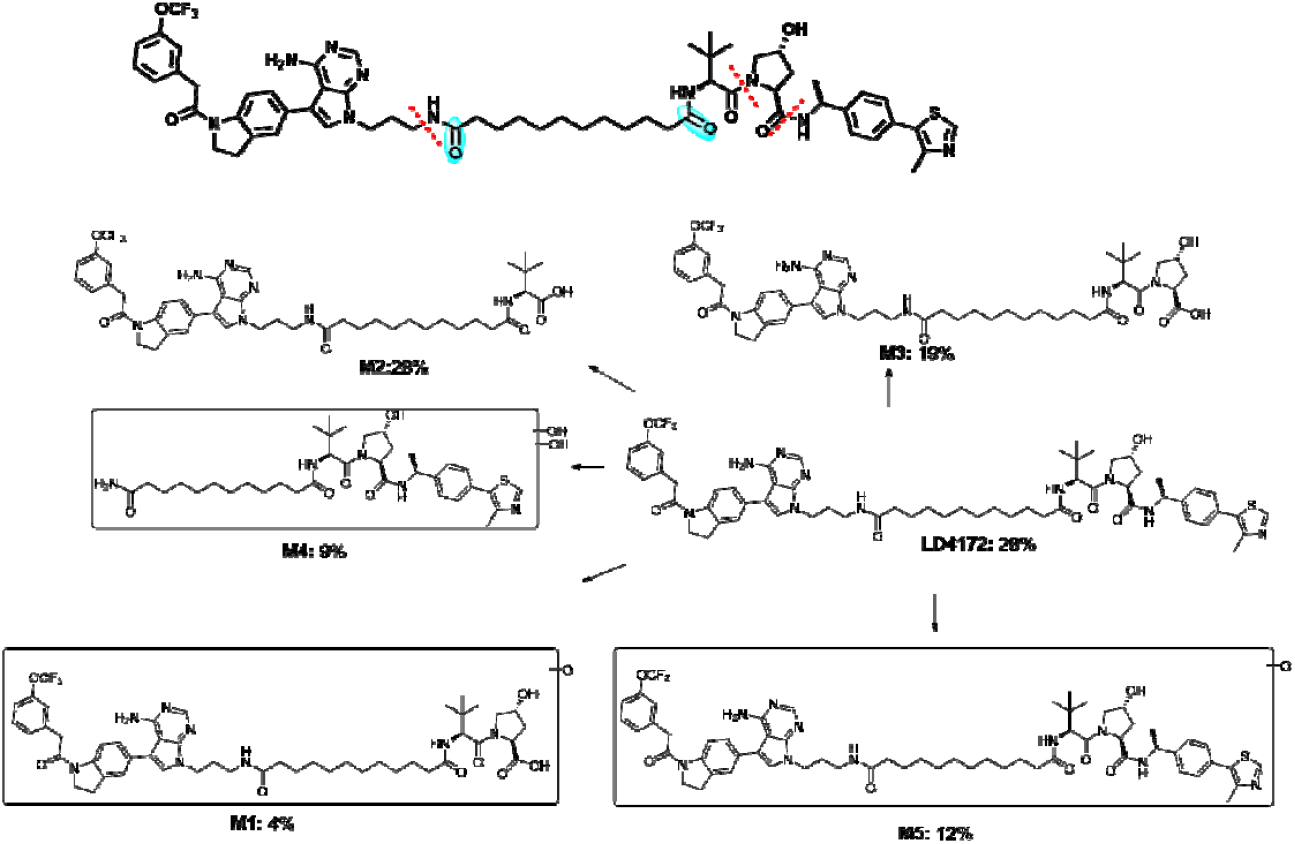
Metabolite identification of **LD4172**.

**Figure 2.**
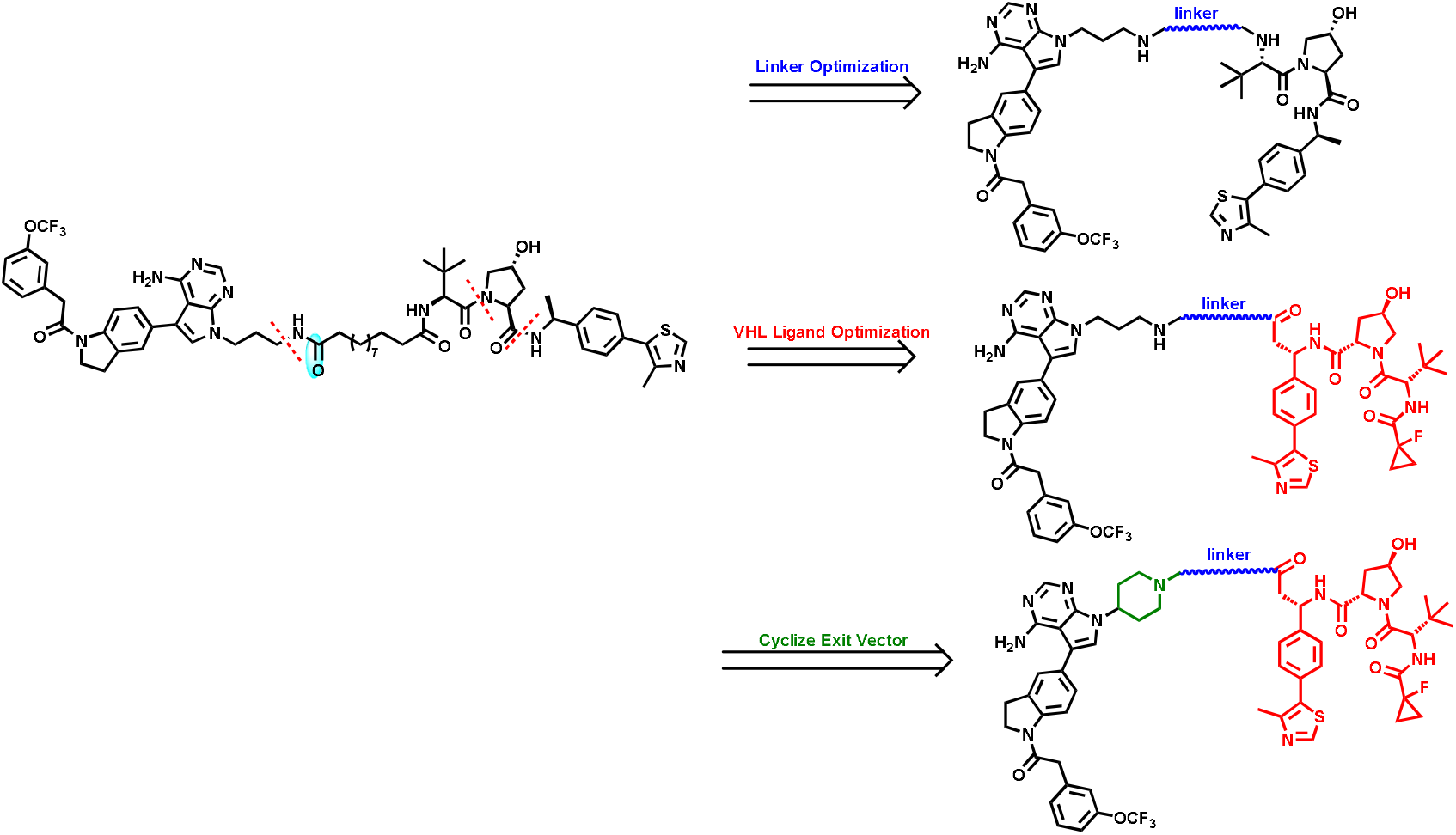
Optimization strategies of **LD4172** to improve metabolic stability.

### Chemistry

The synthesis of linkers **2** and **4** was outlined in Scheme1. The Sonogashira coupling reaction of the starting material ethyl 4-iodobenzoate followed by deprotection yielded the key linker fragment **1**. Alternatively, the nucleophilic aromatic substitution reaction of methyl 6-fluoronicotinate with tert-butyl 6-hydroxy-2-azaspiro[3.3]heptane-2-carboxylate followed by deprotection, resulted in another linker portion **3**. Subsequent nucleophilic substitution of **1** or **3** with different bromides in the presence of K_2_CO_3_ in acetonitrile, followed by ester hydrolysis afforded linkers **2** or **4**.

**Scheme 1.**
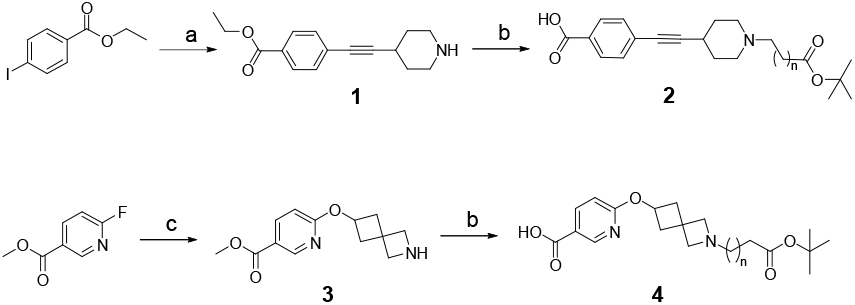
Synthesis Route of Linkers 2 and 4^a^. ^a^Reagents and conditions: (a) (i) Pd(PPh_3_)_2_Cl_2_, CuI, TEA, DMF, 100°C; (ii) TFA, DCM, rt; (b) (i) K_2_CO_3_, KI, MeCN, 80°C; (ii) LiOH, THF/H_2_O = 5/1, rt; (c) (i) NaH, THF, 60°C; (ii) TFA, DCM, rt.

As shown in Scheme 2, the Wittig reaction of 5-bromopicolinaldehyde gave α,β-unsaturated ester **5**. The selective 1,2-addition of **5** in the presence of cobalt chloride hexahydrate and sodium borohydride afforded **6**. The Sonogashira coupling products of **LD4172** involved amide hydrolysis of the VHL **li**gand reaction of **6** or tert-butyl 5-bromopicolinate followed by h**yd**rolysis, gave the key linkers **7-9**.

**Scheme 2.**
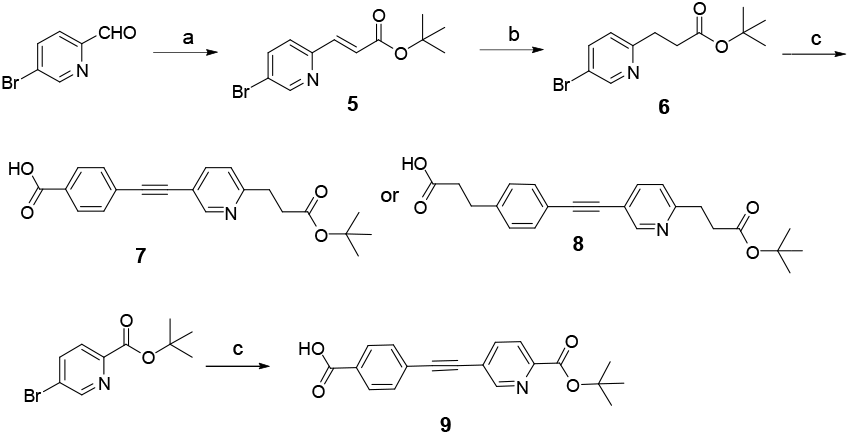
Synthesis Routes of Linkers 7-9^a^. ^a^Reagents and conditions: (a)(tert-Butoxycarbonylmethylene)triphenylphosphorane, THF, **rt;** (b) CoCl_2_.6H_2_O, NaBH_4_, MeOH,0°C-rt; (c) (i) Pd(PPh_3_)_2_Cl_2_, CuI, TEA, DMF,100°C; (ii) LiOH, THF/H_2_O = 5/1, rt.

**Scheme 3.**
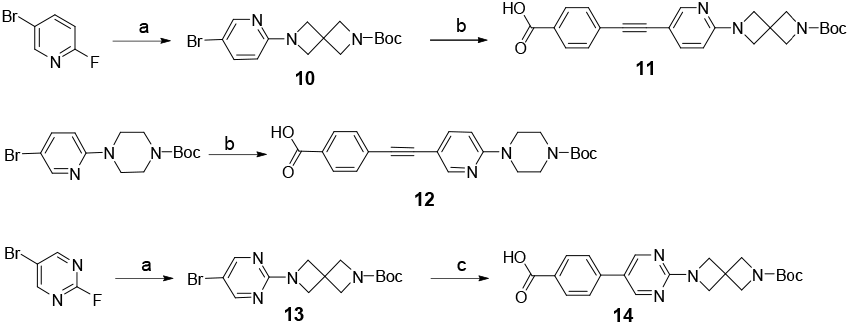
Synthesis Routes of Compounds 11-12and 14^a^. ^a^Reagents and conditions: (a) tert-butyl 2,6-diazaspiro[3.3]heptane-2-carboxylate, TEA, DMF,100°C; (b) (i) Pd(PPh_3_)_2_Cl_2_, CuI, TEA, DMF,100°C; (ii) LiOH, THF/H_2_O = 5/1, rt; (c) (i) Pd(dppf)_2_Cl_2_, K_2_CO_3_, 1,4-dioxane/H_2_O = 10/1,100°C; (ii) LiOH, THF/H_2_O = 5/1, rt.

The synthesis of linkers **11, 12** and **14** was depicted in Scheme 3. Initially, the key intermediate **10** was synthesized through the nucleophilic aromatic substitution reaction of 5-bromo-2-fluoropyridine with tert-butyl 2,6-diazaspiro[3.3]heptane-2-carboxylate. Subsequently, the Sonogashira coupling reaction of **10** or tert-butyl 4-(5-bromopyridin-2-yl)piperazine-1-carboxylate, followed by ester hydrolysis, yielded the key linker portions **11** or **12**, respectively. Moreover, the nucleophilic aromatic substitution reaction of 5-bromo-2-fluoropyrimidine with tert-butyl 2,6-diazaspiro[3.3]heptane-2-carboxylate produced intermediate **13**, which was transformed into linker **14** through Suzuki reaction followed by ester hydrolysis.

**Scheme 4.**
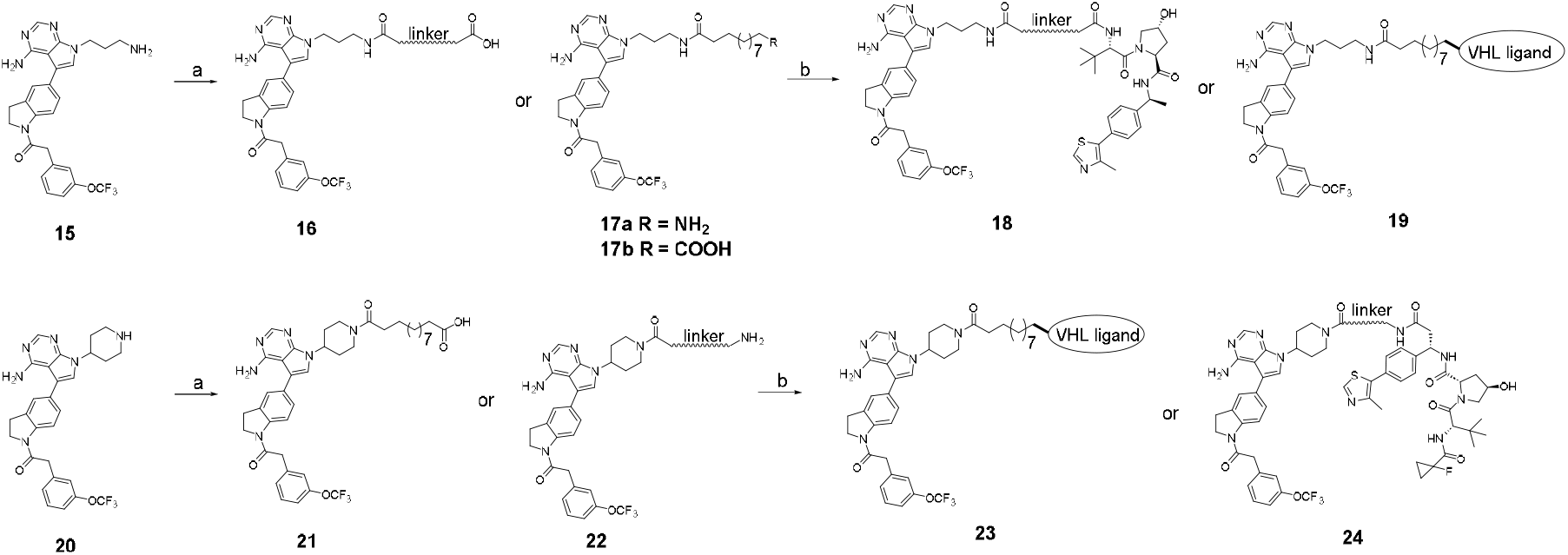
Synthesis Routes of Compounds 18-19 and 23-24^a^. ^a^Reagents and conditions: (a) (i) HATU, DIPEA, DMF, rt; (ii) TFA, DCM, rt; (b) HATU, DIPEA, DMF, rt.

Scheme 4 outlines the synthesis of target compounds **18-19** and **23-24**. Initially, the key intermediates **16** or **17** were synthesized through the coupling of compound **15** with various carboxylic acids, followed by deprotection. Subsequently, these intermediates were transformed into the target compounds **18** and **19** through coupling with different VHL ligands, either commercially available or synthesized following procedures.^23–26^ Intermediates **21** and **22** were obtained through the coupling reaction of compounds **20** with various linker acids, followed by deprotection. Finally, these intermediates were converted into the target compounds **23** and **24** through coupling with different VHL ligands.

### Optimization of the Linker Part of LD4172

It’s widely recognized that linker length, composition, and rigidity play crucial roles in determining the physicochemical and pharmacokinetic (PK) properties of PROTACs, along with their ternary complex formation and activity.^27–31^ Therefore, we systematically explored the impact of different linkers on both the potency of RIPK1 degradation and the metabolic stability in mouse liver S9 fraction. To facilitate high-throughput screening of RIPK1 degradation potency, we established a stable Jurkat cell line expressing nLuc-RIPK1, allowing luminescence-based quantification of RIPK1 levels.^32,33^ Please note that although we did not use CRISPR-Cas9 technology to endogenously knock-in nLuc or HiBiT to RIPK1, the concentration for 50% degradation of RIPK1 (DC_50_) values measured using this Jurkat cell line are consistent with the results from using Western blot. As shown in Table 1 and Figures S1 and S2, replacing the aliphatic chain of **LD4172** with PEG linkers or piperazine containing linkers (compounds **18a-e**) resulted in complete loss of degradation activities without improvement in stability. Although compounds **18f-g**, featuring triazole-containing linkers, exhibited a longer half-life (T_1/2_) in liver S9 fraction compared to **LD4172** (27.3 vs 12.6 mins), they only achieved maximum degradation (D_max_) of 24% and 46% RIPK1 in Jurkat cells, respectively. Similarly, compounds **18h-k**, incorporating saturated rings as linkers with flexible tails, demonstrated increased T_1/2_ in liver S9 fraction, but all exhibited a significant decrease in RIPK1 degradation potency. In contrast, compounds **18l-n**, utilizing unsaturated rings as linkers, displayed potent RIPK1 degradation activities with DC_50_ values of 5.9, 9.5 and 44.1 nM, respectively. Notably, compound **18m** showed enhanced stability compared to **LD4172** in mouse liver S9 fraction, with a T_1/2_ of 64.7 mins, highlighting the potential of introducing rigid linkers to improve the metabolic stability of degraders.

**Table 1.** Impact of Linker Composition on Degradation Potency and Metabolic Stability.

| No. | Linker | DC<sub>50</sub><br>(nM,<br>24h)<sup>a</sup> | D<sub>max</sub><sup>b</sup>,<br>(%,<br>24h) | T<sub>1/2</sub><br>(mins)<sup>c</sup> |
| --- | --- | --- | --- | --- |
| **LD4172** |  | 4.7 ± 0.4 | 94 ± 1 | 12.6<sup>c</sup>; 10.0<sup>d</sup> |
| **18a**<br>**T2P2VHL** |  | >1000 | 3 ± 2 | 9.7 |
| **18b**<br>**T2P3VHL** |  | >1000 | 18 ± 6 | NT<sup>e</sup> |
| **18c**<br>**T2P4VHL** |  | >1000 | 15 ± 2 | NT |
| **18d**<br>**LD5031** |  | >1000 | 32 ± 1 | 9.8<sup>e</sup> |
| **18e**<br>**LD5032** |  | >1000 | 19 ± 1 | NT |
| **18f**<br>**LD5029** |  | >1000 | 24 ± 2 | 27.3<sup>c</sup> |
| **18g**<br>**LD5030** |  | >1000 | 46 ± 1 | NT |
| **18h**<br>**LD5033** |  | >1000 | 47 ± 1 | 15.6<sup>c</sup> |
| **18i**<br>**LD5034** |  | >1000 | 28 ± 2 | NT |
| **18j**<br>**LD5035** |  | >1000 | 47 ± 2 | 26.4<sup>c</sup> |
| **18k**<br>**LD5036** |  | >1000 | 2 ± 1 | NT |
| **18l**<br>**LD5039** |  | 5.9 ± 1.5 | 87 ± 2 | NT |
| **18m**<br>**LD5037** |  | 9.5 ± 1.4 | 89 ± 1 | 64.7<sup>c</sup> |
| **18n**<br>**LD5038** |  | 44.1 ± 4.6 | 86 ± 1 | NT |

<sup>a</sup>The DC<sub>50</sub> of individual compound to nLuc-RIPK1 in Jurkat cells. Data are the mean of triplicate determinations. <sup>b</sup>The maximum degradation level of individual compound to nLuc-RIPK1 in Jurkat cells. Data are the mean of triplicate determinations. <sup>c</sup>The half-life time of individual compound incubated with mouse liver S9 fraction in the presence of NADPH. Data are the means of duplicate determinations. <sup>d</sup>The half-life time of individual compound incubated with human liver microsome in the presence of NADPH. Data are the means of duplicate determinations. <sup>e</sup>Not tested.

### Optimization of the VHL Ligand and Exit Vector of LD4172

Given the identification of major soft spots at the VHL ligand and exit vector, our subsequent efforts were directed towards modifying these two moieties to block the soft spots and enhance metabolic stability. Table 2 and Figures S1 and S2 illustrate our findings. Addition of an amide group at the benzyl site of the VHL ligand resulted in compound **19a**, which exhibited a 30-folds decrease in degradation potency compared to **LD4172**. However, changing the linkage attachment of the VHL ligand from the amine tail in **LD4172** to the benzyl site in compound **19b** maintained degradation potency, with a DC_50_ value of 7.2 nM. Moreover, the half-life (T_1/2_) of **19b** in mouse S9 liver fraction was 18.0 mins, slightly better than that of **LD4172**. Additionally, substitution of the linear amine of the RIPK1 warhead with a piperidine ring as the exit vector yielded compound **23a**, which demonstrated potent degradation activity for nLuc-RIPK1 with a DC_50_ value of 5.1 nM, comparable to that of **LD4172**. Importantly, compound **23a** exhibited significantly enhanced stability compared to **LD4172**, with much longer half-lives (T_1/2_) of 67.3 mins. Finally, attempts to modify the two amide portions within the VHL ligand, known metabolic soft spots, were made. However, any alteration in these regions resulted in a decrease or loss of degradation activity, as evidenced by compounds **23b-e**, although compounds **23b, 23d** and **23e** exhibited longer half-lives (T_1/2_) in liver S9 fraction. These findings underscore that changing the linkage attachment of the VHL ligand or utilizing a cyclic exit vector of the RIPK1 warhead further improved the metabolic stability of the degraders.

**Table 2.**
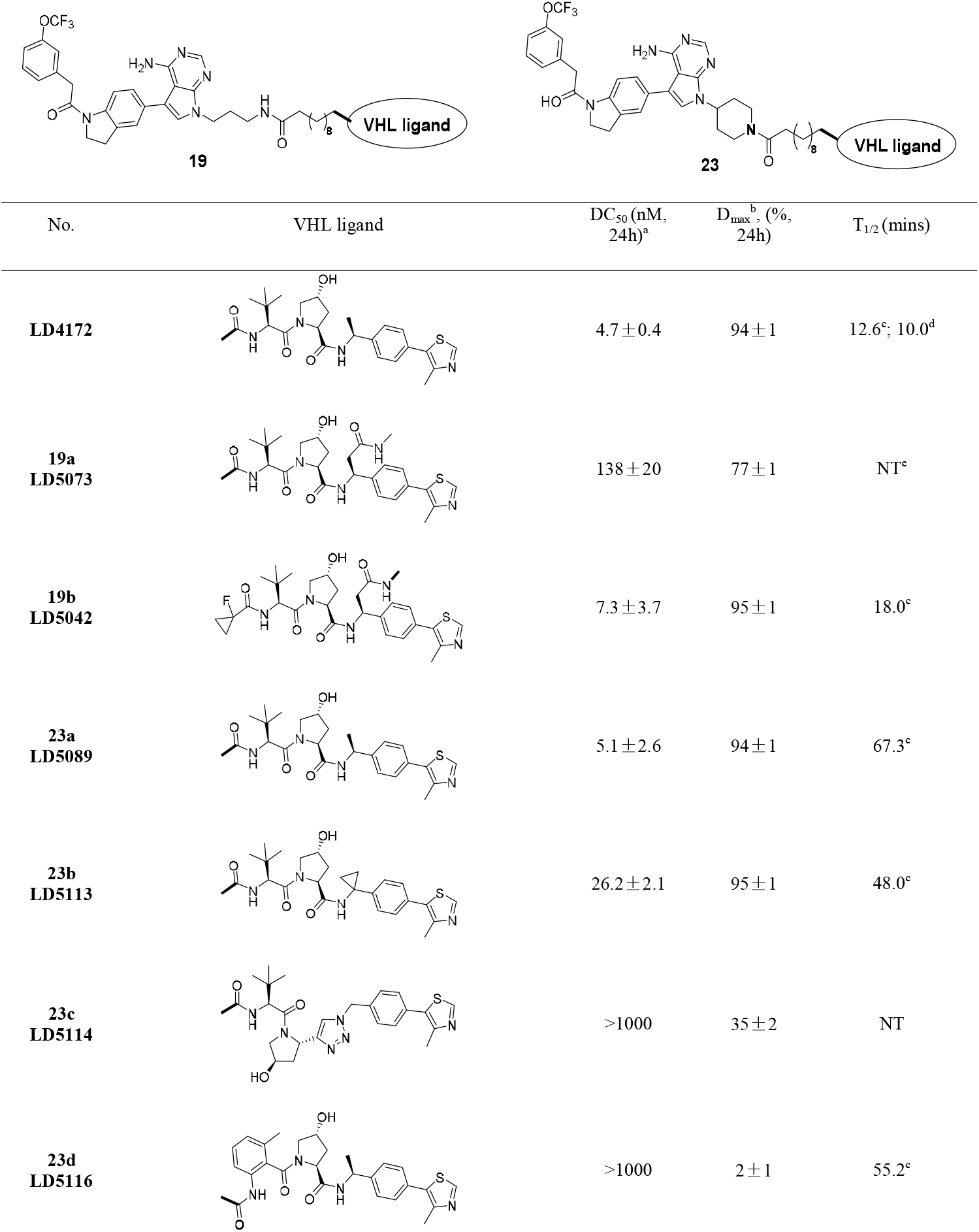
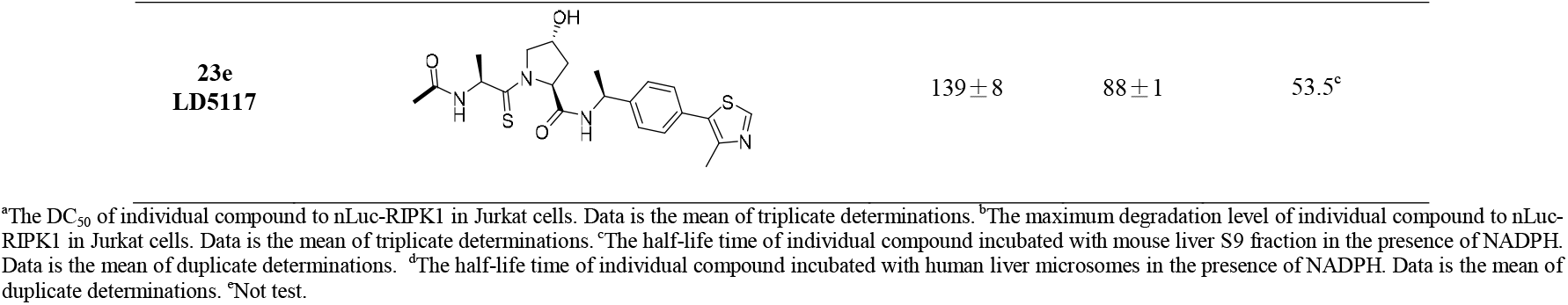
Effect of Exit Vector and VHL Ligand on Degradation Potency and Metabolic Stability.

### Evaluation of Pharmacokinetic Profile of First-generation RIPK1 Degraders

To further evaluate the *in vivo* stability of the optimized compounds, **LD4172, 18m**, and **23a** were selected for comprehensive pharmacokinetic studies. Male C57BL/6J mice (n=3) were intravenously administered at the same dose of 1.0 mg/kg. As summarized in Table 3 and Figure s4, **23a** displayed a clearance rate of 13.3 mL/min/kg, which was lower than that of **LD4172**. Consequently, in comparison to **LD4172, 23a** exhibited a higher plasma drug concentration with a maximum plasma concentration (C_max_) of 3420 ng/mL, and an area under the curve (AUC_0-t_) of 1239 h*ng/mL. Notably, compound **18m**, featuring a rigid linker, demonstrated markedly enhanced *in vivo* stability compared to both **LD4172** and compound **23a**, with a much lower clearance of 2.5 mL/min/kg, an extended plasma half-life (T_1/2_) of 4.6 h, and a higher AUC_0-t_ of 6596 h*ng/mL. Collectively, these findings highlight that optimized degrader with a rigid linker and cyclic exit vector exhibit significantly improved pharmacokinetic profiles.

**Table 3.** Pharmacokinetic Parameters of LD4172, 18m and 23a in C57BL/6J Mice^a^

|  | CL<br>(mL/min/kg) | T<sub>1/2</sub><br>(h) | C<sub>max</sub><br>(ng/mL) | AUC<sub>0-1ast</sub><br>(h*ng/mL) |
| --- | --- | --- | --- | --- |
| **LD4172** | 19.8 ± 1.8 | 3.3 ± 2.1 | 2333 ± 232 | 835 ± 84 |
| **18m** | 2.5 ± 0.3 | 4.6 ± 0.4 | 11867 ± 2178 | 6596 ± 662 |
| **23a** | 13.3 ± 0.8 | 3.1 ± 1.8 | 3420 ± 157 | 1239 ± 79 |

<sup>a</sup>C57BL/6J mice were treated with drug via intravenous administration (1 mg/kg, n=3)

### Design and Evaluation of Second-generation RIPK1 Degraders

Our initial exploration of optimization of exit vector of the RIPK1 binder, linker, and the VHL ligand revealed that introduction of cyclic exit vector, rigid linker and new VHL ligand could significantly enhance the *in vitro* metabolic stability and pharmacokinetic profile. Therefore, to further improve the stability of degraders, we incorporated the optimal moieties into degraders simultaneously to generate the second-generation RIPK1 degraders. As shown in Table 4 and Figure s1, all degraders **24a-c**, with combined optimal moieties exhibited very potent RIPK1 degradation activities with DC_50_ values of 1.8, 0.7 and 1.5 nM, respectively. These compounds could degrade more than 93% of RIPK1. Considering their potent biological activities, we then assessed their *in vitro* metabolic stabilities in both mouse liver S9 fraction and human hepatocytes. As expected, all of these degraders had much longer half-life (T_1/2_) in both mouse liver S9 fraction and human hepatocytes, demonstrating the enhanced *in vitro* metabolic stability (Table 4, Figures S2 and S3).

**Table 4.**
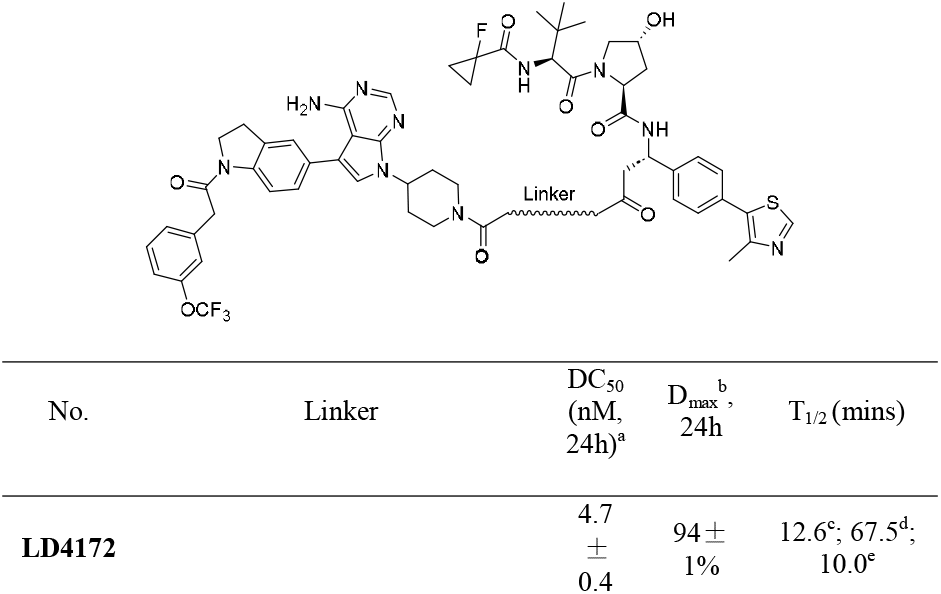
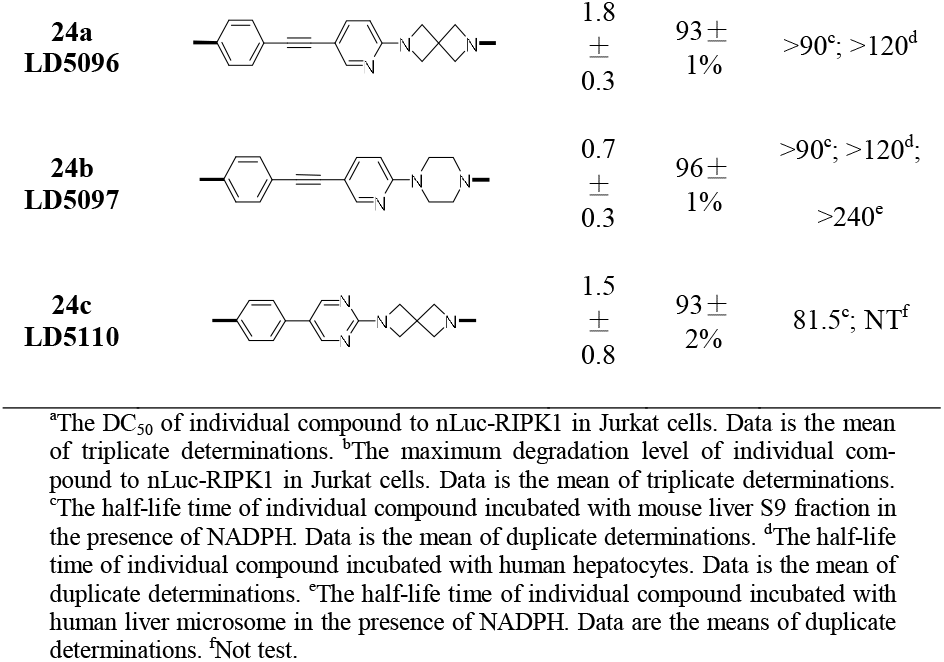
Evaluation of RIPK1 Degraders with Optimized Exit Vector and VHL Ligand

### 24a and 24b Exhibit Excellent Pharmacokinetic Profiles

The full pharmacokinetic profiles of **24a** and **24b** were measured in C57BL/6J mice via intravenous administration (1 mg/kg, n=3). As shown in Table 5 and Figure s4, **24a** and **24b** exhibited a plasma half-life (T_1/2_) of 5.9 and 7.3 h, area under the curve (AUC_0∼t_) of 5915 and 7256 h*ng/mL, and clearance of 2.8 and 2.3 mL/min/kg, respectively. Collectively, our results demonstrated that **24a** and **24b** had significantly improved half-lives and plasma concentration compared to **LD4172**, presenting excellent pharmacokinetic profiles.

**Table 5.** Pharmacokinetic Parameters of 24a and 24b in C57BL/6J Mice^a^

|  | Cl<br>(mL/min/kg) | T<sub>1/2</sub>(h) | C<sub>max</sub><br>(ng/mL) | AUC<sub>0-1ast</sub><br>(h*ng/mL) |
| --- | --- | --- | --- | --- |
| **24a** | 2.8 ± 0.2 | 5.9 ± 0.4 | 19000 ± 1153 | 5915 ± 334 |
| **24b** | 2.3 ± 0.1 | 7.3 ± 1.6 | 19633 ± 929 | 7256 ± 337 |

<sup>a</sup>C57BL/6J mice were treated with drug via intravenous administration (1 mg/kg, n=3)

### Identification of LD5097 (24b) as Potent RIPK1 Degraders Across Various Cancer Cell Lines

Considering the superior degradation activity and favorable pharmacokinetic properties, we designated **24b** as **LD5097** for further evaluation as the lead compound. To assess the potency of **LD5097** in degrading endogenous RIPK1 within cells, we conducted a western blot assay using various wild-type (WT) cancer cell lines. Treatment with increasing concentrations of **LD5097** resulted in a dose-dependent degradation of endogenous RIPK1 after 24 hours in Jurkat cells, with DC_50_ and D_max_ values of 2.6 nM and 98%, respectively (Figure 3a, b). Similarly, in both MOLM14 and U937 cells, **LD5097** exhibited potent degradation of endogenous RIPK1, with DC_50_ values of 2.8 nM and 0.8 nM, respectively. Remarkably, **LD5097** achieved maximal RIPK1 degradation at 200 nM, with D_max_ values of 99% observed in both cell lines (Figure S5).

**Figure 3.**
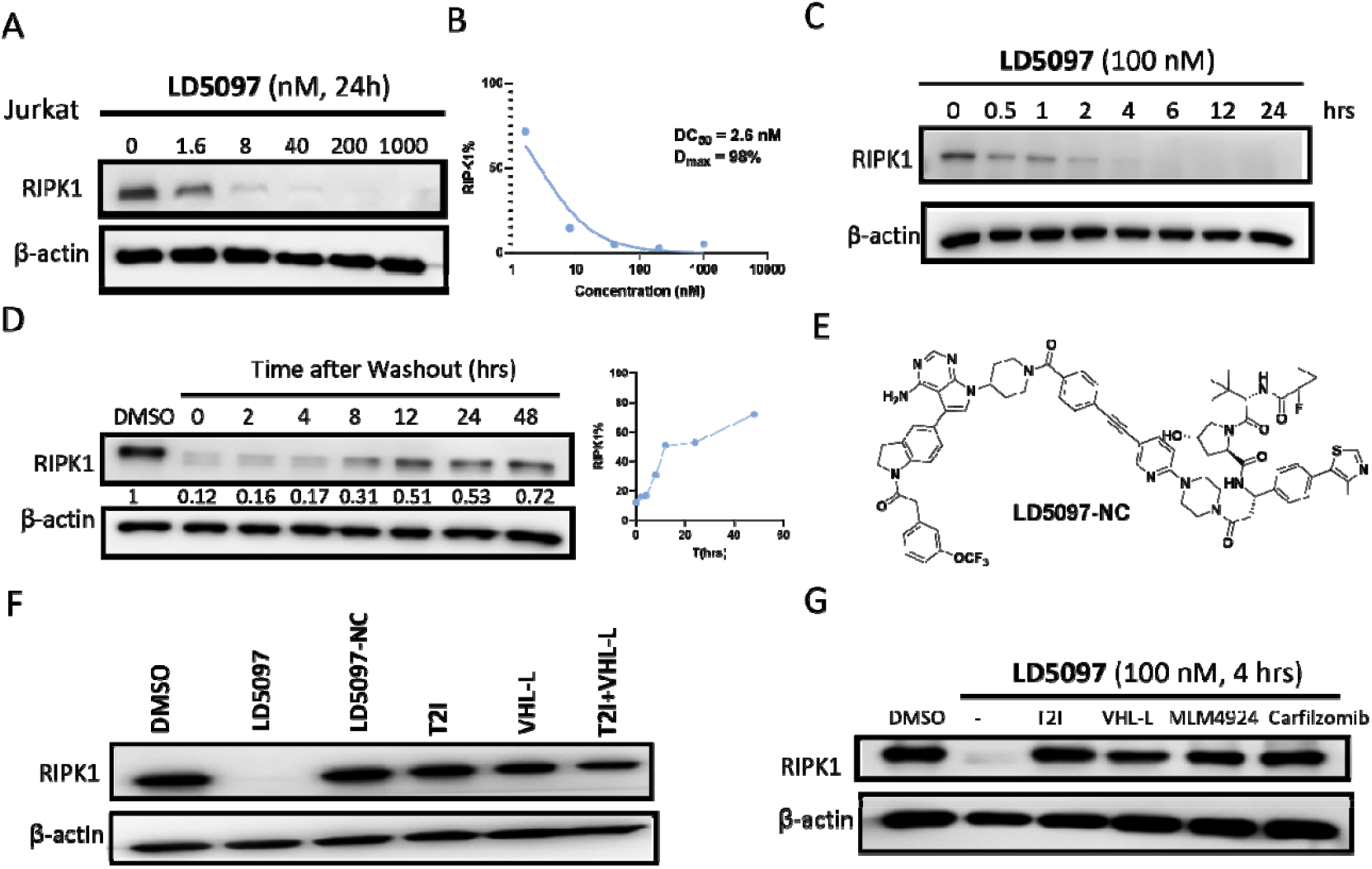
Degradation potency and mechanism validation of **LD5097** (**24b**) *in vitro*. (A) Degradation potency of **LD5097** (**24b**) for endogenous RIPK1 in Jurkat cells. (B) Dose-response of **LD5097** (**24b**) for RIPK1 degradation in both Jurkat cells. (C) Representative wstern blotting of RIPK1 level following treating Jurkat cells with **LD5097** (100 nM) for the indicated time points. (D) RIPK1 level recovery rate of Jurkat cells treated with **LD5097** (100 nM) for 24 hrs, washed 3 times with PBS and replenished with fresh media for indicated time points. (E) Chemical structure of **LD5097-NC. LD5097-NC** has an identical warhead and linker as **LD5097** (**24b**) but contains the inactive VHL ligand and is therefore unable to engage VHL to induce ubiquitination. (F) Representative western blotting of RIPK1 level following treating Jurkat cells with **LD5097** (100 nM), **LD5097-NC** (100 nM), RIPK1 inhibitor T2I (100 nM), VHL ligand (100 nM) or combination of T2I (100 nM) and VHL ligand (100 nM) for 24 hrs. (G) Jurkat cells treated with a proteasome inhibitor (Carfilzomib, 300 nM), a neddylation inhibitor (MLN4924, 1 µM), or excess RIPK1 inhibitor (T2I, 1 µM), VHL ligand (VHL-L, 1 µM) for 2 hours, then quently treated with DMSO or **LD5097** (100 nM) for 4 hours.

### Mechanism Validation of LD5097 *in vitro*

The degradation of RIPK1 induced by **LD5097** occurred rapidly, with complete degradation observed within 2 hours at 100 nM (Figure 3c). A washout experiment performed after a 24-hour treatment demonstrated 53% recovery of RIPK1 levels after 24 hours, suggesting a relatively slow turnover of RIPK1 protein (Figure 3d). To further validate the mechanism action of **LD5097**, we synthesized a negative control, **LD5097-NC**, which possesses a VHL ligand diastereomer that does not bind to VHL (Figure 3e). Treatment with **LD5097-NC**, RIPK1 inhibitor (T2I), VHL ligand (VHL-L)alone, or the combination of T2I and VHL-L did not affect **RI**PK1 levels, confirming that **LD5097** acts as a bifunctional PR**OT**AC, requiring stable ternary complex formation to induce RIPK**1** degradation in cells (Figure 3f). Additionally, RIPK1 degradat**ion** was prevented when cells were pre-treated with carfilzomi**b** (proteasome inhibitor), MLN-4924 (neddylation inhibitor), **10**-fold excess T2I, or VHL-L, providing further evidence that **LD5097** mediates a decrease in RIPK1 levels consistent with a PR**O**TAC mechanism of action (Figure 3g). Collectively, these f**ind**ings confirm that **LD5097** acts as a PROTAC dependent on th**e V**HL E3 ligase and ubiquitin-proteasome system to degrade RI**PK**1 in cells.

### LD5097 is a Highly Selective RIPK1 Degrader

The RIPK1 binder, T2I, utilized in our RIPK1 degraders is recognized as a typical type II kinase inhibitor with binding activity to certain off-targets kinases, including TrkA, Flt1, Flt4, Ret, Met, Mer, Fak, FGFR1, and MLK1.^34^ To assess the specificity of **LD5097**, we conducted mass spectrometry (MS)-based analysis of the entire cellular proteome of MOLM14 cells, a cell line known to express the entire kinome.^35^ MOLM14 cells were treated with either **LD5097** (100 nM) or DMSO for 6 hours, and we succ**ess**fully detected over 17,000 proteome isoforms. Our results demonstrated that **LD5097** degrades RIPK1 with high specificity (red dot in Figure 4), and no degradation of off-target kinases was observed (Table S1). This observation aligns with previous studies suggesting that PROTACs with potentially promiscuous target protein binders may achieve enhanced selectivity through protein-protein interactions involving the E3 ligase. ^35–37^

**Figure 4.**
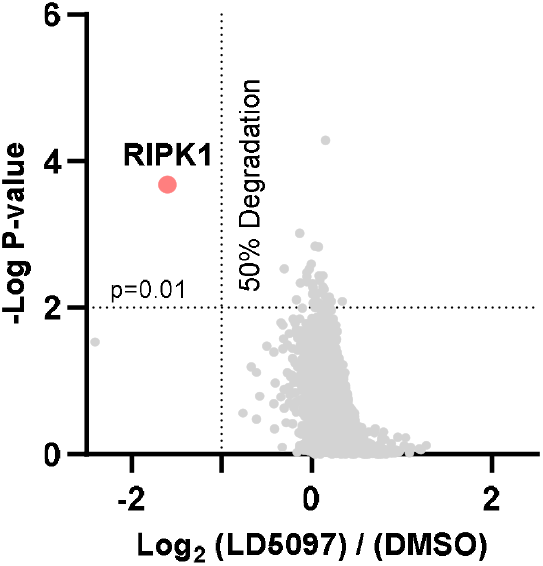
Proteome profiling of **LD5097** (**24b**) induced protein degradation. MOLM14 cells were treated with **LD5097** (**24b**) (100 nM) or DMSO for 6 hrs (n=3). In total, ∼17,000 proteome isoforms were quantified in the proteomics experiment. RIPK1 (red dot) is the only protein showing >50% degradation with P < 0.01.

### LD5097 Enhanced TNFα-mediated Apoptosis *in vitro*

In our previous study, we demonstrated that RIPK1 degraders could enhance cell apoptosis when combined with TNFα treatment, mirroring the effects observed in genetic knockout studies. To evaluate the apoptotic effect of **LD5097**, we treated Jurkat cells with various compounds, including **LD5097**, TNFα, and the pancaspase inhibitor Z-VAD-FMK. Our results revealed significant cell death, particularly apoptosis, induced by the combination of TNFα and **LD5097** (Figure 5a-b). This was further evidenced by elevated expression levels of cleaved caspase 3/7 and PARP, which are markers of apoptosis (Figure 5c). Importantly, this apoptosis could be reversed with the pan-caspase inhibitor, Z-VAD-FMK treatment (Figure 5a-c).

**Figure 5.**
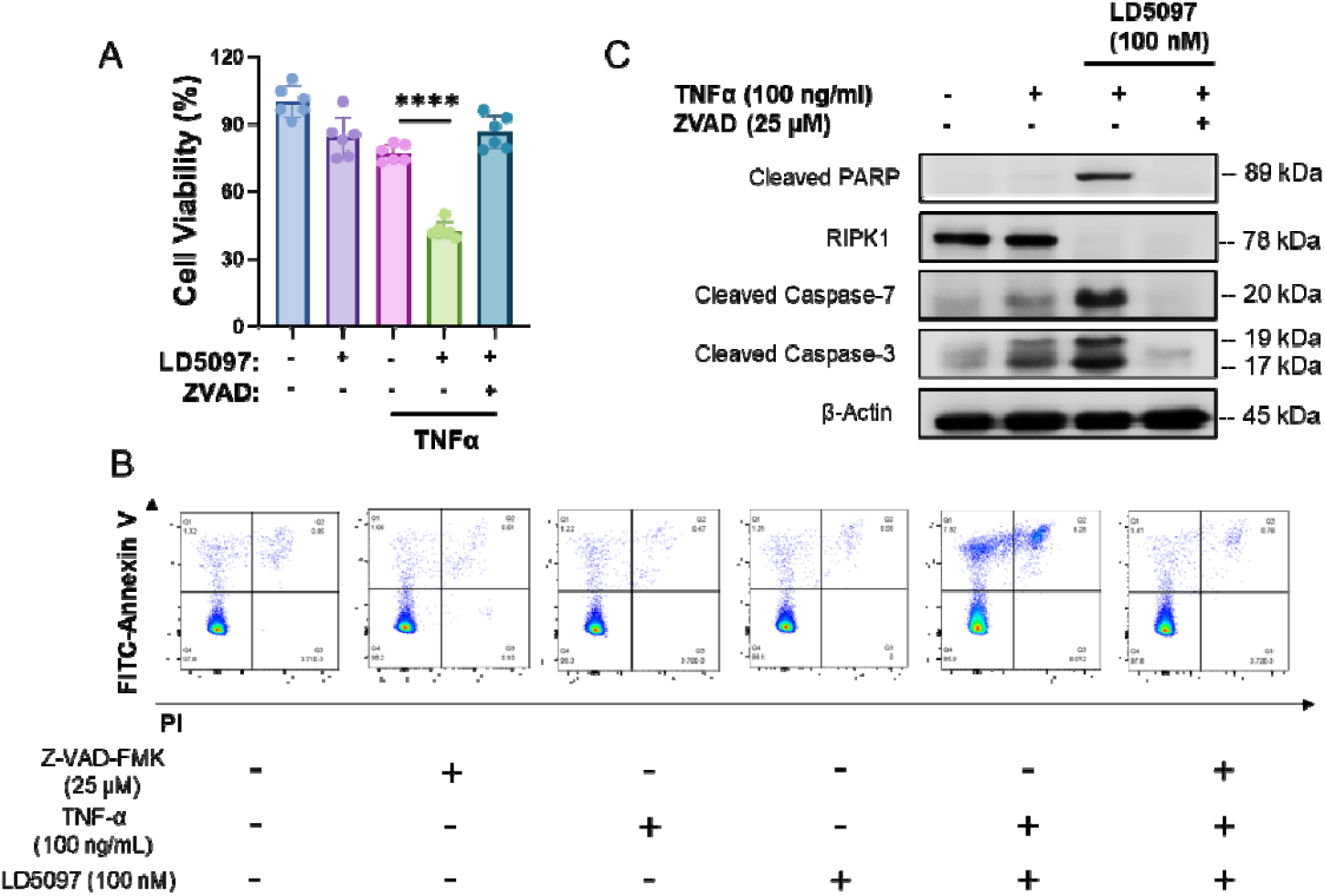
**LD5097** (**24b**) sensitizes Jurkat cells to TNFα-mediated apoptosis. (A) Cell viability of Jurkat cells treated with indicated treatments for 72 hrs (LD5097: 100 nM; TNFα: 100 ng/ml; Z-VAD-FMK: 25 μM). Data are expressed as the mean ± SEM. * p<0.05; ** p<0.01, *** p<0.001, **** p<0.0001, respectively. ns, indicates no statistical significance. (B) Representative flow cytometry dot plots of apoptosis. Jurkat cells were treated with indicated treatment followed by flow cytometry analysis. In all four plots, viable cells are seen in the left lower quadrant (FITC-/PI-), early apoptotic cells in the left upper quadrant (FITC+/PI-), and late apoptotic cells in the right upper quadrant (FITC+/PI+). (C) Western blots for the expression of cleaved caspase-3, cleaved caspase-7, and cleaved PARP in Jurkat cells with indicated treatment for 72 hrs.

### LD5097 Significantly Induced RIPK1 Degradation in Tumors

Subsequently, we assessed both **LD4172** and **LD5097** for their pharmacodynamic effects in Jurkat subcutaneous xenograft tumors in mice (Figure 6). Our findings clearly illustrate that a single intravenous administration of **LD4172** at 10 mg/kg failed to induce effective RIPK1 degradation in tumor tissue. In contrast, treatment with **LD5097** resulted in a reduction of RIPK1 protein levels by more than 70% at both 6- and 24-hour post-administration. These results highlight the superior pharmacodynamic effect of **LD5097**, attributed to its improved pharmacokinetic properties and potent *in vitro* degradation activity.

**Figure 6.**
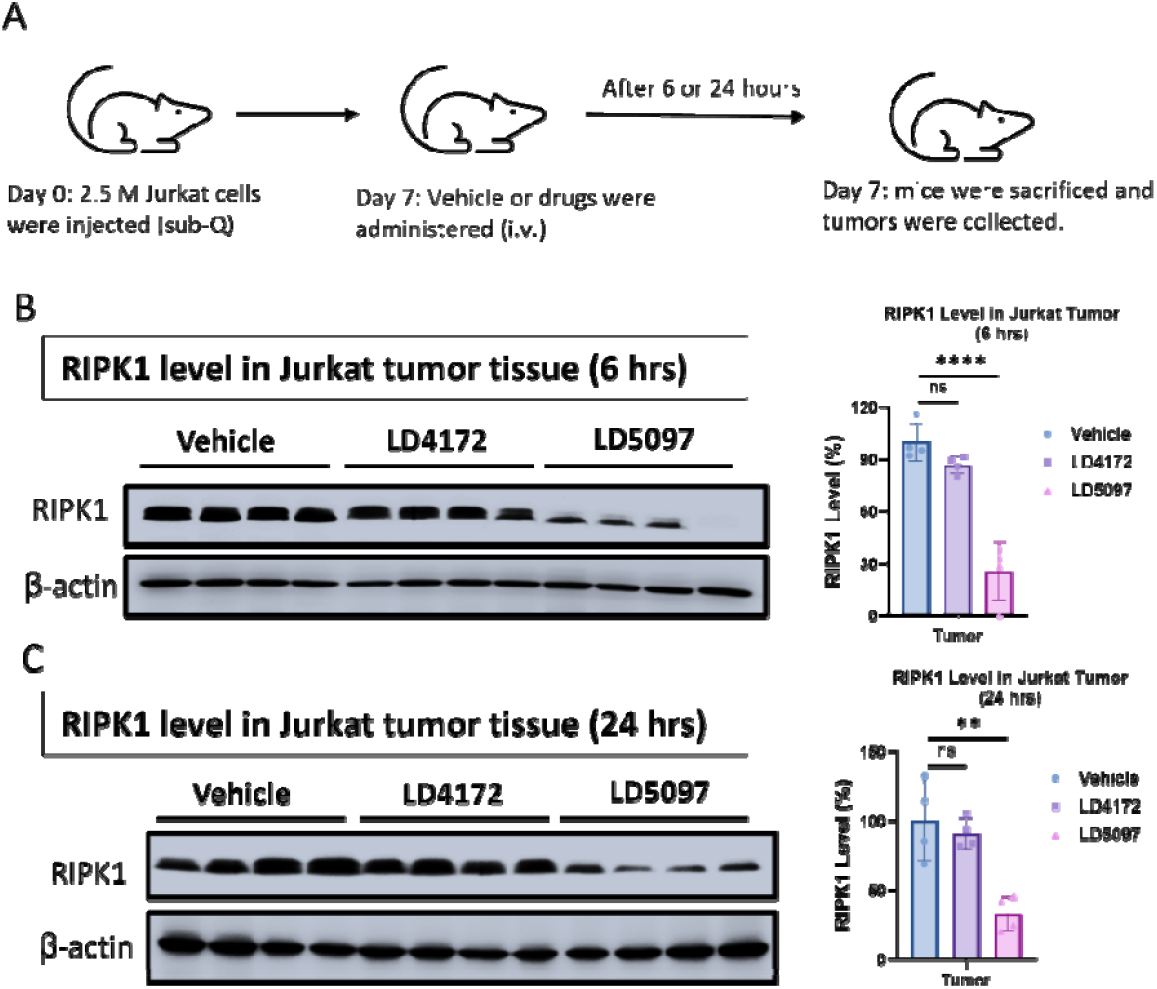
Pharmacodynamic (PD) evaluation of **LD5097** (**24b**) in Jurkat subcutaneous xenograft mouse model. (A) Scheme of PD study. (B) Representative western blotting of RIPK1 levels in tumor tissue. Tumor-bearing nude mice were intravenously administered with vehicle, **LD4172**, or **LD5097** (**24b**) (10 mg/kg, n = 4). Mice were sacrificed at the indicated time point (6 or 24 h), and tumor tissues were collected for analysis. (C) Densitometric analysis of RIPK1 levels in tumor tissue at 24 h (n = 4). Data are expressed as the mean ± SEM. * p<0.05; ** p<0.01, *** p<0.001, **** p<0.0001, respectively. ns, indicates no statistical significance.

### Species-Dependent Activity of LD5097

A critical aspect of the preclinical characterization of LD5097 is its pronounced species-dependent activity (Figure S7). While our previous work utilized the tool compound LD4172 to explore synergistic mechanisms,^19^ our functional assessment of LD5097 revealed a significant discrepancy in its degradative capacity between human and murine cell lines. For instance, LD5097 induced complete degradation of RIPK1 at a concentration of 100 nM in the human Ramos B lymphocyte cell line. In stark contrast, no degradation was observed in the murine A20 B-cell lymphoma line, even at concentrations as high as 10 µM (Figure S7a). In contrast, LD4172 with a flexible linker potently degrades RIPK1 in both human and murine cell lines (Figure S7b). This greater than 100-fold difference in activity underscores a fundamental species-specific limitation of LD5097.

The molecular basis for this species selectivity likely originates from the complex biology of proximity-inducing therapeutics, which rely on the successful formation of a productive ternary complex between the target protein, the E3 ligase, and the degrader molecule. One potential explanation is that amino acid sequence variations between human and mouse RIPK1, or the recruited E3 ligase, may diminish the stability of the ternary complex in the murine system. Such instability would lead to inefficient polyubiquitination and a lack of subsequent proteasomal degradation. Alternatively, the rigid linker of LD5097 imposes specific geometric constraints on the orientation of the E3 ligase relative to the target. It is plausible that sequence divergence leads to altered surface topology and a different spatial presentation of accessible lysine residues on mouse RIPK1. Consequently, the degrader may be unable to position the ligase in a conformation conducive to ubiquitin transfer in the murine context. A detailed investigation into these molecular determinants is beyond the scope of this study but will be a key focus of future work.

This pronounced species-dependent activity has significant translational implications. Primarily, it renders the mouse an inappropriate *in vivo* model for evaluating the preclinical toxicology and efficacy of LD5097. This challenge is analogous to that faced in biologics development, where sequence differences between human proteins and their orthologs in toxicology species often necessitate the creation of surrogate antibodies for safety assessment^38^. While the use of surrogate molecules is not yet standard practice for small molecules, the multi-protein engagement inherent to proximity-inducing therapeutics may amplify the functional consequences of even minor protein sequence variations, presenting a critical consideration for the clinical development of this therapeutic class.

## CONCLUSION

In this study, we undertook the design, synthesis, and evaluation of a series of RIPK1 degraders, building upon the drug-like optimization of our previously identified hit compound, **LD4172**. Our optimization efforts focused on refining the linker, exit vector of the RIPK1 warhead, and the VHL ligand portion in the designed RIPK1 degraders. Through these endeavors, we successfully identified a highly potent and metabolic stable RIPK1 degrader, designated as **LD5097**.

Compound **LD5097** demonstrated remarkable efficacy in inducing RIPK1 degradation, with DC_50_ values of 2.6, 2.8 and 0.8 nM observed in Jurkat, MOM14 and U937 cancer cell lines, respectively. Notably, **LD5097** exhibited rapid and complete degradation of RIPK1 in Jurkat cells within just 2 hours of treatment. Furthermore, proteomic profiling revealed that **LD5097** exhibited high selectivity in degrading RIPK1, with no discernible degradation of off-target proteins associated with the warhead utilized.

In addition to its impressive *in vitro* performance, **LD5097** displayed excellent metabolic stability and pharmacokinetic properties, characterized by low clearance, an extended half-life, and high plasma drug concentrations. Importantly, a single administration of **LD5097** effectively reduced RIPK1 protein levels in Jurkat xenograft tumor tissues in mice for over 24 hours. Collectively, our findings highlight **LD5097** as a promising candidate for RIPK1 degradation, exhibiting potent activity, favorable pharmacokinetic profiles, and notable pharmacodynamic effects.

## EXPERIMENTAL SECTION

### Chemistry

All reagents and solvents employed were purchased commercially and used as received. Reagents were purified prior to use unless otherwise stated. Column chromatography was carried out on a Yamazen Smart Flash EPCLC W-Prep 2XY and/or Agilent 1260 Infinity Preparative LC System. ^1^H NMR and ^13^C NMR spectral data were recorded in CDCl_3_, Acetone-*d*_*6*_, DMSO-*d*_*6*_, CD_3_OD on a Varian Palo Alto 400MHz NMR spectrometer and Chemical shifts (*δ*) were reported in parts per million (ppm), and the signals were described as brs (broad singlet), d (doublet), dd (doublet of doublet), m (multiple), q (quarter), s (singlet), and t (triplet). Coupling constants (J values) were given in Hz. Low-resolution mass spectra (ESI) was obtained using Agilent LC-MS 1200 series (6130 single quad). High-resolution mass spectra (ESI) was obtained using ThermoFisher Orbitrap Fusion Lumos Tribrid Mass Spectrometer. All final compounds had purity >95% determined by using High Pressure Liquid Chromatography (HPLC) using an Agilent Eclipse plus-C18 column eluting with a mixture of MeCN/Water (V:V = 80:20 plus 0.1% FA)

### General Procedures for the preparation of ethyl 4-(piperidin-4-ylethynyl)benzoate (1)

A solution of ethyl 4-iodobenzoate (552 mg, 2.0 mmol) and tert-butyl 4-ethynylpiperidine-1-carboxylate (460 mg, 2.2 mmol) in DMF (20 mL) was added Pd(PPh_3_)_2_Cl_2_ (140 mg, 0.2 mmol), CuI (38 mg, 0.2 mmol) and TEA (0.58 mL, 4.0 mmol). The resulting suspension was stirred for 8 hours at 100 °C. The mixture was cooled to room temperature and was poured into 60 mL water and the compound was extracted with EtOAc. The organic phase was washed with brine and then dried over MgSO_4_, filtered, and concentrated under reduced pressure. The residue was purified by flash chromatography (Hexane/EtOAc=4/1) to afford desired product (515 mg, 72%). MS (ESI): m/z 357.1 [M+1]^+^.

The above product was dissolved in DCM (10 mL) and the solution was added 1 mL TFA. The mixture was stirred for 2 hours at room temperature. The mixture was concentrated under reduced pressure to give compound **1** (326 mg, 88%). ^1^H NMR (400 MHz, DMSO-*d*_*6*_) *δ* 8.51 (m, 1H), 7.90 (d, *J* = 8.0 Hz, 2H), 7.52 (d, *J* = 7.9 Hz, 2H), 4.28 (q, *J* = 6.9 Hz, 2H), 3.20 (s, 2H), 2.96 (d, *J* = 33.8 Hz, 3H), 2.01 (d, *J* = 11.9 Hz, 2H), 1.74 (d, *J* = 10.1 Hz, 2H), 1.28 (t, *J* = 7.0 Hz, 3H). MS (ESI): m/z 257.1 [M+1]^+^.

### General Procedures for the preparation of compound (2)

A solution of **1** (100 mg, 0.39 mmol) in MeCN (5 mL) was added K_2_CO_3_ (64 mg, 0.46 mmol) and bromide (0.46 mmol) at room temperature. Then KI (65 mg, 0.39 mmol) was added and stirring was continued overnight at 80 °C. When TLC showed full conversion, the mixture was poured into 20 mL water and the compound was extracted with EtOAc. The organic phase was washed with brine and then dried over MgSO_4_, filtered, and concentrated under reduced pressure without further purification to afford desired compound.

The above product (0.2 mmol) and LiOH (10 mg, 0.4 mmol) were dissolved in THF (5 mL) and H_2_O (1 mL). This solution was stirred at room temperature overnight. The reaction was concentrated and then was purified by silica gel flash column chromatography (DCM/MeOH=20/1) to give the title compound.

4-((1-(3-(tert-butoxy)-3-oxopropyl)piperidin-4-yl)ethynyl)benzoic acid (**2a**): ^1^H NMR (400 MHz, DMSO-*d*_*6*_) *δ* 7.85 (d, *J* = 8.0 Hz, 2H), 7.45 (d, *J* = 7.6 Hz, 2H), 2.64 (s, 2H), 2.51 (s, 2H), 2.31 (t, *J* = 6.2 Hz, 2H), 2.16 (m, 2H), 1.80 (m, 3H), 1.55 (d, *J* = 10.0 Hz, 2H), 1.36 (s, 9H). MS (ESI): m/z 358.2 [M+1]^+^.

4-((1-(5-(tert-butoxy)-5-oxopentyl)piperidin-4-yl)ethynyl)benzoic acid (**2b**): ^1^H NMR (400 MHz, DMSO-*d*_*6*_) *δ* 7.88 (d, *J* = 7.8 Hz, 2H), 7.49 (d, *J* = 7.4 Hz, 2H), 2.85 (m, 4H), 2.21 (t, *J* = 7.0 Hz, 2H), 2.00 (m, 2H), 1.78 (m, 3H), 1.55 (m, 2H), 1.46 (m, 4H), 1.37 (s, 9H). MS (ESI): m/z 386.2 [M+1]^+^.

### General Procedures for the preparation of methyl 6-((2-azaspiro[3.3]heptan-6-yl)oxy)nicotinate (3)

The mixture of tertbutyl 6-hydroxy-2-azaspiro[3.3]heptane-2-carboxylate (500 mg, 2.3 mmol) in anhydrous THF (15 mL) was slowly added NaH (120 mg, 3 mmol) at 0 °C. The mixture was stirred at room temperature for 10 mins. Then methyl 6-fluoronicotinate (300 mg, 2 mmol) was added and the reaction was heated at 60 °C for 18 hours. After cooling to room temperature, the mixture was poured into 20 mL water and the compound was extracted with EtOAc. The organic phase was washed with brine and then dried over MgSO_4_, filtered, and concentrated under reduced pressure. The residue was purified by flash chromatography (Hexane/EtOAc=4/1) to afford desired compound (441 mg, 55%). MS (ESI): m/z 349.1 [M+1]^+^.

The above product was dissolved in DCM (10 mL) and the solution was added 1 mL TFA. The mixture was stirred for 2 hours at room temperature. The mixture was concentrated under reduced pressure to give compound **3** (266 mg, 85%). ^1^H NMR (400 MHz, CDCl_3_) *δ* 8.77 (s, 1H), 8.23 (d, *J* = 8.5 Hz, 1H), 6.76 (d, *J* = 8.5 Hz, 1H), 5.09 (d, *J* = 5.9 Hz, 1H), 4.19 (d, *J* = 18.4 Hz, 4H), 3.90 (s, 3H), 2.90 (d, *J* = 23.5 Hz, 2H), 2.42 (d, *J* = 6.8 Hz, 2H). MS (ESI): m/z 249.1 [M+1]^+^.

Compound **4** was prepared according to the procedure described above for compound **2**.

*6-((2-(3-(tert-butoxy)-3-oxopropyl)-2-azaspiro[3*.*3]heptan-6-yl)oxy)nicotinic acid (****4a***): ^1^H NMR (400 MHz, DMSO-*d*_*6*_) *δ* 8.65 (s, 1H), 8.11 (d, *J* = 8.1 Hz, 1H), 6.81 (d, *J* = 8.2 Hz, 1H), 5.07 (s, 1H), 2.89 (s, 2H), 2.73 (s, 2H), 2.55 (d, *J* = 7.3 Hz, 4H), 2.17 (d, *J* = 6.3 Hz, 4H), 1.37 (s, 9H). MS (ESI): m/z 363.2 [M+1]^+^.

*6-((2-(5-(tert-butoxy)-5-oxopentyl)-2-azaspiro[3*.*3]heptan-6-yl)oxy)nicotinic acid (****4b***): ^1^H NMR (400 MHz, DMSO-*d*_*6*_) *δ* 8.48 (s, 1H), 7.98 (d, *J* = 8.5 Hz, 1H), 6.59 (d, *J* = 8.4 Hz, 1H), 5.05 – 4.96 (m, 1H), 3.10 (s, 2H), 3.02 (s, 2H), 2.53 (dd, *J* = 10.2, 7.3 Hz, 2H), 2.25 (t, *J* = 6.9 Hz, 2H), 2.15 – 2.10 (m, 2H), 2.06 (d, *J* = 9.5 Hz, 2H), 1.46 – 1.39 (m, 2H), 1.35 (s, 9H), 1.19 (s, 2H). MS (ESI): m/z 391.2 [M+1]^+^.

### General Procedures for the preparation of tert-butyl (E)-3-(5-bromopyridin-2-yl)acrylate (5)

To 5-bromopicolinaldehyde (1 g, 5.0 mmol) in THF (30 mL) was added t-butoxycarbonyl methylene triphenylphosphorane (1.9 g, 5.0 mmol) and the resulting mixture was stirred at room temperature for 16 h. The reaction mixture was diluted with ether (20 mL) and was filtered through a pad of celite. The filtrate was collected and concentrated in vacuo to afford the desired product **5** (800 mg, 56%). ^1^H NMR (400 MHz, CDCl_3_) *δ* 8.65 (s, 1H), 7.80 (dd, *J* = 8.3, 1.9 Hz, 1H), 7.50 (d, *J* = 15.7 Hz, 1H), 7.28 (d, *J* = 8.3 Hz, 1H), 6.80 (d, *J* = 15.7 Hz, 1H), 1.51 (s, 9H). MS (ESI): m/z 284.1, 286.1 [M+1]^+^.

### General Procedures for the preparation of tert-butyl 3-(5-bromopyridin-2-yl)propanoate (6)

To compound **5** (1 g, 3.5 mmol) in methanol (15 ml) was added CoCl_2_.6H_2_O (52 mg, 0.35 mmol) and NaBH_4_ (665 mg, 17.5 mmol) at 0 °C. The resulting mixture was stirred at room temperature for 3 h. The reaction mixture was quenched by adding ice and was extracted with EtOAc. The organic phase was washed with brine and then dried over MgSO_4_, filtered, and concentrated under reduced pressure. The residue was purified by flash chromatography (Hexane/EtOAc=20/1) to afford compound **6** (420 mg, 42%). ^1^H NMR (400 MHz, CDCl_3_) *δ* 8.53 (s, 1H), 7.67 (dd, *J* = 8.3, 2.3 Hz, 1H), 7.06 (d, *J* = 8.2 Hz, 1H), 2.97 (s, 2H), 2.65 (t, *J* = 7.4 Hz, 2H), 1.38 (s, 9H). MS (ESI): m/z 286.1, 288.1 [M+1]^+^.

### General Procedures for the preparation of 4-((6-(3-(tert-butoxy)-3-oxopropyl)pyridin-3-yl)ethynyl)benzoic acid (7)

A solution of compound **6** (420 mg, 1.5 mmol) and methyl 4-ethynylbenzoate (290 mg, 1.8 mmol) in DMF (20 mL) was added Pd(PPh_3_)_2_Cl_2_ (105 mg, 0.15 mmol), CuI (28 mg, 0.15 mmol) and TEA (0.58 mL, 4.0 mmol). The resulting suspension was stirred for 8 hours at 100 °C. The mixture was cooled to room temperature and was poured into 15 mL water and the compound was extracted with EtOAc. The organic phase was washed with brine and then dried over MgSO_4_, filtered, and concentrated under reduced pressure. The residue was purified by flash chromatography (Hexane/EtOAc=4/1) to afford desired product (372 mg, 68%). MS (ESI): m/z 366.1 [M+1]^+^.

The above product (372 mg, 1.0 mmol) and LiOH (46 mg, 2.0 mmol) were dissolved in THF (5 mL) and H_2_O (1 mL). This solution was stirred at room temperature overnight. The reaction was concentrated and then was purified by silica gel flash column chromatography (DCM/MeOH=10/1) to give the title compound (317 mg, 90%). ^1^H NMR (400 MHz, DMSO-*d*_*6*_) *δ* 8.65 (s, 1H), 7.95 (d, *J* = 7.8 Hz, 2H), 7.87 (s, 1H), 7.65 (d, *J* = 7.8 Hz, 2H), 7.34 (d, *J* = 8.3 Hz, 1H), 2.96 (t, *J* = 7.2 Hz, 2H), 2.62 (t, *J* = 7.1Hz, 2H), 1.31 (s, 9H). MS (ESI): m/z 352.1 [M+1]^+^.

Compounds **8** and **9** were prepared according to the procedure described above for compound **7**.

*3-(4-((6-(3-(tert-butoxy)-3-oxopropyl)pyridin-3-yl)ethynyl)phenyl)propanoic acid (****8***): ^1^H NMR (400 MHz, DMSO-*d*_*6*_) *δ* 8.59 (s, 1H), 7.82 (d, *J* = 8.0 Hz, 1H), 7.45 (d, *J* = 6.6 Hz, 2H), 7.30 (d, *J* = 8.0 Hz, 1H), 7.26 (d, *J* = 7.3 Hz, 2H), 2.96 (t, *J* = 6.8 Hz, 2H), 2.82 (t, *J* = 7.1 Hz, 2H), 2.62 (t, *J* = 6.7 Hz, 2H), 2.52 (t, *J* = 7.4 Hz, 3H), 1.31 (s, 9H). MS (ESI): m/z 380.2 [M+1]^+^.

*4-((6-(tert-butoxycarbonyl)pyridin-3-yl)ethynyl)benzoic acid (****9***):

^1^H NMR (400 MHz, DMSO-*d*_*6*_) *δ* 8.73 (s, 1H), 8.04 (d, *J* = 8.0 Hz, 1H), 7.92 (d, *J* = 7.8 Hz, 2H), 7.62 (s, 1H), 7.53 (d, *J* = 7.1 Hz, 2H), 1.33 (s, 9H). MS (ESI): m/z 324.1 [M+1]^+^.

### General Procedures for the preparation of tert-butyl 6-(5-bromopyridin-2-yl)-2,6-diazaspiro[3.3]heptane-2-carboxylate (10)

The mixture of tert-butyl 2,6-diazaspiro[3.3]heptane-2-carboxylate (400 mg, 2.0 mmol) and 5-bromo-2-fluoropyridine (300 mg, 1.7 mmol) in anhydrous DMF (15 mL) was added TEA (0.5 mL, 3.4 mmol) at room temperature. The mixture was stirred at 100 °C for 12 hours. After cooling to room temperature, the mixture was poured into 20 mL water and the compound was extracted with EtOAc. The organic phase was washed with brine and then dried over MgSO_4_, filtered, and concentrated under reduced pressure. The residue was purified by flash chromatography (Hexane/EtOAc=4/1) to afford compound **10** (480 mg, 80%). ^1^H NMR (400 MHz, CDCl_3_) *δ* 8.13 (s, 1H), 7.50 (dd, *J* = 8.8, 2.3 Hz, 1H), 6.17 (d, *J* = 8.8 Hz, 1H), 4.08 (s, 8H), 1.42 (s, 9H). MS (ESI): m/z 354.1, 356.1 [M+1]^+^.

Compound **11-12** was prepared according to the procedure described above for compound **7**.

*4-((6-(6-(tert-butoxycarbonyl)-2,6-diazaspiro[3*.*3]heptan-2-yl)pyridin-3-yl)ethynyl)benzoic acid (****11***). ^1^H NMR (400 MHz, DMSO-*d*_*6*_) *δ* 8.25 (s, 1H), 7.90 (d, *J* = 7.6 Hz, 2H), 7.63 (d, *J* = 7.8 Hz, 1H), 7.56 (d, *J* = 7.8 Hz, 2H), 6.36 (d, *J* = 8.6 Hz, 1H), 4.09 (s, 4H), 4.00 (s, 4H), 1.34 (s, 9H). MS (ESI): m/z 420.2 [M+1]^+^.

*4-((6-(4-(tert-butoxycarbonyl)piperazin-1-yl)pyridin-3-yl)ethynyl)benzoic acid (****12***). ^1^H NMR (400 MHz, CDCl_3_) *δ* 8.35 (s, 1H), 7.99 (d, *J* = 8.2 Hz, 2H), 7.60 (d, *J* = 8.7 Hz, 1H), 7.54 (d, *J* = 8.2 Hz, 2H), 6.59 (d, *J* = 8.9 Hz, 1H), 3.59 (d, *J* = 4.7 Hz, 4H), 3.54 (d, *J* = 4.9 Hz, 4H), 1.47 (s, 9H). MS (ESI): m/z 408.1 [M+1]^+^.

### General Procedures for the preparation of tert-butyl 6-(5-bromopyrimidin-2-yl)-2,6-diazaspiro[3.3]heptane-2-carboxylate (13)

The mixture of tert-butyl 2,6-diazaspiro[3.3]heptane-2-carboxylate (400 mg, 2.0 mmol) and 5-bromo-2-fluoropyrimidine (300 mg, 1.7 mmol) in anhydrous DMF (15 mL) was added TEA (0.5 mL, 3.4 mmol) at room temperature. The mixture was stirred at 100 °C for 12 hours. After cooling to room temperature, the mixture was poured into 20 mL water and filtered to afford compound **13** (500 mg, 84%). ^1^H NMR (400 MHz, CDCl_3_) *δ* 8.29 (s, 2H), 4.20 (s, 4H), 4.09 (s, 4H), 1.43 (s, 9H). MS (ESI): m/z 355.1, 357.1 [M+1]^+^.

General Procedures for the preparation of 4-(6-(6-(tert-butoxycarbonyl)-2,6-diazaspiro[3.3]heptan-2-yl)pyridin-3-yl)benzoic acid (14). A solution of 13 (100 mg, 0.3 mmol), (4-(methoxycarbonyl)phenyl)boronic acid (72 mg, 0.4 mmol), K_2_CO_3_ (83 mg, 0.6 mmol) and Pd(dppf)_2_Cl_2_.CH_2_Cl_2_ (25 mg, 0.03 mmol) in 1,4-dioxane (10 mL) and water (1 mL) was degassed and fulfilled with N_2_ for three times. Then the mixture was stirred at 100 °C for 12 hours before it was quenched with water. The resulting mixture was extracted with EtOAc, the combined organic phases were dried over anhydrous Na_2_SO_4_ and concentrated under reduced pressure. The residue was purified by silica gel flash column chromatography (Hexane/EtOAc=1/1) to give the desired product (87 mg, 71%). MS (ESI): m/z 410.2 [M+1]^+^.

The above product (87 mg, 0.2 mmol) and LiOH (9.6 mg, 0.4 mmol) were dissolved in THF (5 mL) and H_2_O (1 mL). This solution was stirred at room temperature overnight. The reaction was concentrated and then was purified by silica gel flash column chromatography (DCM/MeOH=10/1) to give the title compound **14** (68 mg, 86%). ^1^H NMR (400 MHz, DMSO-*d*_*6*_) *δ* 8.73 (s, 2H), 7.94 (d, *J* = 6.8 Hz, 2H), 7.74 (d, *J* = 8.3 Hz, 2H), 4.19 (s, 4H), 4.01 (s, 4H), 1.34 (s, 9H). MS (ESI): m/z 396.2 [M+1]^+^.

### General Procedures for the preparation of compound (18)

To a solution of **15** (50 mg, 0.1 mmol) and carboxylic acid (0.12 mmol) in DMF (5 mL) was added HATU (57 mg, 0.15 mmol) and DIPEA (86 µL, 0.6 mmol). The mixture was stirred at room temperature for 1 hour. The reaction was concentrated and then was purified by silica gel flash column chromatography (DCM/MeOH=20/1) to give the desired compound.

The compound was then dissolved in DCM (2 mL) and was added TFA (0.5 mL). The mixture was stirred at room temperature for 5 hours before it was concentrated under reduced pressure. Crude product **16** was used directly in the next step.

To a solution of **16** (0.04 mmol) and (S,R,S)-AHPC-Me hydrochloride (0.05 mmol) in DMF (3 mL) was added HATU (28 mg, 0.07 mmol) and DIPEA (34 µL, 0.24 mmol). The mixture was stirred at room temperature for 1 hour. The reaction was concentrated and then was purified by reverse phase preparative HPLC (0-15mins: MeOH/H_2_O =10% → 80%; 15-25mins: MeOH/H_2_O =80%) to provide the title compound.

(2S,4R)-1-((S)-17-(4-amino-5-(1-(2-(3-(trifluoromethoxy)phenyl)acetyl)indolin-5-yl)-7H-pyrrolo[2,3-d]pyrimidin-7-yl)-2-(tert-butyl)-4,13-dioxo-7,10-dioxa-3,14-diazaheptadecanoyl)-4-hydroxy-N-((S)-1-(4-(4-methylthiazol-5-yl)phenyl)ethyl)pyrrolidine-2-carboxamide (**18a**): ^1^H NMR (400 MHz, DMSO-d_6_) *δ* 8.95 (s, 1H), 8.35 (d, J = 7.5 Hz, 1H), 8.08 (d, J = 12.3 Hz, 2H), 7.90 (s, 1H), 7.83 (d, J = 9.0 Hz, 1H), 7.47 – 7.37 (m, 3H), 7.34 (s, 1H), 7.33 – 7.26 (m, 5H), 7.24 – 7.17 (m, 2H), 6.05 (brs, 1H), 5.07 (s, 1H), 4.93 – 4.80 (m, 1H), 4.48 (d, J = 9.2 Hz, 1H), 4.38 (t, J = 8.0 Hz, 1H), 4.25 – 4.18 (m, 3H), 4.12 (s, 2H), 3.93 (s, 2H), 3.54 (m, 8H), 3.42 (s, 4H), 3.20 (t, J = 8.0 Hz, 2H), 3.01 (d, J = 5.4 Hz, 2H), 2.41 (s, 3H), 2.28 (s, 2H), 2.02 – 1.93 (m, 1H), 1.86 (m, 2H), 1.75 (s, 1H), 1.33 (d, J = 6.7 Hz, 3H), 0.88 (s, 9H). ^13^C NMR (100 MHz, DMSO-d_6_) *δ* 171.0, 170.4, 169.8, 168.8, 157.6, 151.9, 150.4, 148.6, 148.1, 145.0, 142.1, 138.4, 130.5 (q, J = 209 Hz), 130.4, 130.3, 130.1, 129.2, 126.8, 123.7, 122.7, 119.4, 116.6, 115.4, 100.3, 69.8, 69.1, 67.3, 58.9, 56.7, 48.2, 42.3, 41.83, 38.1, 36.6, 36.3, 36.0, 35.7, 30.2, 27.9, 26.8, 22.8, 16.4. MS (ESI): m/z 1125.5 [M+1]^+^. HRMS (ESI): m/z Calcd for (C_57_H_68_F_3_N_10_O_9_S) ([M+H]^+^): 1125.4844; found: 1125.4834.

(2S,4R)-1-((S)-20-(4-amino-5-(1-(2-(3-(trifluoromethoxy)phenyl)acetyl)indolin-5-yl)-7H-pyrrolo[2,3-d]pyrimidin-7-yl)-2-(tert-butyl)-4,16-dioxo-7,10,13-trioxa-3,17-diazaicosanoyl)-4-hydroxy-N-((S)-1-(4-(4-methylthiazol-5-yl)phenyl)ethyl)pyrrolidine-2-carboxamide (**18b**): ^1^H NMR (400 MHz, DMSO-d_6_) *δ* 8.94 (s, 1H), 8.35 (d, J = 7.6 Hz, 1H), 8.08 (d, J = 12.4 Hz, 2H), 7.90 (s, 1H), 7.83 (d, J = 9.3 Hz, 1H), 7.43 (dd, J = 15.2, 7.4 Hz, 2H), 7.38 (s, 1H), 7.34 (s, 1H), 7.33 – 7.26 (m, 5H), 7.24 – 7.18 (m, 2H), 6.03 (brs, 1H), 5.07 (s, 1H), 4.95 – 4.81 (m, 1H), 4.48 (d, J = 9.4 Hz, 1H), 4.38 (t, J = 8.0 Hz, 1H), 4.23 – 4.17 (m, 2H), 4.12 (s, 2H), 3.93 (s, 2H), 3.59 – 3.50 (m, 6H), 3.43 (s, 6H), 3.40 (s, 4H), 3.28 (s, 2H), 3.20 (t, J = 7.8 Hz, 2H), 3.01 (d, J = 5.2 Hz, 2H), 2.41 (s, 3H), 2.29 (d, J = 5.5 Hz, 2H), 2.02 – 1.93 (m, 1H), 1.86 (s, 2H), 1.76 (d, J = 8.0 Hz, 1H), 1.33 (d, J = 6.7 Hz, 3H), 0.88 (s, 9H). ^13^C NMR (100 MHz, DMSO-d_6_) *δ* 171.0, 170.5, 170.3, 169.8, 168.9, 157.6, 151.9, 150.4, 148.6, 148.1, 145.1, 142.1, 138.4, 130.5 (q, J = 209 Hz), 130.4, 130.3, 130.1, 129.2, 126.8, 125.3, 123.7, 122.6, 119.4, 116.6, 115.4, 100.3, 70.2, 69.1, 67.3, 58.9, 56.7, 48.2, 41.8, 41.4, 38.1, 36.6, 36.3, 36.1, 35.7, 30.2, 27.9, 26.8, 22.8, 16.4. MS (ESI): m/z 1169.5 [M+1]^+^. HRMS (ESI): m/z Calcd for (C_59_H_72_F_3_N_10_O_10_S) ([M+H]^+^): 1169.5106; found: 1169.5151.

N1-(3-(4-amino-5-(1-(2-(3-(trifluoromethoxy)phenyl)acetyl)indolin-5-yl)-7H-pyrrolo[2,3-d]pyrimidin-7-yl)propyl)-N16-((S)-1-((2S,4R)-4-hydroxy-2-(((S)-1-(4-(4-methylthiazol-5-yl)phenyl)ethyl)carbamoyl)pyrrolidin-1-yl)-3,3-dimethyl-1-oxobutan-2-yl)-4,7,10,13-tetraoxahexadecanediamide (**18c**): ^1^H NMR (400 MHz, DMSO-d_6_) *δ* 8.94 (s, 1H), 8.35 (d, J = 7.4 Hz, 1H), 8.08 (d, J = 11.5 Hz, 2H), 7.90 (s, 1H), 7.84 (d, J = 9.1 Hz, 1H), 7.48 – 7.36 (m, 3H), 7.34 (s, 1H), 7.33 – 7.25 (m, 5H), 7.21 (t, J = 9.2 Hz, 2H), 6.03 (brs, 1H), 5.07 (s, 1H), 4.97 – 4.80 (m, 1H), 4.49 (d, J = 9.1 Hz, 1H), 4.39 (t, J = 7.6 Hz, 1H), 4.21 (d, J = 9.2 Hz, 2H), 4.12 (s, 2H), 3.93 (s, 2H), 3.55 (d, J = 7.3 Hz, 6H), 3.42 (d, J = 8.8 Hz, 12H), 3.29 (s, 2H), 3.20 (t, J = 7.7 Hz, 2H), 3.01 (d, J = 4.6 Hz, 2H), 2.41 (s, 3H), 2.28 (s, 2H), 2.02 – 1.92 (m, 1H), 1.86 (s, 2H), 1.75 (s, 1H), 1.33 (d, J = 6.3 Hz, 3H), 0.89 (s, 9H). ^13^C NMR (100 MHz, DMSO-d_6_) *δ* 171.0, 170.5, 170.3, 169.8, 168.9, 157.6, 151.9, 150.4, 148.6, 148.2, 145.0, 142.2, 138.4, 130.5 (q, J = 209 Hz), 130.4, 130.3, 130.1, 129.2, 126.8, 125.3, 123.7, 122.6, 121.8, 119.3, 116.6, 115.3, 100.3, 70.3, 69.1, 67.3, 66.3, 58.9, 56.7, 51.7, 48.2, 41.8, 41.6, 38.1, 36.6, 36.3, 36.0, 35.7, 34.8, 30.2, 27.9, 26.8, 22.8, 16.4. MS (ESI): m/z 1213.5 [M+1]^+^. HRMS (ESI): m/z Calcd for (C_61_H_76_F_3_N_10_O_11_S) ([M+H]^+^): 1213.5368; found: 1213.5348.

(2S,4R)-1-((S)-2-(4-(4-(4-((3-(4-amino-5-(1-(2-(3-(trifluoromethoxy)phenyl)acetyl)indolin-5-yl)-7H-pyrrolo[2,3-d]pyrimidin-7-yl)propyl)amino)-4-oxobutyl)piperazin-1-yl)butanamido)-3,3-dimethylbutanoyl)-4-hydroxy-N-((S)-1-(4-(4-methylthiazol-5-yl)phenyl)ethyl)pyrrolidine-2-carboxamide (**18d**): ^1^H NMR (400 MHz, DMSO-d_6_) *δ* 8.94 (s, 1H), 8.35 (d, J = 7.3 Hz, 1H), 8.22 (s, 2H), 8.11 – 8.06 (m, 1H), 7.84 (s, 1H), 7.78 (t, J = 8.3 Hz, 1H), 7.42 (d, J = 13.1 Hz, 1H), 7.37 (d, J = 14.7 Hz, 3H), 7.32 (d, J = 8.6 Hz, 1H), 7.28 (s, 2H), 7.25 – 7.15 (m, 2H), 6.04 (brs, 1H), 4.87 (d, J = 5.9 Hz, 1H), 4.48 (d, J = 8.6 Hz, 1H), 4.39 (d, J = 7.6 Hz, 1H), 4.23 (s, 1H), 4.19 (d, J = 8.0 Hz, 1H), 4.12 (s, 2H), 3.93 (s, 2H), 3.56 (s, 2H), 3.33 (t, J = 6.1 Hz, 1H), 3.20 (t, J = 7.6 Hz, 1H), 3.13 (s, 1H), 2.99 (s, 2H), 2.41 (s, 3H), 2.41 – 2.20 (m, 8H), 2.16 (d, J = 15.1 Hz, 2H), 2.14 – 2.06 (m, 1H), 2.03 (d, J = 6.5 Hz, 2H), 1.96 (d, J = 8.5 Hz, 1H), 1.85 (d, J = 5.9 Hz, 2H), 1.75 (s, 2H), 1.60 (s, 4H), 1.35 – 1.30 (m, 3H), 0.89 (s, 10H). ^13^C NMR (100 MHz, DMSO-d_6_) *δ* 172.4, 172.3, 171.0, 170.0, 168.8, 157.6, 151.8, 145.0, 142.2, 138.4, 133.2, 130.5 (q, J = 209 Hz), 130.4, 130.1, 129.2, 126.7, 125.3, 123.6, 122.6, 119.4, 116.6, 69.1, 58.9, 57.6, 56.75, 52.9, 48.2, 42.0, 41.6, 40.4, 40.2, 40.1, 39.9, 39.7, 39.5, 39.3, 38.1, 36.4, 35.63, 33.5, 33.8, 33.2, 30.3, 27.9, 26.8, 22.8, 16.4. MS (ESI): m/z 1177.5 [M+1]^+^. HRMS (ESI): m/z Calcd for (C_61_H_76_F_3_N_12_O_7_S) ([M+H]^+^): 1177.5633; found: 1177.5623.

(2S,4R)-1-((2S)-2-(4-(4-(5-((3-(4-amino-5-(1-(2-(3-(trifluoromethoxy)phenyl)acetyl)indolin-5-yl)-7H-pyrrolo[2,3-d]pyrimidin-7-yl)propyl)amino)-5-oxopentyl)piperazin-1-yl)-3-methylbutanamido)-3,3-dimethylbutanoyl)-4-hydroxy-N-((S)-1-(4-(4-methylthiazol-5-yl)phenyl)ethyl)pyrrolidine-2-carboxamide (**18e**): ^1^H NMR (400 MHz, DMSO-d_6_) *δ* 8.94 (s, 1H), 8.35 (d, J = 7.4 Hz, 1H), 8.08 (d, J = 8.8 Hz, 2H), 7.83 (s, 1H), 7.76 (d, J = 9.0 Hz, 1H), 7.47 – 7.37 (m, 3H), 7.35 – 7.26 (m, 6H), 7.24 – 7.17 (m, 2H), 6.03 (brs, 1H), 4.94 – 4.80 (m, 1H), 4.47 (d, J = 8.9 Hz, 1H), 4.38 (t, J = 7.8 Hz, 1H), 4.26 – 4.17 (m, 3H), 4.11 (s, 2H), 3.93 (s, 2H), 3.23 – 3.17 (m, 2H), 3.00 (d, J = 4.9 Hz, 2H), 2.41 (s, 3H), 2.30 (s, 2H), 2.21 (d, J = 5.2 Hz, 6H), 2.02 (d, J = 6.7 Hz, 3H), 1.85 (s, 2H), 1.75 (s, 1H), 1.43 (s, 4H), 1.33 (d, J = 6.7 Hz, 7H), 0.89 (s, 9H). ^13^C NMR (100 MHz, DMSO-d_6_) *δ* 172.5, 172.4, 171.0, 168.9, 157.6, 151.9, 150.4, 148.6, 148.2, 145.1, 142.2, 138.4, 130.5 (q, J = 210 Hz), 130.3, 130.1, 126.8, 125.3, 123.6, 122.6, 119.4, 116.6, 115.5, 100.4, 69.1, 58.9, 57.7, 56.8, 56.7, 52.9, 48.3, 48.1, 42.1, 41.6, 38.1, 36.3, 35.7, 35.6, 30.3, 27.9, 26.8, 26.1, 23.8, 23.6, 22.8, 16.4. MS (ESI): m/z 1205.5 [M+1]^+^. HRMS (ESI): m/z Calcd for (C_63_H_80_F_3_N_12_O_7_S) ([M+H]^+^): 1205.5946; found: 1205.5928.

(2S,4R)-1-((S)-2-(5-(4-(4-((3-(4-amino-5-(1-(2-(3-(trifluoromethoxy)phenyl)acetyl)indolin-5-yl)-7H-pyrrolo[2,3-d]pyrimidin-7-yl)propyl)amino)-4-oxobutyl)-1H-1,2,3-triazol-1-yl)pentanamido)-3,3-dimethylbutanoyl)-4-hydroxy-N-((S)-1-(4-(4-methylthiazol-5-yl)phenyl)ethyl)pyrrolidine-2-carboxamide (**18f**): ^1^H NMR (400 MHz, DMSO-d_6_) *δ* 8.94 (s, 1H), 8.34 (d, J = 6.8 Hz, 1H), 8.08 (d, J = 9.5 Hz, 2H), 7.89 – 7.80 (m, 2H), 7.79 (s, 1H), 7.45 – 7.37 (m, 3H), 7.35 – 7.26 (m, 6H), 7.25 – 7.16 (m, 2H), 6.09 (brs, 1H), 5.07 (s, 1H), 4.87 (s, 1H), 4.46 (d, J = 9.1 Hz, 1H), 4.38 (t, J = 7.3 Hz, 1H), 4.24 (d, J = 5.3 Hz, 2H), 4.19 (d, J = 7.9 Hz, 2H), 4.12 (s, 2H), 3.93 (s, 2H), 3.55 (s, 2H), 3.20 (d, J = 7.5 Hz, 2H), 3.00 (s, 2H), 2.54 (d, J = 6.7 Hz, 2H), 2.41 (s, 3H), 2.23 (d, J = 6.7 Hz, 1H), 2.11 (d, J = 19.9 Hz, 4H), 1.97 (s, 1H), 1.86 (s, 2H), 1.74 (dd, J = 13.8, 7.7 Hz, 4H), 1.40 (s, 2H), 1.33 (d, J = 6.1 Hz, 3H), 1.15 (dd, J = 16.8, 10.7 Hz, 2H), 0.88 (s, 9H). ^13^C NMR (100 MHz DMSO-d_6_) *δ* 172.15 (d, J = 13.5 Hz), 151.89 (s), 130.53 – 129.95 (m), 129.33 (d, J = 19.8 Hz), 127.56 (s), 126.79 (s), 125.35 (s), 123.77 (s), 122.66 (s), 122.17 (s), 119.42 (s), 116.60 (s), 69.18 (s), 58.96 (s), 56.76 (d, J = 14.0 Hz), 49.26 (s), 48.19 (d, J = 16.5 Hz), 42.09 (s), 41.65 (s), 40.45 (d, J = 21.0 Hz), 40.16 (s), 40.14 (s), 39.93 (s), 39.72 (s), 39.51 (s), 39.30 (s), 38.14 (s), 36.38 (s), 35.59 (s), 35.32 (s), 34.52 (s), 30.33 (s), 29.76 (s), 27.92 (s), 26.84 (s), 25.64 (s), 25.08 (s), 22.82 (d, J = 5.8 Hz), 16.40 (s). MS (ESI): m/z 1174.5 [M+1]^+^. HRMS (ESI): m/z Calcd for (C_60_H_71_F_3_N_13_O_7_S) ([M+H]^+^): 1174.5272; found: 1174.5261.

(2S,4R)-1-((S)-2-(6-(4-(5-((3-(4-amino-5-(1-(2-(3-(trifluoromethoxy)phenyl)acetyl)indolin-5-yl)-7H-pyrrolo[2,3-d]pyrimidin-7-yl)propyl)amino)-5-oxopentyl)-1H-1,2,3-triazol-1-yl)hexanamido)-3,3-dimethylbutanoyl)-4-hydroxy-N-((S)-1-(4-(4-methylthiazol-5-yl)phenyl)ethyl)pyrrolidine-2-carboxamide (**18g**): ^1^H NMR (400 MHz, DMSO-d_6_) *δ* 8.94 (s, 1H), 8.34 (d, J = 7.6 Hz, 1H), 8.12 – 8.05 (m, 2H), 7.84 (s, 1H), 7.77 (d, J = 7.9 Hz, 2H), 7.46 – 7.37 (m, 3H), 7.31 (dd, J = 15.6, 6.7 Hz, 6H), 7.24 – 7.17 (m, 2H), 6.12 (brs, 1H), 5.08 (s, 1H), 4.92 – 4.83 (m, 1H), 4.47 (d, J = 9.2 Hz, 1H), 4.38 (t, J = 7.9 Hz, 1H), 4.21 (d, J = 7.7 Hz, 4H), 4.12 (s, 2H), 3.93 (s, 2H), 3.56 (s, 2H), 3.22 – 3.17 (m, 2H), 3.00 (d, J = 5.1 Hz, 2H), 2.55 (s, 2H), 2.41 (s, 3H), 2.18 (dd, J = 13.9, 7.2 Hz, 1H), 2.06 (s, 2H), 2.01 – 1.92 (m, 1H), 1.85 (d, J = 6.4 Hz, 2H), 1.73 (d, J = 7.8 Hz, 4H), 1.54 – 1.41 (m, 6H), 1.33 (d, J = 6.8 Hz, 3H), 1.21 – 1.09 (m, 4H), 0.88 (s, 9H). ^13^C NMR (100 MHz, DMSO-d_6_) *δ* 172.4, 172.3, 171.0, 170.0, 168.9, 157.2, 151.9, 150.3, 148.6, 148.1, 147.0, 145.0, 142.2, 138.4, 130.5 (q, J = 210 Hz), 130.4, 130.1, 129.2, 127.6, 126.8, 125.3, 123.8, 122.7, 121.9, 119.4, 116.6, 115.6, 69.2, 58.9, 56.8, 49.4, 48.3, 48.1, 42.1, 41.6, 38.1, 36.3, 35.5, 35.0, 30.3, 29.8, 29.0, 27.9, 26.8, 25.9, 25.2, 22.8, 16.4. MS (ESI): m/z 1202.5 [M+1]^+^. HRMS (ESI): m/z Calcd for (C_62_H_75_F_3_N_13_O_7_S) ([M+H]^+^): 1202.5585; found: 1202.5583.

(2S,4R)-1-((S)-2-(3-(4-((4-((3-(4-amino-5-(1-(2-(3-(trifluoromethoxy)phenyl)acetyl)indolin-5-yl)-7H-pyrrolo[2,3-d]pyrimidin-7-yl)propyl)carbamoyl)phenyl)ethynyl)piperidin-1-yl)propanamido)-3,3-dimethylbutanoyl)-4-hydroxy-N-((S)-1-(4-(4-methylthiazol-5-yl)phenyl)ethyl)pyrrolidine-2-carboxamide (**18h**): ^1^H NMR (400 MHz, DMSO-d_6_) *δ* 8.97 (s, 1H), 8.61 (s, 1H), 8.44 (dd, J = 31.2, 7.9 Hz, 1H), 8.13 (d, J = 23.6 Hz, 4H), 7.81 (d, J = 7.3 Hz, 1H), 7.44 (dd, J = 18.6, 7.1 Hz, 4H), 7.34 (d, J = 21.7 Hz, 5H), 7.24 (dd, J = 15.7, 8.1 Hz, 2H), 6.07 (brs, 1H), 4.90 (s, 1H), 4.53 (d, J = 8.5 Hz, 1H), 4.43 (s, 1H), 4.29 – 4.18 (m, 5H), 3.97 (s, 2H), 3.59 (s, 4H), 3.24 (d, J = 6.6 Hz, 4H), 3.10 – 3.04 (m, 2H), 2.44 (s, 3H), 2.03 (s, 2H), 1.92 (s, 2H), 1.74 (m, 2H), 1.46 (s, 1H), 1.41 – 1.32 (m, 4H), 1.25 (d, J = 5.9 Hz, 3H), 1.17 (s, 2H), 0.95 (s, 9H). ^13^C NMR (100 MHz, DMSO-d_6_) *δ* 171.0, 169.9, 168.9, 165.9, 163.6, 162.7, 157.6, 151.9, 150.5, 148.7, 148.1, 145.1, 142.2, 138.4, 130.5 (q, J = 209 Hz), 130.4, 130.3, 129.2, 127.8, 126.8, 122.7, 119.4, 116.6, 115.5, 69.2, 58.9, 56.7, 53.6, 53.1, 48.3, 48.1, 46.0, 42.0, 41.6, 38.2, 37.2, 36.2, 35.8, 32.2, 31.2, 30.2, 28.2, 27.9, 26.8, 22.8, 17.9, 16.4, 12.9, 9.0. MS (ESI): m/z 1220.5 [M+1]^+^. HRMS (ESI): m/z Calcd for (C_66_H_73_F_3_N_11_O_7_S) ([M+H]^+^): 1220.5367; found: 1220.5315.

(2S,4R)-1-((S)-2-(5-(4-((4-((3-(4-amino-5-(1-(2-(3-(trifluoromethoxy)phenyl)acetyl)indolin-5-yl)-7H-pyrrolo[2,3-d]pyrimidin-7-yl)propyl)carbamoyl)phenyl)ethynyl)piperidin-1-yl)pentanamido)-3,3-dimethylbutanoyl)-4-hydroxy-N-((S)-1-(4-(4-methylthiazol-5-yl)phenyl)ethyl)pyrrolidine-2-carboxamide (**18i**): ^1^H NMR (400 MHz, DMSO-d_6_) *δ* 8.95 (s, 1H), 8.55 (s, 1H), 8.34 (d, J = 7.5 Hz, 1H), 8.07 (d, J = 8.3 Hz, 2H), 7.77 (d, J = 7.7 Hz, 3H), 7.45 – 7.38 (m, 5H), 7.34 (d, J = 8.8 Hz, 4H), 7.27 (s, 2H), 7.21 (dd, J = 16.3, 8.1 Hz, 2H), 6.05 (brs, 1H), 4.87 (d, J = 7.1 Hz, 1H), 4.48 (d, J = 9.1 Hz, 1H), 4.38 (t, J = 7.8 Hz, 1H), 4.25 – 4.15 (m, 5H), 3.93 (s, 2H), 3.56 (s, 4H), 3.21 (d, J = 7.5 Hz, 4H), 2.66 (d, J = 21.9 Hz, 2H), 2.41 (s, 3H), 2.28 (s, 2H), 2.12 (s, 2H), 1.99 (d, J = 6.8 Hz, 2H), 1.83 (s, 2H), 1.75 (s, 1H), 1.59 (d, J = 9.0 Hz, 2H), 1.41 (m, 4H), 1.33 (d, J = 6.7 Hz, 3H), 1.16 (m, 2H), 0.90 (s, 9H). ^13^C NMR (100 MHz, DMSO-d_6_) *δ* 172.4, 171.0, 170.0, 168.9, 165.9, 151.9, 150.5, 148.2, 142.2, 138.4, 130.5 (q, J = 212 Hz), 130.4, 130.1, 129.2, 127.8, 126.8, 125.3, 123.7, 122.7, 119.4, 95.6, 69.2, 59.0, 58.0, 56.8, 48.3, 48.1, 41.7, 38.2, 35.6, 35.1, 31.6, 30.2, 27.9, 26.9, 26.1, 23.8, 22.8, 16.4KK. MS (ESI): m/z 1248.5 [M+1]^+^. HRMS (ESI): m/z Calcd for (C_68_H_77_F_3_N_11_O_7_S) ([M+H]^+^): 1248.5680; found: 1248.5636.

N-(3-(4-amino-5-(1-(2-(3-(trifluoromethoxy)phenyl)acetyl)indolin-5-yl)-7H-pyrrolo[2,3-d]pyrimidin-7-yl)propyl)-6-((2-(3-(((S)-1-((2S,4R)-4-hydroxy-2-(((S)-1-(4-(4-methylthiazol-5-yl)phenyl)ethyl)carbamoyl)pyrrolidin-1-yl)-3,3-dimethyl-1-oxobutan-2-yl)amino)-3-oxopropyl)-2-azaspiro[3.3]heptan-6-yl)amino)nicotinamide (**18j**): ^1^H NMR (400 MHz, DMSO-d_6_) *δ* 8.98 (s, 1H), 8.59 (s, 1H), 8.52 (s, 1H), 8.41 (t, J = 10.0 Hz, 2H), 8.13 (s, 1H), 8.11 – 8.04 (m, 2H), 7.48 (t, J = 7.9 Hz, 1H), 7.42 (d, J = 7.1 Hz, 2H), 7.36 (d, J = 8.0 Hz, 3H), 7.31 (s, 3H), 7.27 (d, J = 8.2 Hz, 1H), 7.22 (d, J = 8.3 Hz, 1H), 6.82 (d, J = 8.5 Hz, 1H), 6.07 (brs, 1H), 5.13 – 5.03 (m, 1H), 4.96 – 4.83 (m, 1H), 4.48 (d, J = 9.0 Hz, 1H), 4.42 (t, J = 8.1 Hz, 1H), 4.26 (s, 1H), 4.23 (d, J = 7.4 Hz, 4H), 3.97 (s, 2H), 3.60 (s, 2H), 3.24 (d, J = 7.0 Hz, 4H), 3.17 (s, 2H), 3.12 – 3.06 (m, 2H), 2.60 (d, J = 5.9 Hz, 2H), 2.13 (d, J = 7.4 Hz, 4H), 2.02 (d, J = 6.8 Hz, 3H), 1.78 (s, 1H), 1.36 (d, J = 6.7 Hz, 3H), 0.95 (s, 9H). ^13^C NMR (100 MHz, DMSO-d_6_) *δ* 171.1, 171.0, 169.9, 168.9, 164.9, 164.4, 157.6, 151.9, 150.5, 148.6, 148.1, 147.4, 145.1, 142.1, 138.6, 138.4, 130.5 (q, J = 208 Hz), 130.4, 130.0, 129.2, 126.7, 125.3, 124.2, 123.7, 122.6, 119.4, 116.6, 115.4, 110.4, 100.3, 69.1, 66.7, 65.8, 65.2, 58.9, 56.8, 56.7, 55.2, 48.2, 48.1, 42.2, 41.7, 40.8, 38.1, 37.1, 35.6, 33.7, 32.1, 30.1, 27.9, 26.8, 22.8, 16.4. MS (ESI): m/z 1225.5 [M+1]^+^. HRMS (ESI): m/z Calcd for (C_64_H_72_F_3_N_12_O_8_S) ([M+H]^+^): 1225.5269; found: 1225.5244.

N-(3-(4-amino-5-(1-(2-(3-(trifluoromethoxy)phenyl)acetyl)indolin-5-yl)-7H-pyrrolo[2,3-d]pyrimidin-7-yl)propyl)-6-((2-(5-(((S)-1-((2S,4R)-4-hydroxy-2-(((S)-1-(4-(4-methylthiazol-5-yl)phenyl)ethyl)carbamoyl)pyrrolidin-1-yl)-3,3-dimethyl-1-oxobutan-2-yl)amino)-5-oxopentyl)-2-azaspiro[3.3]heptan-6-yl)amino)nicotinamide (**18k**): ^1^H NMR (400 MHz, DMSO-d_6_) *δ* 8.94 (s, 1H), 8.55 (s, 1H), 8.49 (s, 1H), 8.35 (d, J = 7.4 Hz, 1H), 8.09 (s, 1H), 8.03 (d, J = 8.8 Hz, 1H), 7.76 (d, J = 9.1 Hz, 1H), 7.41 (dd, J = 14.9, 7.8 Hz, 3H), 7.33 (d, J = 11.5 Hz, 4H), 7.27 (s, 2H), 7.23 (d, J = 7.7 Hz, 1H), 7.18 (d, J = 7.9 Hz, 1H), 6.78 (d, J = 8.7 Hz, 1H), 6.03 (brs, 1H), 5.08 – 4.99 (m, 1H), 4.93 – 4.80 (m, 1H), 4.47 (d, J = 9.1 Hz, 1H), 4.38 (t, J = 8.1 Hz, 1H), 4.23 (s, 2H), 4.19 (d, J = 7.6 Hz, 3H), 3.93 (s, 2H), 3.56 (s, 2H), 3.20 (s, 2H), 3.13 (s, 2H), 3.07 (s, 2H), 2.99 (s, 2H), 2.54 (s, 2H), 2.41 (s, 3H), 2.23 (s, 2H), 2.09 (d, J = 8.4 Hz, 2H), 1.98 (d, J = 6.7 Hz, 2H), 1.97 – 1.92 (m, 1H), 1.75 (s, 1H), 1.41 (d, J = 7.0 Hz, 2H), 1.33 (d, J = 6.7 Hz, 3H), 1.17 (d, J = 14.7 Hz, 4H), 0.89 (s, 9H). ^13^C NMR (100 MHz, DMSO-d_6_) *δ* 172.3, 171.0, 170.0, 168.8, 164.9, 164.4, 157.6, 151.9, 150.5, 148.1, 147.4, 145.1, 142.2, 138.6, 138.4, 130.5 (q, J = 210 Hz), 130.4, 130.1, 129.3, 127.5, 126.8, 125.3, 124.2, 123.7, 122.7, 119.4, 116.5, 110.4, 100.3, 69.1, 66.7, 66.3, 65.7, 59.3, 58.9, 56.7, 48.2, 42.1, 41.7, 40.9, 38.1, 37.1, 35.6, 35.2, 32.1, 30.1, 29.4, 27.9, 27.3, 26.8, 23.7, 22.8, 16.4. MS (ESI): m/z 1253.5 [M+1]^+^. HRMS (ESI): m/z Calcd for (C_66_H_76_F_3_N_12_O_8_S) ([M+H]^+^): 1253.5582; found: 1253.5557.

5-((4-((3-(4-amino-5-(1-(2-(3-(trifluoromethoxy)phenyl)acetyl)indolin-5-yl)-7H-pyrrolo[2,3-d]pyrimidin-7-yl)propyl)carbamoyl)phenyl)ethynyl)-N-((S)-1-((2S,4R)-4-hydroxy-2-(((S)-1-(4-(4-methylthiazol-5-yl)phenyl)ethyl)carbamoyl)pyrrolidin-1-yl)-3,3-dimethyl-1-oxobutan-2-yl)picolinamide (**18l**): ^1^H NMR (400 MHz, DMSO-d_6_) *δ* 8.95 (s, 1H), 8.85 (s, 1H), 8.64 (s, 1H), 8.51 – 8.43 (m, 2H), 8.19 (d, J = 8.1 Hz, 1H), 8.11 (s, 1H), 8.08 (d, J = 7.9 Hz, 2H), 7.88 (d, J = 8.0 Hz, 2H), 7.70 (d, J = 8.0 Hz, 2H), 7.43 (dd, J = 16.6, 8.1 Hz, 3H), 7.35 (d, J = 8.2 Hz, 3H), 7.29 (d, J = 13.5 Hz, 3H), 7.25 – 7.17 (m, 2H), 6.06 (brs, 1H), 5.11 (s, 1H), 4.94 – 4.84 (m, 1H), 4.68 (d, J = 9.5 Hz, 1H), 4.43 (t, J = 8.2 Hz, 1H), 4.27 (s, 1H), 4.20 (d, J = 7.1 Hz, 3H), 3.93 (s, 2H), 3.61 (s, 2H), 3.28 (s, 3H), 3.26 – 3.17 (m, 4H), 2.42 (s, 3H), 2.02 (s, 3H), 1.73 (d, J = 11.8 Hz, 1H), 1.36 (d, J = 6.8 Hz, 3H). ^13^C NMR (100 MHz, DMSO-d_6_) *δ* 170.8, 169.3, 168.8, 165.7, 162.4, 157.6, 152.0, 151.9, 151.1, 150.5, 148.6, 148.4, 148.1, 145.1, 142.1, 141.0, 138.4, 135.4, 132.0, 130.5 (q, J = 210 Hz), 130.4, 130.3, 130.1, 129.3, 128.0, 127.5, 126.7, 125.3, 124.3, 123.7, 122.7, 122.4, 122.1, 119.4, 116.6, 115.4, 100.3, 94.2, 87.8, 69.2, 59.0, 57.0, 48.2, 42.1, 41.6, 38.1, 37.3, 36.7, 30.1, 27.9, 26.7, 22.9, 16.4. MS

(ESI): m/z 1186.4 [M+1]^+^. HRMS (ESI): m/z Calcd for (C_64_H_63_F_3_N_11_O_7_S) ([M+H]^+^): 1186.4585; found: 1186.4562.

(2S,4R)-1-((S)-2-(3-(5-((4-((3-(4-amino-5-(1-(2-(3-(trifluoromethoxy)phenyl)acetyl)indolin-5-yl)-7H-pyrrolo[2,3-d]pyrimidin-7-yl)propyl)carbamoyl)phenyl)ethynyl)pyridin-2-yl)propanamido)-3,3-dimethylbutanoyl)-4-hydroxy-N-((S)-1-(4-(4-methylthiazol-5-yl)phenyl)ethyl)pyrrolidine-2-carboxamide (**18m**): ^1^H NMR (400 MHz, DMSO-d_6_) *δ* 8.98 (s, 1H), 8.66 (d, J = 6.6 Hz, 2H), 8.38 (d, J = 7.7 Hz, 1H), 8.14 (s, 1H), 8.11 (d, J = 8.2 Hz, 1H), 7.93 (d, J = 9.5 Hz, 1H), 7.88 (s, 2H), 7.67 (d, J = 8.0 Hz, 2H), 7.48 (t, J = 7.9 Hz, 1H), 7.43 (d, J = 8.1 Hz, 2H), 7.38 (d, J = 6.3 Hz, 4H), 7.33 (d, J = 11.0 Hz, 4H), 7.29 – 7.21 (m, 2H), 6.08 (brs, 1H), 4.91 (t, J = 7.0 Hz, 1H), 4.52 (d, J = 9.4 Hz, 1H), 4.43 (t, J = 8.1 Hz, 1H), 4.26 (s, 2H), 4.24 (s, 2H), 3.97 (s, 2H), 3.60 (s, 2H), 3.27 (s, 2H), 3.23 (d, J = 8.0 Hz, 2H), 3.00 (d, J = 7.7 Hz, 2H), 2.77 – 2.65 (m, 1H), 2.58 (d, J = 6.8 Hz, 1H), 2.45 (s, 3H), 2.05 (m, 3H), 1.79 (s, 1H), 1.37 (d, J = 6.8 Hz, 3H), 0.91 (s, 9H). ^13^C NMR (100 MHz, DMSO-d_6_) *δ* 171.4, 171.0, 169.9, 168.9, 165.8, 161.3, 157.6, 151.9, 151.4 150.5, 148.1, 145.0, 142.1, 139.2, 138.4, 134.9, 131.76 (s), 130.5 (q, J = 209 Hz), 130.4, 130.3, 130.1, 129.2, 127.9, 126.8, 125.3, 124.9, 123.7, 123.0, 122.6, 119.4, 116.8, 116.5, 115.4, 100.3, 91.5, 88.7, 69.2, 58.9, 56.9, 48.3, 48.1, 42.2, 41.6, 38.1, 37.3, 35.6, 34.4, 33.7, 30.1, 27.9, 26.8, 22.8, 16.4. MS (ESI): m/z 1214.5 [M+1]^+^. HRMS (ESI): m/z Calcd for (C_66_H_67_F_3_N_11_O_7_S) ([M+H]^+^): 1214.4898; found: 1214.4882.

(2S,4R)-1-((S)-2-(3-(5-((4-(3-((3-(4-amino-5-(1-(2-(3-(trifluoromethoxy)phenyl)acetyl)indolin-5-yl)-7H-pyrrolo[2,3-d]pyrimidin-7-yl)propyl)amino)-3-oxopropyl)phenyl)ethynyl)pyridin-2-yl)propanamido)-3,3-dimethylbutanoyl)-4-hydroxy-N-((S)-1-(4-(4-methylthiazol-5-yl)phenyl)ethyl)pyrrolidine-2-carboxamide (**18n**): ^1^H NMR (400 MHz, DMSO-d_6_) *δ* 8.95 (s, 1H), 8.56 (s, 1H), 8.36 (d, J = 7.5 Hz, 1H), 8.10 (s, 1H), 8.08 (d, J = 8.3 Hz, 1H), 7.90 (d, J = 7.7 Hz, 2H), 7.77 (d, J = 7.9 Hz, 1H), 7.43 (d, J = 7.5 Hz, 4H), 7.39 (s, 1H), 7.35 (s, 1H), 7.32 (d, J = 8.4 Hz, 1H), 7.28 (s, 4H), 7.21 (d, J = 21.8 Hz, 7H), 6.03 (brs, 1H), 5.10 (s, 1H), 4.94 – 4.80 (m, 1H), 4.48 (d, J = 9.0 Hz, 1H), 4.40 (t, J = 7.9 Hz, 1H), 4.25 (s, 1H), 4.20 (t, J = 8.1 Hz, 2H), 4.06 (s, 2H), 3.93 (s, 3H), 3.57 (s, 2H), 3.18 (t, J = 7.8 Hz, 2H), 2.99 (s, 2H), 2.93 (d, J = 6.7 Hz, 2H), 2.81 (d, J = 6.9 Hz, 2H), 2.42 (s, 3H), 2.36 (d, J = 7.0 Hz, 2H), 2.03 – 1.93 (m, 1H), 1.86 – 1.81 (m, 2H), 1.76 (d, J = 4.9 Hz, 1H), 1.33 (s, 3H), 0.87 (s, 9H). ^13^C NMR (100 MHz, DMSO-d_6_) *δ* 171.5, 171.0, 169.9, 168.8, 160.9, 157.6, 151.9, 151.8, 151.2, 150.4, 148.6, 148.2, 145.1, 143.0, 142.1, 139.0, 138.4, 133.2, 131.8, 130.5 (q, J = 210 Hz), 130.4, 130.1, 129.2, 129.1, 126.8, 125.3, 123.6, 122.9, 122.6, 119.8, 119.4, 117.2, 116.6, 115.4, 100.4, 92.2, 86.3, 69.2, 58.9, 56.9, 56.7, 48.3, 48.1, 42.0, 41.7 38.1, 36.9, 36.4, 35.6, 35.3, 34.4, 33.7, 33.1, 31.7, 31.3, 30.2, 27.9, 26.8, 22.8, 16.4. MS (ESI): m/z 1242.5 [M+1]^+^. HRMS (ESI): m/z Calcd for (C_68_H_71_F_3_N_11_O_7_S) ([M+H]^+^): 1242.5211; found: 1242.5201.

Compound **19** was prepared according to the procedure described above for compound **18**.

N1-(3-(4-amino-5-(1-(2-(3-(trifluoromethoxy)phenyl)acetyl)indolin-5-yl)-7H-pyrrolo[2,3-d]pyrimidin-7-yl)propyl)-N12-((S)-1-((2S,4R)-4-hydroxy-2-(((S)-3-(methylamino)-1-(4-(4-methylthiazol-5-yl)phenyl)-3-oxopropyl)carbamoyl)pyrrolidin-1-yl)-3,3-dimethyl-1-oxobutan-2-yl)dodecanediamide (**19a**): ^1^H NMR (400 MHz, DMSO-d_6_) *δ* 8.94 (s, 1H), 8.52 (d, J = 7.9 Hz, 1H), 8.07 (d, J = 9.3 Hz, 2H), 7.81 (s, 1H), 7.74 (s, 1H), 7.44 (t, J = 8.1 Hz, 1H), 7.37 (d, J = 6.7 Hz, 1H), 7.35 (s, 1H), 7.30 (d, J = 13.5 Hz, 6H), 7.21 (dd, J = 13.8, 7.7 Hz, 3H), 6.03 (brs, 1H), 5.12 (d, J = 7.3 Hz, 1H), 4.53 (d, J = 9.0 Hz, 1H), 4.40 (t, J = 8.2 Hz, 1H), 4.26 – 4.16 (m, 3H), 4.11 (s, 2H), 3.93 (s, 2H), 3.54 (d, J = 10.1 Hz, 2H), 3.19 (s, 2H), 2.99 (d, J = 6.0 Hz, 2H), 2.84 (s, 1H), 2.53 (d, J = 7.6 Hz, 1H), 2.40 (s, 3H), 2.29 (s, 1H), 2.00 (t, J = 7.3 Hz, 2H), 1.85 (s, 2H), 1.72 (s, 1H), 1.42 (s, 2H), 1.33 (d, J = 5.0 Hz, 2H), 1.23 – 1.06 (m, 16H),0.96 (s, 9H), 0.89 (s, 2H). ^13^C NMR (100 MHz, DMSO-d_6_) *δ* 172.6, 172.5, 171.1, 170.1, 169.9, 168.9, 151.9, 148.6, 148.1, 142.8, 138.3, 130.5 (q, J = 196 Hz), 130.4, 130.3, 129.0, 127.6, 125.3, 122.6, 119.4, 116.7, 99.7, 69.1, 59.1, 56.7, 50.1, 48.2, 42.5, 42.2, 41.7, 38.1, 36.2, 35.9, 35.5, 35.3, 30.2, 29.3, 29.2, 29.1, 29.0, 27.9, 26.8, 25.9, 25.7, 16.4. MS (ESI): m/z 1206.6 [M+1]^+^. HRMS (ESI): m/z Calcd for (C_63_H_79_F_3_N_11_O_8_S) ([M+H]^+^): 1206.5786; found: 1206.5778.

(2S,4R)-N-((S)-3-((11-((3-(4-amino-5-(1-(2-(3-(trifluoromethoxy)phenyl)acetyl)indolin-5-yl)-7H-pyrrolo[2,3-d]pyrimidin-7-yl)propyl)amino)-11-oxoundecyl)amino)-1-(4-(4-methylthiazol-5-yl)phenyl)-3-oxopropyl)-1-((S)-2-(1-fluorocyclopropane-1-carboxamido)-3,3-dimethylbutanoyl)-4-hydroxypyrrolidine-2-carboxamide (**19b**): ^1^H NMR (400 MHz, DMSO-d_6_) *δ* 8.97 (s, 1H), 8.56 (d, J = 7.6 Hz, 1H), 8.12 (d, J = 9.2 Hz, 2H), 7.85 (s, 1H), 7.79 (s, 1H), 7.48 (t, J = 7.9 Hz, 1H), 7.40 (d, J = 7.0 Hz, 2H), 7.35 (d, J = 7.6 Hz, 2H), 7.32 (s, 3H), 7.24 (t, J = 9.4 Hz, 3H), 6.08 (brs, 1H), 5.16 (d, J = 9.6 Hz, 1H), 4.58 (d, J = 8.9 Hz, 1H), 4.44 (t, J = 8.2 Hz, 1H), 4.29 – 4.21 (m, 3H), 4.15 (s, 2H), 3.97 (s, 2H), 3.58 (q, J = 11.2 Hz, 2H), 3.23 (t, J = 7.8 Hz, 2H), 3.03 (d, J = 5.7 Hz, 2H), 2.98 (d, J = 6.2 Hz, 1H), 2.86 (d, J = 6.6 Hz, 1H), 2.57 (d, J = 6.9 Hz, 2H), 2.44 (s, 3H), 2.04 (t, J = 6.9 Hz, 3H), 1.93 – 1.84 (m, 2H), 1.74 (s, 1H), 1.46 (s, 2H), 1.39 – 1.31 (m, 2H), 1.17 (m, 14H), 1.03 (m, 2H), 0.97 (s, 9H). ^13^C NMR (100 MHz, DMSO-d_6_) *δ* 172.5, 170.8, 169.2, 168.9, 168.5, 157.6, 151.9, 150.5, 148.6, 148.1, 142.5, 142.2, 138.4, 130.5 (q, J = 206 Hz), 130.4, 130.3, 128.9, 125.3, 123.6, 122.7, 119.4, 116.6, 115.4, 100.3, 79.7, 77.4, 69.2, 59.2, 56.9, 50.4, 48.3, 42.3, 42.1, 41.6, 38.7, 38.1, 36.5, 36.4, 35.9, 30.3, 29.4, 29.2, 29.1, 27.9, 26.7, 26.6, 25.7, 16.4, 13.4 (d, J = 11 Hz), 13.1 (d, J = 10 Hz). MS (ESI): m/z 1250.6 [M+1]^+^. HRMS (ESI): m/z Calcd for (C_65_H_80_F_4_N_11_O_8_S) ([M+H]^+^): 1250.5848; found: 1250.5867.

General Procedures for the preparation of 12-(4-(4-amino-5-(1-(2-(3-(trifluoromethoxy)phenyl)acetyl)indolin-5-yl)-7H-pyrrolo[2,3-d]pyrimidin-7-yl)piperidin-1-yl)-12-oxododecanoic acid (21). To a solution of 20 (50 mg, 0.1 mmol), 12-(tert-butoxy)-12-oxododecanoic acid (24 mg, 0.12 mmol) in DMF (5 mL) was added HATU (57 mg, 0.15 mmol) and DIPEA (43 µL, 0.3 mmol). The mixture was stirred at room temperature for 1 hour and was concentrated and then was purified by silica gel flash column chromatography (DCM/MeOH=20/1) to give desired compound. MS (ESI): m/z 805.5 [M+1]^+^.

The above compound was dissolved in DCM (2 mL) and TFA (0.5 mL). The mixture was stirred at room temperature for 2 hours before it was concentrated under reduced pressure to give compound **21** without further purification. ^1^H NMR (400 MHz, DMSO-*d*_*6*_) *δ* 11.93 (s, 1H), 8.17 (s, 1H), 8.07 (d, *J* = 8.6 Hz, 1H), 7.49 (s, 1H), 7.43 (d, *J* = 7.2 Hz, 1H), 7.30 (s, 3H), 7.26 – 7.17 (m, 2H), 4.82 (m, 1H), 4.21 (t, *J* = 8.3 Hz, 2H), 3.94 (s, 2H), 3.23 – 3.16 (m, 2H), 2.65 (d, *J* = 12.1 Hz, 2H), 2.29 (s, 2H), 2.14 (t, *J* = 7.2 Hz, 2H), 1.98 – 1.83 (m, 2H), 1.44 (d, *J* = 6.1 Hz, 4H), 1.21 (m, 16H). MS (ESI): m/z 749.3 [M+1]^+^.

### General Procedures for the preparation of compound (23)

To a solution of **21** (20 mg, 0.03 mmol) and VHL ligand acid (0.04 mmol) in DMF (5 mL) was added HATU (19 mg, 0.05 mmol) and DIPEA (22 µL, 0.15 mmol). The mixture was stirred at room temperature for 1 hour. The reaction was concentrated and then was purified by reverse phase preparative HPLC (0-20mins: MeOH/H_2_O =10% → 80%; 20-30mins: MeOH/H_2_O =80%) to provide the title compound.

(2S,4R)-1-((S)-2-(10-(4-(4-amino-5-(1-(2-(3-(trifluoromethoxy)phenyl)acetyl)indolin-5-yl)-7H-pyrrolo[2,3-d]pyrimidin-7-yl)piperidin-1-yl)-10-oxodecanamido)-3,3-dimethylbutanoyl)-4-hydroxy-N-((S)-1-(4-(4-methylthiazol-5-yl)phenyl)ethyl)pyrrolidine-2-carboxamide (**23a**). ^1^H NMR (400 MHz, DMSO-d_6_) *δ* 8.95 (s, 1H), 8.35 (d, J = 7.5 Hz, 1H), 8.12 – 8.03 (m, 2H), 7.76 (d, J = 9.0 Hz, 1H), 7.46 – 7.42 (m, 1H), 7.39 (d, J = 5.3 Hz, 3H), 7.34 (d, J = 8.1 Hz, 2H), 7.29 (s, 4H), 7.24 – 7.17 (m, 2H), 6.04 (brs, 1H), 5.06 (s, 1H), 4.87 (d, J = 6.8 Hz, 1H), 4.81 (s, 1H), 4.54 (d, J = 12.4 Hz, 1H), 4.48 (d, J = 9.1 Hz, 1H),4.38 (t, J = 7.7 Hz, 1H), 4.27 – 4.15 (m, 4H), 3.93 (s, 2H), 3.56 (s, 2H), 3.18 (d, J = 7.3 Hz, 2H), 2.41 (s, 3H), 2.30 (s, 2H), 2.24 – 2.16 (m, 1H), 2.08 (d, J = 6.1 Hz, 1H), 1.93 (d, J = 12.1 Hz, 4H), 1.86 – 1.81 (m, 1H), 1.75 (d, J = 4.6 Hz, 1H), 1.46 (s, 4H), 1.33 (d, J = 6.7 Hz, 3H), 1.23 (s, 8H), 0.89 (s, 9H). ^13^C NMR (100 MHz, DMSO-d_6_) *δ* 172.5, 171.0, 170.9, 170.0, 168.8, 157.6, 151.9, 151.7, 150.1, 148.1, 145.1, 142.1, 138.4, 130.5 (q, J = 209 Hz), 130.4, 130.1, 129.2, 126.8, 125.3, 122.6, 120.8, 119.4, 116.5, 115.8, 100.4, 69.2, 58.9, 56.7, 51.5, 48.2, 48.1, 41.6, 38.1, 35.6, 35.3, 32.8, 32.6, 31.9, 29.3, 29.2, 29.1, 27.9, 26.8, 25.8, 25.3, 22.8, 16.4. MS (ESI): m/z 1147.5 [M+1]^+^. HRMS (ESI): m/z Calcd for (C_61_H_74_F_3_N_10_O_7_S) ([M+H]^+^): 1147.5415; found: 1147.5402.

(2S,4R)-1-((S)-2-(12-(4-(4-amino-5-(1-(2-(3-(trifluoromethoxy)phenyl)acetyl)indolin-5-yl)-7H-pyrrolo[2,3-d]pyrimidin-7-yl)piperidin-1-yl)-12-oxododecanamido)-3,3-dimethylbutanoyl)-4-hydroxy-N-(1-(4-(4-methylthiazol-5-yl)phenyl)cyclopropyl)pyrrolidine-2-carboxamide (**23b**): ^1^H NMR (400 MHz, DMSO-d_6_) *δ* 8.92 (s, 1H), 8.78 (s, 1H), 8.27 (s, 1H), 8.11 – 8.03 (m, 2H), 7.87 (d, J = 9.2 Hz, 1H), 7.44 (t, J = 7.8 Hz, 1H), 7.39 (s, 1H), 7.30 – 7.25 (m, 6H), 7.19 (d, J = 9.7 Hz, 2H), 6.04 (brs, 1H), 4.80 (s, 1H), 4.52 (t, J = 11.7 Hz, 2H), 4.35 (t, J = 7.9 Hz, 1H), 4.31 (s, 1H), 4.20 (t, J = 8.0 Hz, 2H), 3.99 (s, 1H), 3.93 (s, 2H), 3.60 (s, 2H), 3.17 (d, J = 7.8 Hz, 2H), 2.66 (t, J = 10.4 Hz, 1H), 2.39 (s, 3H), 2.30 (s, 2H), 2.22 (dd, J = 14.2, 6.8 Hz, 1H), 2.12 – 2.03 (m, 1H), 1.94 (d, J = 9.2 Hz, 1H), 1.85 (s, 2H), 1.46 (d, J = 5.2 Hz, 4H), 1.21 (m, 20H), 0.89 (s, 9H). ^13^C NMR (100 MHz, DMSO-d_6_) *δ* 173.0, 172.5, 170.9, 170.1, 168.8, 157.6, 151.6, 150.1, 148.6, 147.8, 144.1, 142.1, 138.4, 130.5 (q, J = 190 Hz), 130.4, 130.3, 128.7, 128.6, 125.3, 125.2, 122.6, 120.8, 119.4, 116.5, 115.8, 100.4, 69.4, 59.2, 56.8, 56.6, 51.6, 49.0, 48.2, 44.7, 41.6, 40.9, 38.0, 35.7, 35.3, 34.0, 32.7, 32.6, 31.9, 29.4, 29.3, 29.2, 29.1, 27.9, 26.7, 25.8, 25.3, 20.3, 19.4, 16.3. MS (ESI): m/z 1187.6 [M+1]^+^. HRMS (ESI): m/z Calcd for (C_64_H_78_F_3_N_10_O_7_S) ([M+H]^+^): 1187.5728; found: 1187.5713.

12-(4-(4-amino-5-(1-(2-(3-(trifluoromethoxy)phenyl)acetyl)indolin-5-yl)-7H-pyrrolo[2,3-d]pyrimidin-7-yl)piperidin-1-yl)-N-((S)-1-((2S,4R)-4-hydroxy-2-(1-(4-(4-methylthiazol-5-yl)benzyl)-1H-1,2,3-triazol-4-yl)pyrrolidin-1-yl)-3,3-dimethyl-1-oxobutan-2-yl)-12-oxododecanamide (**23c**): ^1^H NMR (400 MHz, DMSO-d_6_) *δ* 8.96 (s, 1H), 8.08 (d, J = 6.7 Hz, 1H), 8.02 (d, J = 24.8 Hz, 1H), 7.90 (s, 1H), 7.71 (d, J = 9.3 Hz, 1H), 7.41 (dd, J = 12.7, 5.2 Hz, 3H), 7.32 – 7.17 (m, 6H), 6.94 (s, 1H), 6.05 (brs, 1H), 5.58 (s, 1H), 5.11 (s, 1H), 4.80 (s, 1H), 4.54 (d, J = 12.9 Hz, 1H), 4.41 (d, J = 9.5 Hz, 2H), 4.24 – 4.16 (m, 2H), 3.93 (s, 3H), 3.70 (s, 1H), 3.56 (d, J = 10.3 Hz, 1H), 3.23 – 3.15 (m, 2H), 3.13 (s, 1H), 2.89 (s, 1H), 2.75 (s, 1H), 2.64 (d, J = 9.7 Hz, 1H), 2.38 (d, J = 11.6 Hz, 3H), 2.29 (s, 2H), 2.19 (td, J = 13.5, 6.8 Hz, 2H), 2.10 – 2.00 (m, 2H), 1.92 (m, 4H), 1.46 (d, J = 5.3 Hz, 4H), 1.19 (m, 16H), 0.92 – 0.78 (m, 2H), 0.68 (s, 9H). ^13^C NMR (100 MHz, DMSO-d_6_) *δ* 172.5, 170.9, 169.8, 168.8, 157.6, 152.2, 151.7, 150.1, 148.7, 148.5, 148.2, 142.1, 138.4, 136.4, 131.5, 130.5 (q, J = 190 Hz), 130.3, 129.6, 128.6, 125.3, 123.8, 122.6, 120.8, 119.4, 116.5, 115.8, 100.4, 69.0, 56.7, 52.5, 51.5, 51.1, 48.2, 41.6, 35.3, 35.2, 32.7, 31.9, 29.4, 29.2, 29.0, 27.9, 26.6, 25.8, 25.3, 25.1, 16.3. MS (ESI): m/z 1185.5 [M+1]^+^. HRMS (ESI): m/z Calcd for (C_63_H_76_F_3_N_12_O_6_S) ([M+H]^+^): 1185.5684; found: 1185.5669.

(2S,4R)-1-(2-(10-(4-(4-amino-5-(1-(2-(3-(trifluoromethoxy)phenyl)acetyl)indolin-5-yl)-7H-pyrrolo[2,3-d]pyrimidin-7-yl)piperidin-1-yl)-10-oxodecanamido)-6-methylbenzoyl)-4-hydroxy-N-((S)-1-(4-(4-methylthiazol-5-yl)phenyl)ethyl)pyrrolidine-2-carboxamide (**23d**): ^1^H NMR (400 MHz, DMSO-d_6_) *δ* 9.29 (s, 1H), 8.93 (s, 1H), 8.90 (d, J = 7.0 Hz, 1H), 8.16 (d, J = 8.5 Hz, 1H), 8.08 (d, J = 7.6 Hz, 2H), 7.42 (d, J = 8.4 Hz, 3H), 7.37 (t, J = 9.3 Hz, 4H), 7.28 (s, 3H), 7.19 (d, J = 8.1 Hz, 3H), 6.90 (s, 1H), 6.05 (brs, 1H), 4.98 – 4.89 (m, 1H), 4.79 (s, 1H), 4.68 (t, J = 8.0 Hz, 1H), 4.52 (s, 2H), 4.20 (d, J = 8.1 Hz, 3H), 4.14 (s, 1H), 3.92 (s, 3H), 3.18 (s, 2H), 3.12 (s, 2H), 2.86 (d, J = 10.3 Hz, 1H), 2.63 (s, 1H), 2.42 (s, 3H), 2.32 (d, J = 6.7 Hz, 4H), 2.22 (s, 2H), 2.15 (s, 3H), 1.84 (m, 8H), 1.43 (m, 4H), 1.41 (d, J = 6.6 Hz, 3H), 1.26 (m, 4H), 1.14 (s, 9H). ^13^C NMR (100 MHz, DMSO-d_6_) *δ* 172.7, 171.7, 170.9, 170.8, 168.8, 166.8, 157.6, 151.9, 151.7, 150.1, 148.6, 144.7, 142.1, 138.4, 135.4, 133.7, 130.5 (q, J = 202 Hz), 130.4, 130.3, 129.3, 126.7, 125.3, 122.6, 120.9, 119.4, 117.8, 116.5, 115.8, 100.4, 68.9, 57.9, 56.5, 51.5, 48.8, 48.2, 44.7, 41.6, 38.5, 36.8, 32.7, 31.9, 29.3, 29.1, 29.0, 27.9, 25.8, 25.3, 22.6, 19.1, 16.4. MS (ESI): m/z 1167.5 [M+1]^+^. HRMS (ESI): m/z Calcd for (C_63_H_70_F_3_N_10_O_7_S) ([M+H]^+^): 1167.5102; found: 1167.5088.

(2S,4R)-1-((S)-2-(12-(4-(4-amino-5-(1-(2-(3-(trifluoromethoxy)phenyl)acetyl)indolin-5-yl)-7H-pyrrolo[2,3-d]pyrimidin-7-yl)piperidin-1-yl)-12-oxododecanamido)propanethioyl)-4-hydroxy-N-((S)-1-(4-(4-methylthiazol-5-yl)phenyl)ethyl)pyrrolidine-2-carboxamide (**23e**): ^1^H NMR (400 MHz, DMSO-d_6_) *δ* 8.99 (s, 1H), 8.46 (s, 1H), 8.40 (d, J = 21.0 Hz, 1H), 8.14 (d, J = 8.3 Hz, 1H), 8.04 (d, J = 7.4 Hz, 1H), 7.82 (s, 1H), 7.48 (t, J = 7.9 Hz, 1H), 7.43 (d, J = 7.7 Hz, 2H), 7.37 (d, J = 10.4 Hz, 4H), 7.31 (s, 1H), 7.27 (d, J = 7.9 Hz, 2H), 4.91 (dd, J = 14.9, 7.1 Hz, 4H), 4.84 – 4.77 (m, 2H), 4.59 (d, J = 12.7 Hz, 1H), 4.36 (s, 1H), 4.25 (t, J = 8.4 Hz, 2H), 3.98 (s, 3H), 3.82 (t, J = 11.2 Hz, 2H), 3.24 (d, J = 8.8 Hz, 2H), 2.45 (s, 3H), 2.34 (d, J = 7.5 Hz, 2H), 2.10 (t, J = 6.8 Hz, 4H), 1.92 (d, J = 5.7 Hz, 2H), 1.46 (d, J = 8.4 Hz, 4H), 1.37 (d, J = 6.8 Hz, 3H), 1.24 (m, 16H), 1.20 (d, J = 6.7 Hz, 3H). ^13^C NMR (100 MHz, DMSO-d_6_) *δ* 203.6, 171.8, 171.0, 169.22, 169.1, 162.7, 151.9, 148.1, 145.0, 142.8, 138.3, 133.6, 131.52 (s), 130.5 (q, J = 212 Hz), 130.4, 130.1, 129.2, 127.9, 126.8, 125.3, 122.6, 119.5, 118.9, 116.8, 98.9, 68.7, 65.0, 58.9, 52.6, 51.6, 48.3, 48.0, 41.6, 38.0, 36.2, 35.4, 32.7, 29.3, 29.2, 29.1, 27.9, 25.5, 25.3, 22.8, 20.6, 16.4. MS (ESI): m/z 1149.5 [M+1]^+^. HRMS (ESI): m/z Calcd for (C_60_H_72_F_3_N_10_O_6_S_2_) ([M+H]^+^): 1149.5030; found: 1149.5018.

### General Procedures for the preparation of compound (24)

To a solution of **20** (50 mg, 0.1 mmol), carboxylic acid (0.12 mmol) in DMF (5 mL) was added HATU (57 mg, 0.15 mmol) and DIPEA (43 µL, 0.3 mmol). The mixture was stirred at room temperature for 1 hour and was concentrated and then was purified by silica gel flash column chromatography (DCM/MeOH=20/1) to give the title compound. The compound was dissolved in DCM (2 mL) and TFA (0.5 mL). The mixture was stirred at room temperature for 2 hours before it was concentrated under reduced pressure to give compound **22** without further purification.

To a solution of **22** (0.04 mmol) and VH101 acid (0.05 mmol) in DMF (5 mL) was added HATU (28 mg, 0.07 mmol) and DIPEA (34 µL, 0.24 mmol). The mixture was stirred at room temperature for 1 hour. The reaction was concentrated and then was purified by reverse phase preparative HPLC (0-20mins: MeOH/H_2_O =10% →80%; 20-30mins: MeOH/H_2_O =80%) to provide compound **24**.

(2S,4R)-N-((S)-3-(6-(5-((4-(4-(4-amino-5-(1-(2-(3-(trifluoromethoxy)phenyl)acetyl)indolin-5-yl)-7H-pyrrolo[2,3-d]pyrimidin-7-yl)piperidine-1-carbonyl)phenyl)ethynyl)pyridin-2-yl)-2,6-diazaspiro[3.3]heptan-2-yl)-1-(4-(4-methylthiazol-5-yl)phenyl)-3-oxopropyl)-1-((S)-2-(1-fluorocyclopropane-1-carboxamido)-3,3-dimethylbutanoyl)-4-hydroxypyrrolidine-2-carboxamide (**24a**): ^1^H NMR (400 MHz, DMSO-d_6_) *δ* 8.97 (s, 1H), 8.55 (d, J = 7.4 Hz, 1H), 8.22 (s, 1H), 8.08 (d, J = 9.2 Hz, 2H), 7.61 (d, J = 8.2 Hz, 1H), 7.56 – 7.50 (m, 3H), 7.47 (s, 2H), 7.42 (d, J = 6.4 Hz, 2H), 7.35 (d, J = 7.0 Hz, 2H), 7.30 (d, J = 12.3 Hz, 3H), 7.22 (s, 3H), 6.31 (d, J = 8.1 Hz, 1H), 6.07 (brs, 1H), 5.12 (d, J = 6.4 Hz, 1H), 4.89 (s, 1H), 4.63 (s, 1H), 4.55 (d, J = 8.8 Hz, 1H), 4.42 (t, J = 7.4 Hz, 1H), 4.23 (d, J = 17.3 Hz, 5H), 4.10 – 3.96 (m, 7H), 3.94 (s, 2H), 3.54 (m, 2H), 3.20 (s, 2H), 3.13 (s, 1H), 2.57 (d, J = 9.1 Hz, 1H), 2.43 (s, 3H), 2.03 (m, 4H), 1.84 (s, 1H), 1.72 (m, 1H), 1.34 (m, 2H), 1.19 (m, 4H), 0.94 (s, 9H). ^13^C NMR (100 MHz, DMSO-d_6_) *δ* 170.8, 169.5, 169.2, 168.9, 168.7, 168.5, 168.3, 159.0, 157.6, 152.0, 151.7, 151.5, 150.1, 148.6, 148.2, 142.8, 142.1, 139.8, 138.4, 136.1, 130.4, 130.4 (q, J = 198 Hz), 130.3, 129.2, 127.6, 127.4, 125.3, 124.2, 122.6, 120.9, 119.4, 116.5, 115.9, 107.3, 105.9, 100.3, 89.6, 89.5, 79.7, 77.4, 69.2, 60.6, 60.4, 59.2, 57.9, 57.0, 51.1, 49.9, 48.3, 41.6, 38.1, 36.5, 32.9, 27.9, 26.6, 16.4, 13.4 (d, J = 10 Hz), 13.2 (d, J = 10 Hz). MS (ESI): m/z 1394.6 [M+1]^+^. HRMS (ESI): m/z Calcd for (C_75_H_76_F_4_N_13_O_8_S) ([M+H]^+^): 1394.5597; found: 1394.5587.

(2S,4R)-N-((S)-3-(4-(5-((4-(4-(4-amino-5-(1-(2-(3-(trifluoromethoxy)phenyl)acetyl)indolin-5-yl)-7H-pyrrolo[2,3-d]pyrimidin-7-yl)piperidine-1-carbonyl)phenyl)ethynyl)pyridin-2-yl)piperazin-1-yl)-1-(4-(4-methylthiazol-5-yl)phenyl)-3-oxopropyl)-1-((S)-2-(1-fluorocyclopropane-1-carboxamido)-3,3-dimethylbutanoyl)-4-hydroxypyrrolidine-2-carboxamide (**LD5097** (**24b)**): ^1^H NMR (400 MHz, DMSO-d_6_) *δ* 8.93 (s, 1H), 8.51 (d, J = 7.7 Hz, 1H), 8.27 (s, 1H), 8.09 (d, J = 11.7 Hz, 2H), 7.64 (d, J = 8.6 Hz, 1H), 7.55 (d, J = 7.7 Hz, 2H), 7.51 (s, 1H), 7.45 (dd, J = 13.8, 7.8 Hz, 3H), 7.39 (s, 4H), 7.30 (d, J = 12.2 Hz, 3H), 7.22 (s, 3H), 6.80 (d, J = 8.4 Hz, 1H), 6.07 (brs, 1H), 5.20 (d, J = 6.9 Hz, 1H), 5.12 (s, 1H), 4.90 (s, 1H), 4.65 (d, J = 8.1 Hz, 1H), 4.53 (d, J = 8.6 Hz, 1H), 4.42 (t, J = 8.0 Hz, 1H), 4.23 (s, 1H), 4.20 (d, J = 7.7 Hz, 2H), 3.94 (s, 2H), 3.59 – 3.52 (m, 4H), 3.42 (dd, J = 24.6, 14.6 Hz, 4H), 3.28 (s, 2H), 3.20 (t, J = 7.7 Hz, 2H), 2.86 (d, J = 5.9 Hz, 2H), 2.36 (s, 3H), 2.02 (s, 2H), 1.86 (s, 1H), 1.73 (s, 1H), 1.32 (m, 2H), 1.17 (d, J = 9.9 Hz, 4H), 0.92 (s, 9H), 0.89 (s, 2H). ^13^C NMR (100 MHz, DMSO-d_6_) *δ* 170.9, 169.2, 168.9, 168.8, 168.5, 168.4, 168.3, 157.9, 157.7, 151.9, 151.8, 151.2, 150.1, 148.6, 148.2, 142.8, 142.1, 140.4, 138.4, 136.1, 130.5 (q, J = 198 Hz), 130.4, 129.2, 127.9, 127.6, 125.3, 124.2, 122.6, 120.9, 119.4, 116.5, 115.9, 107.4, 107.0, 100.3, 89.6, 89.4, 78.5 (d, J = 231 Hz), 69.2, 59.2, 56.9, 51.1, 50.3, 48.2, 45.1, 44.4, 41.6, 38.1, 36.5, 27.9, 26.6, 16.3, 13.4 (d, J = 10 Hz), 13.2 (d, J = 11 Hz). MS (ESI): m/z 1382.6 [M+1]^+^. HRMS (ESI): m/z Calcd for (C_74_H_76_F_4_N_13_O_8_S) ([M+H]^+^): 1382.5597; found: 1382.5590.

N-((S)-1-((2S,4R)-2-((S)-4-(6-(5-(4-(4-(4-amino-5-(1-(2-(3-(trifluoromethoxy)phenyl)acetyl)indolin-5-yl)-7H-pyrrolo[2,3-d]pyrimidin-7-yl)piperidine-1-carbonyl)phenyl)pyrimidin-2-yl)-2,6-diazaspiro[3.3]heptan-2-yl)-2-(4-(4-methylthiazol-5-yl)phenyl)-4-oxobutanoyl)-4-hydroxypyrrolidin-1-yl)-3,3-dimethyl-1-oxobutan-2-yl)-1-fluorocyclopropane-1-carboxamide (**24c**): ^1^H NMR (400 MHz, DMSO-d_6_) *δ* 8.97 (s, 1H), 8.69 (s, 2H), 8.53 (d, J = 7.2 Hz, 1H), 8.08 (d, J = 11.2 Hz, 2H), 7.68 (d, J = 7.1 Hz, 2H), 7.51 (d, J = 6.7 Hz, 2H), 7.45 – 7.39 (m, 3H), 7.35 (d, J = 7.3 Hz, 2H), 7.30 (d, J = 12.0 Hz, 3H), 7.23 (d, J = 7.0 Hz, 3H), 6.07 (brs, 1H), 5.13 (s, 1H), 5.10 (s, 1H), 4.89 (s, 1H), 4.54 (d, J = 8.7 Hz, 1H), 4.42 (t, J = 7.7 Hz, 1H), 4.24 (s, 2H), 4.09 (m, 8H), 3.94 (s, 3H), 3.53 (d, J = 10.6 Hz, 2H), 3.21 (d, J = 7.7 Hz, 2H), 3.13 (s, 1H), 2.57 (d, J = 14.3 Hz, 1H), 2.43 (s, 3H), 2.03 (s, 3H), 1.85 (s, 2H), 1.71 (s, 1H), 1.32 (m, 2H), 1.19 (m, 4H), 0.94 (s, 9H). ^13^C NMR (100 MHz, DMSO-d_6_) *δ* 170.8, 169.5, 169.2, 169.1, 168.9, 162.1, 157.6, 156.3, 152.0, 151.7, 150.1, 148.6, 148.2, 142.8, 142.1, 138.4, 136.5, 135.2, 130.5 (q, J = 200 Hz), 130.4, 130.3, 129.1, 128.0, 127.5, 125.8, 125.3, 122.6, 122.3, 120.9, 119.4, 116.5, 115.9, 100.3, 69.3, 60.1, 59.2, 57.9, 57.0, 51.2, 49.9, 48.3, 41.6, 38.1, 36.5, 32.7, 27.9, 26.6, 16.4, 13.3 (d, J = 10 Hz), 13.2 (d, J = 11 Hz). MS (ESI): m/z 1356.5 [M+1]^+^. HRMS (ESI): m/z Calcd for (C_72_H_74_F_4_N_13_O_8_S) ([M+H]^+^): 1356.5440; found: 1356.5497.

(2R,4S)-N-((S)-3-(4-(5-((4-(4-(4-amino-5-(1-(2-(3-(trifluoromethoxy)phenyl)acetyl)indolin-5-yl)-7H-pyrrolo[2,3-d]pyrimidin-7-yl)piperidine-1-carbonyl)phenyl)ethynyl)pyridin-2-yl)piperazin-1-yl)-1-(4-(4-methylthiazol-5-yl)phenyl)-3-oxopropyl)-1-((S)-2-(1-fluorocyclopropane-1-carboxamido)-3,3-dimethylbutanoyl)-4-hydroxypyrrolidine-2-carboxamide

**(LD5097-NC**): ^1^H NMR (400 MHz, DMSO-d_6_) *δ* 8.94 (s, 1H), 8.43 (s, 1H), 8.28 (s, 1H), 8.08 (d, J = 9.5 Hz, 2H), 7.66 (d, J = 8.6 Hz, 1H), 7.55 (d, J = 7.2 Hz, 2H), 7.51 (s, 1H), 7.44 (t, J = 6.9 Hz, 5H), 7.39 (d, J = 7.7 Hz, 2H), 7.30 (d, J = 13.0 Hz, 3H), 7.23 (s, 2H), 6.83 (d, J = 8.1 Hz, 1H), 6.06 (brs, 1H), 5.21 (d, J = 6.5 Hz, 1H), 4.89 (m, 1H), 4.56 (d, J = 8.8 Hz, 1H), 4.40 – 4.29 (m, 2H), 4.20 (d, J = 8.2 Hz, 2H), 3.94 (s, 2H), 3.61 (s, 2H), 3.44 (m, 8H), 3.21 (d, J = 7.7 Hz, 2H), 2.81 (s, 2H), 2.37 (s, 3H), 2.03 (s, 2H), 1.95 (s, 2H), 1.36 – 1.24 (m, 2H), 1.19 (s, 2H), 1.15 (d, J = 6.7 Hz, 2H), 0.92 (s, 9H). ^13^C NMR (100 MHz, DMSO-d_6_) *δ* 171.1, 169.0, 168.8, 168.6, 168.4, 157.8, 157.7, 152.0, 151.8, 151.1, 150.1, 148.6, 148.2, 142.7, 142.1, 140.4, 138.4, 136.1, 130.5 (q, J = 200 Hz), 130.4, 129.1, 127.6, 125.3, 124.2, 122.7, 120.9, 119.4, 116.5, 115.9, 107.4, 107.0, 100.3, 89.6, 89.4, 79.5, 77.2, 68.9, 59.1, 56.7, 55.8, 51.1, 49.8, 48.3, 45.1, 44.5, 44.3, 41.6, 41.1, 38.1, 36.1, 29.4, 27.9, 26.8, 26.5, 16.3, 13.4 (d, J = 10 Hz), 13.2 (d, J = 11 Hz). MS (ESI): m/z 1382.6 [M+1]^+^. HRMS (ESI): m/z Calcd for (C_74_H_76_F_4_N_13_O_8_S) ([M+H]^+^): 1382.5597; found: 1382.5590.

### nLuc-RIPK1 degradation evaluation in Jurkat cells

Jurkat cells expressing nLuc-RIPK1 were seeded into 96-well white plates (Corning) at 1□× □10^4^ cells/well in 99□μL of DMEM/FCS, followed by overnight incubation. Cells were then treated with either DMSO or degrader compound as indicated. After 24 hours, 100 uL of the Nano-Glo Luciferase assay system (Promega) was added into each well. Plates were incubated for 10□min at room temperature. Luminescence was then measured on a BioTek Synergy H1 plate.

### Metabolites identification in mouse liver S9 fraction

Mix the 5 uL of MgCl_2_ (50 mM), 3 uL of S9 fraction (20 mg/ml) and 37 uL PSB with 5 uL of NADPH (10 mM) or 5 μL of ultra-pure water. The final concentration of S9 fraction was 1 mg/mL. The mixture was pre-warmed at 37°C for 10 minutes. 30 mins after 1 μL of 100 μM test compound solutions, the reaction was stopped by the addition of 10 volumes of cold acetonitrile. Samples were centrifuged at 20,000 g for 20 minutes. Aliquot of 100 µL of the supernatant was mixed with 100 µL of ultra-pure water and then injected into Orbitrap Fusion™ Lumos™ Tribrid™ Mass Spectrometer. Metabolites were then analyzed using Compound Discoverer 3.1.

### Metabolic stability in mouse liver S9 fraction

Mix the 25.5 uL of MgCl_2_ (50 mM), 15 uL of S9 fraction (20 mg/ml) and 186.5 uL PSB with 25.5 uL of NADPH (10 mM) or 25.5 μL of ultra-pure water. The final concentration of S9 fraction was 1 mg/mL. The mixture was pre-warmed at 37°C for 10 minutes.

The reaction was started with the addition of 2.55 μL of 100 μM control compound or test compound solutions. Verapamil was used as positive control in this study. The final concentration of control compound and test compounds were 1 μM. The incubation solution was incubated in water batch at 37°C.

Aliquots of 25 µL were taken from the reaction solution at 0, 15, 30, 45, 60 and 90 minutes. The reaction was stopped by the addition of 10 volumes of cold acetonitrile with IS (100 nM imipramine). Samples were centrifuged at 12,000 g for 10 minutes. Aliquot of 100 µL of the supernatant was mixed with 100 µL of ultra-pure water and then used for LC-MS/MS analysis. All calculations were carried out using GraphPad Prism 10.00.

### Metabolic stability in human hepatocytes

Pipette 198 μL of hepatocytes into each well of a 96-well non-coated plate with 5×10^5^ cells/mL. Added 2 μL of the 100 μM test compound or positive control into respective wells of the 96-well non-coated plate to start the reaction. Transfer well contents in 25 μL aliquots at time points of 0.5, 15, 30, 60, 90 and 120 minutes. The aliquots were then mixed with 6 volumes (150 μL) of acetonitrile containing with internal standard, IS (100 nM alprazolam, 200 nM caffeine and 100 nM tolbutamide) to terminate the reaction. Vortex for 5 minutes. Samples were centrifuges for 45 minutes at 3,220 g. Aliquot of 100 µL of the supernatant was diluted by 100 µL ultrapure water, and the mixture was used for LC/MS/MS analysis. All incubations were performed in duplicate.

### Metabolic stability in human liver microsomes

Mix 168 µL of phosphate buffer, 10 µL of human microsomes (20 mg/mL), 20 µL of NADPH (10 mM), or 20 µL of ultra-pure water. The final concentration of the human liver microsome fraction was 1 mg/mL. The mixture was pre-warmed at 37°C for 10 minutes. The reaction was started with the addition of 2 µL of 100 µM control compound or test compounds. Transfer 30 µL of the reaction solution at 0, 20, 40, 60, 120, and 240 minutes and mix with 120 µL of methanol containing 200 nM imipramine as an internal reference. Samples were centrifuged at 20,000 g for 20 minutes. An aliquot of 100 µL of the supernatant was mixed with 100 µL of ultra-pure water and then injected into an Orbitrap Fusion™ Lumos™ Tribrid™ Mass Spectrometer. All incubations were performed in triplicates.

### Pharmacokinetics assay

For IV injection, drug (Formulation: 0.4 mg/mL in cosolvent which content 10% of DMSO, 30% of PEG400, 5% of Tween80 in PBS; Dosing: 1 mg/kg) were injected into C57BL/6J male mice (n = 3). 10 µL of blood was collected via the tail vein at 2, 5, 15, 30 min, and 1, 2, 4, 8, 24 and 48 h.

The desired serial concentrations of working solutions were achieved by diluting stock solution of analyte with 100% acetonitrile solution. 5 µL of working solutions (1, 2, 4, 10, 20, 100, 200, 1000, 2000 ng/mL) were added to 10 μL of the blank C57BL/6J mouse plasma to achieve calibration standards of 0.5∼1000 ng/mL (0.5, 1, 2, 5, 10, 50, 100, 500, 1000 ng/mL) in a total volume of 15 μL. Four quality control samples at 1 ng/mL, 2 ng/mL, 50 ng/mL and 800 ng/mL for plasma were prepared independently of those used for the calibration curves. These QC samples were prepared on the day of analysis in the same way as calibration standards.

15 µL standards and 15 µL QC samples and 15 µL unknown samples(10 µL plasma with 5 µL blank solution)were added to 200 µL of acetonitrile containing IS mixture for precipitating protein respectively. Then the samples were vortexed for 30 s. After centrifugation at 4 °C, 3900 rpm for 15 min, the supernatant was diluted 3 times with MeCN. 5 µL of supernatant was injected into the LC/MS/MS system for quantitative analysis.

The pharmacokinetic parameters of drug were analyzed using PK Solver Excel® Add-in. The PK trace was fitted into a non-compartmental extravascular model.

### Western blotting

Cells were seeded into six-well plates at a density of 5×10^5^ cells/mL in 2 mL of complete culture medium. Following an overnight adaptation period, cells were treated with serially diluted compounds for 24 h. After treatment, whole-cell lysates were prepared using a lysis buffer (1×RIPA supplemented with protease and phosphatase inhibitor cocktail). Protein concentrations in the lysates were measured using the BCA protein assay. Subsequently, equal amounts of protein (20 µg) from each sample were loaded onto a sodium dodecyl sulfate-polyacrylamide gel and separated by electrophoresis (Bio-Rad) at 120 V for 1.5 hours. The separated proteins were then transferred to a polyvinylidene fluoride (PVDF) membrane using a Transblot Turbo system (Bio-Rad). After blocking for 2 h at room temperature in 5% BSA-TBST, the membranes were incubated overnight at 4°C with specific primary antibodies (diluted at 1:1000 in TBST) targeting the proteins of interest, including anti-RIPK1 (3493, Cell Signaling Technology (CST)), anti-cleaved caspase 3 (9661, CST), anti-cleaved caspase7 (8438, CST), anti-cleaved PARP (5625,CST), and anti-β-actin (4970, CST). The membranes were then incubated with horseradish peroxidase-conjugated secondary antibodies (1:1000, 7074, CST) for 1 h at room temperature. Immunoblots were imaged using ECL Prime chemiluminescent western blot detection reagent (R1100, Kindle Biosciences,) and visualized using an Imager (D1001, Kindle Biosciences). All western blots were processed and quantified using ImageJ software, and protein levels were normalized to β-actin loading controls.

### Cell viability assay

urkat cells (7,500 cells per well) were seeded into 96-well plates containing 150 μL of RPMI medium supplemented with 10% FBS. On the following day, DMSO or drug was added. After 72 hrs, cell viability was measured by Alamar Blue assay. For statistical significance, six replications were tested in each treatment. Relative cell viabilities, referred to DMSO control cells, were plotted using Graphpad Prism.

### Apoptosis detection using FITC-conjugated Annexin V/PI

Apoptosis quantification was conducted utilizing a FITC-conjugated Annexin V/PI assay kit (556547, BD Biosciences) and analyzed through flow cytometry. Briefly, 2×10^5^ of Jurkat cells were seeded onto six-well plates and treated as specified for 72 hours at 37°C. Treated and untreated cells were harvested, washed with PBS, and resuspended in 100 µl of binding buffer. Subsequently, cells were stained with PI (50 µg/ml) and FITC-conjugated Annexin V (10 mg/ml) for 15 minutes at room temperature in the dark. After adding another 400 µl of binding buffer, the cells were subjected to LSR II Flow cytometer (BD Biosciences) for analysis, and flow cytometry data were processed using the FlowJo software.

### Pharmacodynamics study

Jurkat cells (2.5×10^6^ cells per mouse) were implanted subcutaneously into right side of male nude mice (6-weeks, the Jackson laboratory). After 7 days after tumor inoculation, mice (n = 4 per group) were treated with drugs (10 mg/kg) or the solvent vehicle via intravenous injection. After 6 hours or 24 hours, mice were euthanized and tumor tissues were collect for Western blot.

## ABBREVIATIONS

CoCl_2_.6H_2_O: Cobalt chloride hexahydrate
CuI: Copper(I) iodide
DCM: dichloromethane
DIPEA: N,N-Diisopropylethylamine
DMF: N,N-Dimethylformamide
EtOAc: ethyl acetate
FA: formic acid
HATU: 2-(7-Aza-1H-benzotriazole-1-yl)-1,1,3,3-tetraMethyluroniuM hexafluorophosphate
IV: intravenous
K_2_CO_3_: Potassium carbonate
KI: Potassium iodide
LiOH: Lithium hydroxide
nLuc: NanoLuc luciferase
MeCN: Acetonitrile
MeOH: methanol
MgSO_4_: magnesium sulfate
NaBH_4_: Sodium borohydride
N_2_: nitrogen gas, NADPH, nicotinamide adenine dinucleotide phosphate
NaH: sodium hydride
NF-κB: Nuclear factor kappa-light-chain-enhancer of activated B cells
PD: pharmacodynamics
PD-1: programmed cell death protein 1
Pd(PPh_3_)_2_Cl_2_: Bis(triphenylphosphine)palladium(II) chloride
PK: pharmacokinetics
PROTAC: proteolysis targeting chimera
RT: room temperature
TEA: triethylamine
TFA: Trifluoroacetic acid
THF: Tetrahydrofuran
VHL: von Hippel-Lindau Cullin RING E3 ligase
WT: wild-type.

## ASSOCIATED CONTENT

### Supporting Information

Supporting Information is available free of charge via the Internet at http://pubs.acs.org.

Figure S1-S7, Table S1, spectra and purity (PDF)

Molecular formula strings (CSV).

Research involving animals was performed in accordance with institutional guidelines as defined by Institutional Animal Care and Use Committee (AN-6075).

## Notes

J.W. is the co-founder of Chemical Biology Probes LLC. J. W. has stock ownership in CoRegen Inc and serves as a consultant for this company. J.W., X.Y. and B.Y. are the co-founders of Fortitude Biomedicines, Inc. and hold equity interest in this company. X.Y., D.L., and J.W. are inventors on a patent covering RIPK1 degraders reported in this work, titled “Novel RIPK1 Kinase-Targeting PROTACs and Methods of Use Thereof”, with the identification number WO2022120118A1. The remaining authors declare no competing interests.

## ACKNOWLEDGMENT

The research was supported in part by National Institute of Health (R01-268518 to J.W.), Cancer Prevention & Research Institute of Texas (CPRIT, RP220480 to J.W.), and the Michael E. DeBakey, M.D., Professorship in Pharmacology (to J.W.).

## Insert Table of Contents artwork here

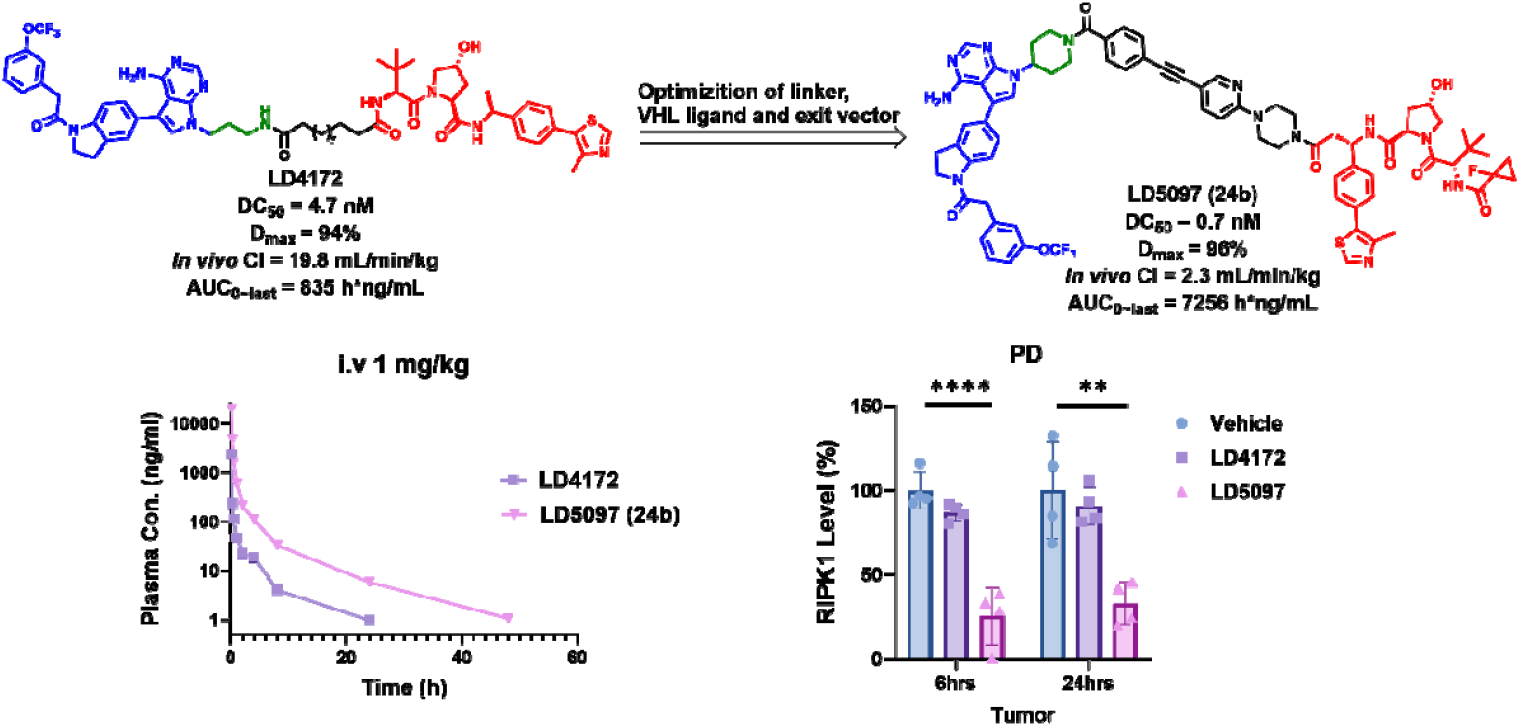

## REFERENCES

(1) Liu, L.; Lalaoui, N. 25 Years of Research Put RIPK1 in the Clinic. Seminars in Cell & Developmental Biology 2020, S1084952120301142. 10.1016/j.semcdb.2020.08.007.

(2) Silke, J.; Rickard, J. A.; Gerlic, M. The Diverse Role of RIP Kinases in Necroptosis and Inflammation. Nat Immunol 2015, 16 (7), 689–697. 10.1038/ni.3206.

(3) Vandenabeele, P.; Declercq, W.; Van Herreweghe, F.; Vanden Berghe, T. The Role of the Kinases RIP1 and RIP3 in TNF-Induced Necrosis. Science Signaling 2010, 3 (115), re4–re4. 10.1126/scisignal.3115re4.

(4) Yuan, J.; Amin, P.; Ofengeim, D. Necroptosis and RIPK1-Mediated Neuroinflammation in CNS Diseases. Nat Rev Neurosci 2019, 20 (1), 19–33. 10.1038/s41583-018-0093-1.

(5) Samson, A. L.; Garnish, S. E.; Hildebrand, J. M.; Murphy, J. M. Location, Location, Location: A Compartmentalized View of TNF-Induced Necroptotic Signaling. Sci. Signal. 2021, 14 (668), eabc6178. 10.1126/scisignal.abc6178.

(6) Ofengeim, D.; Yuan, J. Regulation of RIP1 Kinase Signal-ling at the Crossroads of Inflammation and Cell Death. Nat Rev Mol Cell Biol 2013, 14 (11), 727–736. 10.1038/nrm3683.

(7) Mifflin, L.; Ofengeim, D.; Yuan, J. Receptor-Interacting Protein Kinase 1 (RIPK1) as a Therapeutic Target. Nat Rev Drug Discov 2020, 19 (8), 553–571. 10.1038/s41573-020-0071-y.

(8) Ofengeim, D.; Mazzitelli, S.; Ito, Y.; DeWitt, J. P.; Mifflin, L.; Zou, C.; Das, S.; Adiconis, X.; Chen, H.; Zhu, H.; Kelliher, M. A.; Levin, J. Z.; Yuan, J. RIPK1 Mediates a Disease-Associated Microglial Response in Alzheimer’s Disease. Proc Natl Acad Sci USA 2017, 114 (41), E8788–E8797. 10.1073/pnas.1714175114.

(9) He, S.; Wang, X. RIP Kinases as Modulators of Inflammation and Immunity. Nat Immunol 2018, 19 (9), 912–922. 10.1038/s41590-018-0188-x.

(10) Xia, X.; Lei, L.; Wang, S.; Hu, J.; Zhang, G. Necroptosis and Its Role in Infectious Diseases. Apoptosis 2020, 25 (3–4), 169–178. 10.1007/s10495-019-01589-x.

(11) Guo, H.; Omoto, S.; Harris, P. A.; Finger, J. N.; Bertin, J.; Gough, P. J.; Kaiser, W. J.; Mocarski, E. S. Herpes Simplex Virus Suppresses Necroptosis in Human Cells. Cell Host & Microbe 2015, 17 (2), 243–251. 10.1016/j.chom.2015.01.003.

(12) Strilic, B.; Yang, L.; Albarrán-Juárez, J.; Wachsmuth, L.; Han, K.; Müller, U. C.; Pasparakis, M.; Offermanns, S. Tumour-Cell-Induced Endothelial Cell Necroptosis via Death Receptor 6 Promotes Metastasis. Nature 2016, 536 (7615), 215–218. 10.1038/nature19076.

(13) Jiao, D.; Cai, Z.; Choksi, S.; Ma, D.; Choe, M.; Kwon, H.-J.; Baik, J. Y.; Rowan, B. G.; Liu, C.; Liu, Z. Necroptosis of Tumor Cells Leads to Tumor Necrosis and Promotes Tumor Metastasis. Cell Res 2018, 28 (8), 868–870. 10.1038/s41422-018-0058-y.

(14) Krysko, O.; Aaes, T. L.; Kagan, V. E.; D’Herde, K.; Bachert, C.; Leybaert, L.; Vandenabeele, P.; Krysko, D. V. Necroptotic Cell Death in Anti-Cancer Therapy. Immunological Reviews 2017, 280 (1), 207–219. 10.1111/imr.12583.

(15) Hou, J.; Ju, J.; Zhang, Z.; Zhao, C.; Li, Z.; Zheng, J.; Sheng, T.; Zhang, H.; Hu, L.; Yu, X.; Zhang, W.; Li, Y.; Wu, M.; Ma, H.; Zhang, X.; He, S. Discovery of Potent Necroptosis Inhibitors Targeting RIPK1 Kinase Activity for the Treatment of Inflammatory Disorder and Cancer Metastasis. Cell Death Dis 2019, 10 (7), 493. 10.1038/s41419-019-1735-6.

(16) Manguso, R. T.; Pope, H. W.; Zimmer, M. D.; Brown, F. D.; Yates, K. B.; Miller, B. C.; Collins, N. B.; Bi, K.; LaFleur, M. W.; Juneja, V. R.; Weiss, S. A.; Lo, J.; Fisher, D. E.; Miao, D.; Van Allen, E.; Root, D. E.; Sharpe, A. H.; Doench, J. G.; Haining, W. N. In Vivo CRISPR Screening Identifies Ptpn2 as a Cancer Immunotherapy Target. Nature 2017, 547 (7664), 413–418. 10.1038/nature23270.

(17) Cucolo, L.; Chen, Q.; Qiu, J.; Yu, Y.; Klapholz, M.; Budinich, K. A.; Zhang, Z.; Shao, Y.; Brodsky, I. E.; Jordan, M. S.; Gilliland, D. G.; Zhang, N. R.; Shi, J.; Minn, A. J. The Interferon-Stimulated Gene RIPK1 Regulates Cancer Cell Intrinsic and Extrinsic Resistance to Immune Checkpoint Blockade. Immunity 2022, 55 (4), 671-685.e10. 10.1016/j.immuni.2022.03.007.

(18) Hou, J.; Wang, Y.; Shi, L.; Chen, Y.; Xu, C.; Saeedi, A.; Pan, K.; Bohat, R.; Egan, N. A.; McKenzie, J. A.; Mbofung, R. M.; Williams, L. J.; Yang, Z.; Sun, M.; Liang, X.; Rodon Ahnert, J.; Varadarajan, N.; Yee, C.; Chen, Y.; Hwu, P.; Peng, W. Integrating Genome-Wide CRISPR Immune Screen with Multi-Omic Clinical Data Reveals Distinct Classes of Tumor Intrinsic Immune Regulators. J Immunother Cancer 2021, 9 (2), e001819. 10.1136/jitc-2020-001819.

(19) Yu, X.; Lu, D.; Qi, X.; Paudel, R. R.; Lin, H.; Holloman, B. L.; Jin, F.; Xu, L.; Ding, L.; Peng, W.; Wang, M. C.; Chen, X.; Wang, J. Development of a RIPK1 Degrader to Enhance Antitumor Immunity. Nat Commun 2024, 15 (1), 10683. 10.1038/s41467-024-55006-2.

(20) Zhang, Z.; Li, C.; Hawkins, N. J.; Mudududdla, R.; Nie, Y.; Liu, P.-K.; Huang, P.; Del Rio, N. M.; Chang, H.; Brown, M. E.; Li, L.; Tang, W. Development of Potent and Selective RIPK1 De-graders Targeting Its Nonenzymatic Function for Cancer Treatment. J. ed. Chem. 2025, 68 (14), 15120–15136. 10.1021/acs.jmedchem.5c01340.

(21) Mannion, J.; Gifford, V.; Bellenie, B.; Fernando, W.; Ramos Garcia, L.; Wilson, R.; John, S. W.; Udainiya, S.; Patin, E. C.; Tiu, C.; Smith, A.; Goicoechea, M.; Craxton, A.; Moraes De Vasconcelos, N.; Guppy, N.; Cheung, K.-M. J.; Cundy, N. J.; Pierrat, O.; Brennan, A.; Roumeliotis, T. I.; Benstead-Hume, G.; Alexander, J.; Muirhead, G.; Layzell, S.; Lyu, W.; Roulstone, V.; Allen, M.; Baldock, H.; Legrand, A.; Gabel, F.; Serrano-Aparicio, N.; Starling, C.; Guo, H.; Upton, J.; Gyrd-Hansen, M.; MacFarlane, M.; Seddon, B.; Raynaud, F.; Roxanis, I.; Harrington, K.; Haider, S.; Choudhary, J. S.; Hoelder, S.; Tenev, T.; Meier, P. A RIPK1-Specific PROTAC Degrader Achieves Potent Antitumor Activity by Enhancing Immunogenic Cell Death. Immunity 2024, S1074761324002309. 10.1016/j.immuni.2024.04.025.

(22) Inuzuka, H.; Qian, C.; Qi, Y.; Xiong, Y.; Wang, C.; Wang, Z.; Zhang, D.; Zhang, C.; Jin, J.; Wei, W. Targeted Degradation of Receptor-Interacting Protein Kinase 1 to Modulate the Necroptosis Pathway. ACS Pharmacol. Transl. Sci. 2024, 7 (11), 3518–3526. 10.1021/acsptsci.4c00421.

(23) Han, X.; Wang, C.; Qin, C.; Xiang, W.; Fernandez-Salas, E.; Yang, C.-Y.; Wang, M.; Zhao, L.; Xu, T.; Chinnaswamy, K.; Delproposto, J.; Stuckey, J.; Wang, S. Discovery of ARD-69 as a Highly Potent Proteolysis Targeting Chimera (PROTAC) Degrader of Androgen Receptor (AR) for the Treatment of Prostate Cancer. J. Med. Chem. 2019, 62 (2), 941–964. 10.1021/acs.jmedchem.8b01631.

(24) Buckley, D. L.; Gustafson, J. L.; Van Molle, I.; Roth, A. G.; Tae, H. S.; Taylor, P. C.; Jorgensen, W. L.; Ciulli, A.; Crews, C. M. Small-Molecule Inhibitors of the Interaction between the E3 LigaseVHL and HIF1a. Angew. Chem. Int. Ed. 2012, 51, 11463–11467.

(25) Soares, P.; Lucas, X.; Ciulli, A. Thioamide Substitution to Probe the Hydroxyproline Recognition of VHL Ligands. Bioorganic & Medicinal Chemistry 2018, 26 (11), 2992–2995. 10.1016/j.bmc.2018.03.034.

(26) Soares, P.; Gadd, M. S.; Frost, J.; Galdeano, C.; Ellis, L.; Epemolu, O.; Rocha, S.; Read, K. D.; Ciulli, A. Group-Based Optimization of Potent and Cell-Active Inhibitors of the von Hippel–Lindau (VHL) E3 Ubiquitin Ligase: Structure–Activity Relationships Leading to the Chemical Probe (2 S,4 R)-1-((S)-2-(1-Cyanocyclopropanecarboxamido)-3,3-Dimethylbutanoyl)-4-Hydroxy-N -(4-(4-Methylthiazol-5-Yl)Benzyl)Pyrrolidine-2-Carboxamide (VH298). J. Med. Chem. 2018, 61 (2), 599–618. 10.1021/acs.jmedchem.7b00675.

(27) Liu, X.; Kalogeropulou, A. F.; Domingos, S.; Makukhin, N.; Nirujogi, R. S.; Singh, F.; Shpiro, N.; Saalfrank, A.; Sammler, E.; Ganley, I. G.; Moreira, R.; Alessi, D. R.; Ciulli, A. Discovery of XL01126: A Potent, Fast, Cooperative, Selective, Orally Bioavailable, and Blood–Brain Barrier Penetrant PROTAC Degrader of Leucine-Rich Repeat Kinase 2. J. Am. Chem. Soc. 2022, jacs.2c05499. 10.1021/jacs.2c05499.

(28) Han, X.; Zhao, L.; Xiang, W.; Qin, C.; Miao, B.; Xu, T.; Wang, M.; Yang, C.-Y.; Chinnaswamy, K.; Stuckey, J.; Wang, S. Discovery of Highly Potent and Efficient PROTAC Degraders of Androgen Receptor (AR) by Employing Weak Binding Affinity VHL E3 Ligase Ligands. J. Med. Chem. 2019, 62 (24), 11218–11231. 10.1021/acs.jmedchem.9b01393.

(29) Troup, R. I.; Fallan, C.; Baud, M. G. J. Current Strategies for the Design of PROTAC Linkers: A Critical Review. Exploration of Targeted Anti-tumor Therapy 2020, 1 (5). 10.37349/etat.2020.00018.

(30) Bemis, T. A.; La Clair, J. J.; Burkart, M. D. Unraveling the Role of Linker Design in Proteolysis Targeting Chimeras: Miniperspective. J. Med. Chem. 2021, acs.jmedchem.1c00482. 10.1021/acs.jmedchem.1c00482.

(31) Goracci, L.; Desantis, J.; Valeri, A.; Castellani, B.; Eleuteri, M.; Cruciani, G. Understanding the Metabolism of Proteolysis Targeting Chimeras (PROTACs): The Next Step toward Pharmaceutical Applications. J. Med. Chem. 2020, acs.jmedchem.0c00793. 10.1021/acs.jmedchem.0c00793.

(32) Riching, K. M.; Mahan, S.; Corona, C. R.; McDougall, M.; Vasta, J. D.; Robers, M. B.; Urh, M.; Daniels, D. L. Quantitative Live-Cell Kinetic Degradation and Mechanistic Profiling of PROTAC Mode of Action. ACS Chem. Biol. 2018, 13 (9), 2758–2770. 10.1021/acschembio.8b00692.

(33) Grohmann, C.; Magtoto, C. M.; Walker, J. R.; Chua, N. K.; Gabrielyan, A.; Hall, M.; Cobbold, S. A.; Mieruszynski, S.; Brzozowski, M.; Simpson, D. S.; Dong, H.; Dorizzi, B.; Jacobsen, A. V.; Morrish, E.; Silke, N.; Murphy, J. M.; Heath, J. K.; Testa, A.; Maniaci, C.; Ciulli, A.; Lessene, G.; Silke, J.; Feltham, R. Development of NanoLuc-Targeting Protein Degraders and a Universal Reporter System to Benchmark Tag-Targeted Degradation Platforms. Nat Commun 2022, 13 (1), 2073. 10.1038/s41467-022-29670-1.

(34) Li, Y.; Xiong, Y.; Zhang, G.; Zhang, L.; Yang, W.; Yang, J.; Huang, L.; Qiao, Z.; Miao, Z.; Lin, G.; Sun, Q.; Niu, T.; Chen, L.; Niu, D.; Li, L.; Yang, S. Identification of 5-(2,3-Dihydro-1 H -Indol-5-Yl)-7 H -Pyrrolo[2,3-d]Pyrimidin-4-Amine Derivatives as a New Class of Receptor-Interacting Protein Kinase 1 (RIPK1) Inhibitors, Which Showed Potent Activity in a Tumor Metastasis Model. J. Med. Chem. 2018, 61 (24), 11398–11414. 10.1021/acs.jmedchem.8b01652.

(35) Huang, H.-T.; Dobrovolsky, D.; Paulk, J.; Yang, G.; Weisberg, E. L.; Doctor, Z. M.; Buckley, D. L.; Cho, J.-H.; Ko, E.; Jang, J.; Shi, K.; Choi, H. G.; Griffin, J. D.; Li, Y.; Treon, S. P.; Fischer, E. S.; Bradner, J. E.; Tan, L.; Gray, N. S. A Chemoproteomic Approach to Query the Degradable Kinome Using a Multi-Kinase Degrader. Cell Chemical Biology 2018, 25 (1), 88-99.e6. 10.1016/j.chembiol.2017.10.005.

(36) Donovan, K. A.; Ferguson, F. M.; Bushman, J. W.; Eleuteri, N. A.; Bhunia, D.; Ryu, S.; Tan, L.; Shi, K.; Yue, H.; Liu, X.; Dobrovolsky, D.; Jiang, B.; Wang, J.; Hao, M.; You, I.; Teng, M.; Liang, Y.; Hatcher, J.; Li, Z.; Manz, T. D.; Groendyke, B.; Hu, W.; Nam, Y.; Sengupta, S.; Cho, H.; Shin, I.; Agius, M. P.; Ghobrial, I. M.; Ma, M. W.; Che, J.; Buhrlage, S. J.; Sim, T.; Gray, N. S.; Fischer, E. S. Mapping the Degradable Kinome Provides a Resource for Expedited Degrader Development. Cell 2020, 183 (6), 1714-1731.e10. 10.1016/j.cell.2020.10.038.

(37) Bondeson, D. P.; Smith, B. E.; Burslem, G. M.; Buhimschi, A. D.; Hines, J.; Jaime-Figueroa, S.; Wang, J.; Hamman, B. D.; Ishchenko, A.; Crews, C. M. Lessons in PROTAC Design from Selective Degradation with a Promiscuous Warhead. Cell Chemical Biology 2018, 25 (1), 78-87.e5. 10.1016/j.chembiol.2017.09.010.

(38) Bussiere, J. L.; Martin, P.; Horner, M.; Couch, J.; Flaherty, M.; Andrews, L.; Beyer, J.; Horvath, C. Alternative Strategies for Toxicity Testing of Species-Specific Biopharmaceuticals. Int J Toxicol 2009, 28 (3), 230–253. 10.1177/1091581809337262.

