## Supplemental Info for "Design, Optimization and Development of RIPK1 Degraders with Improved Pharmacokinetic and Pharmacodynamic Properties"

### Supporting Information

#### Table of Contents

**Fitting curve of tested compounds for nLuc-RIPK1 degradation ……………...…S2**
**Time-Remaining drug curves of tested compounds in mouse liver S9 fraction .....S3**
**Time-Remaining drug curves of tested compounds in human hepatocytes ……..S4**
**Time-Remaining drug curves of tested compounds in human microsomes ……..S4**
**Plasma concentration-time curves of tested compounds in C57BL/6J mice …....S5**
**Degradation potency of LD5097 in MOLM14 and U937 cells ………….…...…S6**
**Quantification of RIPK1 levels in Ramos and A20 cells…………..…..…………...S7**
**Purity of target compounds……………….…………………...…..……………...S8**
**$^1$H NMR, $^{13}$C NMR spectral data of target compounds…………….……………..S9**
**HPLC chromatography of representative compounds…………………….……....S59**

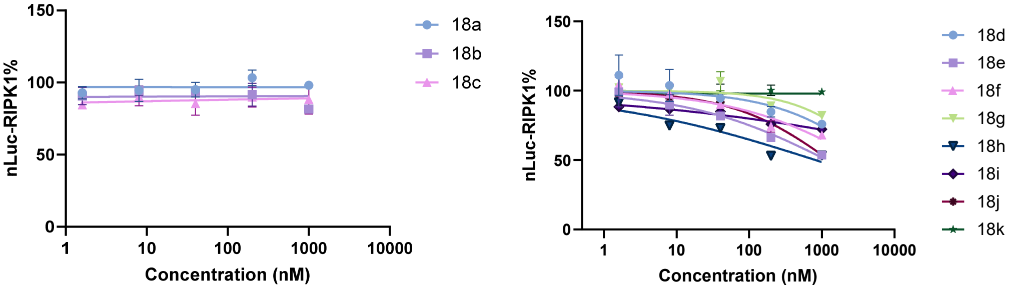

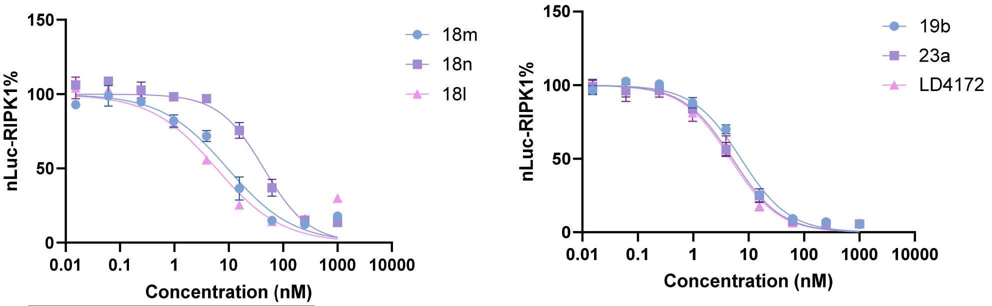

| | 12m | 12n | 12l |
|---|---|---|---|
| IC50 | 9.490 | 44.09 | 5.907 |

| | 19b | 23a | LD4172 |
|---|---|---|---|
| IC50 | 7.269 | 5.133 | 4.701 |

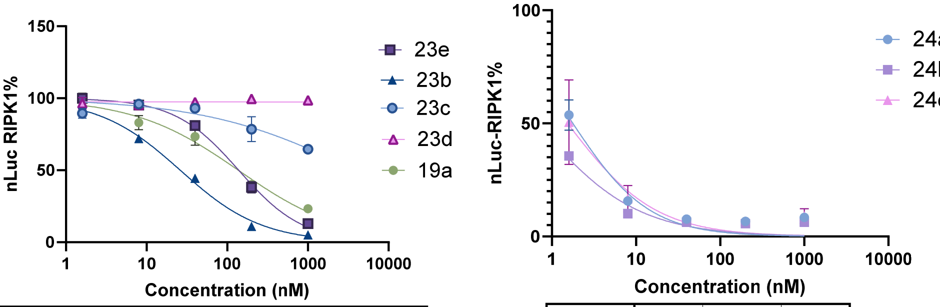

| | 23b | 23c | 23d | 19a | 23e |
|---|---|---|---|---|---|
| IC50 | 26.19 | 3866 | 1.336e+151 | 137.5 | 138.8 |

| | 24a | 24b | 24c |
|---|---|---|---|
| IC50 | 1.775 | 0.6626 | 1.536 |

**Figure. S1.** Fitting curve of tested compounds for nLuc-RIPK1 degradation in Jurkat cells.

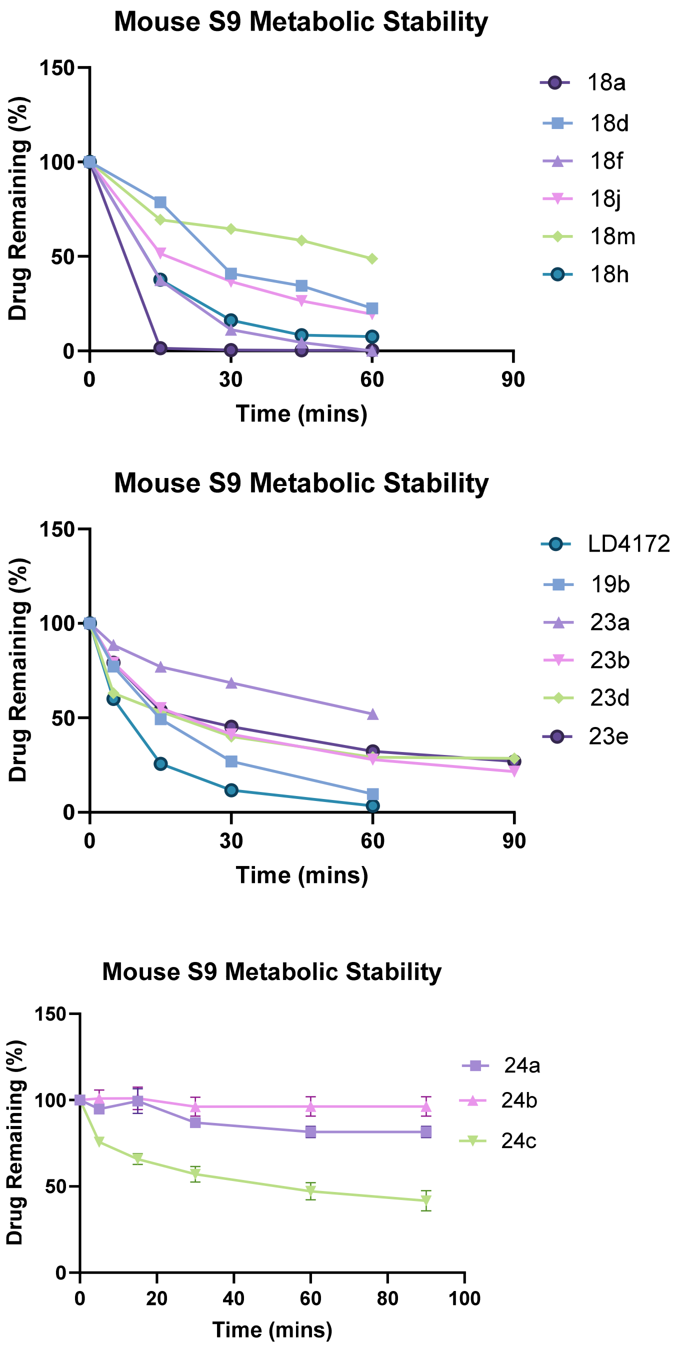

**Figure. S2.** Time-Remaining drug curves of tested compounds in mouse liver S9 fraction.

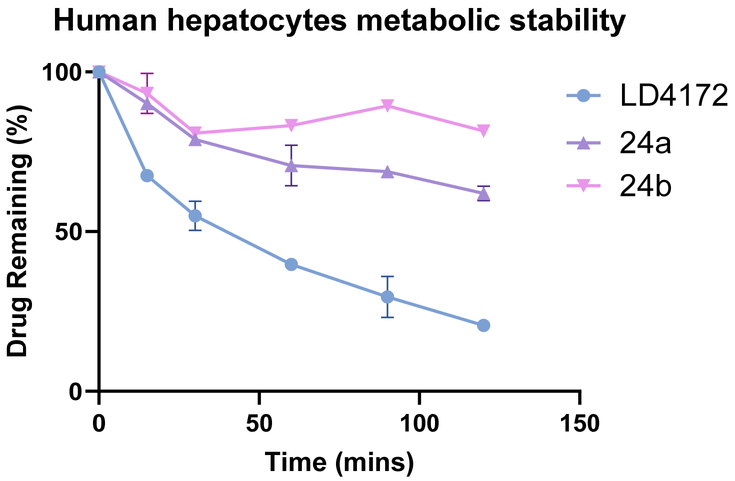

**Figure. S3.** Time-Remaining drug curves of tested compounds in human hepatocytes.

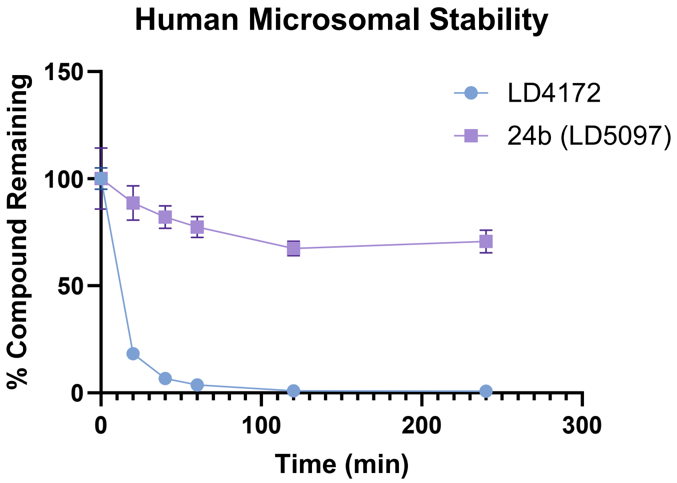

**Figure. S4.** Time-Remaining drug curves of tested compounds in human microsomes.

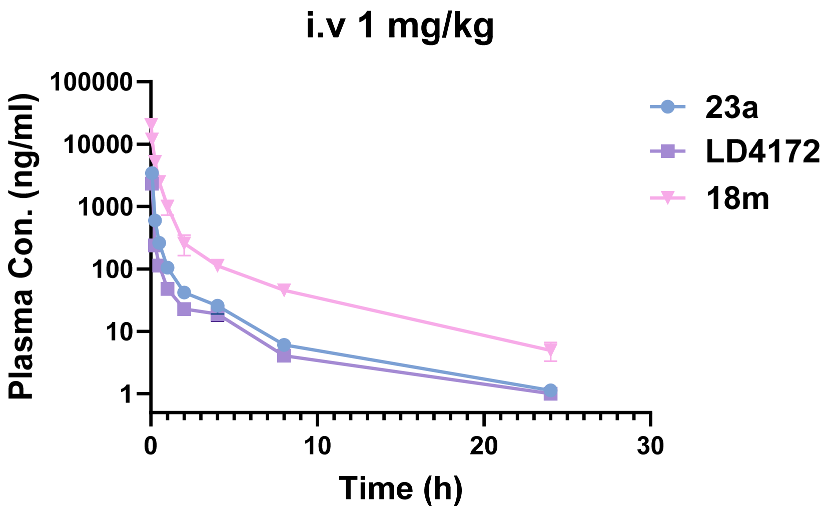

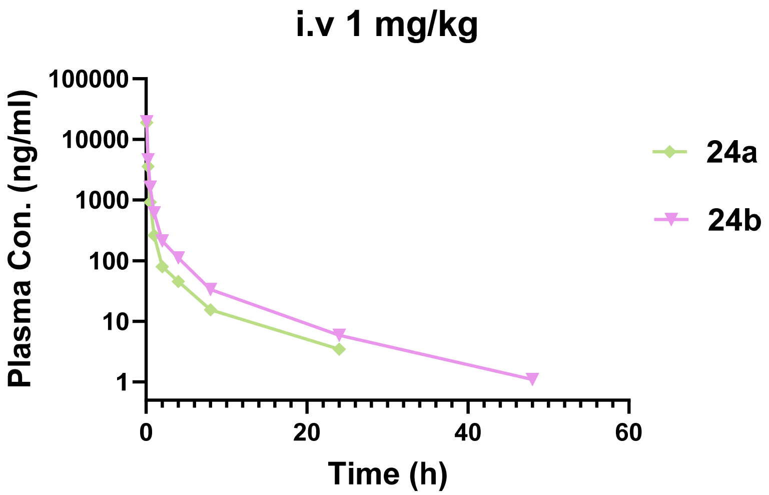

**Figure. S5.** Plasma concentration-time curves of tested compounds in C57BL/6J mice.

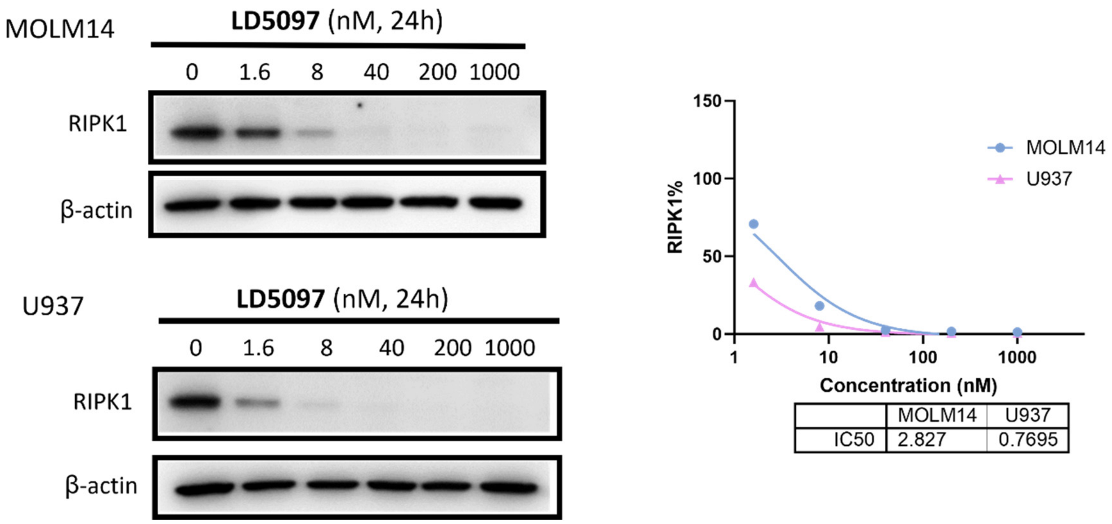

**Figure S6**. Degradation potency of **LD5097** (**24b**) in MOLM14 and U937 cells.

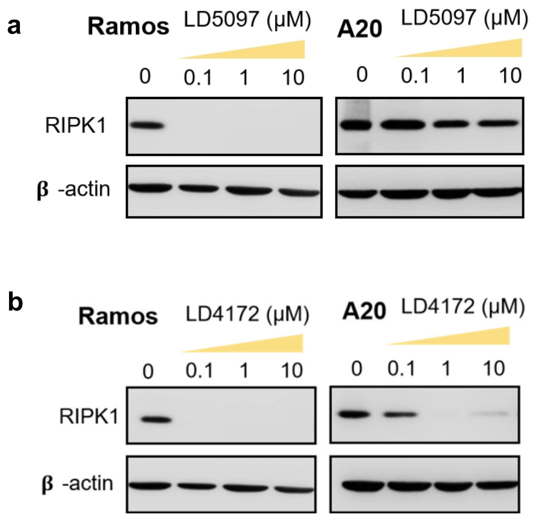

**Figure S7.** Quantification of RIPK1 levels in Ramos and A20 cells treated with LD5097 (a) and LD4172 (b) for 24 h at 0, 0.1, 1 and 10 μM, followed by Western blotting.

HPLC analyses were performed on an Agilent 1260 Infinity LC/MS System (column, Agilent Eclipse plus C18, 4.6 mm × 100 mm, 3.5 μm; with the solvent system of MeCN/$H_2O$/FA, 0.7 mL/min; UV wavelength, maximal absorbance at 254 nm; temperature, ambient; injection volume, 5 μL).

Table S1: Purity of target compounds

| Sample | Purity % | Sample | Purity % |
|---|---|---|---|
| **18a** | 95.7 | **18n** | 98.4 |
| **18b** | >99.0 | **19a** | 95.6 |
| **18c** | 96.4 | **19b** | 98.9 |
| **18d** | >99.0 | **23a** | 95.3 |
| **18e** | 97.8 | **23b** | 97.8 |
| **18f** | >99.0 | **23c** | 98.9 |
| **18g** | 95.6 | **23d** | >99.0 |
| **18h** | 96.6 | **23e** | 99.0 |
| **18i** | >99.0 | **24a** | 95.0 |
| **18j** | 98.0 | **24b** | 98.1 |
| **18k** | 95.9 | **24c** | 95.3 |
| **18l** | 95.6 | **LD5097-NC** | >99.0 |

| | |
|---|---|
| **18m** | 95.7 |

### $^{1}H$ NMR, $^{13}C$ NMR spectra of representative compounds

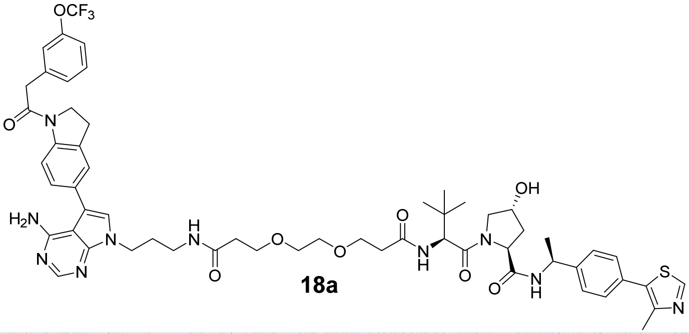

**18a**

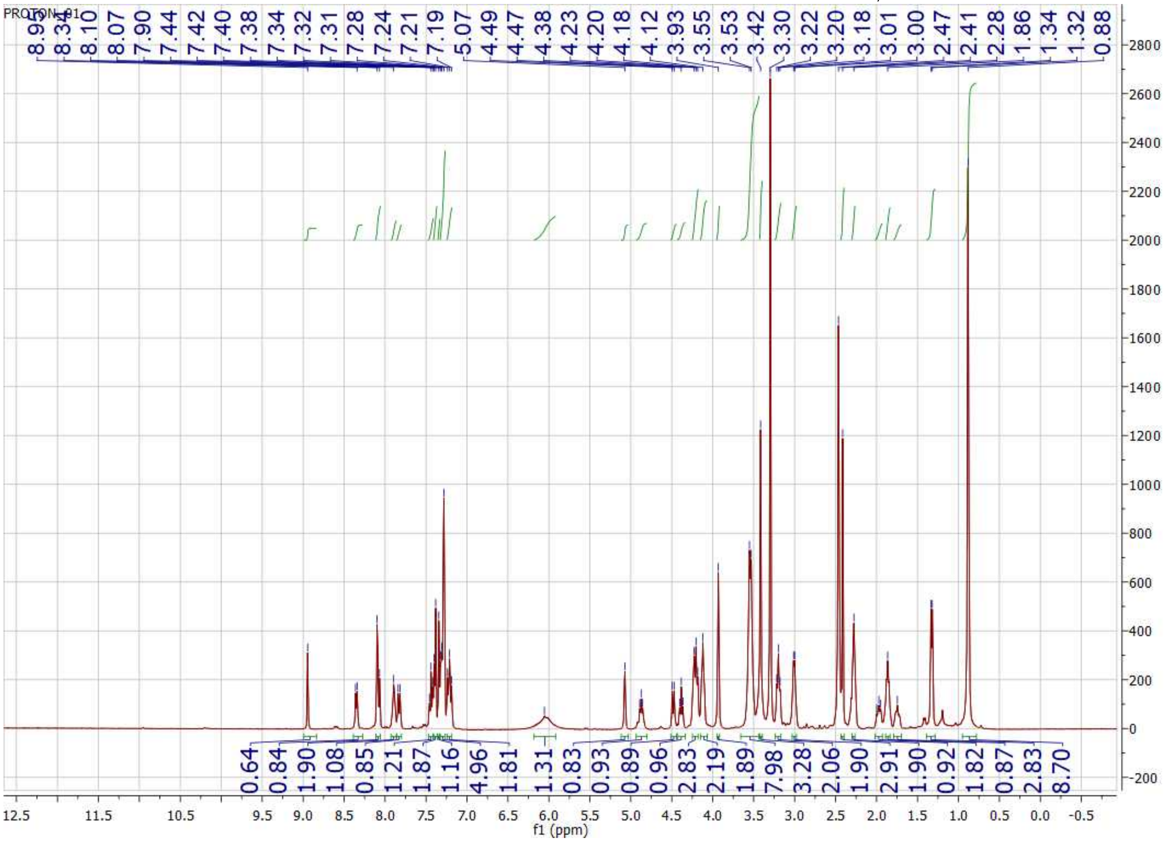

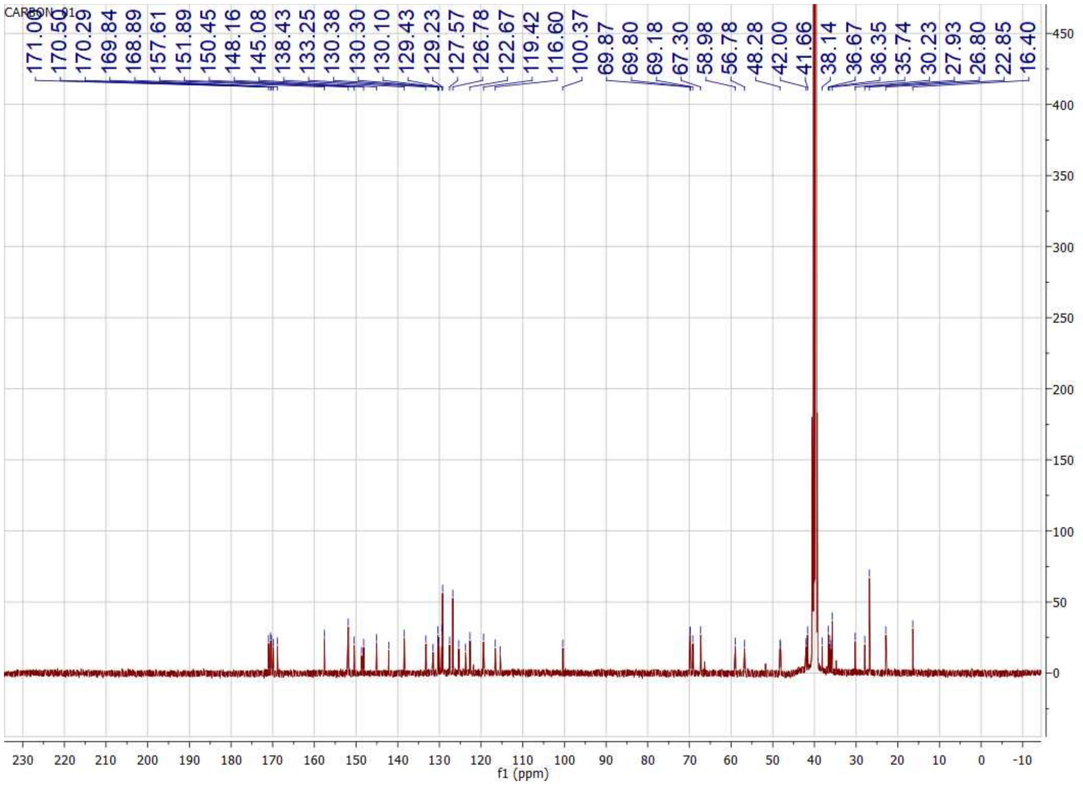
CARBON_01
171.06
170.50
170.29
169.84
168.89
157.61
151.89
150.45
148.16
145.08
138.43
133.25
130.38
130.30
130.10
129.43
129.23
127.57
126.78
122.67
119.42
116.60
100.37
69.87
69.80
69.18
67.30
58.98
56.78
48.28
42.00
41.66
38.14
36.67
36.35
35.74
30.23
27.93
26.80
22.85
16.40
f1 (ppm)

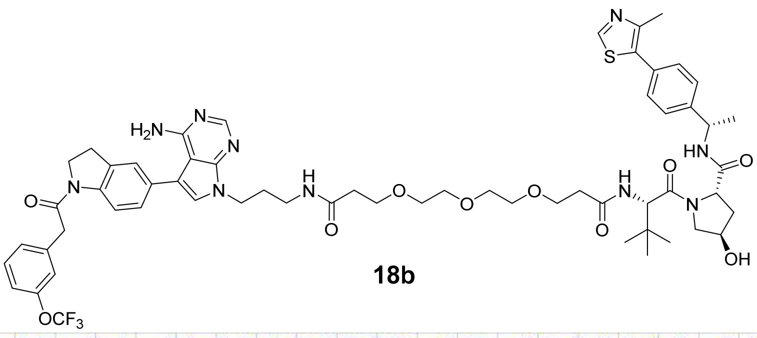

**18b**

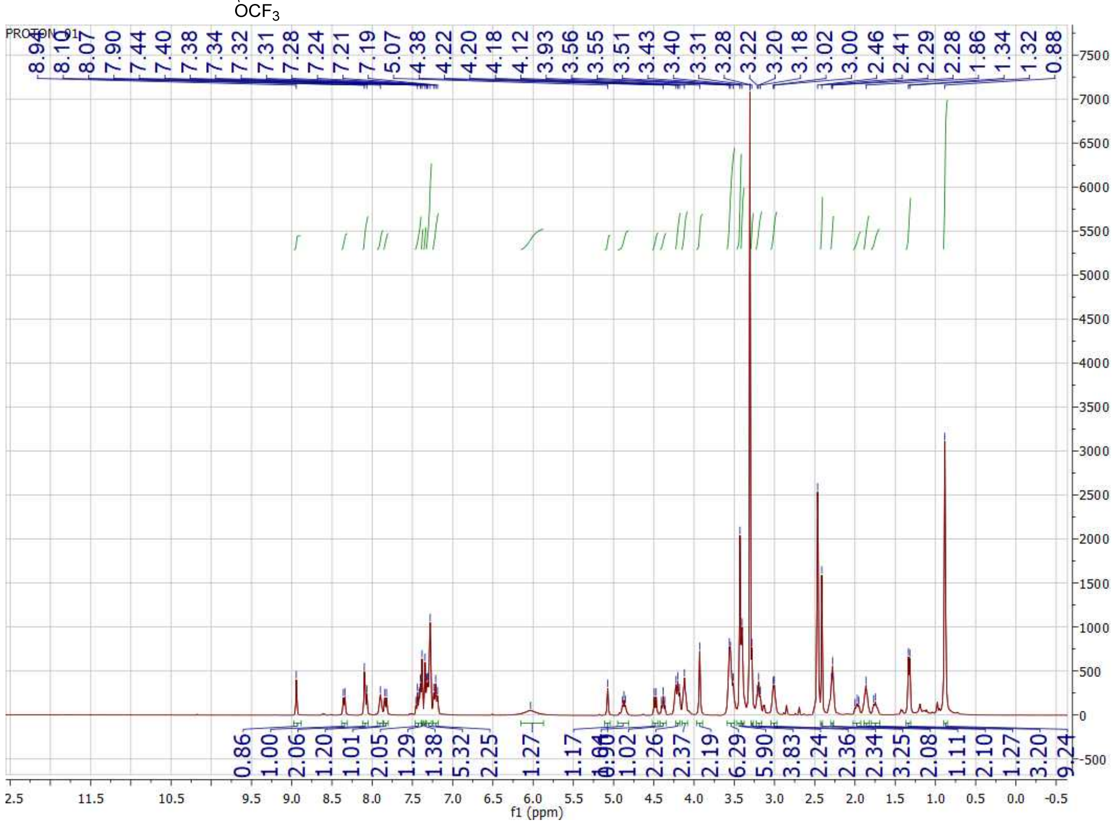

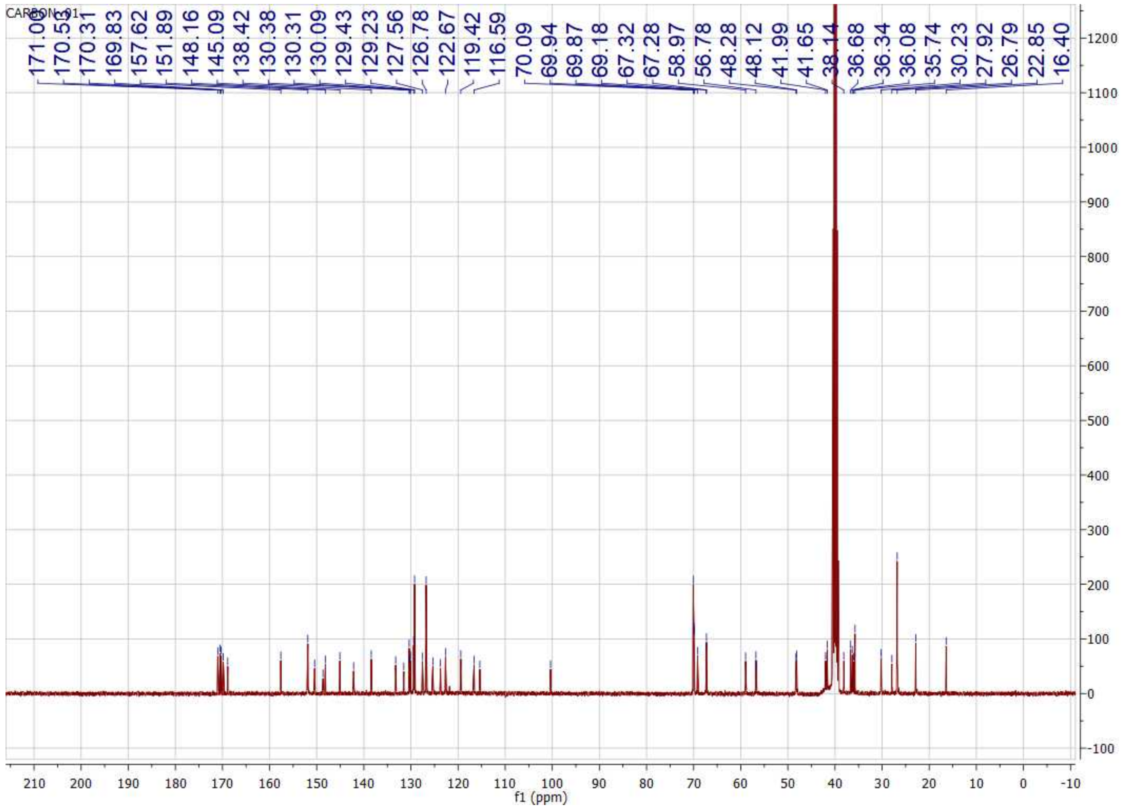
CARBON_01
171.00
170.52
170.31
169.83
157.62
151.89
148.16
145.09
138.42
130.38
130.31
130.09
129.43
129.23
127.56
126.78
122.67
119.42
116.59
70.09
69.94
69.87
69.18
67.32
67.28
58.97
56.78
48.28
48.12
41.99
41.65
38.14
36.68
36.34
36.08
35.74
30.23
27.92
26.79
22.85
16.40
f1 (ppm)

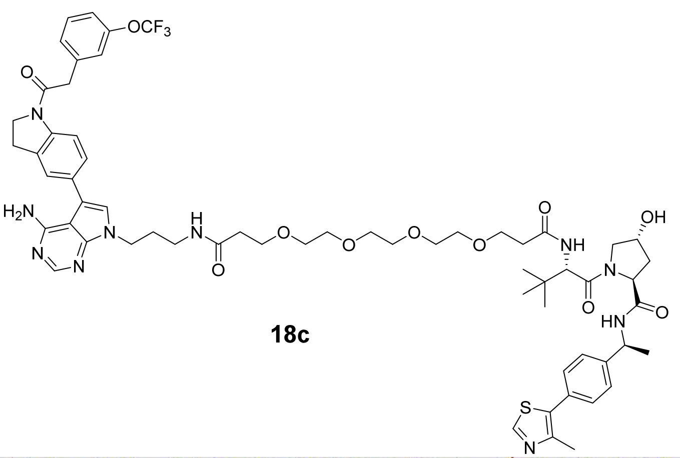

**18c**

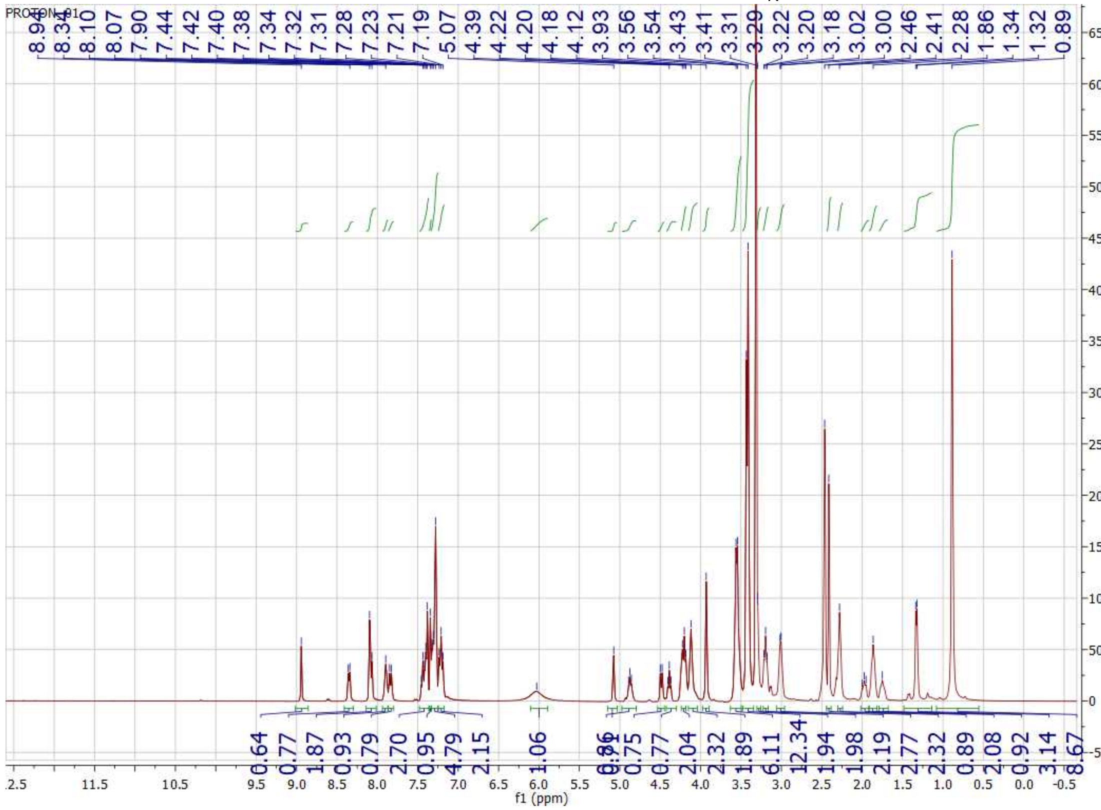

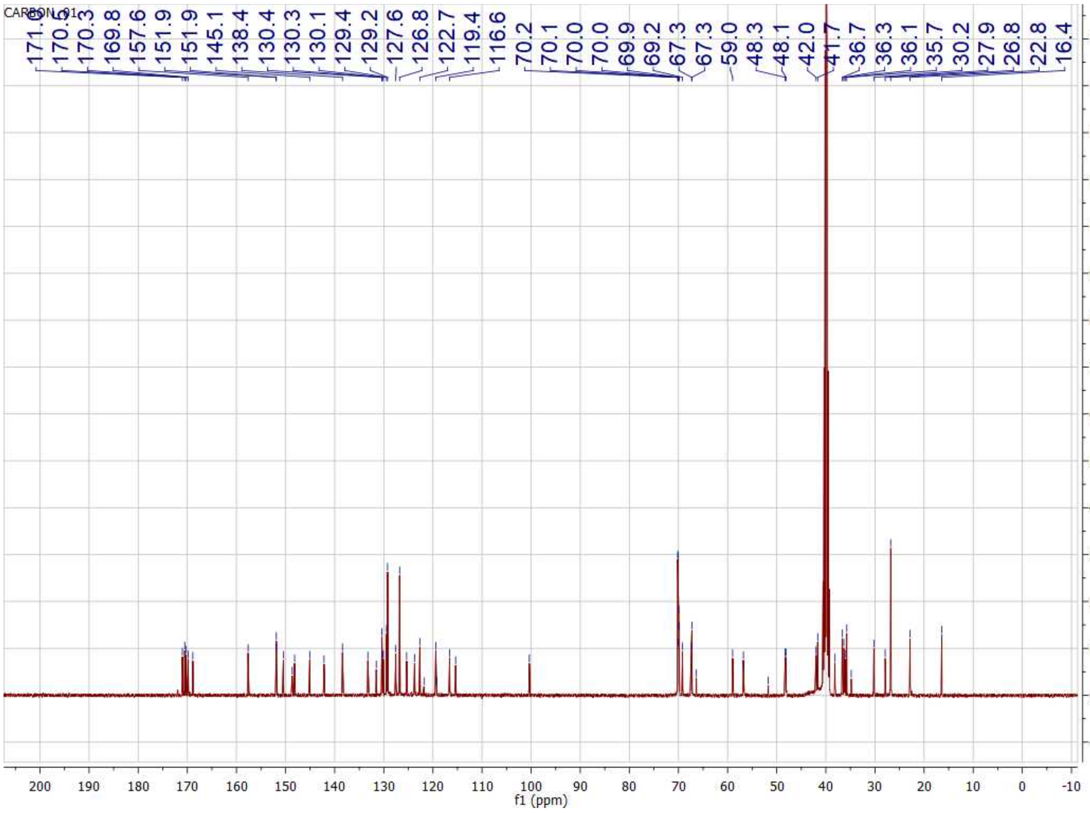

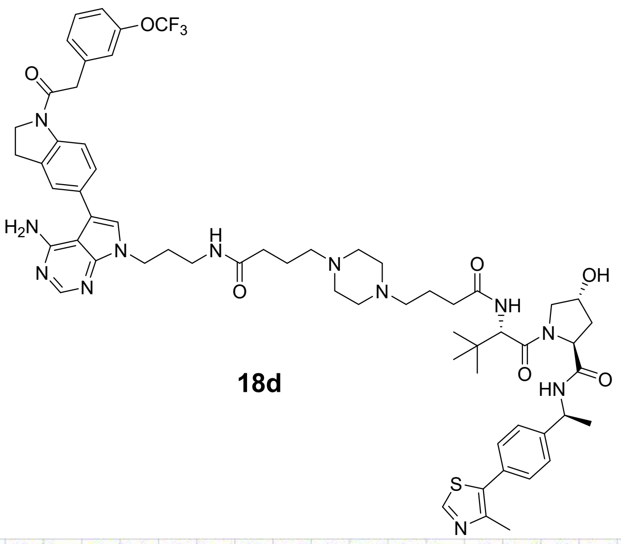

**18d**

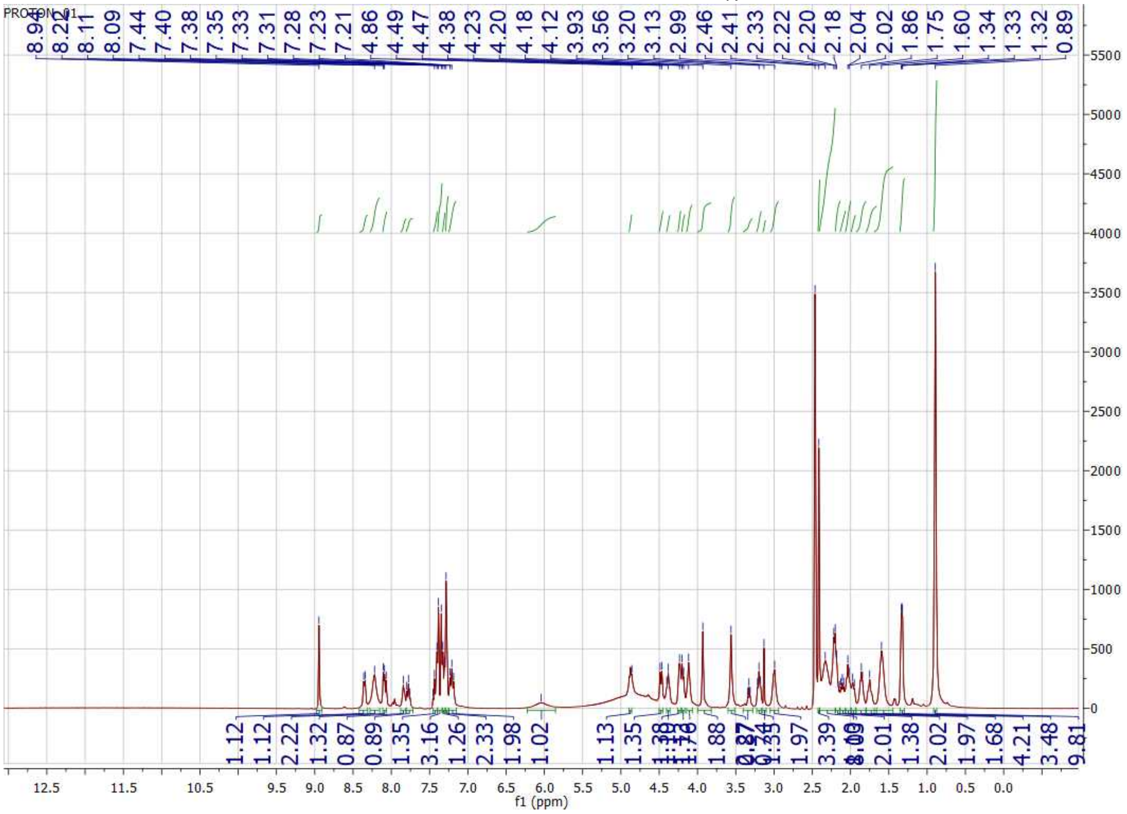

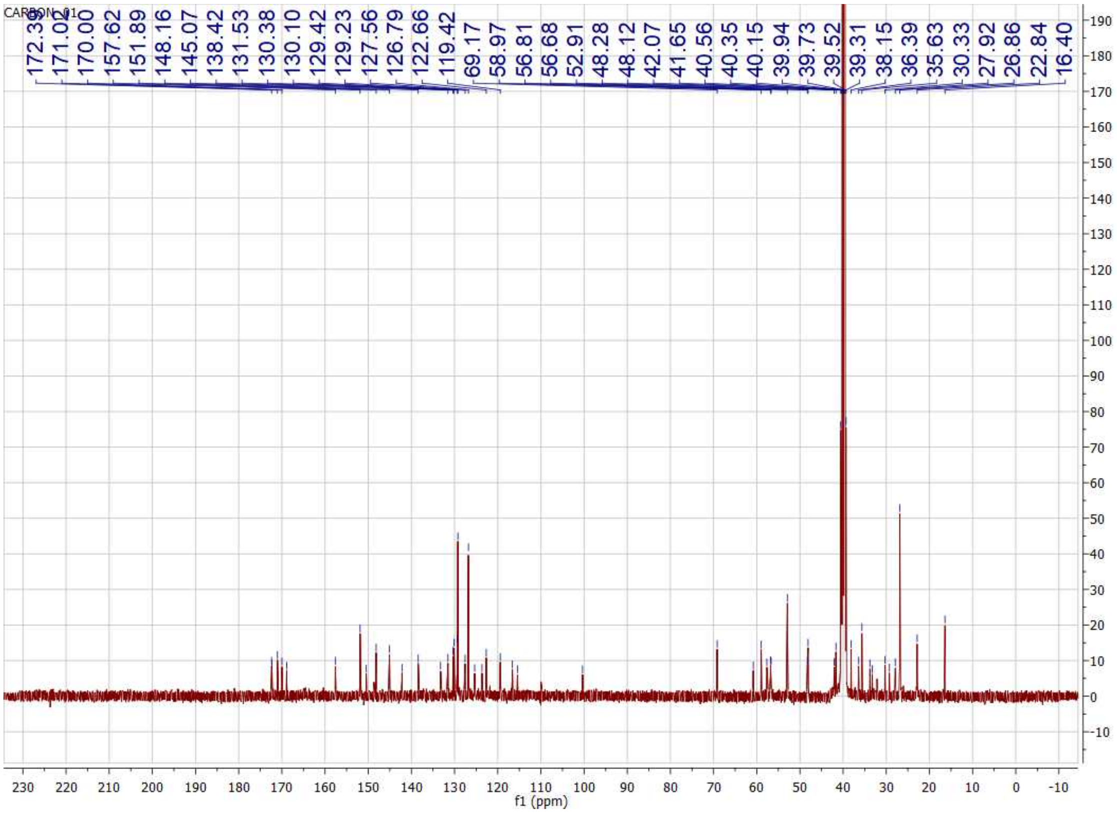
CARBON_01
172.39
171.02
170.00
157.62
151.89
148.16
145.07
138.42
131.53
130.38
130.10
129.42
129.23
127.56
126.79
122.66
119.42
69.17
58.97
56.81
56.68
52.91
48.28
48.12
42.07
41.65
40.56
40.35
40.15
39.94
39.73
39.52
39.31
38.15
36.39
35.63
30.33
27.92
26.86
22.84
16.40
f1 (ppm)

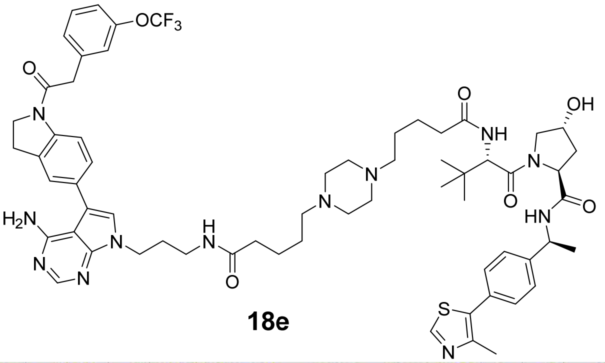

**18e**

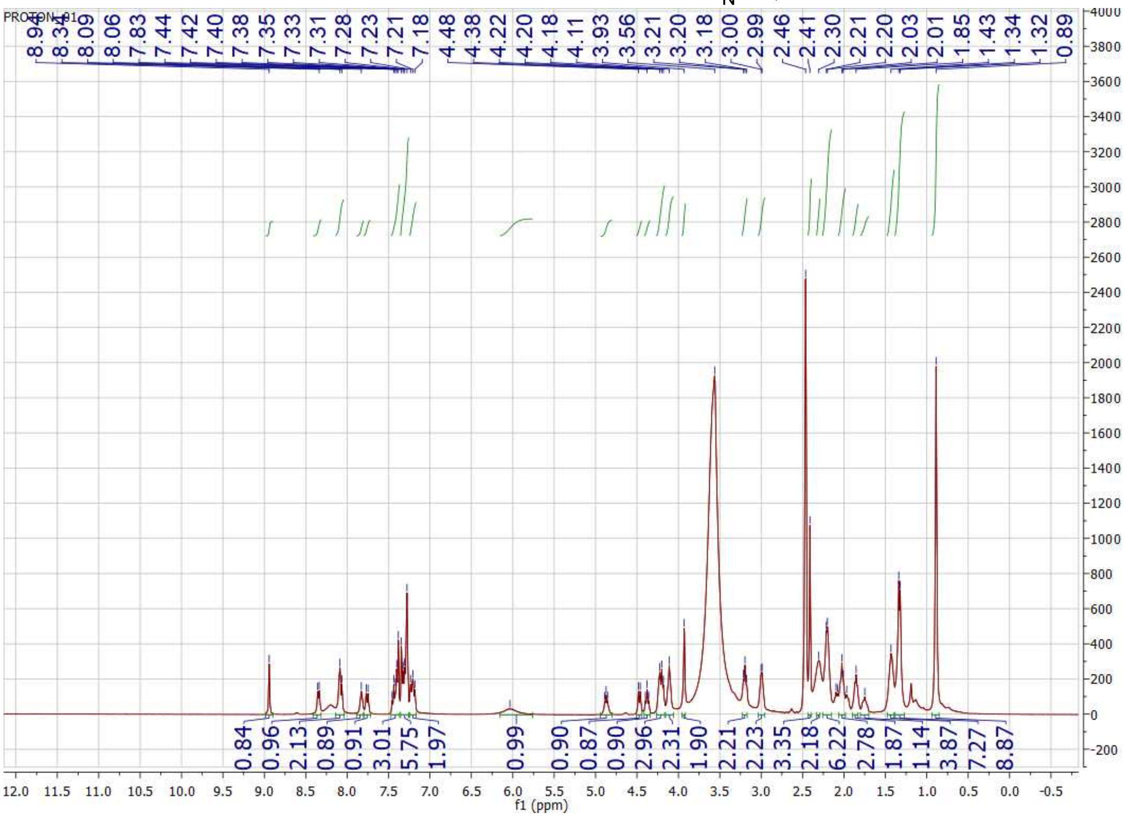

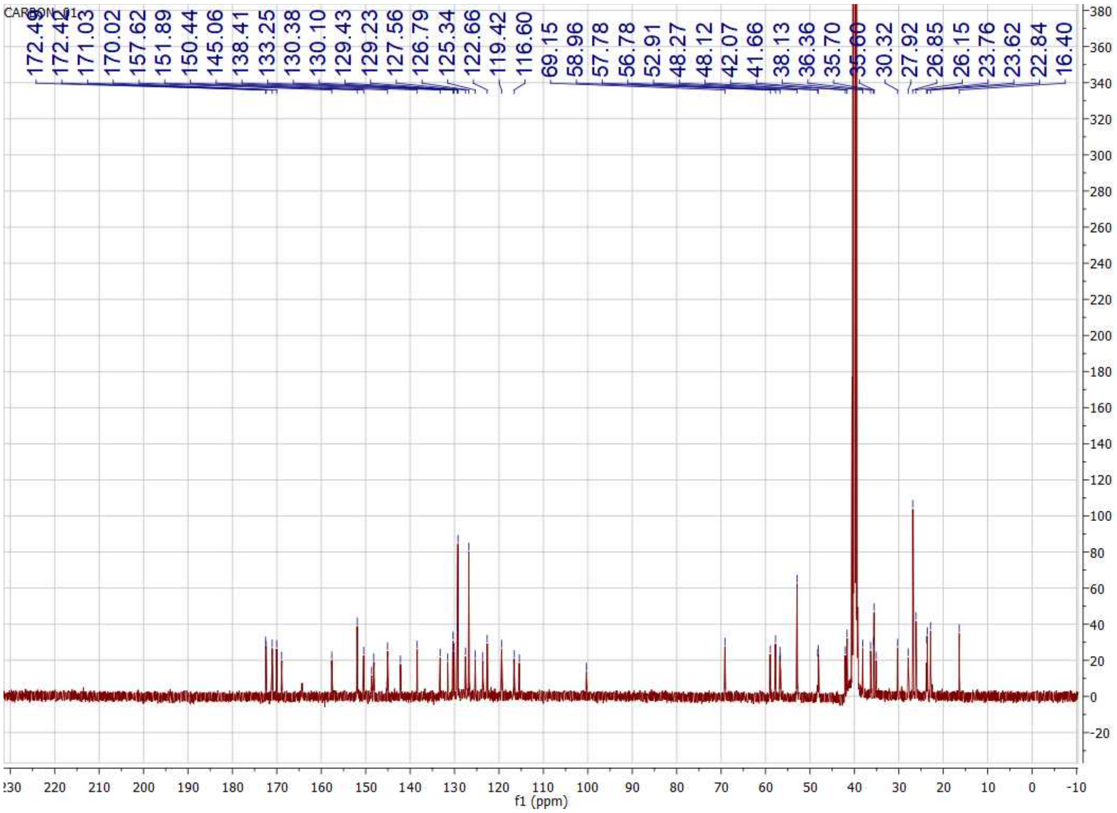
CARBON_01
172.42
171.03
170.02
157.62
151.89
150.44
145.06
138.41
133.25
130.38
130.10
129.43
129.23
127.56
126.79
125.34
122.66
119.42
116.60
69.15
58.96
57.78
56.78
52.91
48.27
48.12
42.07
41.66
38.13
36.36
35.70
30.32
27.92
26.85
26.15
23.76
23.62
22.84
16.40
f1 (ppm)

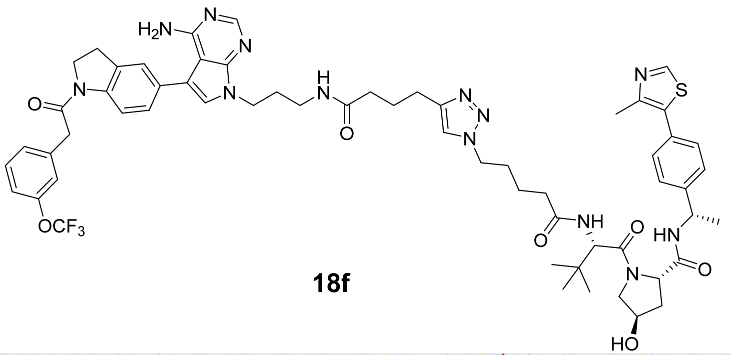

**18f**

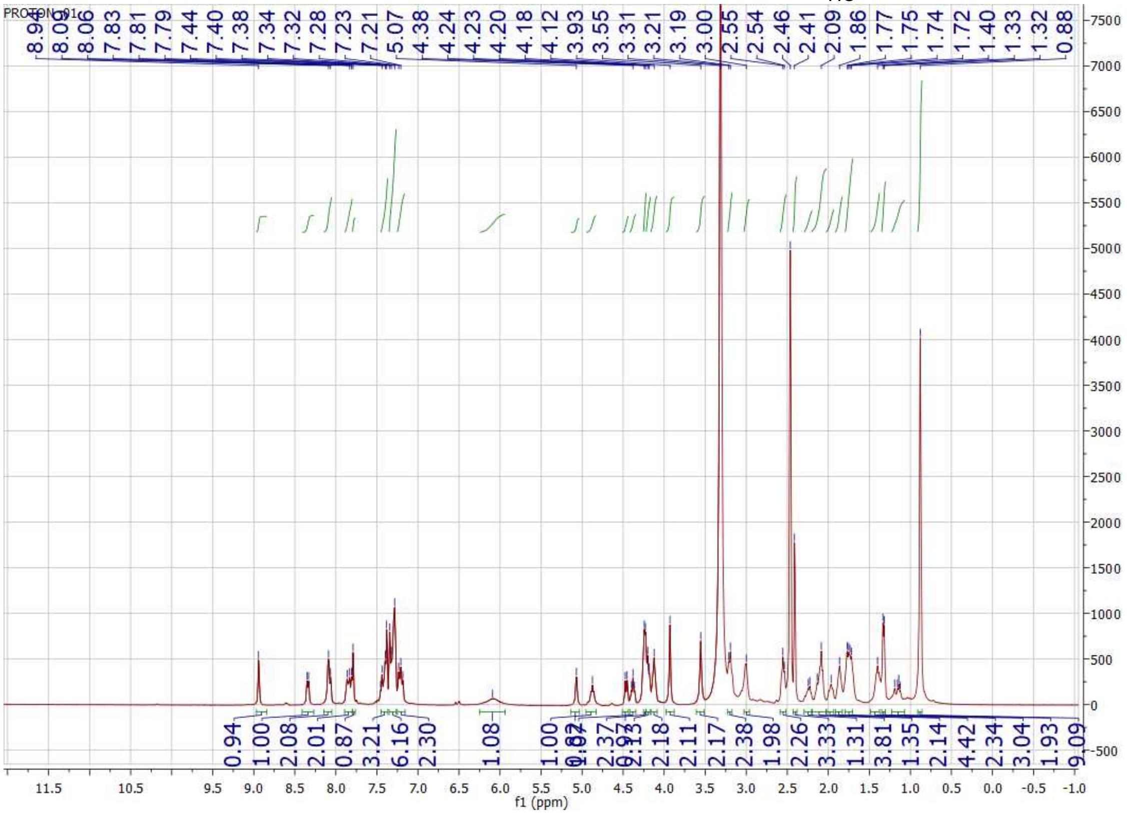

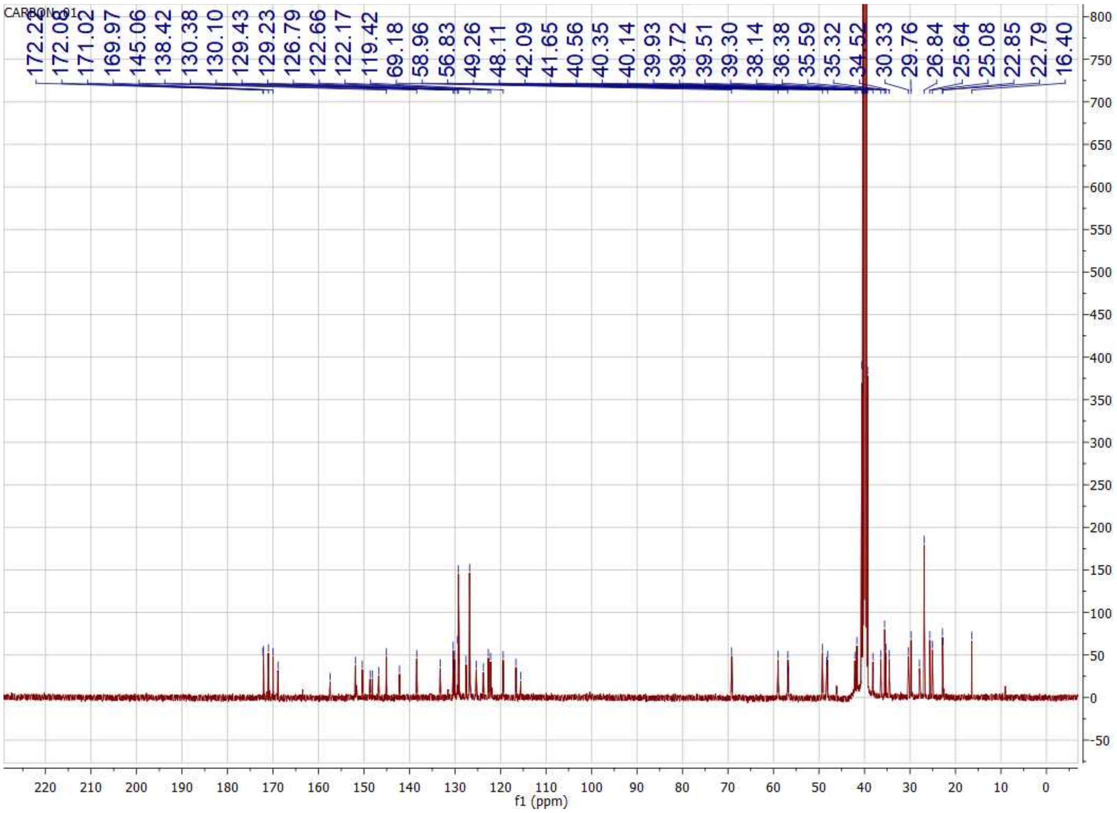
CARBON_01
172.22
172.08
171.02
169.97
145.06
138.42
130.38
130.10
129.43
129.23
126.79
122.66
122.17
119.42
69.18
58.96
56.83
49.26
48.11
42.09
41.65
40.56
40.35
40.14
39.93
39.72
39.51
39.30
38.14
36.38
35.59
35.32
34.52
30.33
29.76
26.84
25.64
25.08
22.85
22.79
16.40
f1 (ppm)

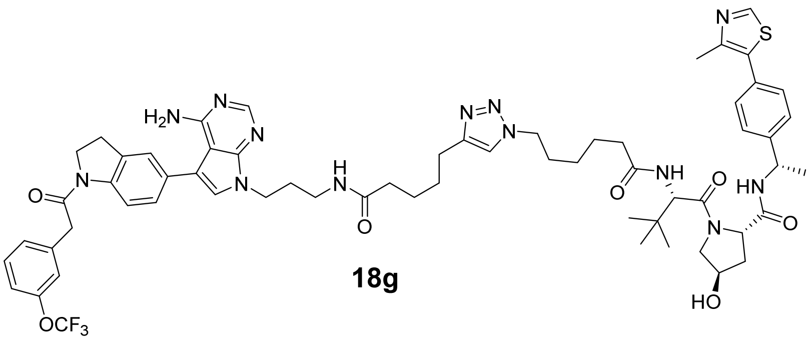

**18g**

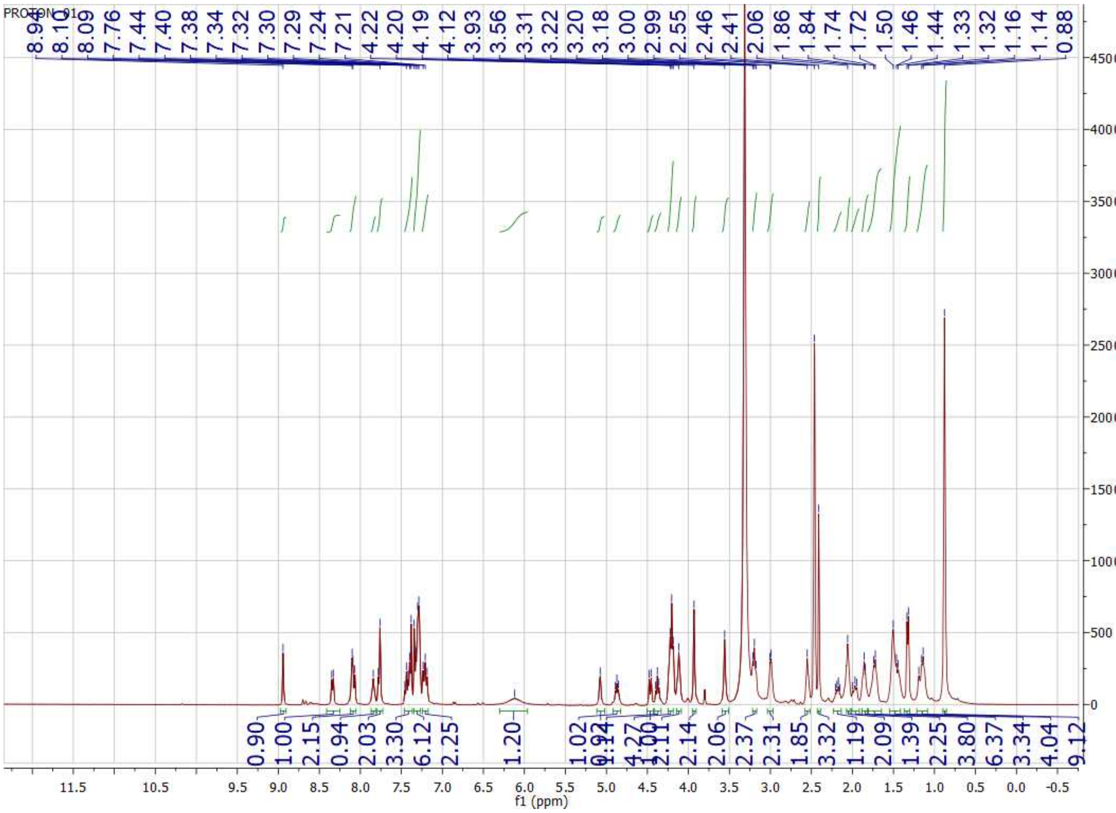

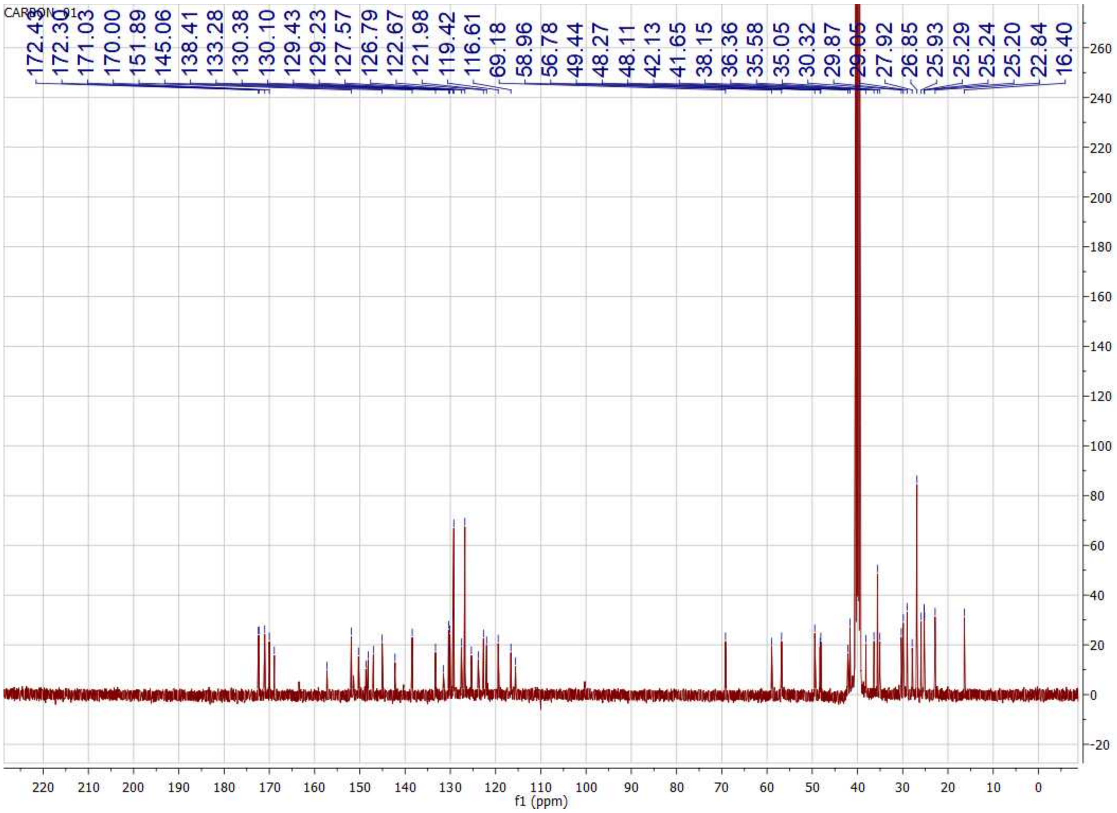
CARBON_01
172.30
171.03
170.00
151.89
145.06
138.41
133.28
130.38
130.10
129.43
129.23
127.57
126.79
122.67
121.98
119.42
116.61
69.18
58.96
56.78
49.44
48.27
48.11
42.13
41.65
38.15
36.36
35.58
35.05
30.32
29.87
27.92
26.85
25.93
25.29
25.24
25.20
22.84
16.40
260
240
220
200
180
160
140
120
100
80
60
40
20
0
-20
220
210
200
190
180
170
160
150
140
130
120
110
100
90
80
70
60
50
40
30
20
10
0
f1 (ppm)

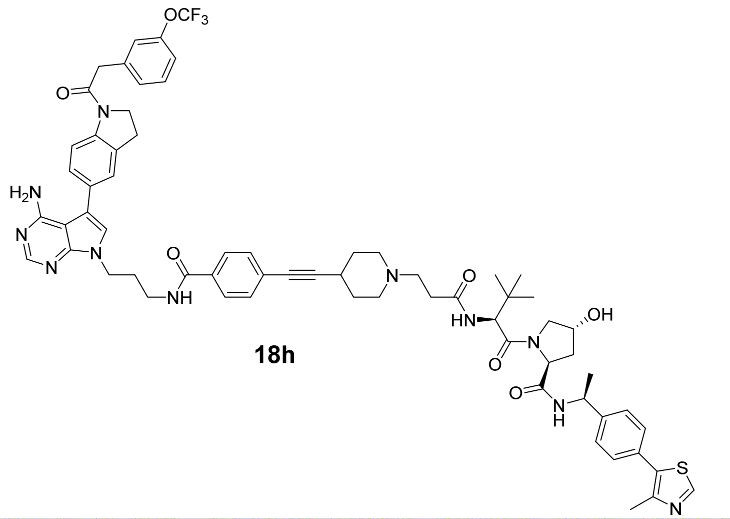
OCF3
O
N
H2N
N
N
N
O
NH
N
O
HN
O
N
OH
O
N
H
S
N

**18h**

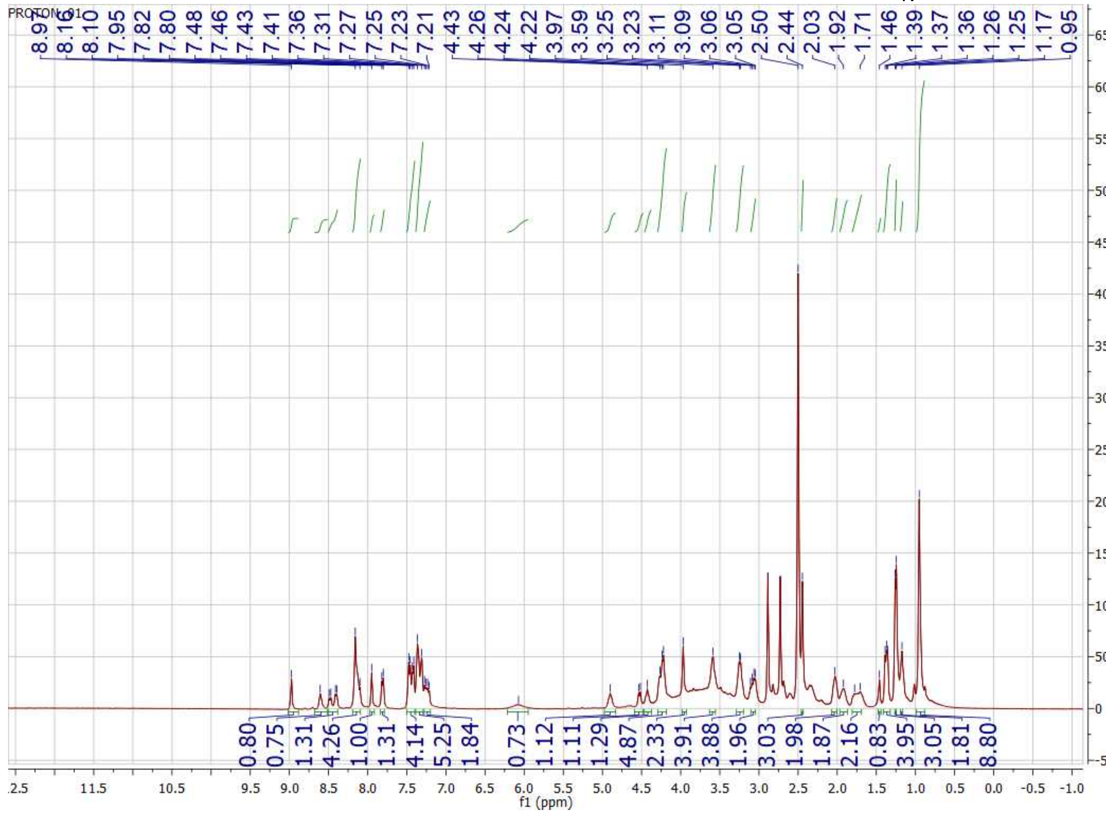
PROTON_01
8.97 8.16 8.10 7.95 7.82 7.80 7.48 7.46 7.43 7.41 7.36 7.31 7.27 7.25 7.23 7.21 4.43 4.26 4.24 4.22 3.97 3.59 3.25 3.23 3.11 3.09 3.06 3.05 2.50 2.44 2.03 1.92 1.71 1.46 1.39 1.37 1.36 1.26 1.25 1.17 0.95
0.80 0.75 1.31 4.26 1.00 1.31 4.14 5.25 1.84 0.73 1.12 1.11 1.29 4.87 2.33 3.91 3.88 1.96 3.03 1.98 1.87 2.16 0.83 3.95 3.05 1.81 8.80
f1 (ppm)

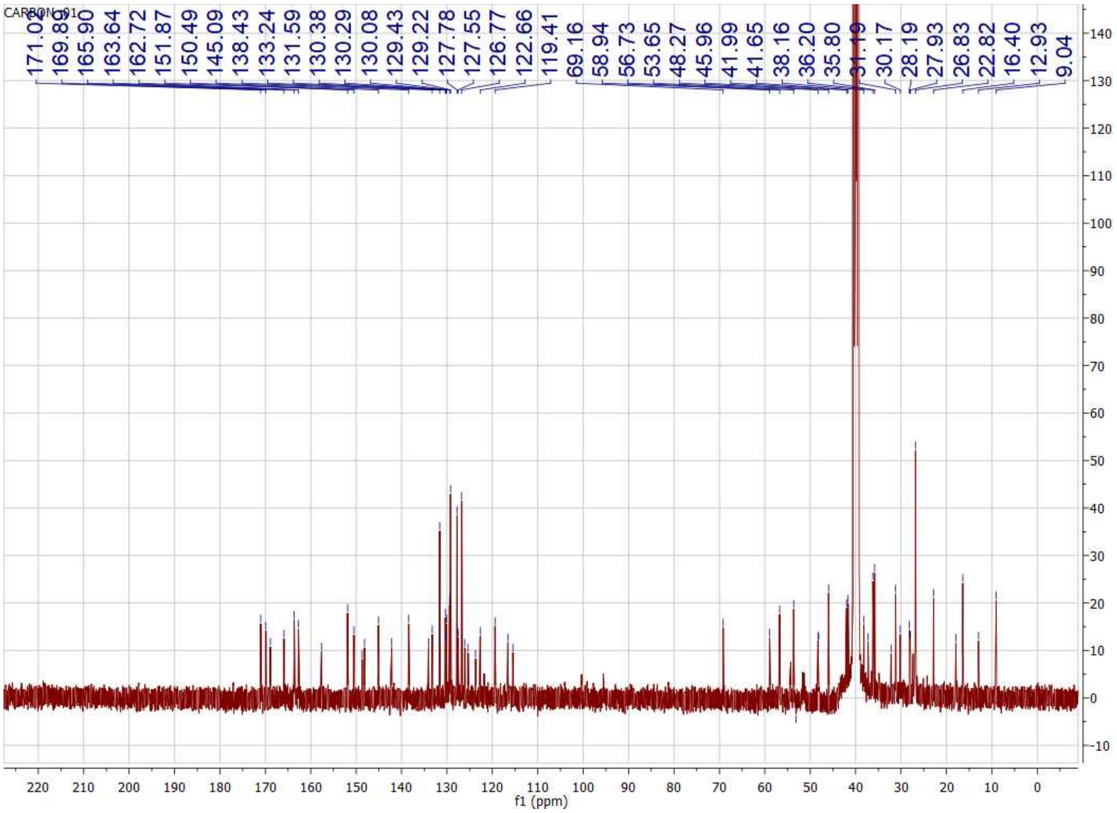
171.02
169.89
165.90
163.64
162.72
151.87
150.49
145.09
138.43
133.24
131.59
130.38
130.29
130.08
129.43
129.22
127.78
127.55
126.77
122.66
119.41
69.16
58.94
56.73
53.65
48.27
45.96
41.99
41.65
38.16
36.20
35.80
30.17
28.19
27.93
26.83
22.82
16.40
12.93
9.04
f1 (ppm)

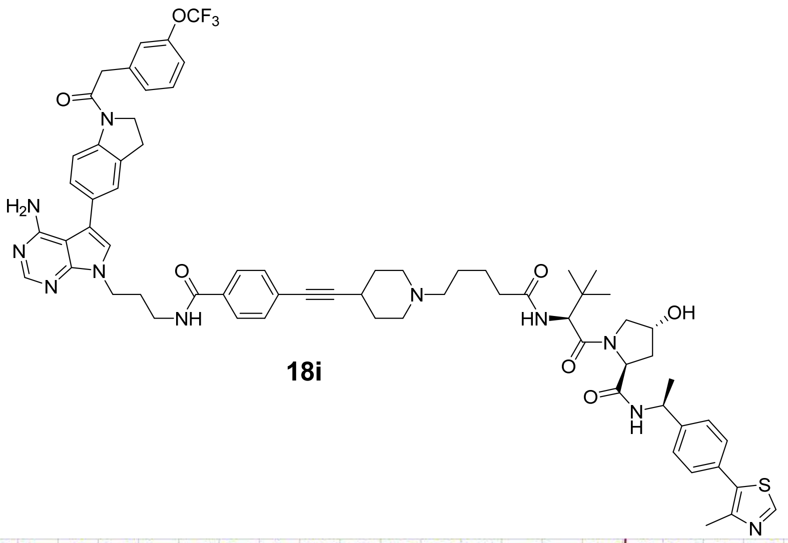
OCF3
O
N
H2N
N
N
N
O
NH
N
O
HN
O
N
OH
O
N
H
S
N

**18i**

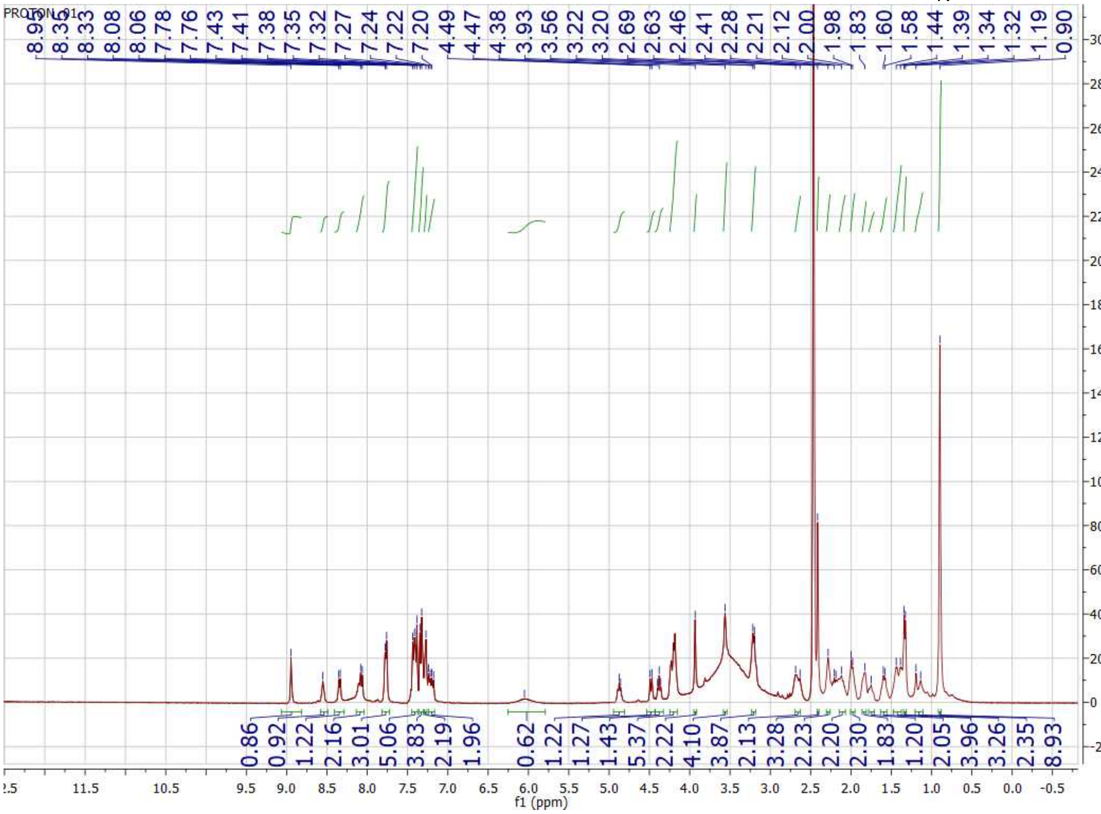
PROTON_01
8.33
8.08
8.06
7.78
7.76
7.43
7.41
7.38
7.35
7.32
7.27
7.24
7.22
7.20
4.49
4.47
4.38
3.93
3.56
3.22
3.20
2.69
2.63
2.46
2.41
2.28
2.21
2.12
2.00
1.98
1.83
1.60
1.58
1.44
1.39
1.34
1.32
1.19
0.90
0.86
0.92
1.22
2.16
3.01
5.06
3.83
2.19
1.96
0.62
1.22
1.27
1.43
5.37
2.22
4.10
3.87
2.13
3.28
2.23
2.20
2.30
1.83
1.20
2.05
3.96
3.26
2.35
8.93
f1 (ppm)

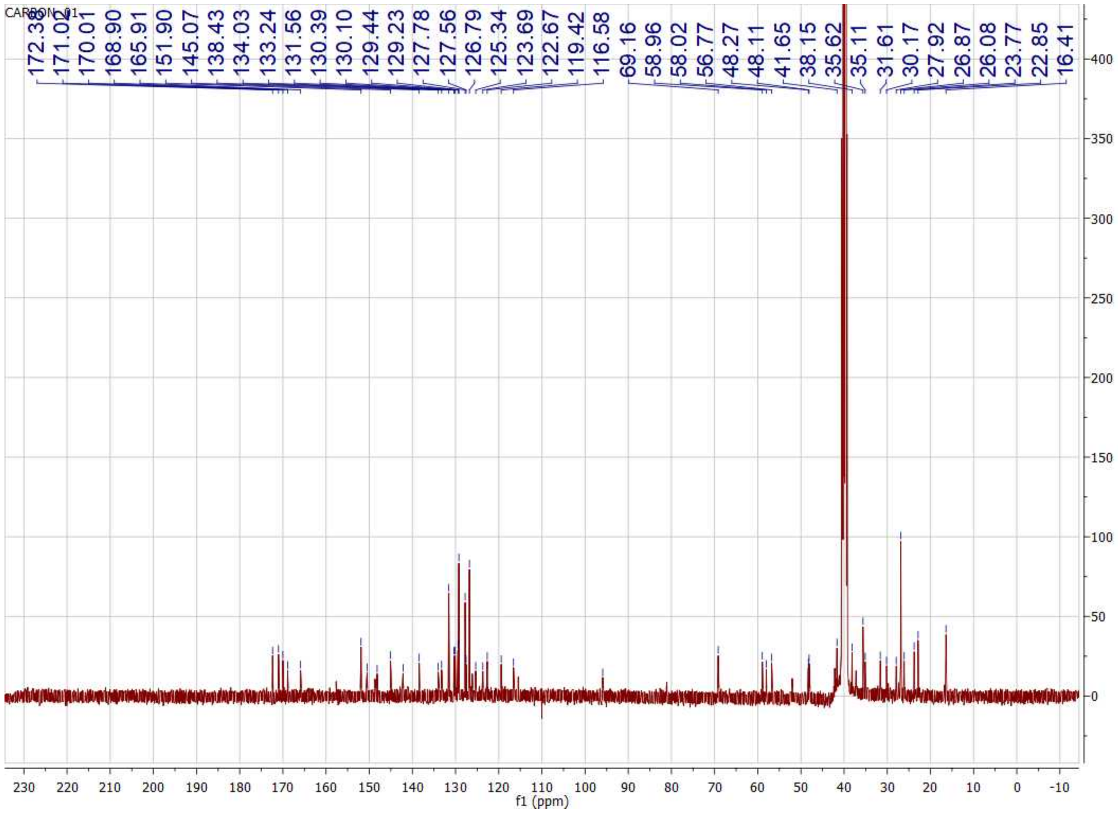
CARBON_01
172.30
171.02
170.01
168.90
165.91
151.90
145.07
138.43
134.03
133.24
131.56
130.39
130.10
129.44
129.23
127.78
127.56
126.79
125.34
123.69
122.67
119.42
116.58
69.16
58.96
58.02
56.77
48.27
48.11
41.65
38.15
35.62
35.11
31.61
30.17
27.92
26.87
26.08
23.77
22.85
16.41
f1 (ppm)

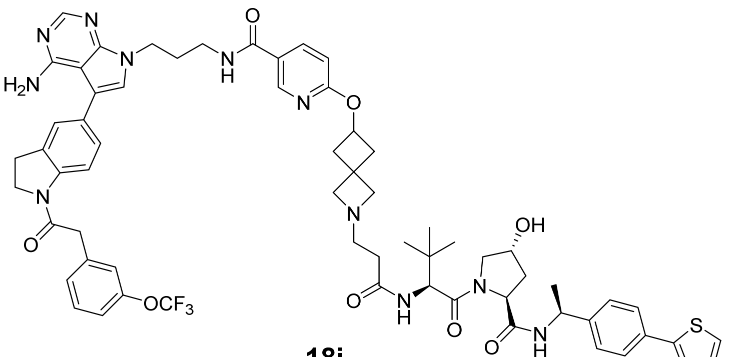

N
N
N
H2N
O
N
H
N
O
N
OCF3
O
N
O
N
H
O
N
OH
O
N
H
S
N

**18j**

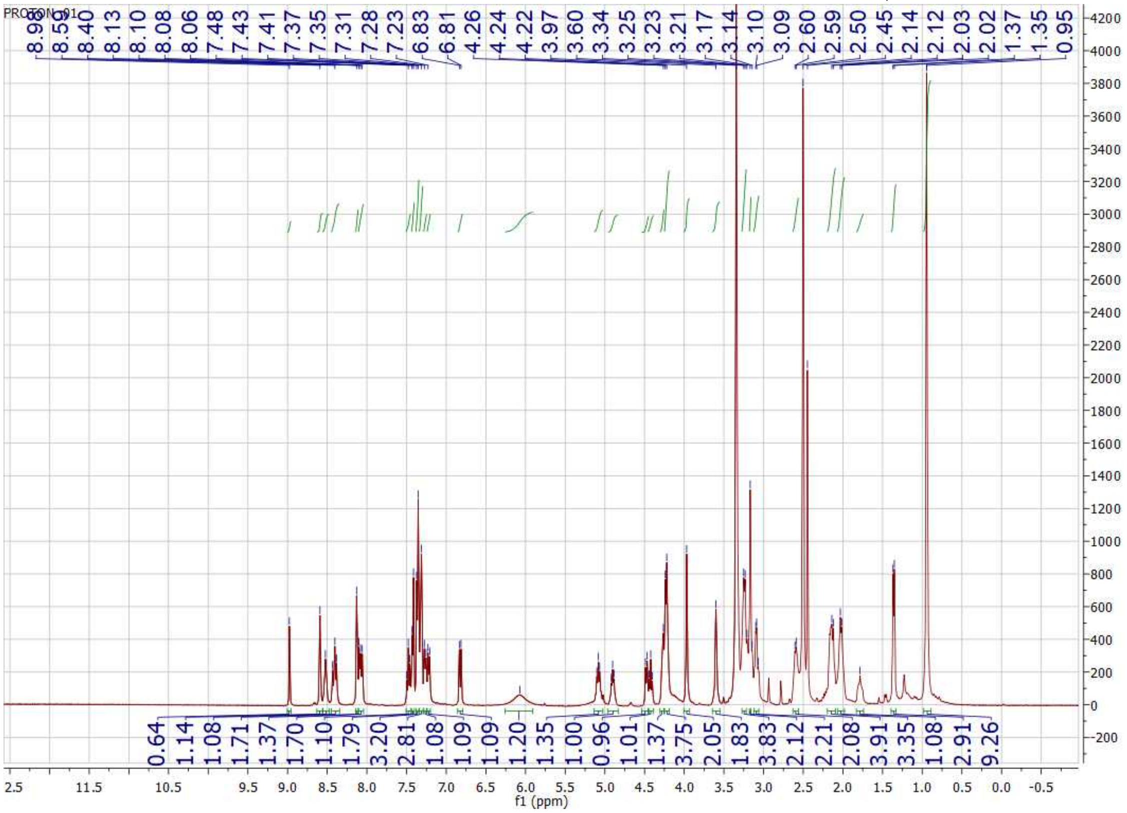

PROTON_01
8.98 8.59 8.40 8.13 8.10 8.08 8.06 7.48 7.43 7.41 7.37 7.35 7.31 7.28 7.23 6.83 6.81 4.26 4.24 4.22 3.97 3.60 3.34 3.25 3.23 3.21 3.17 3.14 3.10 3.09 2.60 2.59 2.50 2.45 2.14 2.12 2.03 2.02 1.37 1.35 0.95
0.64 1.14 1.08 1.71 1.37 1.70 1.10 1.79 3.20 2.81 2.08 1.08 1.09 1.09 1.20 1.35 1.00 0.96 0.96 1.01 1.37 1.37 3.75 2.05 1.83 3.83 2.12 2.21 2.08 3.91 3.35 1.08 2.91 9.26
f1 (ppm)

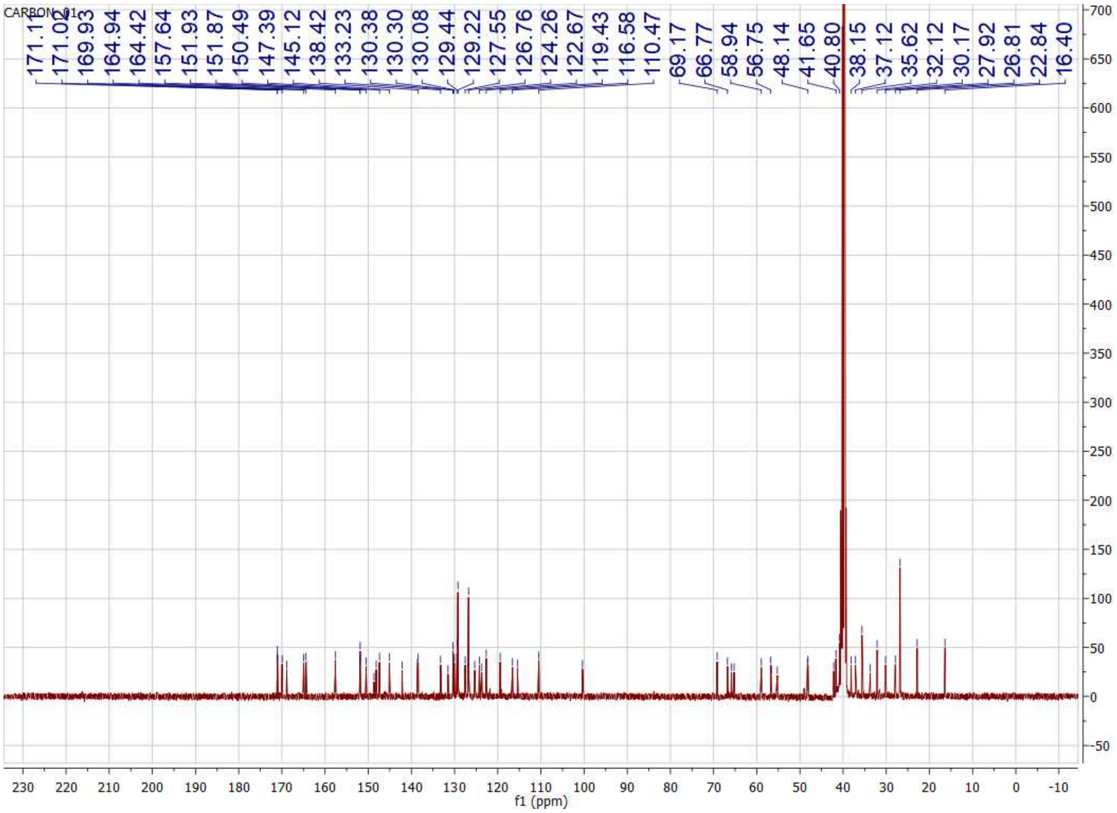

CARBON_01
171.11
171.02
169.93
164.94
164.42
157.64
151.93
151.87
150.49
147.39
145.12
138.42
133.23
130.38
130.30
130.08
129.44
129.22
127.55
126.76
124.26
122.67
119.43
116.58
110.47
69.17
66.77
58.94
56.75
48.14
41.65
40.80
38.15
37.12
35.62
32.12
30.17
27.92
26.81
22.84
16.40
f1 (ppm)

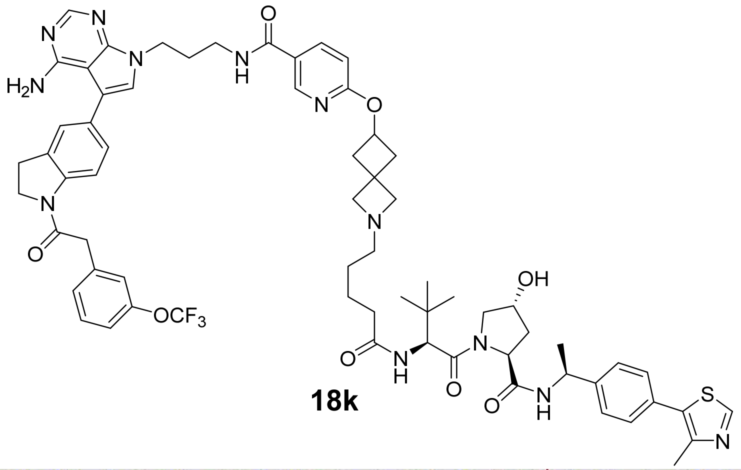

**18k**

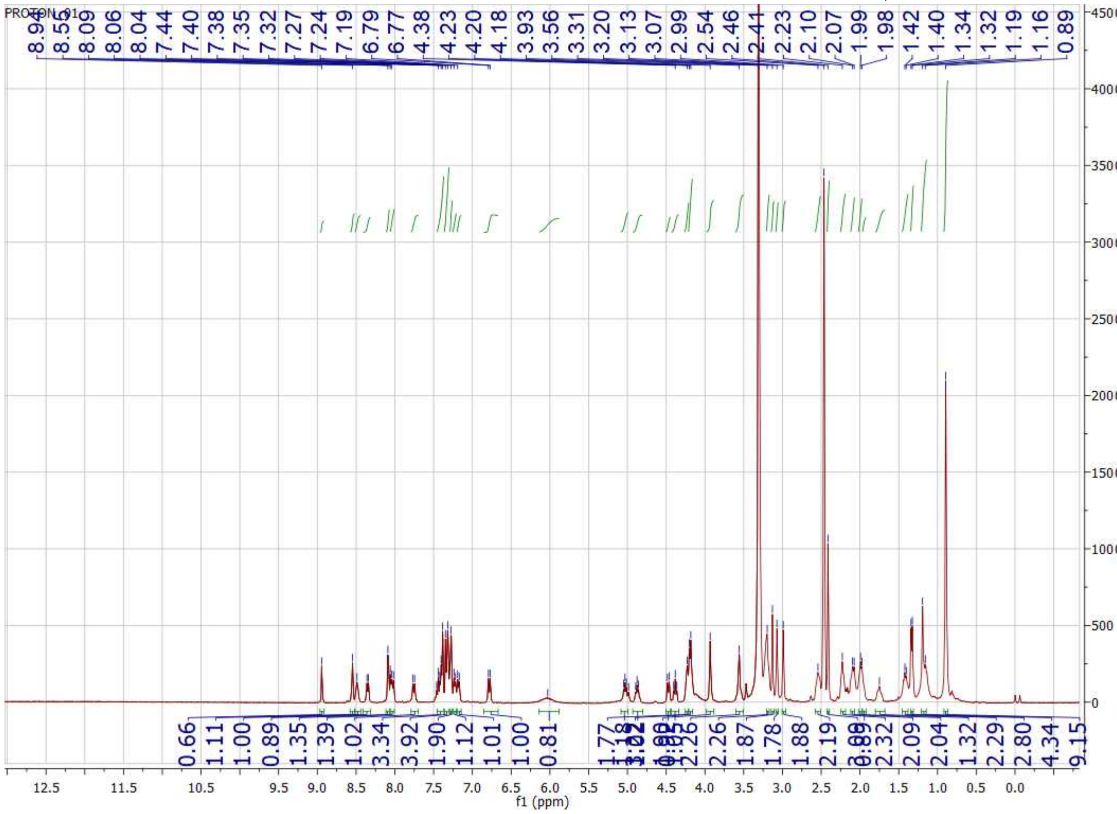

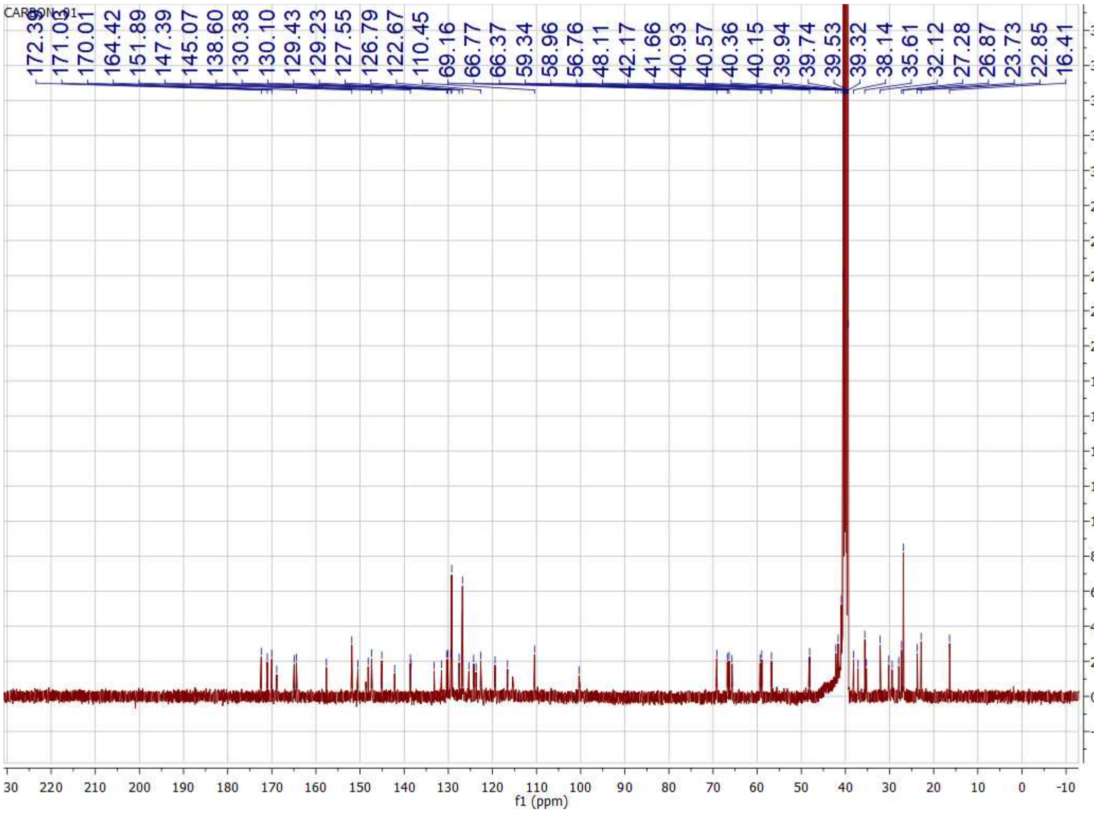
CARBON_01
172.35
171.03
170.01
164.42
151.89
147.39
145.07
138.60
130.38
130.10
129.43
129.23
127.55
126.79
122.67
110.45
69.16
66.77
66.37
59.34
58.96
56.76
48.11
42.17
41.66
40.93
40.57
40.36
40.15
39.94
39.74
39.53
39.32
38.14
35.61
32.12
27.28
26.87
23.73
22.85
16.41
f1 (ppm)

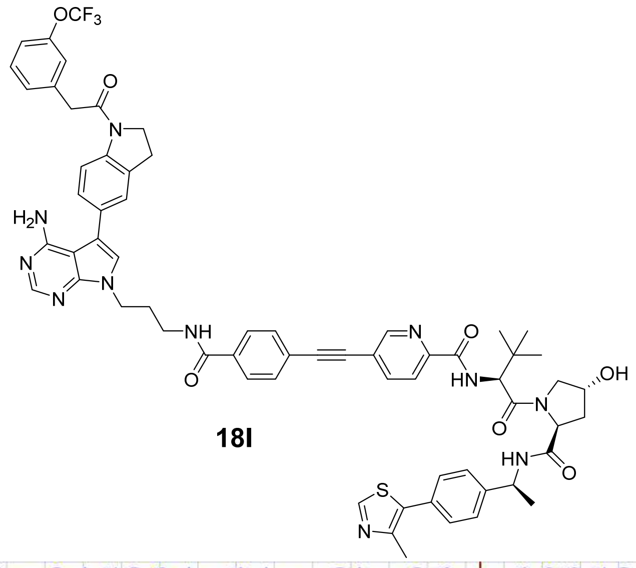

OCF3
O
N
H2N
N
N
N
NH
O
N
O
HN
O
N
OH
HN
O
S
N

**18l**

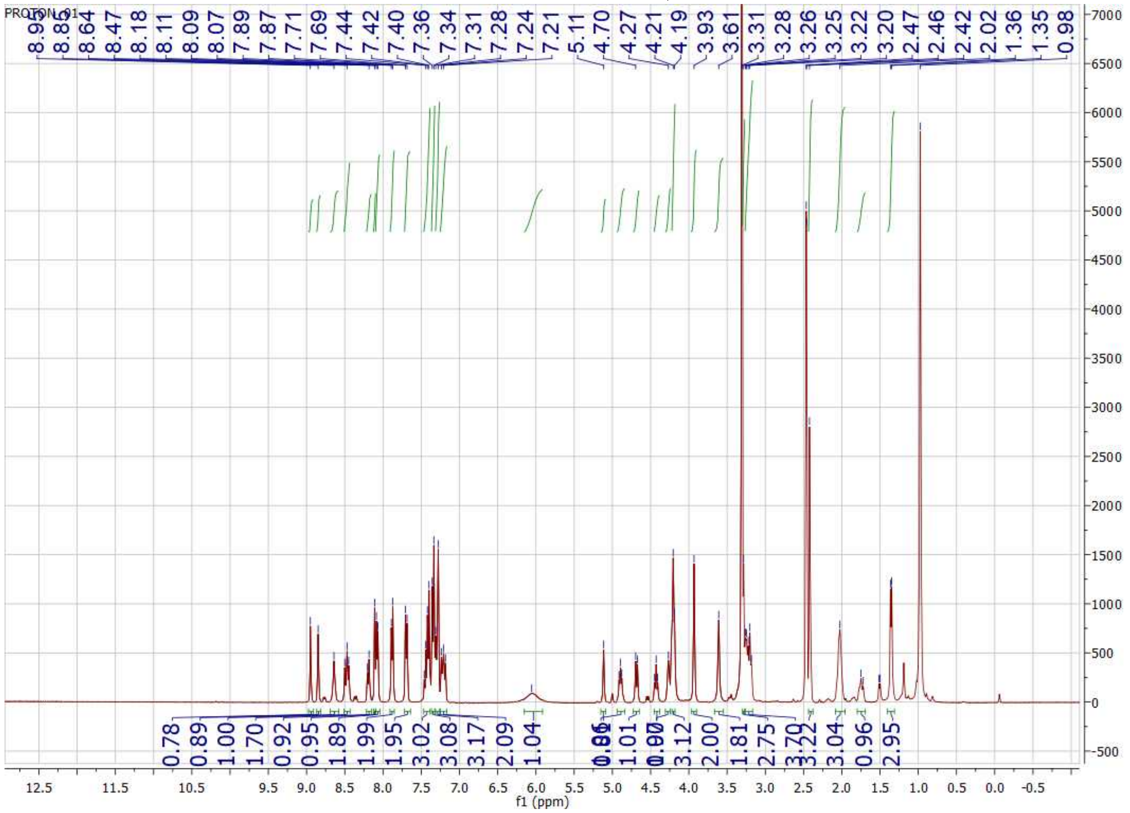

PROTON_01
8.95 [illegible] 8.88 8.64 8.47 8.18 8.11 8.09 8.07 7.89 7.87 7.71 7.69 7.44 7.42 7.40 7.36 7.34 7.31 7.28 7.24 7.21 5.11 4.70 4.27 4.21 4.19 3.93 3.61 3.31 3.28 3.26 3.25 3.22 3.20 2.47 2.46 2.42 2.02 1.36 1.35 0.98
0.78 0.89 1.00 1.70 0.92 0.95 1.89 1.99 1.95 3.02 3.08 3.17 2.09 1.04 [illegible] 1.01 [illegible] 3.12 2.00 1.81 2.75 3.70 3.22 3.04 0.96 2.95
f1 (ppm)

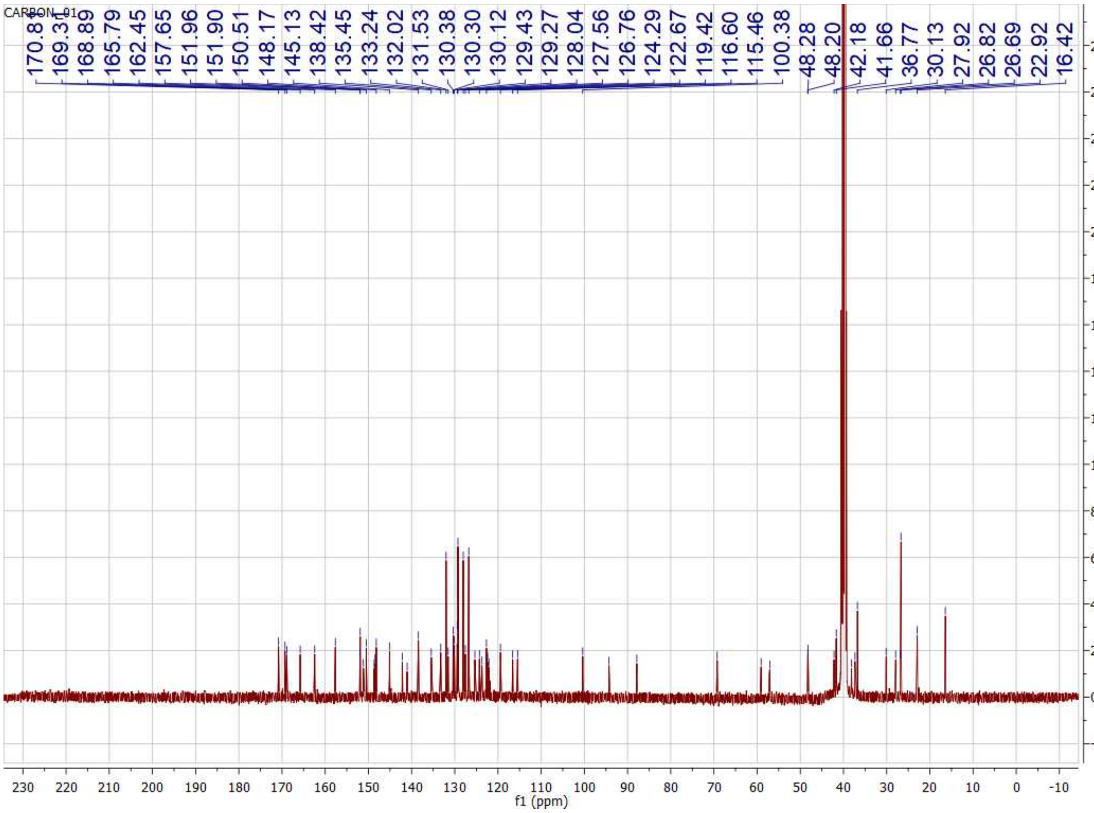

CARBON_01
170.8
169.31
168.89
165.79
162.45
157.65
151.96
151.90
150.51
148.17
145.13
138.42
135.45
133.24
132.02
131.53
130.38
130.30
130.12
129.43
129.27
128.04
127.56
126.76
124.29
122.67
119.42
116.60
115.46
100.38
48.28
48.20
42.18
41.66
36.77
30.13
27.92
26.82
26.69
22.92
16.42
f1 (ppm)

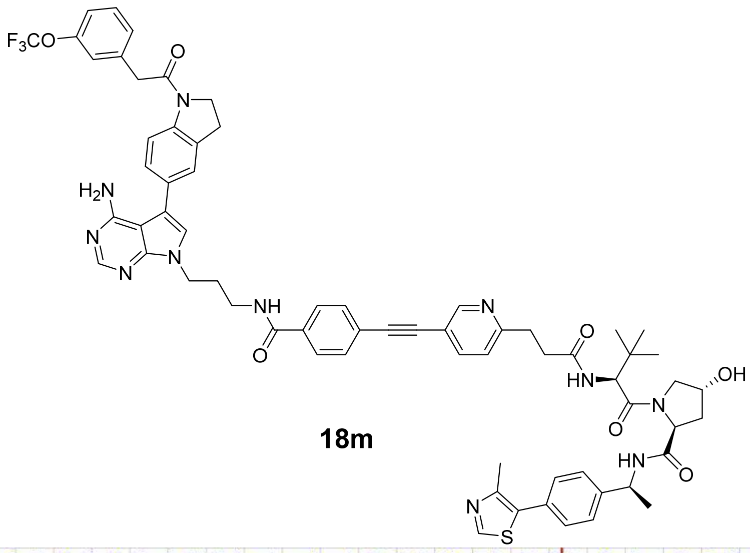
F3CO
O
N
H2N
N
N
N
NH
O
N
O
HN
OH
N
O
HN
O
N
S

**18m**

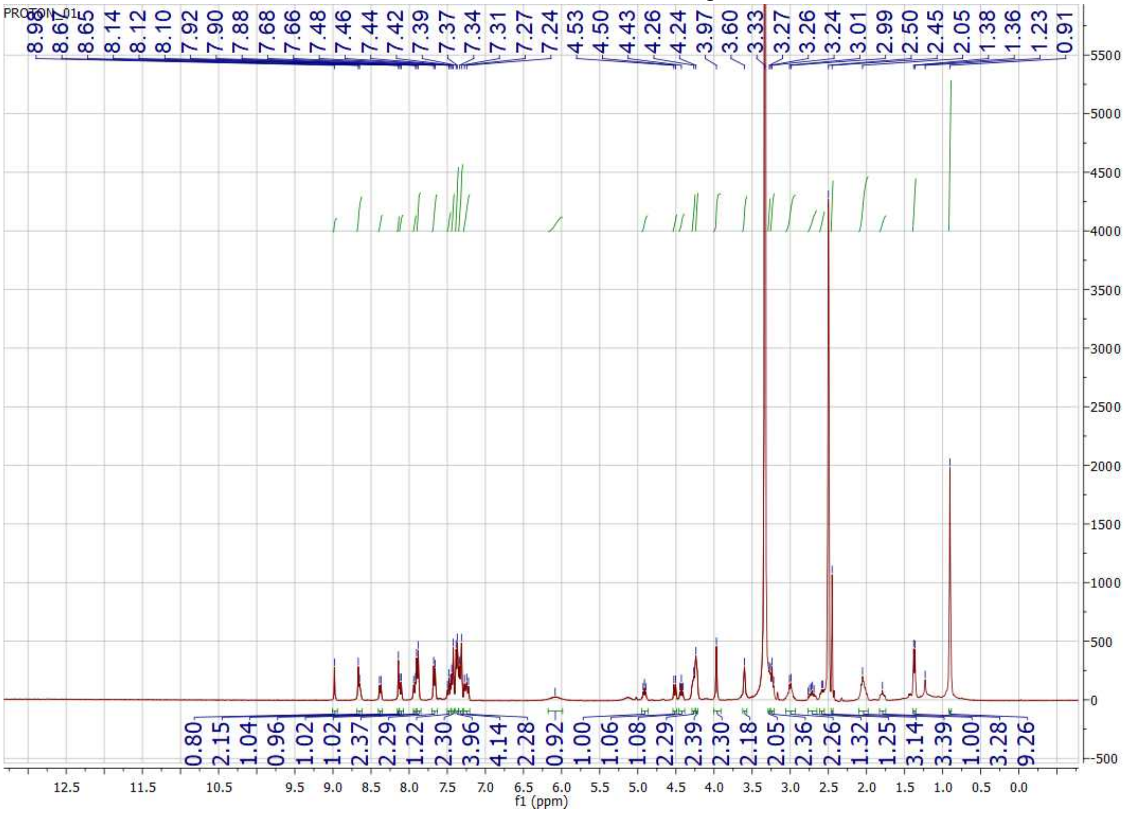
PROTON_01
8.99 8.67 8.65 8.14 8.12 8.10 7.92 7.90 7.88 7.68 7.66 7.48 7.46 7.44 7.42 7.39 7.37 7.34 7.31 7.27 7.24 4.53 4.50 4.43 4.26 4.24 3.97 3.60 3.33 3.27 3.26 3.24 3.01 2.99 2.50 2.45 2.05 1.38 1.36 1.23 0.91
0.80 2.15 1.04 0.96 1.02 1.02 2.37 2.29 1.22 2.30 3.96 4.14 2.28 0.92 1.00 1.06 1.08 2.29 2.39 2.30 2.18 2.05 2.36 2.26 1.32 1.25 3.14 3.39 1.00 3.28 9.26
f1 (ppm)

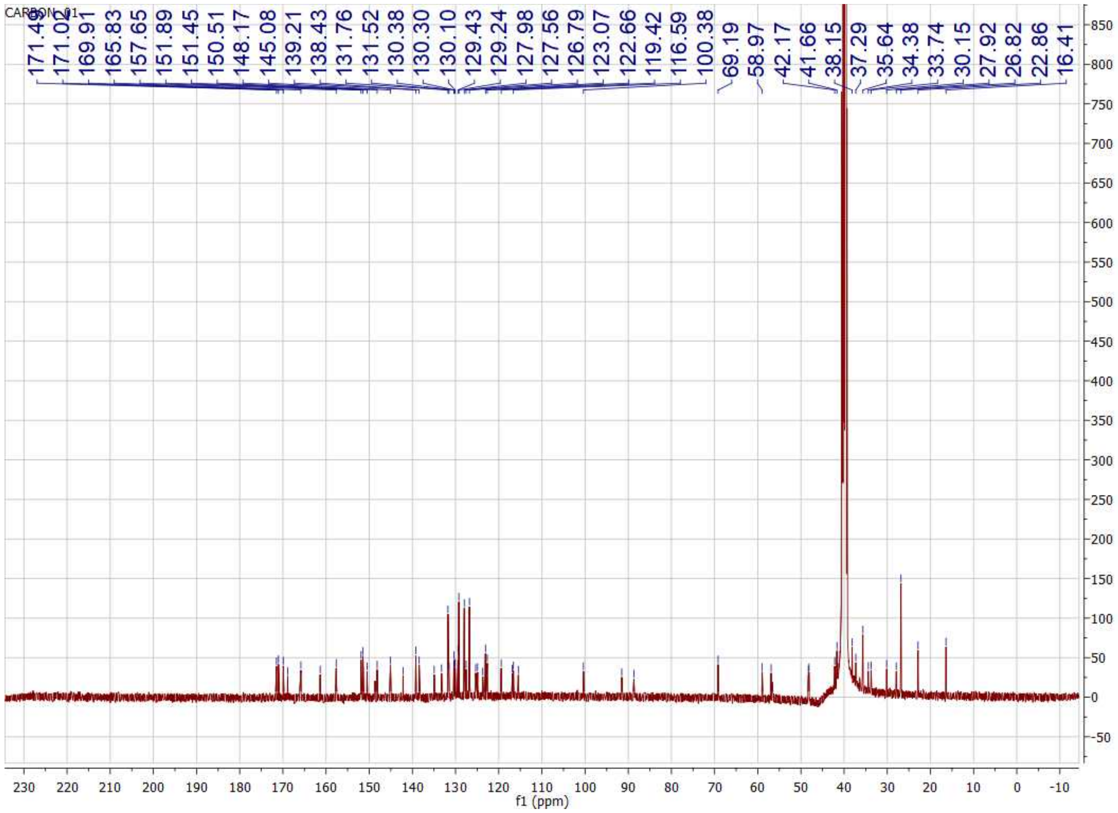
CARBON_01
171.49
171.02
169.91
165.83
157.65
151.89
151.45
150.51
148.17
145.08
139.21
138.43
131.76
131.52
130.38
130.30
130.10
129.43
129.24
127.98
127.56
126.79
123.07
122.66
119.42
116.59
100.38
69.19
58.97
42.17
41.66
38.15
37.29
35.64
34.38
33.74
30.15
27.92
26.82
22.86
16.41
f1 (ppm)

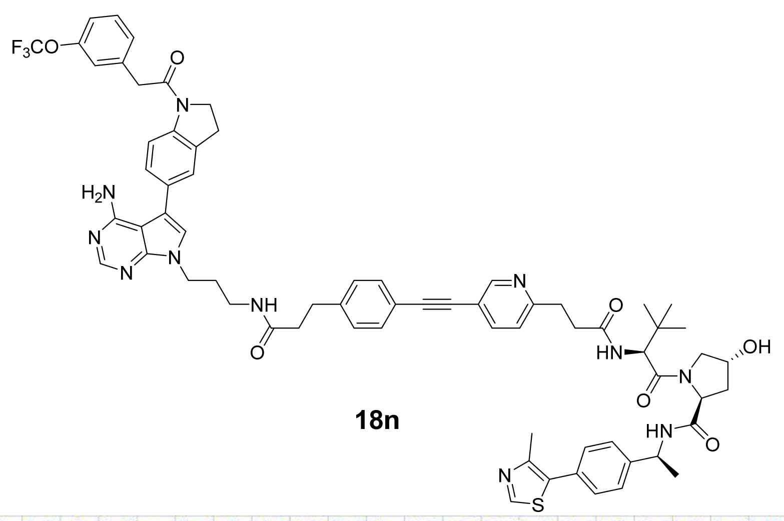

**18n**

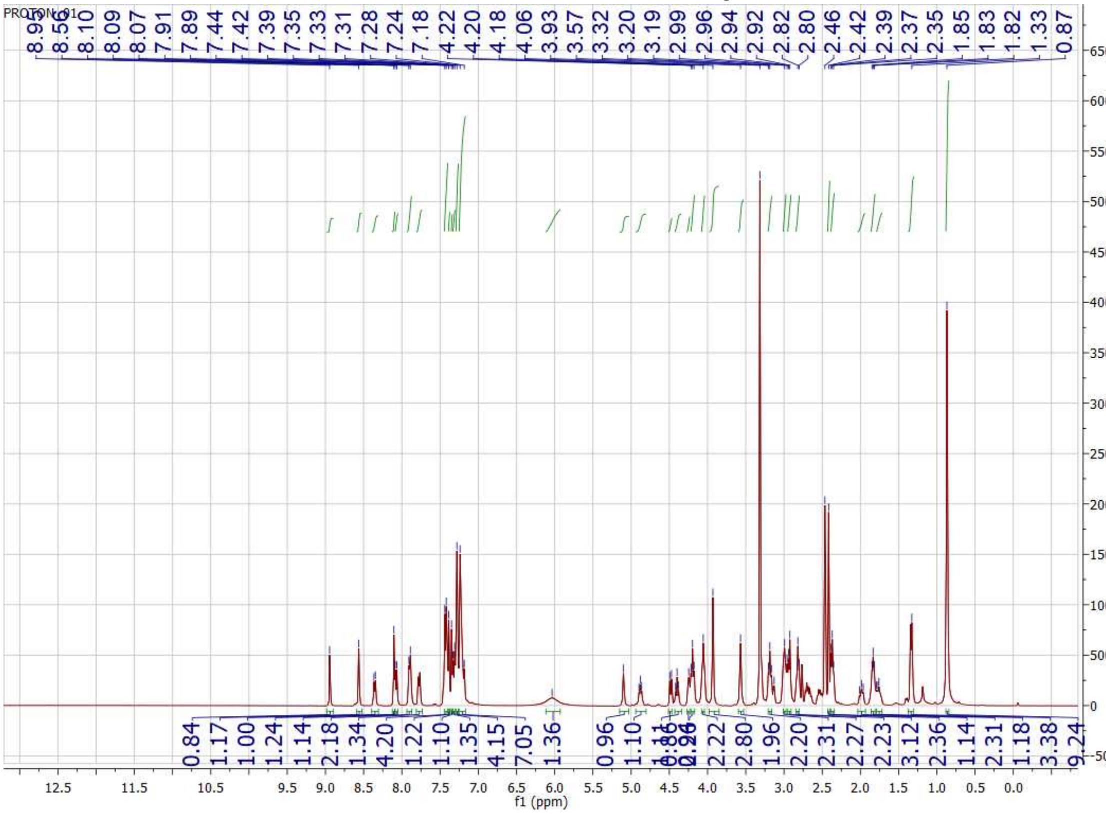

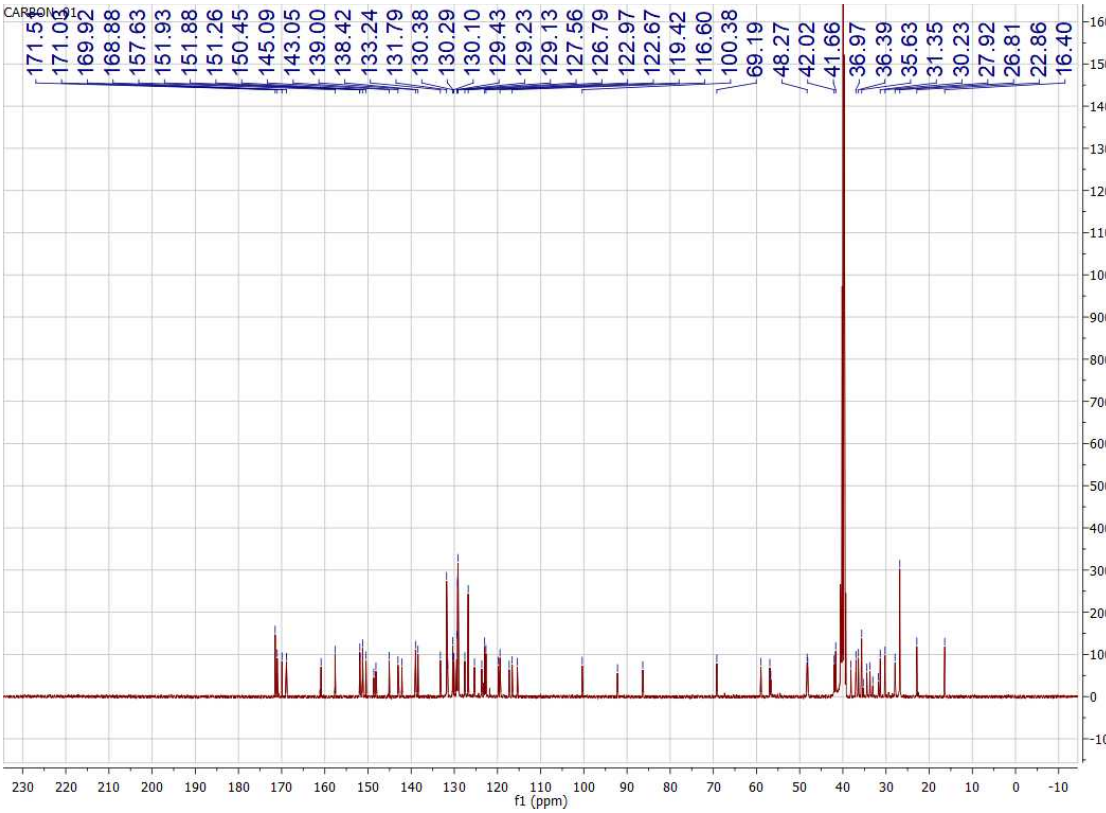
CARBON_01
171.54
171.03
169.92
168.88
157.63
151.93
151.88
151.26
150.45
145.09
143.05
139.00
138.42
133.24
131.79
130.38
130.29
130.10
129.43
129.23
129.13
127.56
126.79
122.97
122.67
119.42
116.60
100.38
69.19
48.27
42.02
41.66
36.97
36.39
35.63
31.35
30.23
27.92
26.81
22.86
16.40
f1 (ppm)

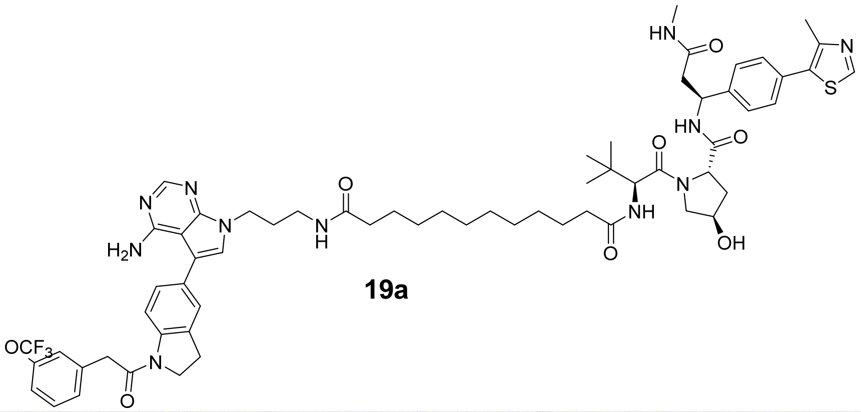

**19a**

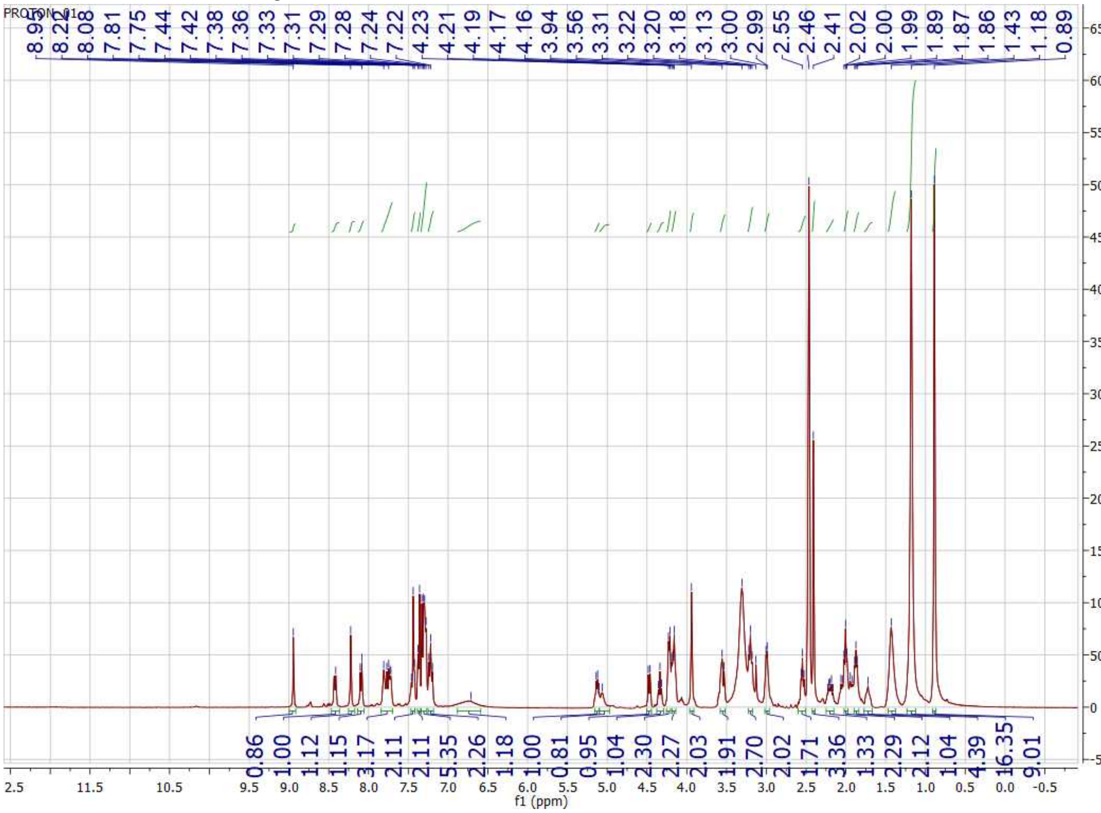

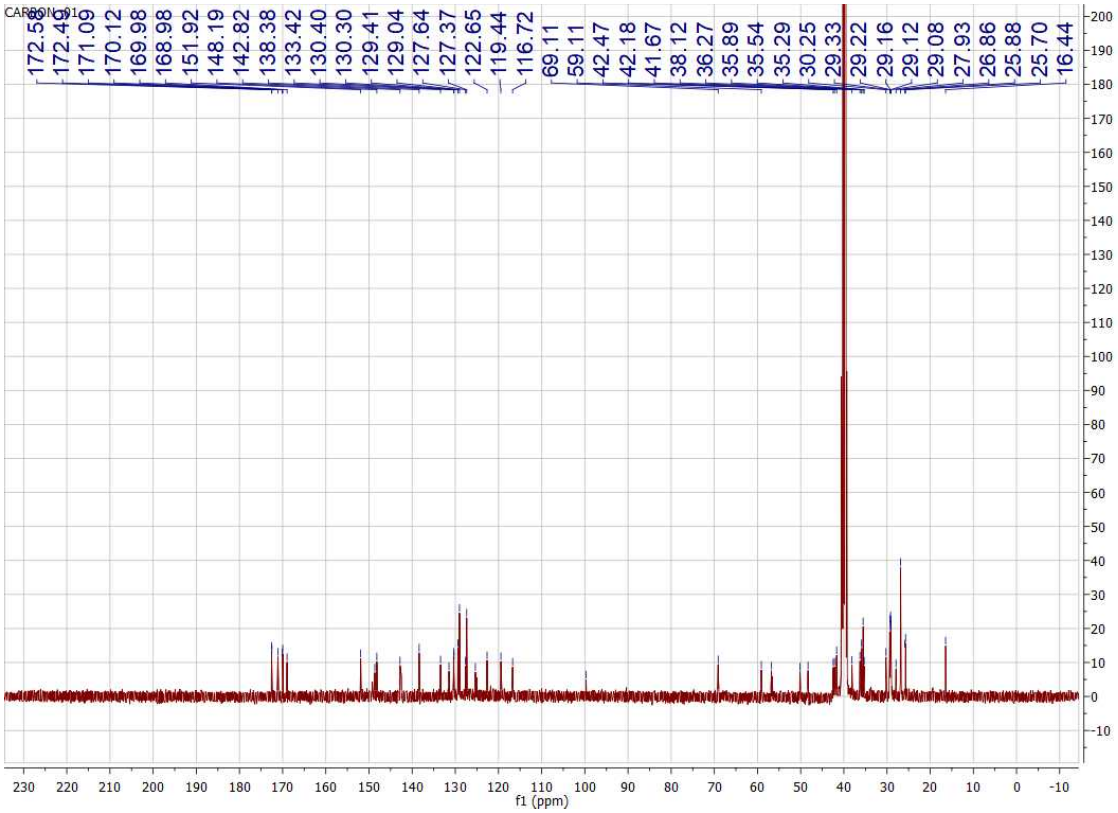

CARBON_01
172.56
172.49
171.09
170.12
169.98
168.98
151.92
148.19
142.82
138.38
133.42
130.40
130.30
129.41
129.04
127.64
127.37
122.65
119.44
116.72
69.11
59.11
42.47
42.18
41.67
38.12
36.27
35.89
35.54
35.29
30.25
29.33
29.22
29.16
29.12
29.08
27.93
26.86
25.88
25.70
16.44
f1 (ppm)

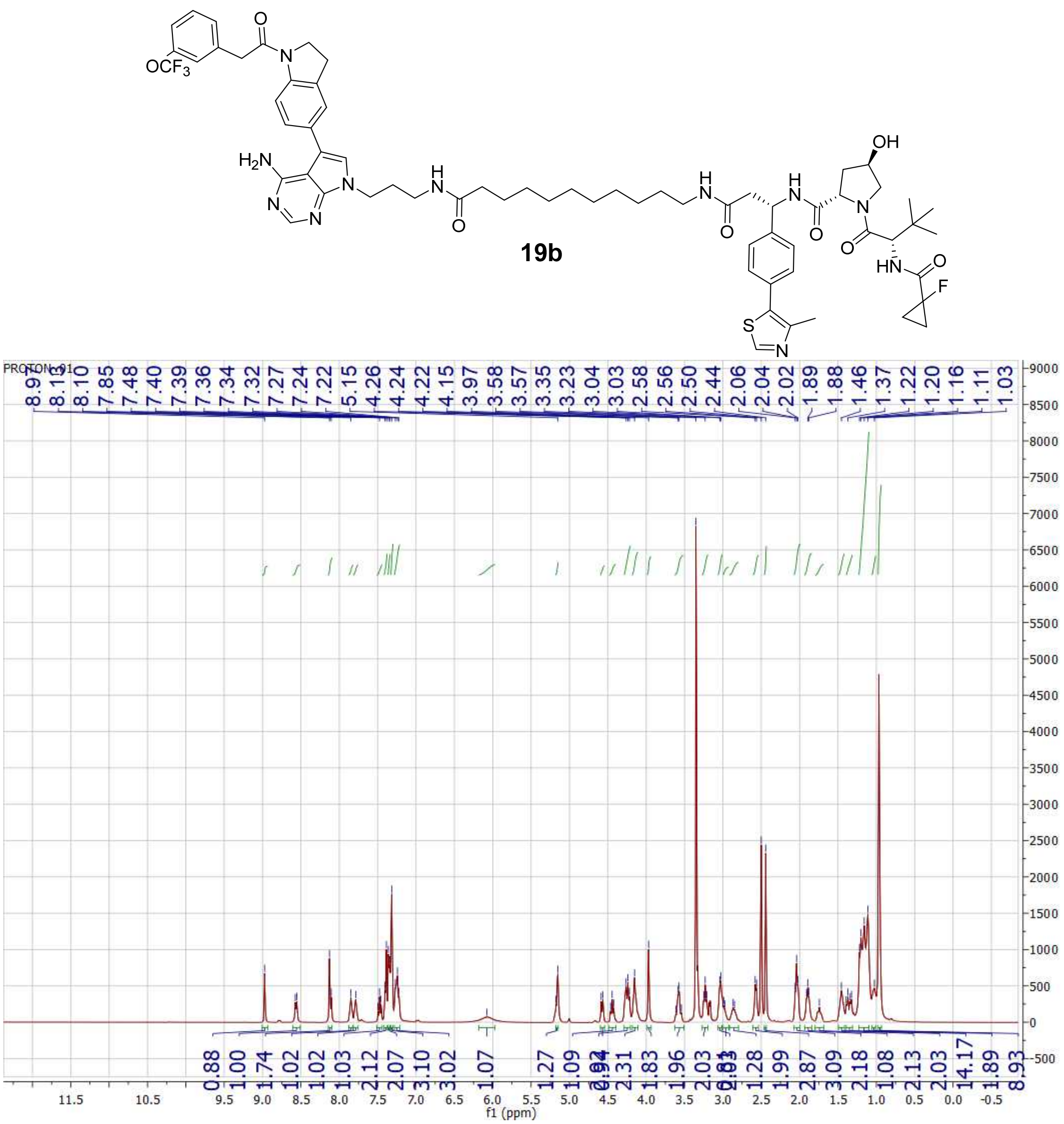

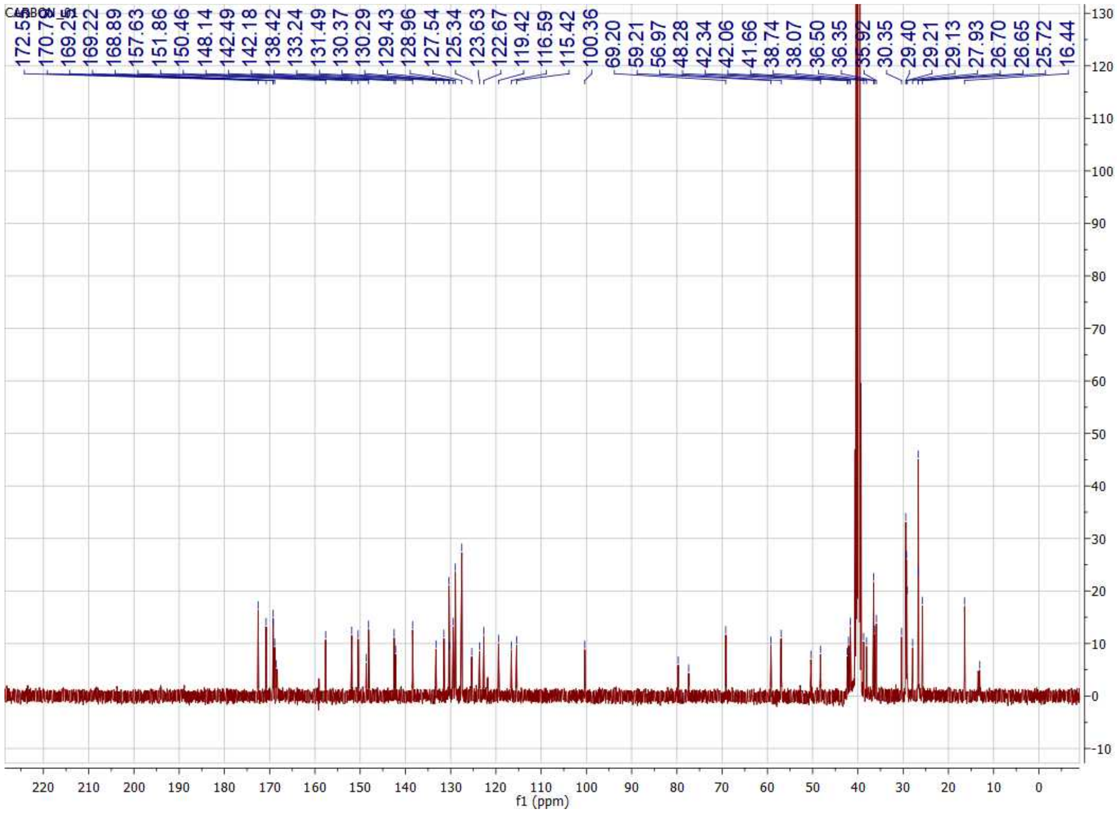

CARBON_01
172.53
170.75
169.26
169.22
168.89
157.63
151.86
150.46
148.14
142.49
142.18
138.42
133.24
131.49
130.37
130.29
129.43
128.96
127.54
125.34
123.63
122.67
119.42
116.59
115.42
100.36
69.20
59.21
56.97
48.28
42.34
42.06
41.66
38.74
38.07
36.50
36.35
35.92
30.35
29.40
29.21
29.13
27.93
26.70
26.65
25.72
16.44
f1 (ppm)

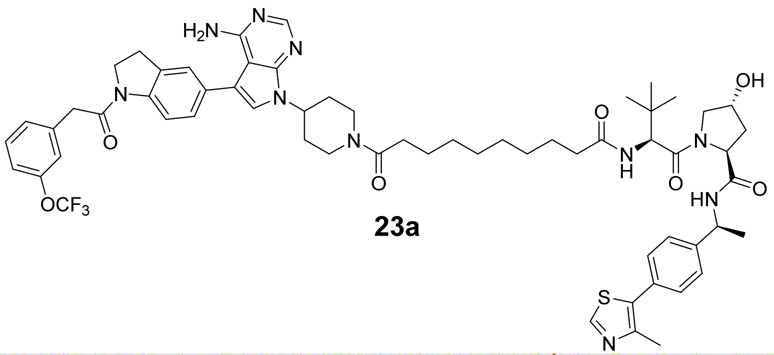
H2N
N
N
N
N
N
O
OCF3
N
O
O
N
H
O
N
OH
HN
O
S
N

**23a**

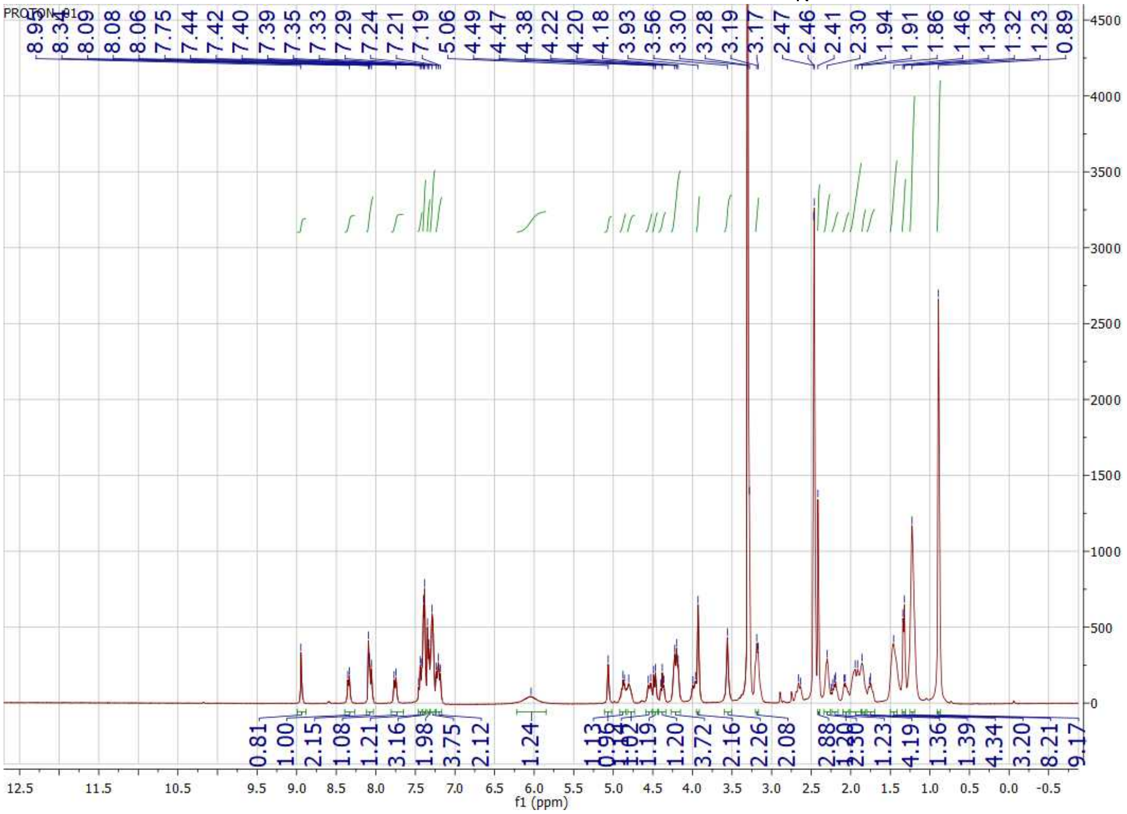
PROTON_01
8.95 8.34 8.09 8.08 8.06 7.75 7.44 7.42 7.40 7.39 7.35 7.33 7.29 7.24 7.21 7.19 5.06 4.49 4.47 4.38 4.22 4.20 4.18 3.93 3.56 3.30 3.28 3.19 3.17 2.47 2.46 2.41 2.30 1.94 1.91 1.86 1.46 1.34 1.32 1.23 0.89
f1 (ppm)

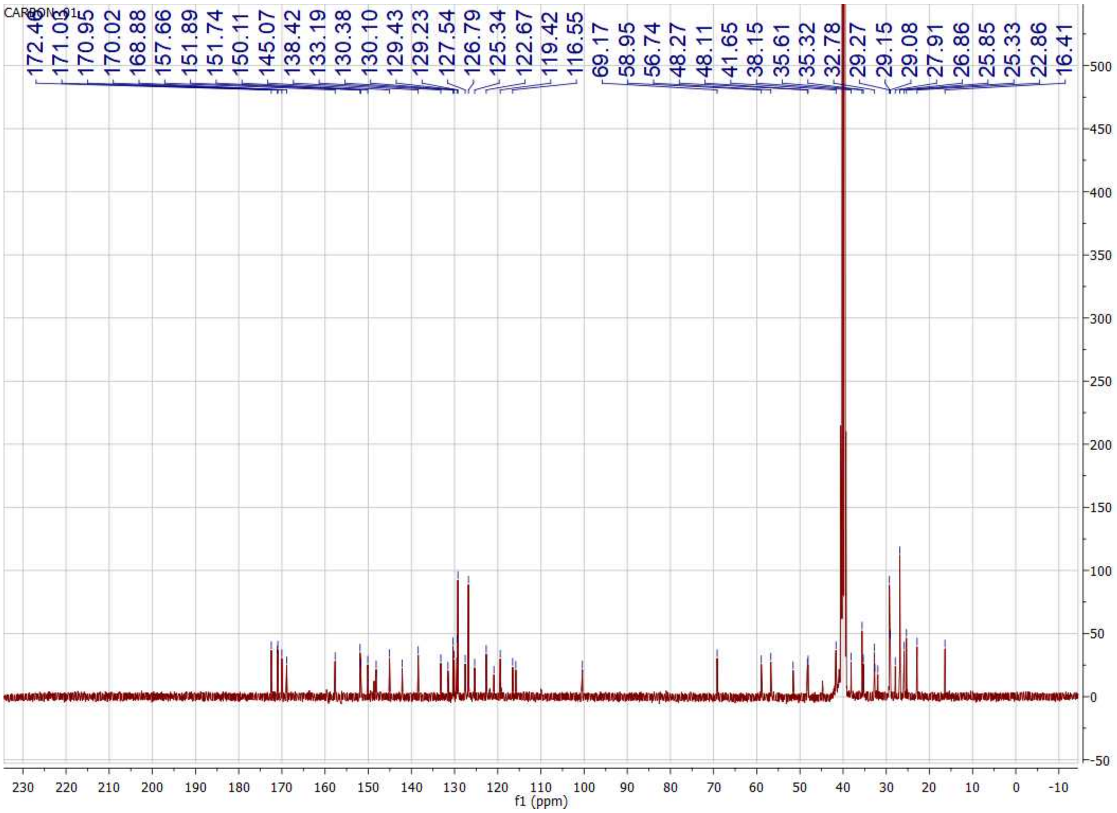
CARBON_01
172.40
171.03
170.95
170.02
168.88
157.66
151.89
151.74
150.11
145.07
138.42
133.19
130.38
130.10
129.43
129.23
127.54
126.79
125.34
122.67
119.42
116.55
69.17
58.95
56.74
48.27
48.11
41.65
38.15
35.61
35.32
32.78
29.27
29.15
29.08
27.91
26.86
25.85
25.33
22.86
16.41
f1 (ppm)

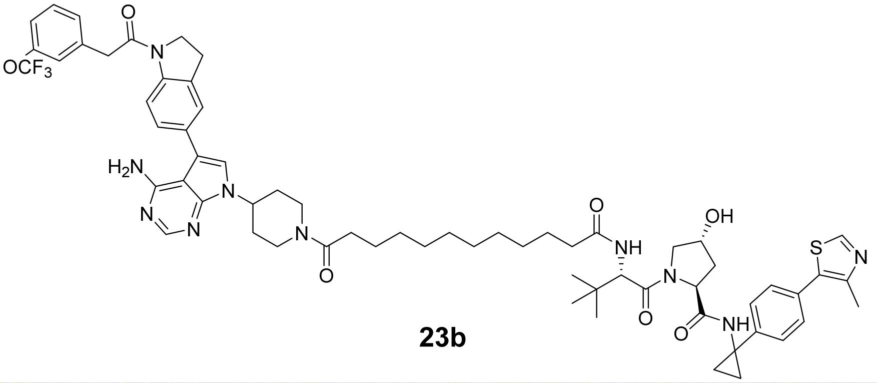

OCF3
H2N
N
N
N
N
O
O
O
NH
N
OH
O
O
NH
S
N

**23b**

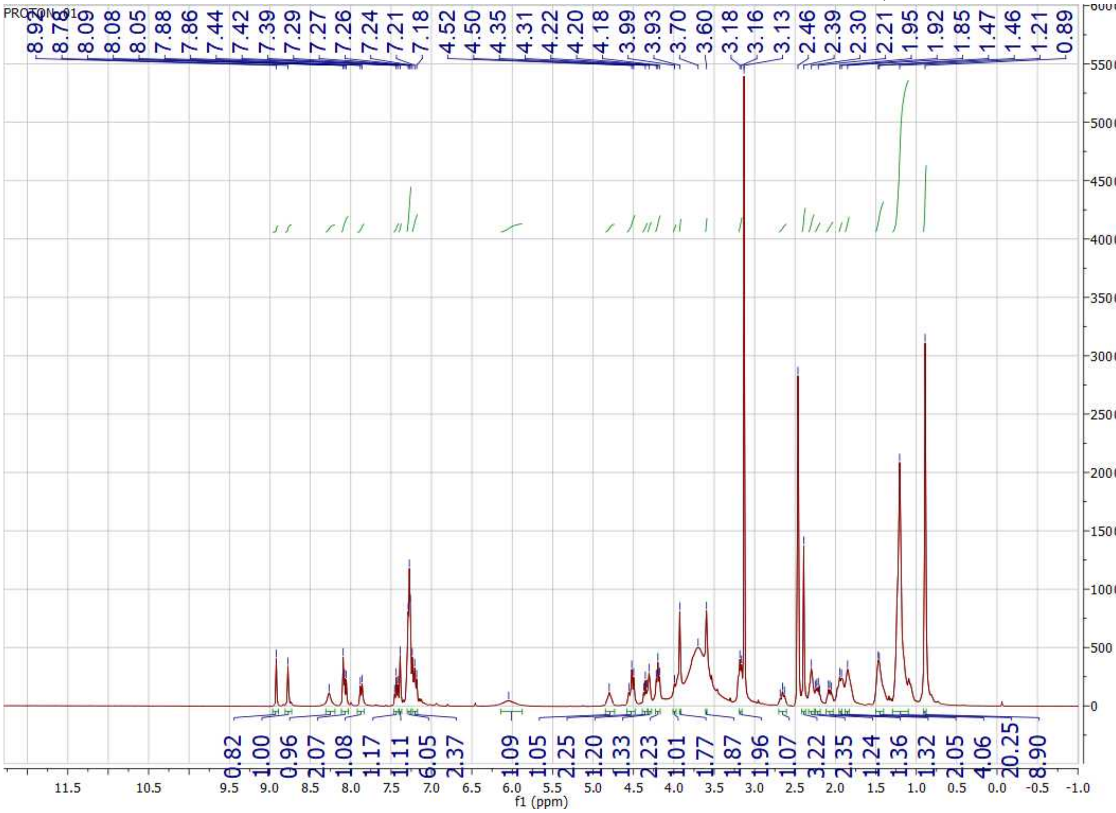

PROTON_01
8.92
8.78
8.09
8.08
8.05
7.88
7.86
7.44
7.42
7.39
7.29
7.27
7.26
7.24
7.21
7.18
4.52
4.50
4.35
4.31
4.22
4.20
4.18
3.99
3.93
3.70
3.60
3.18
3.16
3.13
2.46
2.39
2.30
2.21
1.95
1.92
1.85
1.47
1.46
1.21
0.89
0.82
1.00
0.96
2.07
1.08
1.17
1.11
6.05
2.37
1.09
1.05
2.25
1.20
1.33
2.23
1.01
1.77
1.87
1.96
1.07
3.22
2.35
1.24
1.36
1.32
2.05
4.06
20.25
8.90
f1 (ppm)

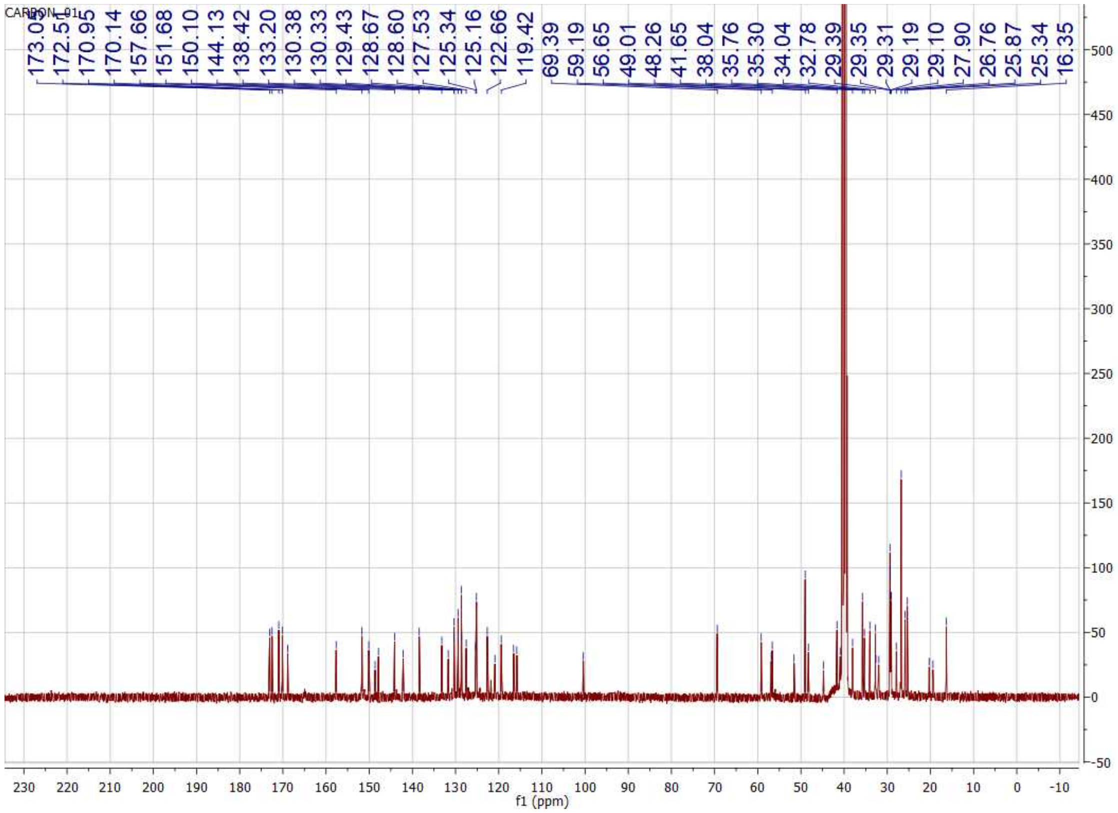
CARBON_01
173.03
172.51
170.95
170.14
157.66
151.68
150.10
144.13
138.42
133.20
130.38
130.33
129.43
128.67
128.60
127.53
125.34
125.16
122.66
119.42
69.39
59.19
56.65
49.01
48.26
41.65
38.04
35.76
35.30
34.04
32.78
29.39
29.35
29.31
29.19
29.10
27.90
26.76
25.87
25.34
16.35
f1 (ppm)

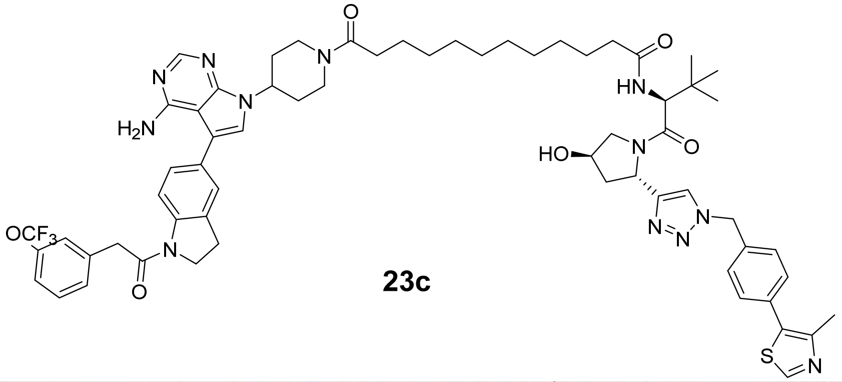

**23c**

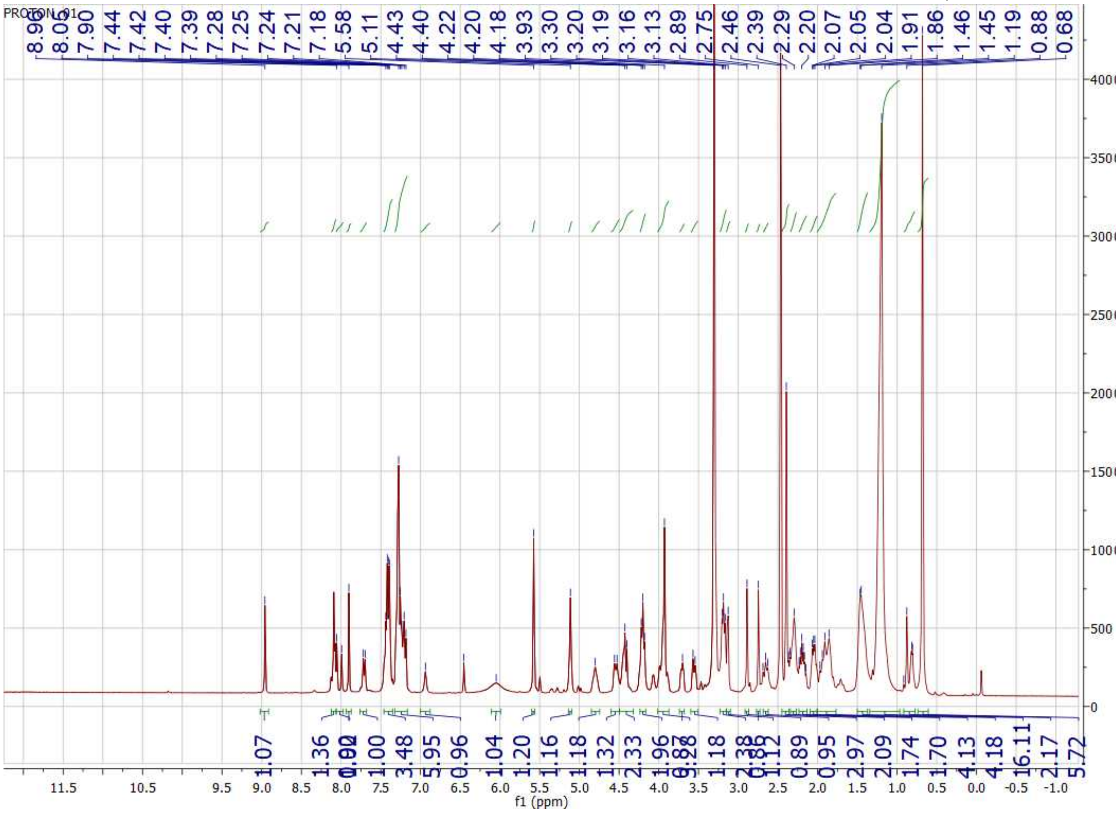

PROTON_01
8.96 8.05 7.90 7.44 7.42 7.40 7.39 7.28 7.25 7.24 7.21 7.18 5.58 5.11 4.43 4.40 4.22 4.20 4.18 3.93 3.30 3.20 3.19 3.16 3.13 2.89 2.75 2.46 2.39 2.29 2.20 2.07 2.05 2.04 1.91 1.86 1.46 1.45 1.19 0.88 0.68
1.07 1.36 1.00 3.48 5.95 0.96 1.04 1.20 1.16 1.18 1.32 2.33 1.96 1.18 1.12 0.89 0.95 2.97 2.09 1.74 1.70 4.13 4.18 16.11 2.17 5.72
f1 (ppm)

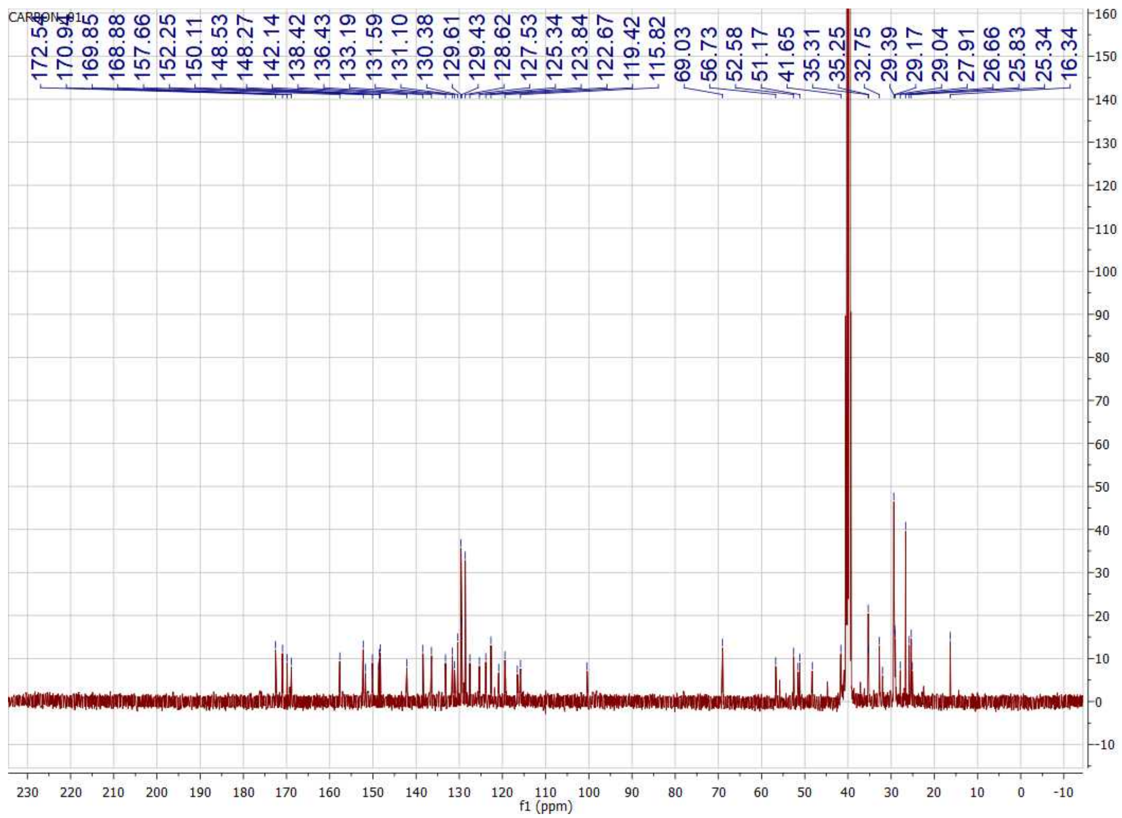
CARBON_01
172.54
170.94
169.85
168.88
157.66
152.25
150.11
148.53
148.27
142.14
138.42
136.43
133.19
131.59
131.10
130.38
129.61
129.43
128.62
127.53
125.34
123.84
122.67
119.42
115.82
69.03
56.73
52.58
51.17
41.65
35.31
35.25
32.75
29.39
29.17
29.04
27.91
26.66
25.83
25.34
16.34
f1 (ppm)

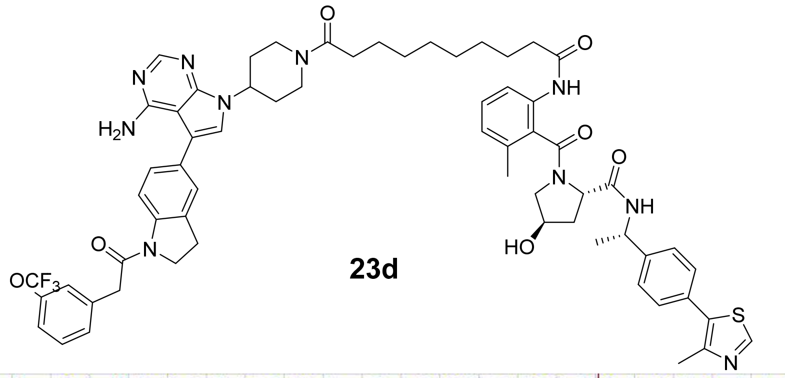

**23d**

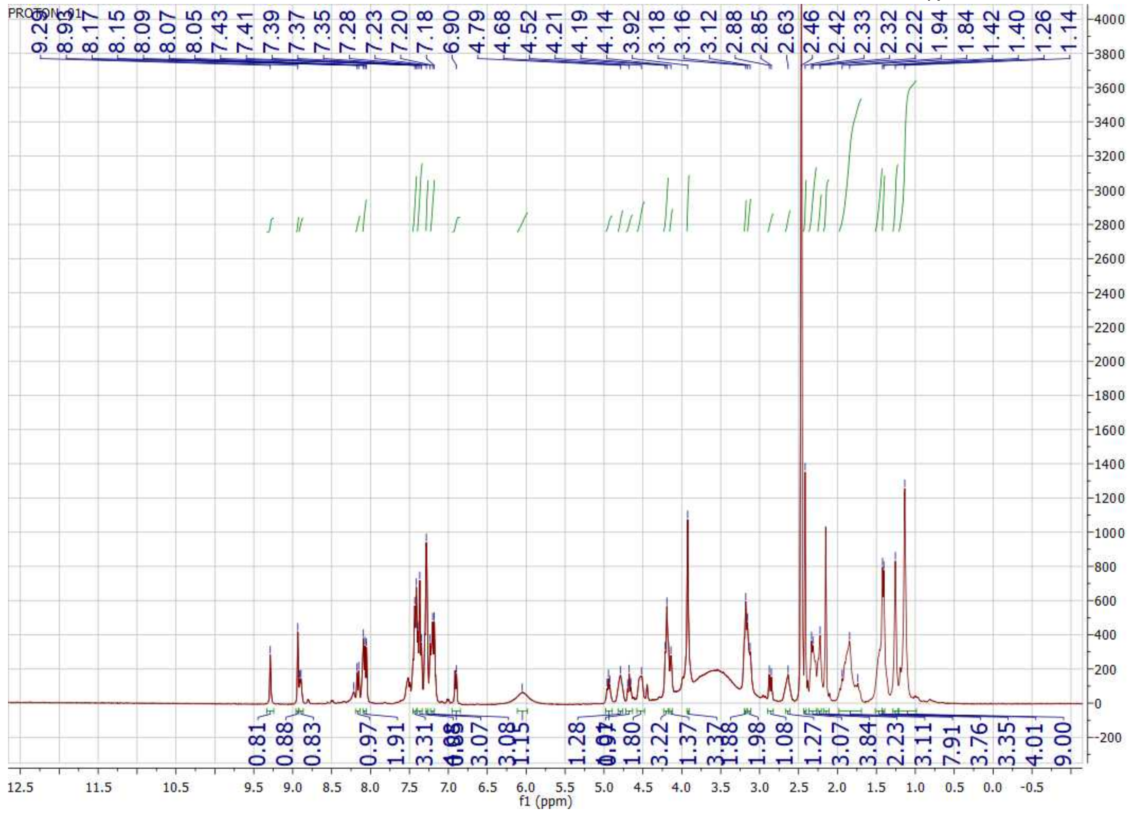

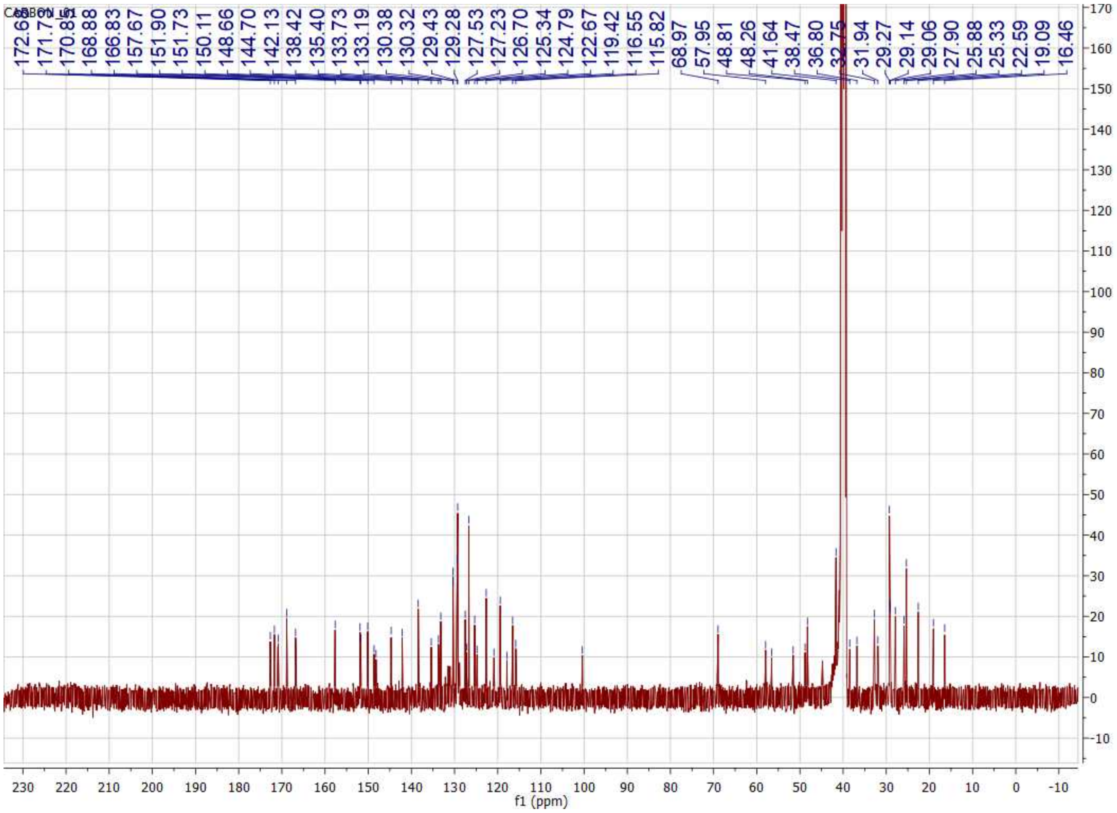

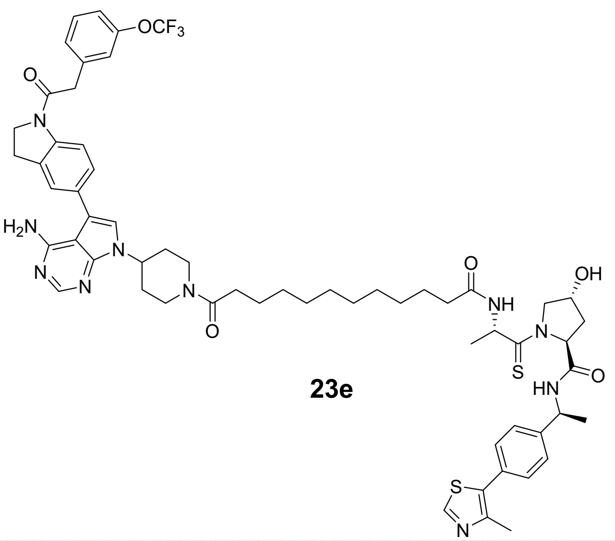

**23e**

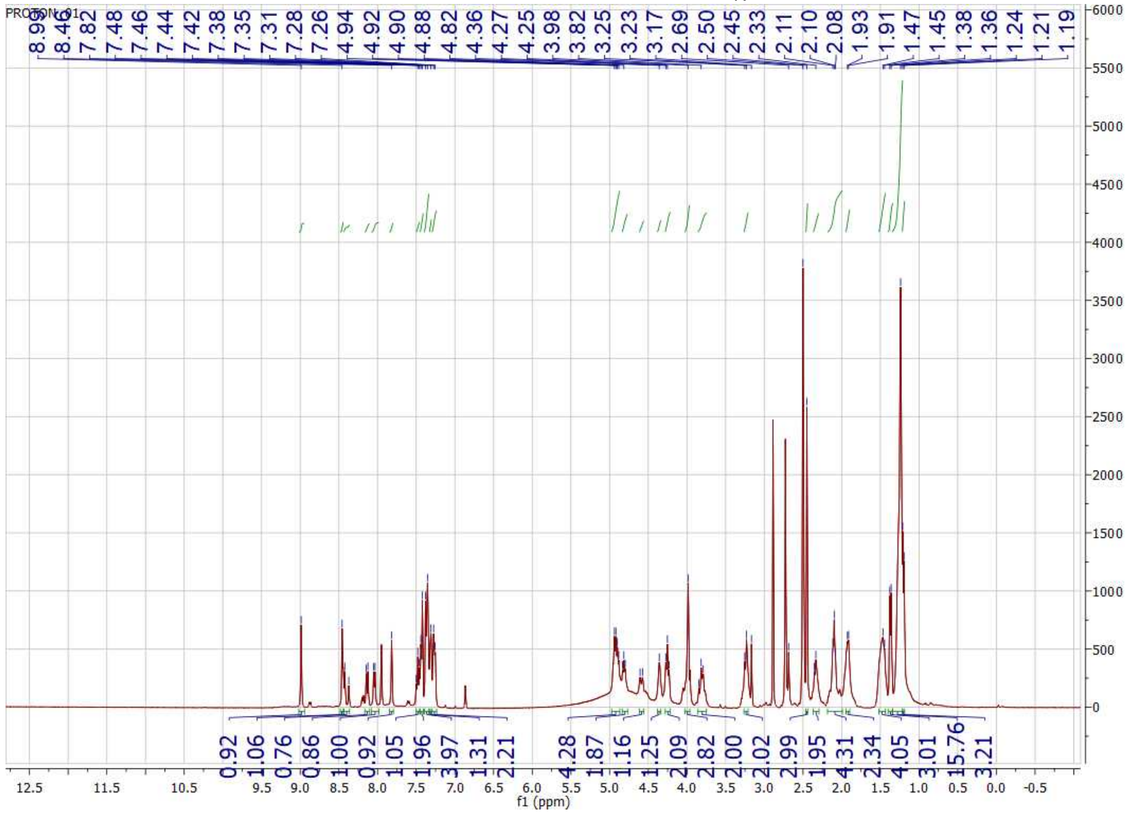

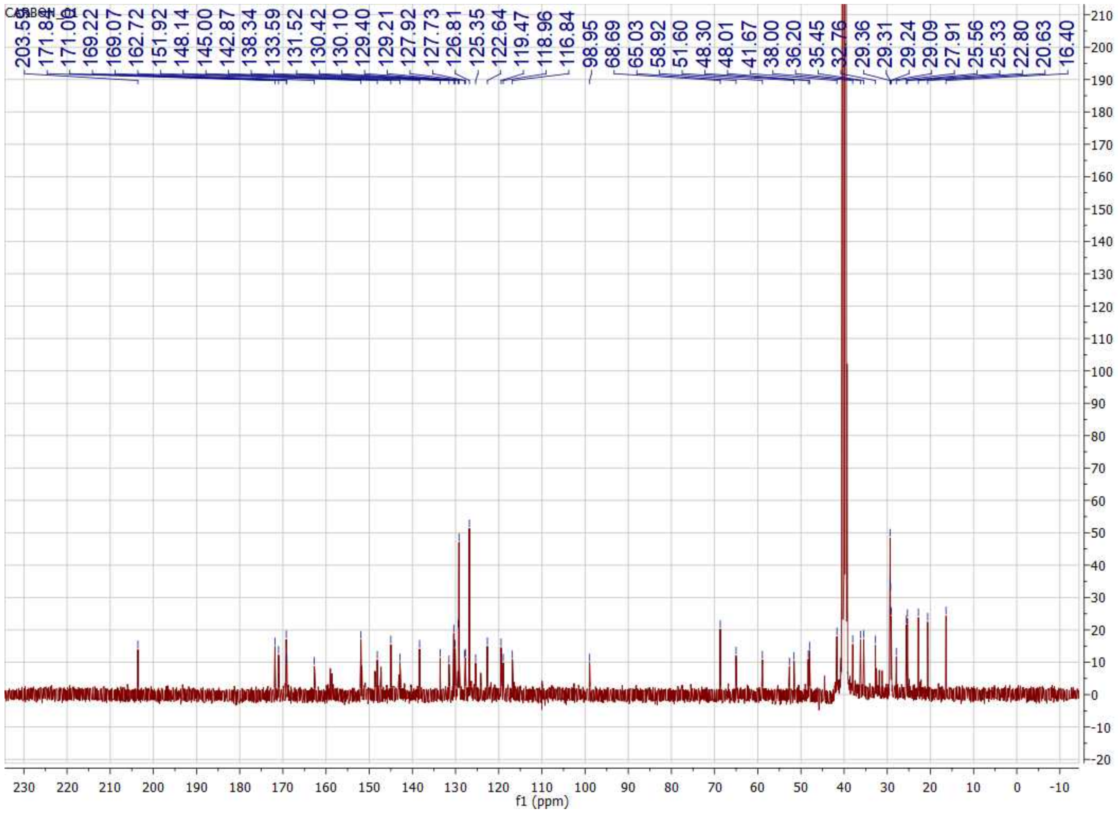
CARBON_01
203.55
171.84
171.01
169.22
169.07
162.72
151.92
148.14
145.00
142.87
138.34
133.59
131.52
130.42
130.10
129.40
129.21
127.92
127.73
126.81
125.35
122.64
119.47
118.96
116.84
98.95
68.69
65.03
58.92
51.60
48.30
48.01
41.67
38.00
36.20
35.45
32.76
29.36
29.31
29.24
29.09
27.91
25.56
25.33
22.80
20.63
16.40
f1 (ppm)

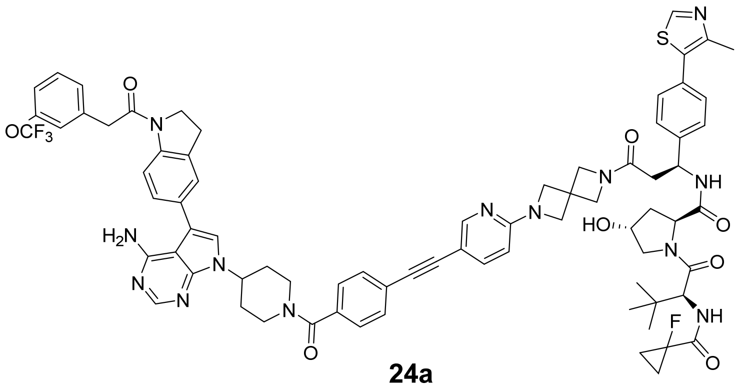

**24a**

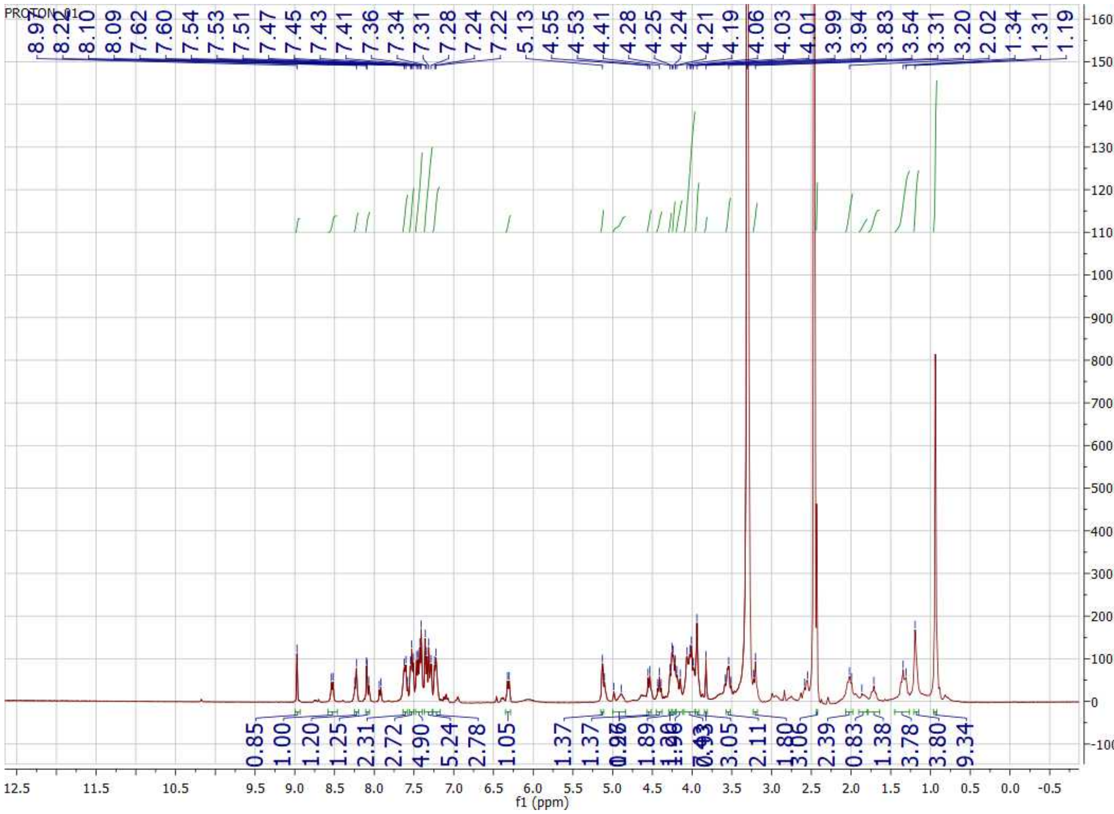
PROTON_01
8.97
8.22
8.10
8.09
7.62
7.60
7.54
7.53
7.51
7.47
7.45
7.43
7.41
7.36
7.34
7.31
7.28
7.24
7.22
5.13
4.55
4.53
4.41
4.28
4.25
4.24
4.21
4.19
4.06
4.03
4.01
3.99
3.94
3.83
3.54
3.31
3.20
2.02
1.34
1.31
1.19
0.85
1.00
1.20
1.25
2.31
2.72
4.90
5.24
2.78
1.05
1.37
1.37
1.89
3.05
2.11
1.80
3.06
2.39
0.83
1.38
3.78
3.80
9.34
f1 (ppm)

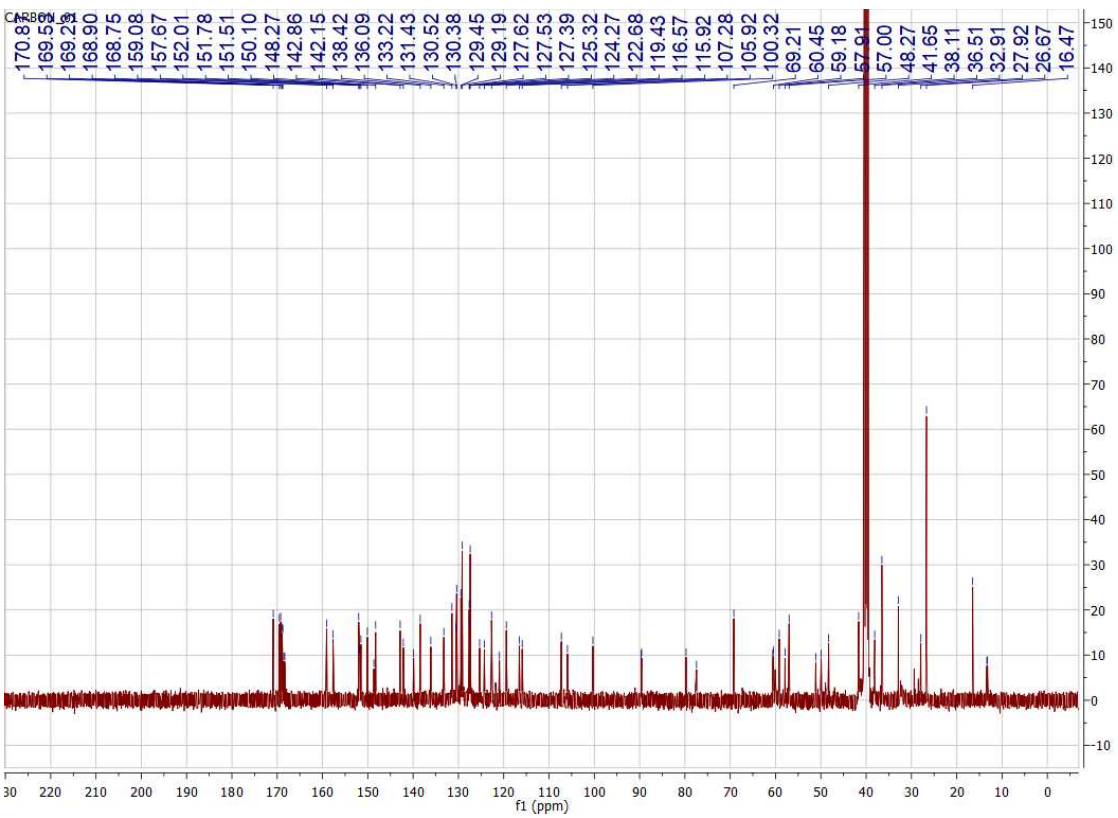
CARBON_01
170.8
169.53
169.29
168.90
168.75
159.08
157.67
152.01
151.78
151.51
150.10
148.27
142.86
142.15
138.42
136.09
133.22
131.43
130.52
130.38
129.45
129.19
127.62
127.53
127.39
125.32
124.27
122.68
119.43
116.57
115.92
107.28
105.92
100.32
69.21
60.45
59.18
57.91
57.00
48.27
41.65
38.11
36.51
32.91
27.92
26.67
16.47
f1 (ppm)

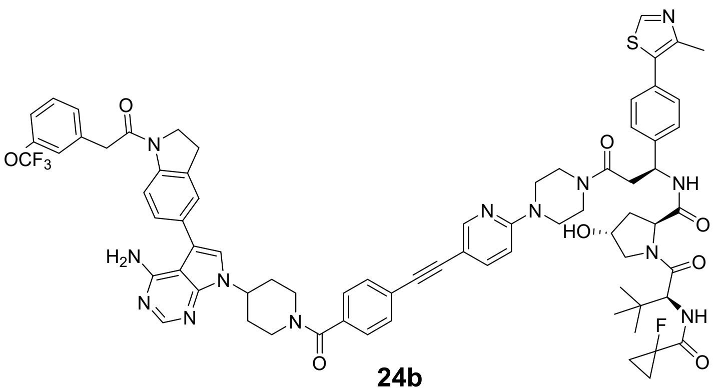

**24b**

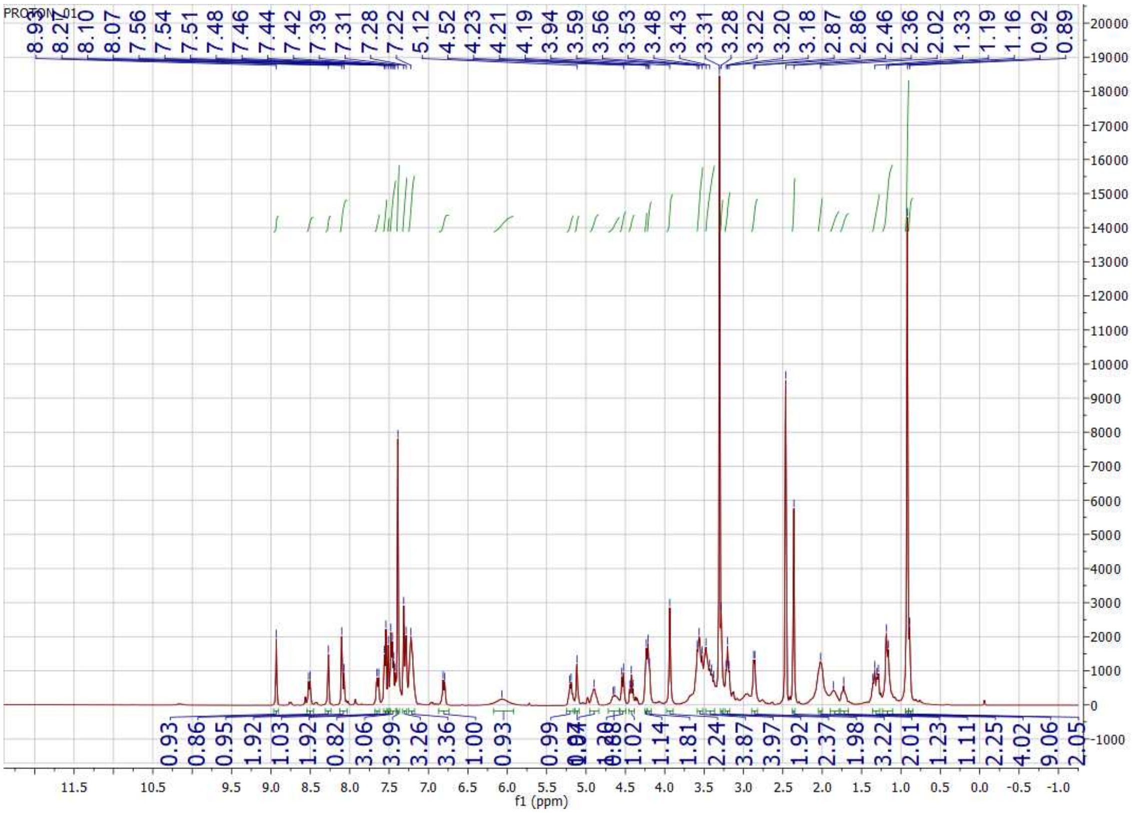

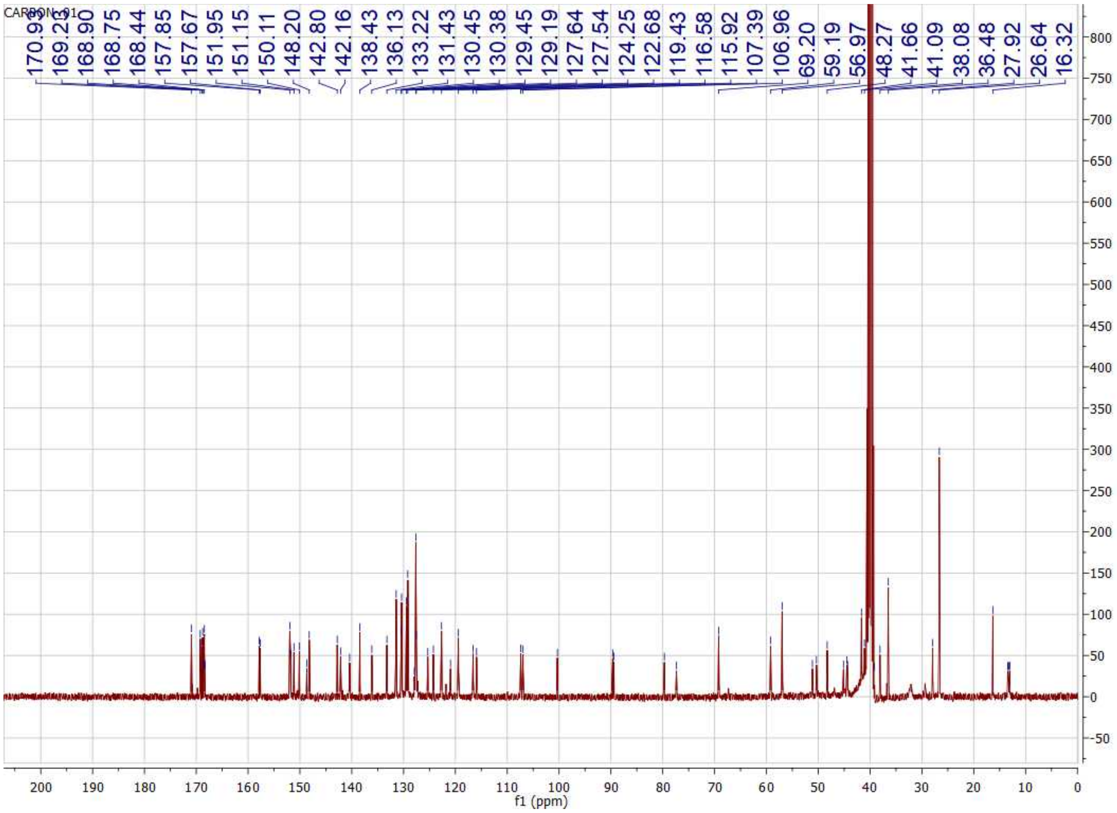
CARBON_01
170.93
169.22
168.90
168.75
168.44
157.85
157.67
151.95
151.15
150.11
148.20
142.80
142.16
138.43
136.13
133.22
131.43
130.45
130.38
129.45
129.19
127.64
127.54
124.25
122.68
119.43
116.58
115.92
107.39
106.96
69.20
59.19
56.97
48.27
41.66
41.09
38.08
36.48
27.92
26.64
16.32
f1 (ppm)

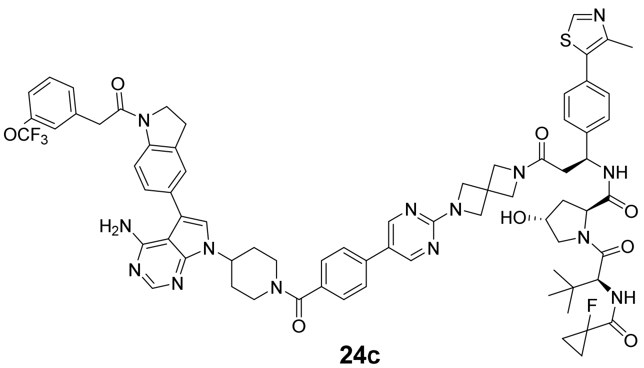

OCF3
H2N
HO
NH
F

**24c**

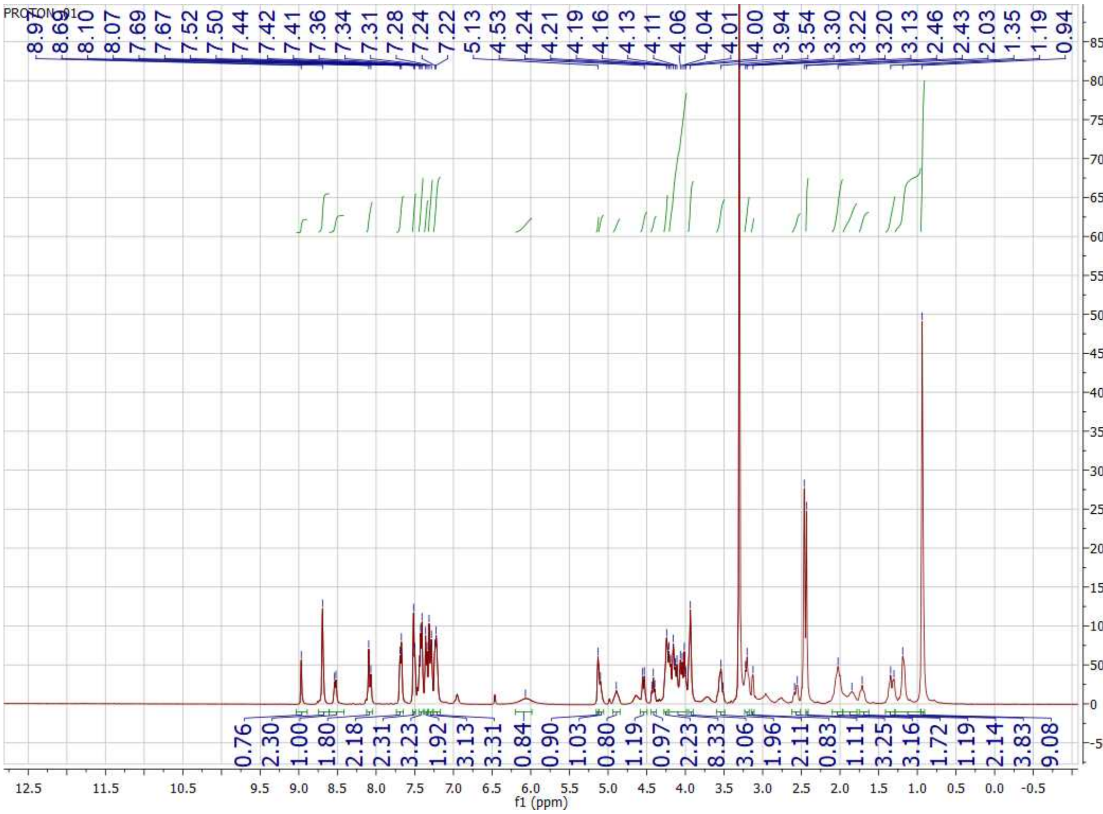

PROTON_01
8.62 8.10 8.07 7.69 7.67 7.52 7.50 7.44 7.42 7.41 7.36 7.34 7.31 7.28 7.24 7.22 5.13 4.53 4.24 4.21 4.19 4.16 4.13 4.11 4.06 4.04 4.01 4.00 3.94 3.54 3.30 3.22 3.20 3.13 2.46 2.43 2.03 1.35 1.19 0.94
0.76 2.30 1.00 1.80 2.18 2.31 3.23 1.92 3.13 3.31 0.84 0.90 1.03 0.80 1.19 0.97 2.23 8.33 3.06 1.96 2.11 0.83 1.11 3.25 3.16 1.72 1.19 2.14 3.83 9.08
f1 (ppm)

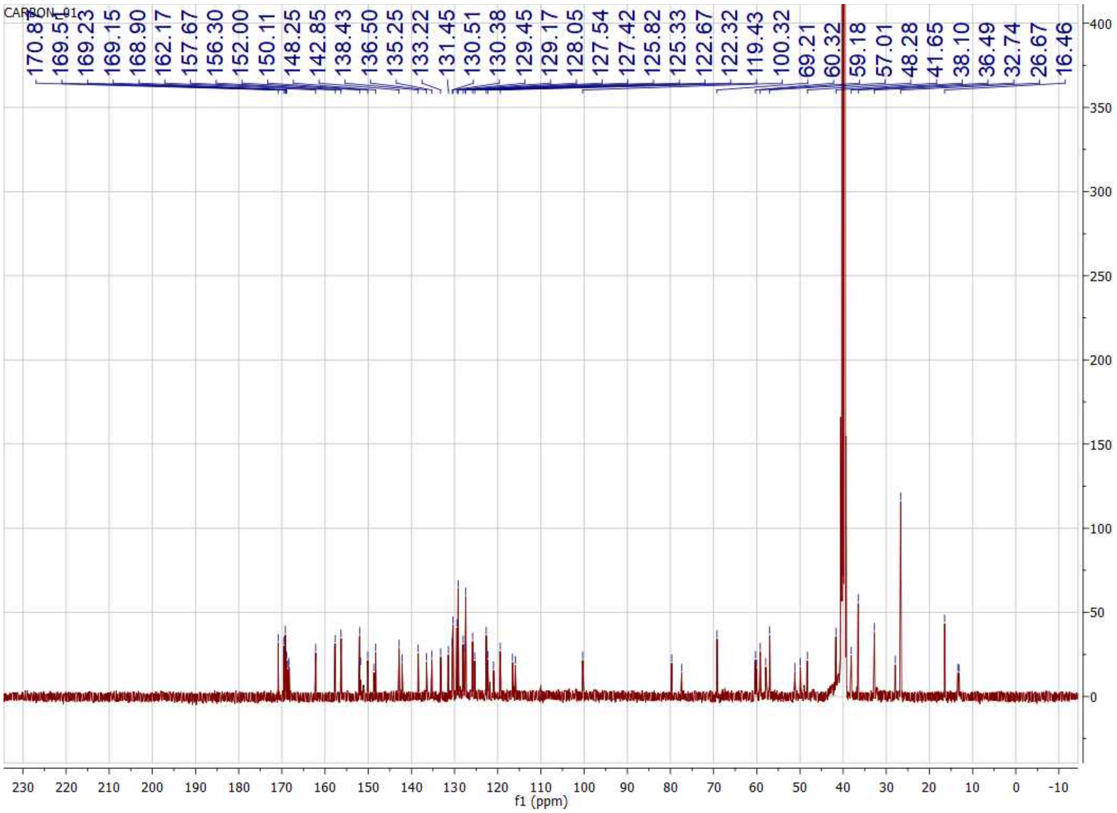
CARBON_01
170.81
169.51
169.23
169.15
168.90
162.17
157.67
156.30
152.00
150.11
148.25
142.85
138.43
136.50
135.25
133.22
131.45
130.51
130.38
129.45
129.17
128.05
127.54
127.42
125.82
125.33
122.67
122.32
119.43
100.32
69.21
60.32
59.18
57.01
48.28
41.65
38.10
36.49
32.74
26.67
16.46
f1 (ppm)

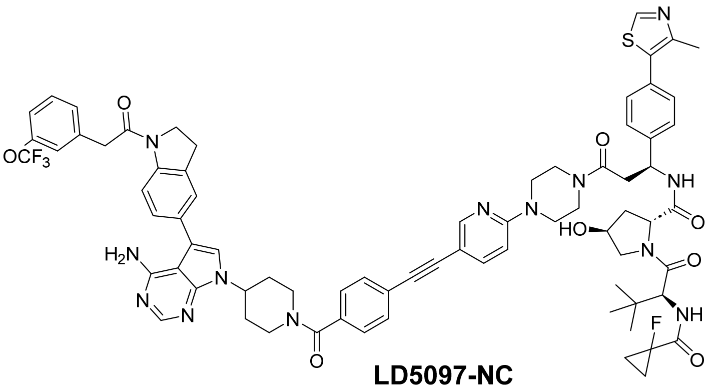

**LD5097-NC**

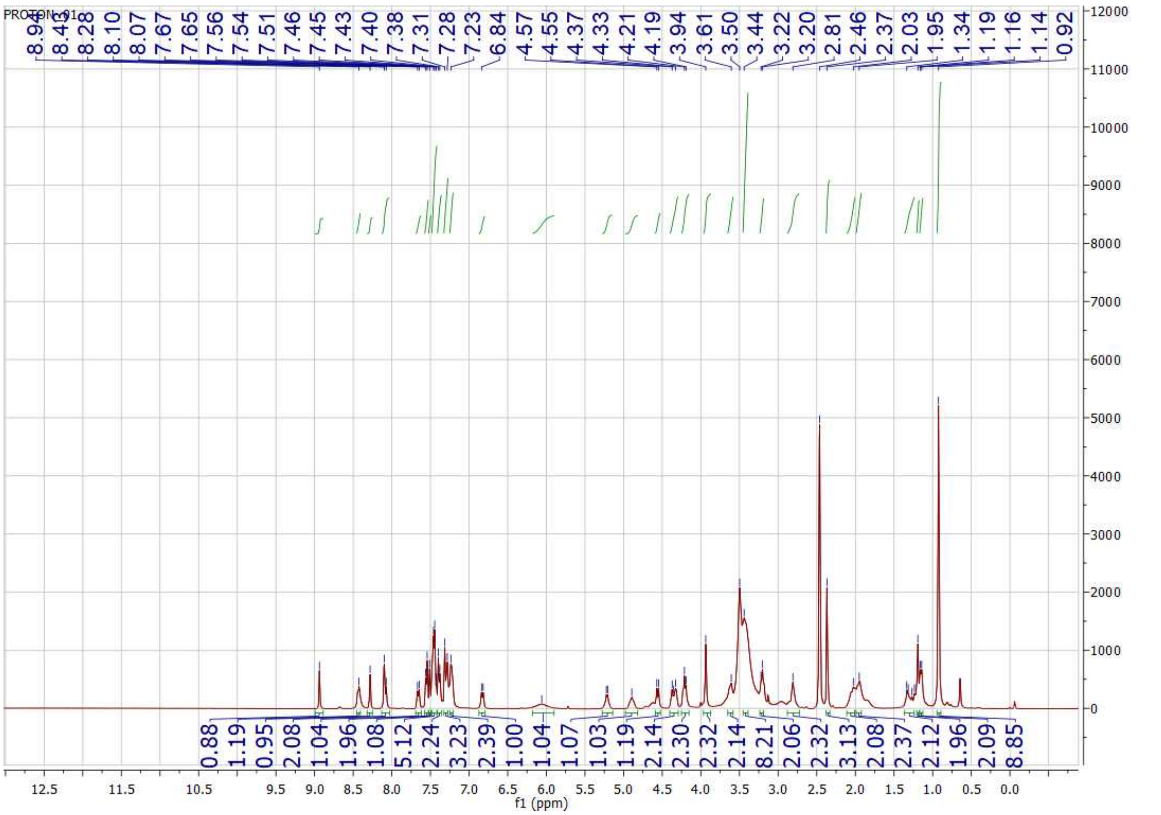

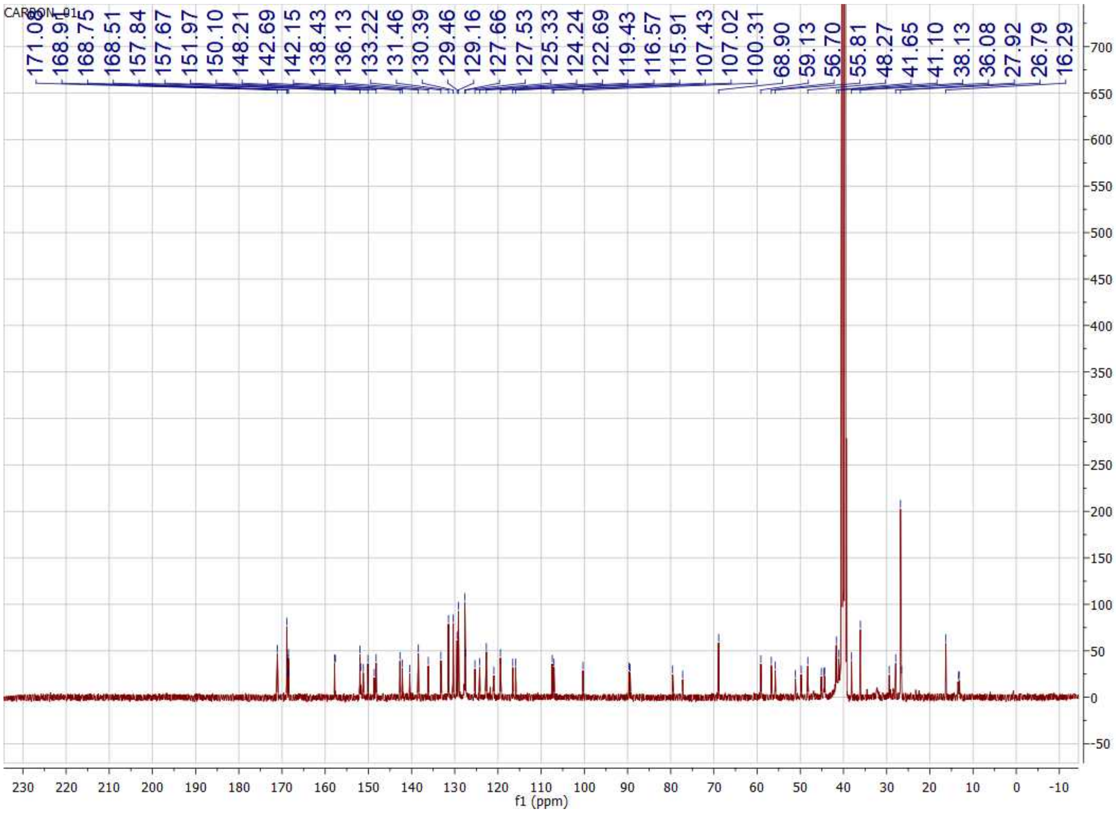
CARBON_01
171.08
168.91
168.75
168.51
157.84
157.67
151.97
150.10
148.21
142.69
142.15
138.43
136.13
133.22
131.46
130.39
129.46
129.16
127.66
127.53
125.33
124.24
122.69
119.43
116.57
115.91
107.43
107.02
100.31
68.90
59.13
56.70
55.81
48.27
41.65
41.10
38.13
36.08
27.92
26.79
16.29
f1 (ppm)

### HPLC chromatography of representative compounds

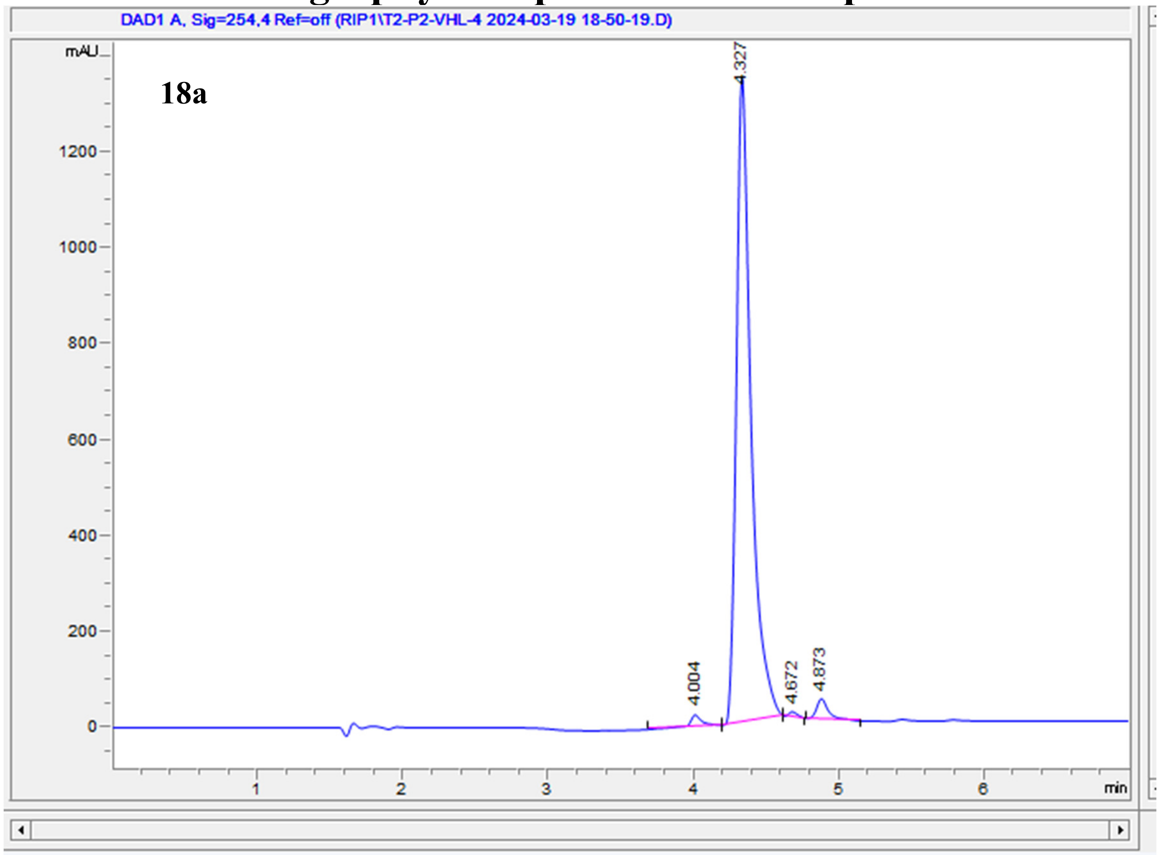

| # | Time | Area | Height | Width | Area% | Symmetry |
|---|---|---|---|---|---|---|
| 1 | 4.004 | 144.5 | 24.7 | 0.0823 | 1.458 | 0.753 |
| 2 | 4.327 | 9482.2 | 1352.6 | 0.1018 | 95.729 | 0.572 |
| 3 | 4.672 | 42.9 | 10.7 | 0.0626 | 0.433 | 0.612 |
| 4 | 4.873 | 235.8 | 42.4 | 0.0809 | 2.380 | 0.626 |

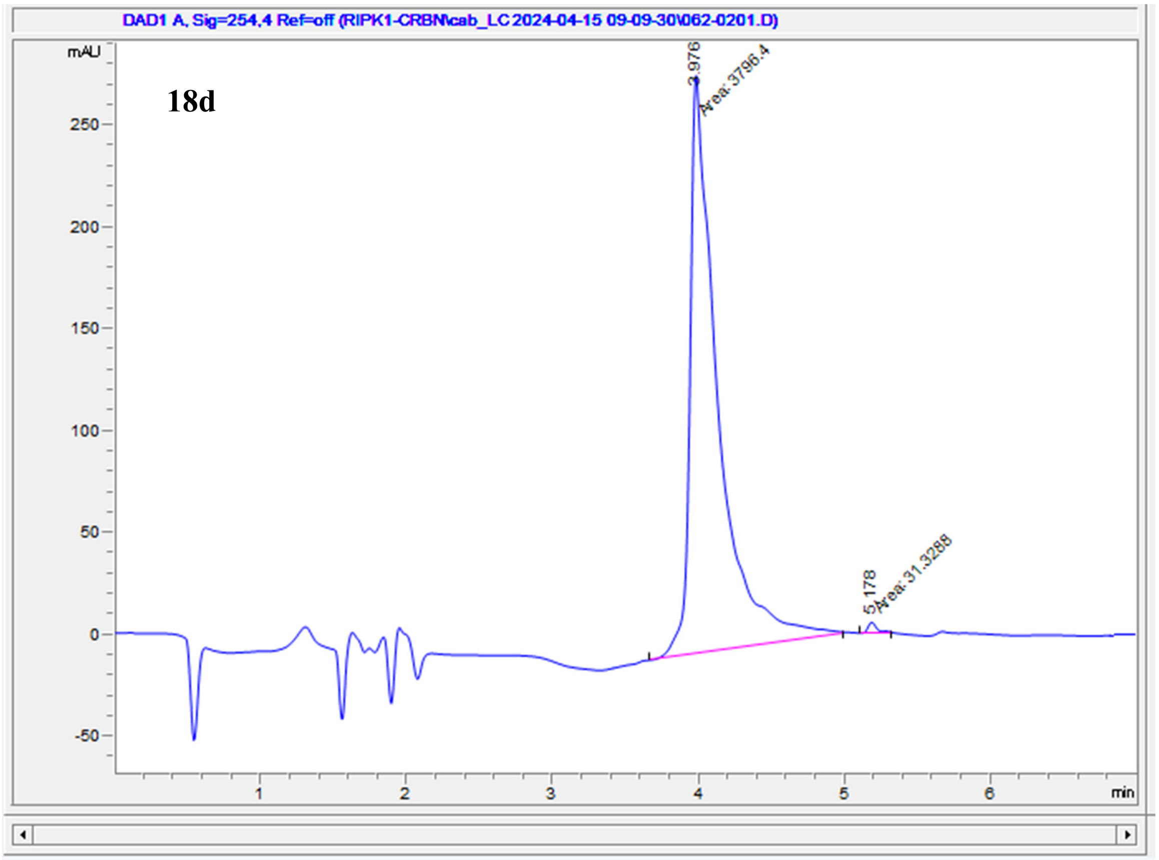

| # | Time | Area | Height | Width | Area% | Symmetry |
|---|---|---|---|---|---|---|
| 1 | 3.976 | 3796.4 | 284.4 | 0.2225 | 99.182 | 0.281 |
| 2 | 5.178 | 31.3 | 5.9 | 0.0884 | 0.818 | 0.589 |

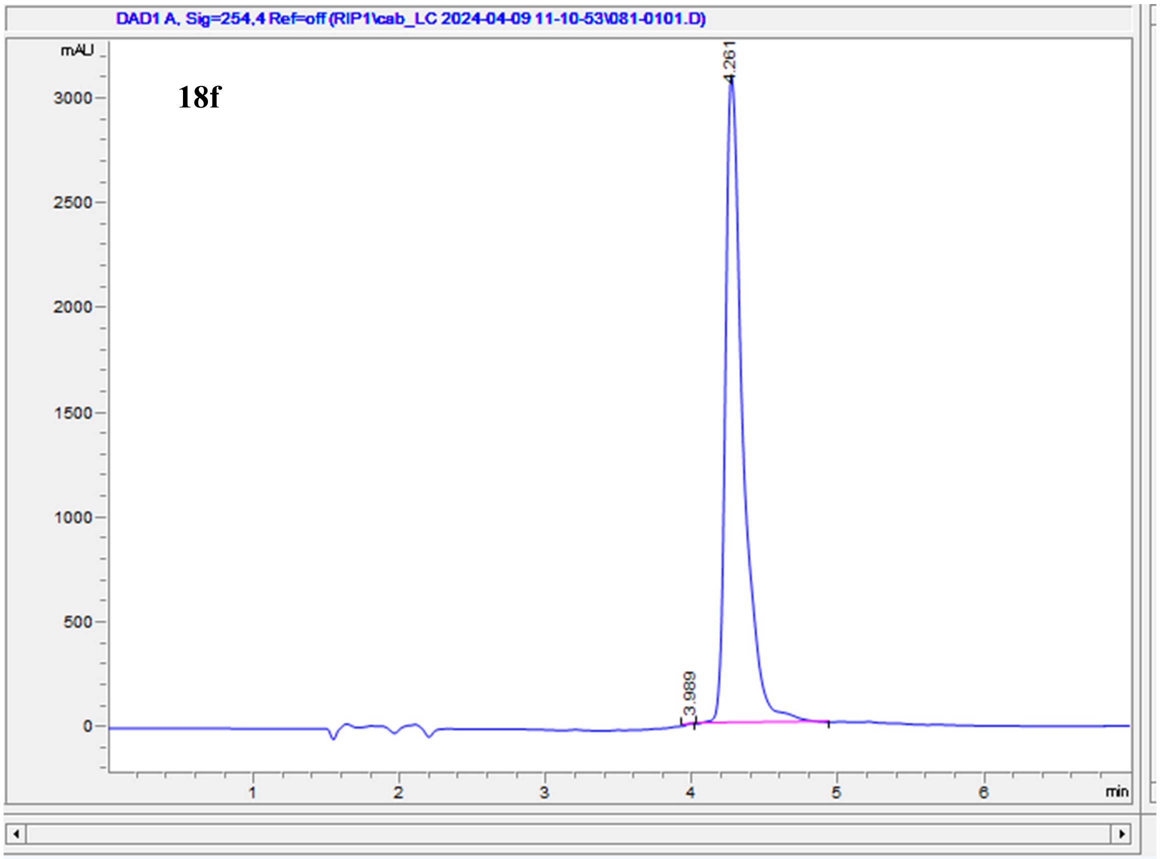

| # | Time | Area | Height | Width | Area% | Symmetry |
|---|---|---|---|---|---|---|
| 1 | 3.989 | 18.2 | 5.3 | 0.0536 | 0.070 | 2.121 |
| 2 | 4.261 | 25977.9 | 3098.1 | 0.1238 | 99.930 | 0.477 |

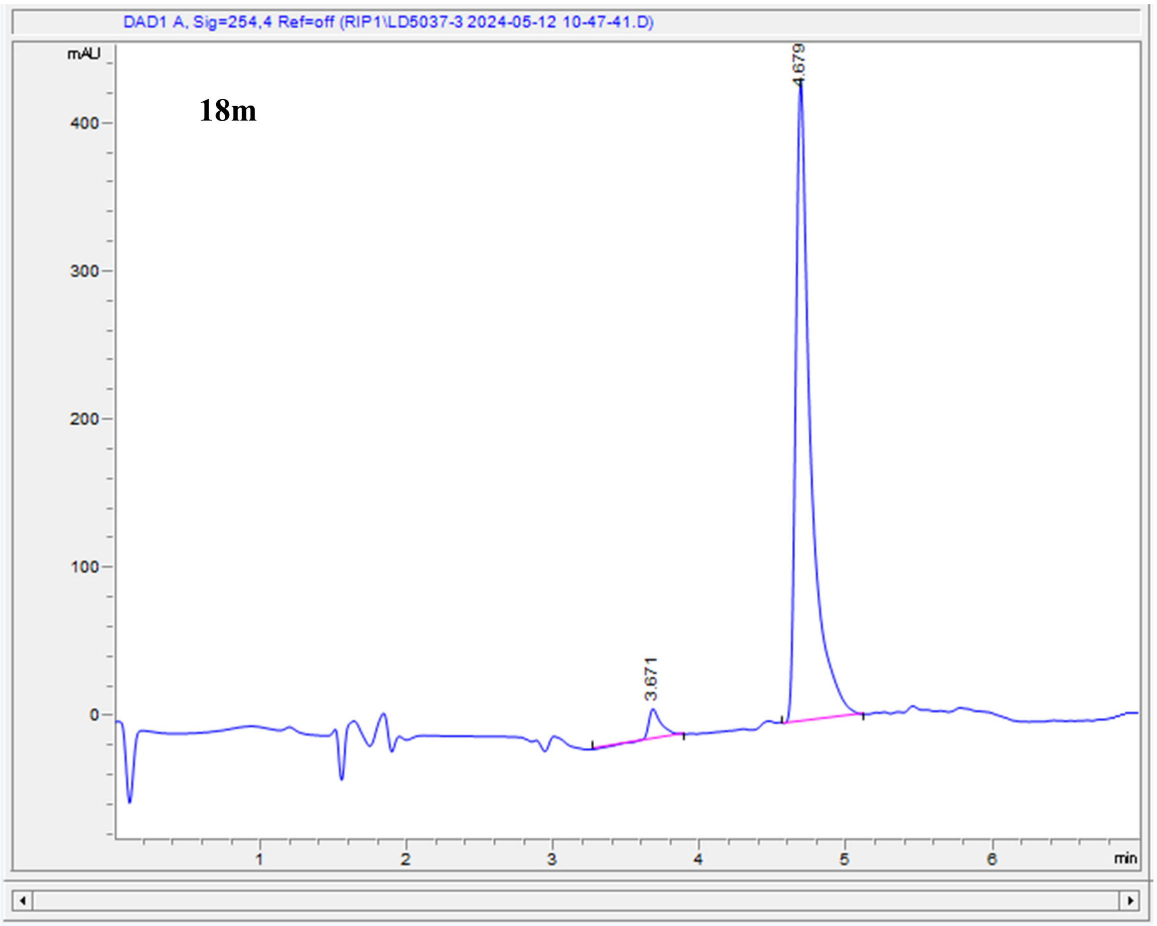

| # | Time | Area | Height | Width | Area% | Symmetry |
|---|---|---|---|---|---|---|
| 1 | 3.671 | 136.5 | 20.8 | 0.0925 | 4.274 | 0.601 |
| 2 | 4.679 | 3058 | 432.4 | 0.1005 | 95.726 | 0.441 |

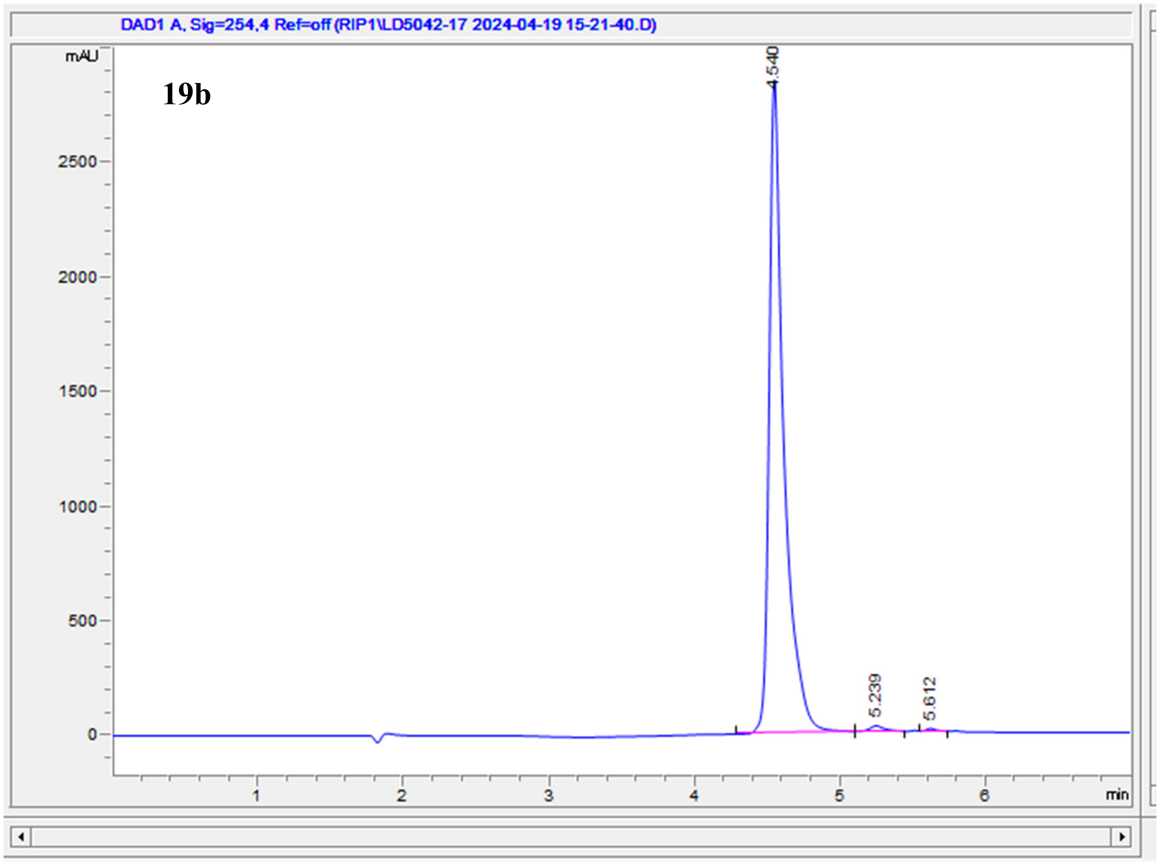

| # | Time | Area | Height | Width | Area% | Symmetry |
|---|---|---|---|---|---|---|
| 1 | 4.54 | 19959.1 | 2851.4 | 0.0997 | 98.898 | 0.502 |
| 2 | 5.239 | 172.2 | 24.5 | 0.1 | 0.853 | 0.638 |
| 3 | 5.612 | 50.3 | 11.3 | 0.0657 | 0.249 | 0.627 |

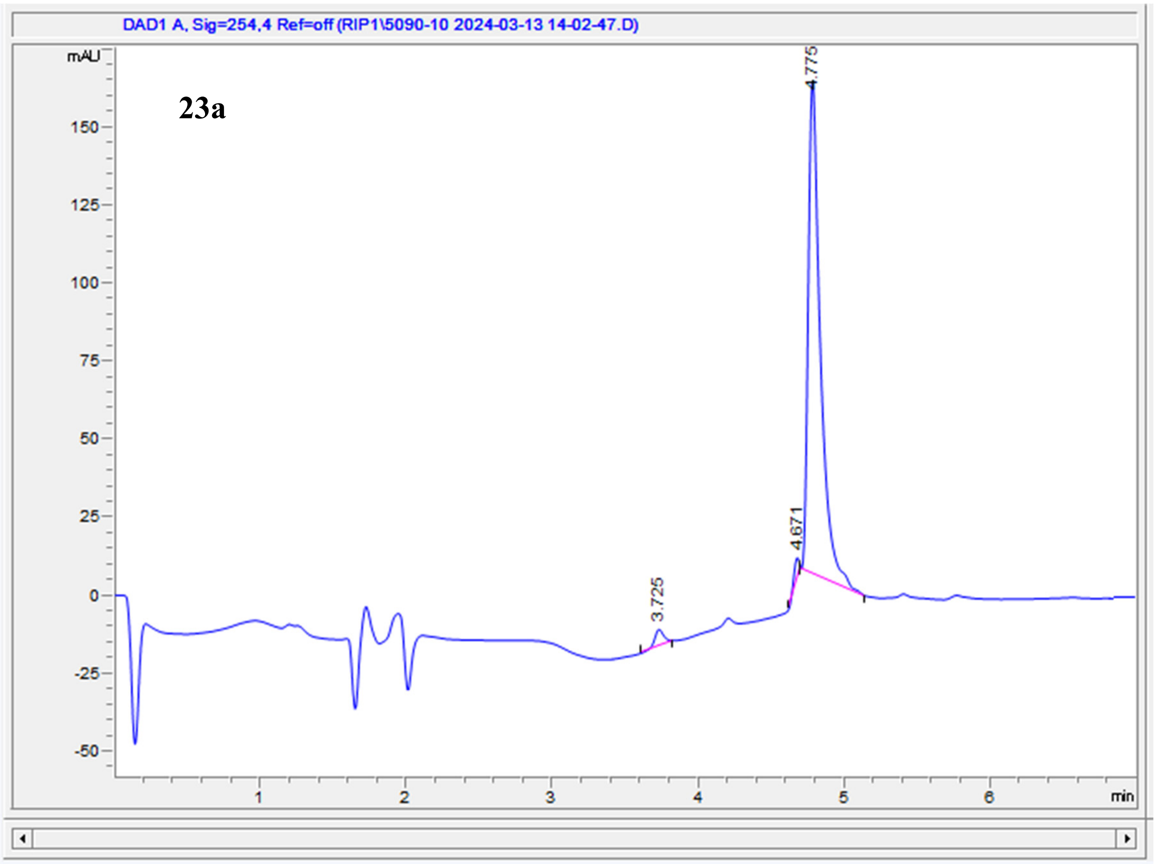

| # | Time | Area | Height | Width | Area% | Symmetry |
|---|---|---|---|---|---|---|
| 1 | 3.725 | 23.7 | 5.4 | 0.0674 | 2.503 | 0.91 |
| 2 | 4.671 | 20.7 | 7.5 | 0.0461 | 2.186 | 2.164 |
| 3 | 4.775 | 903.4 | 157.8 | 0.0828 | 95.311 | 0.497 |

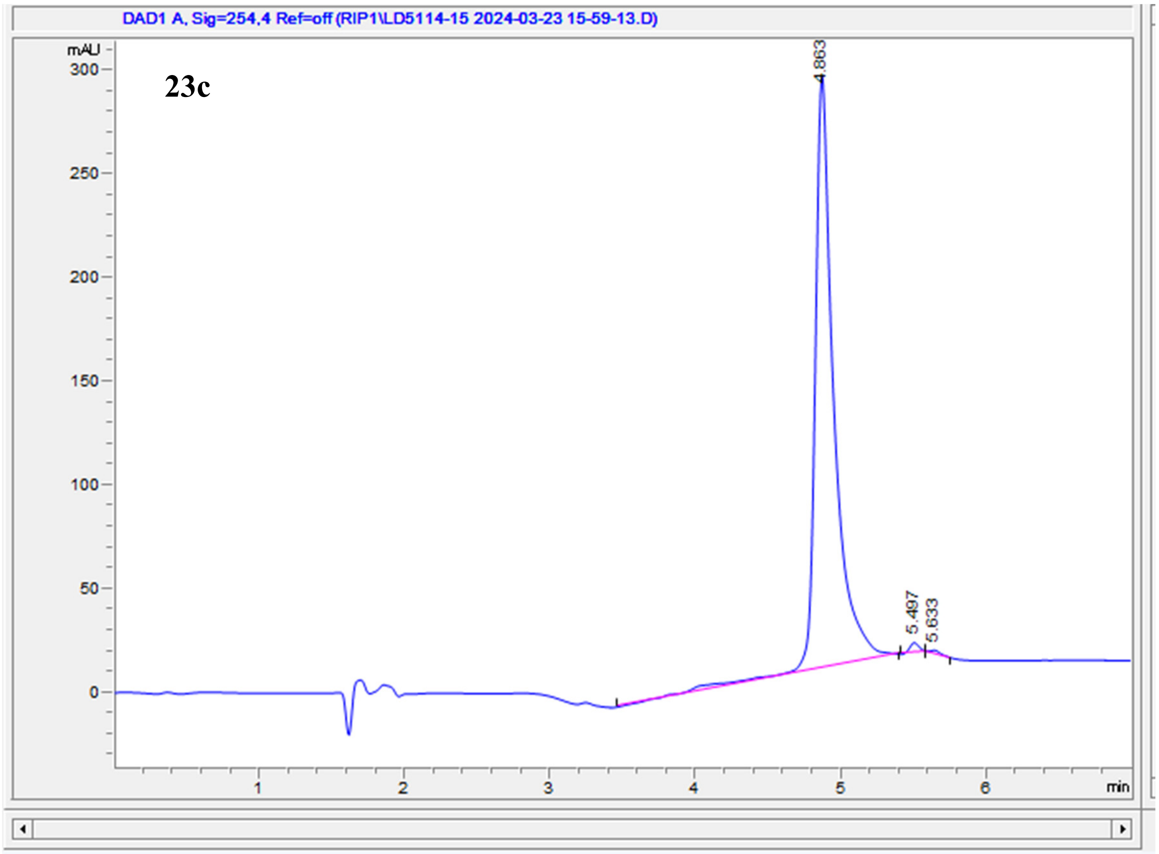

| # | Time | Area | Height | Width | Area% | Symmetry |
|---|---|---|---|---|---|---|
| 1 | 4.863 | 2640 | 287.1 | 0.1293 | 98.958 | 0.586 |
| 2 | 5.497 | 21.9 | 4.9 | 0.0686 | 0.822 | 0.943 |
| 3 | 5.633 | 5.9 | 1.3 | 0.0685 | 0.221 | 0.399 |

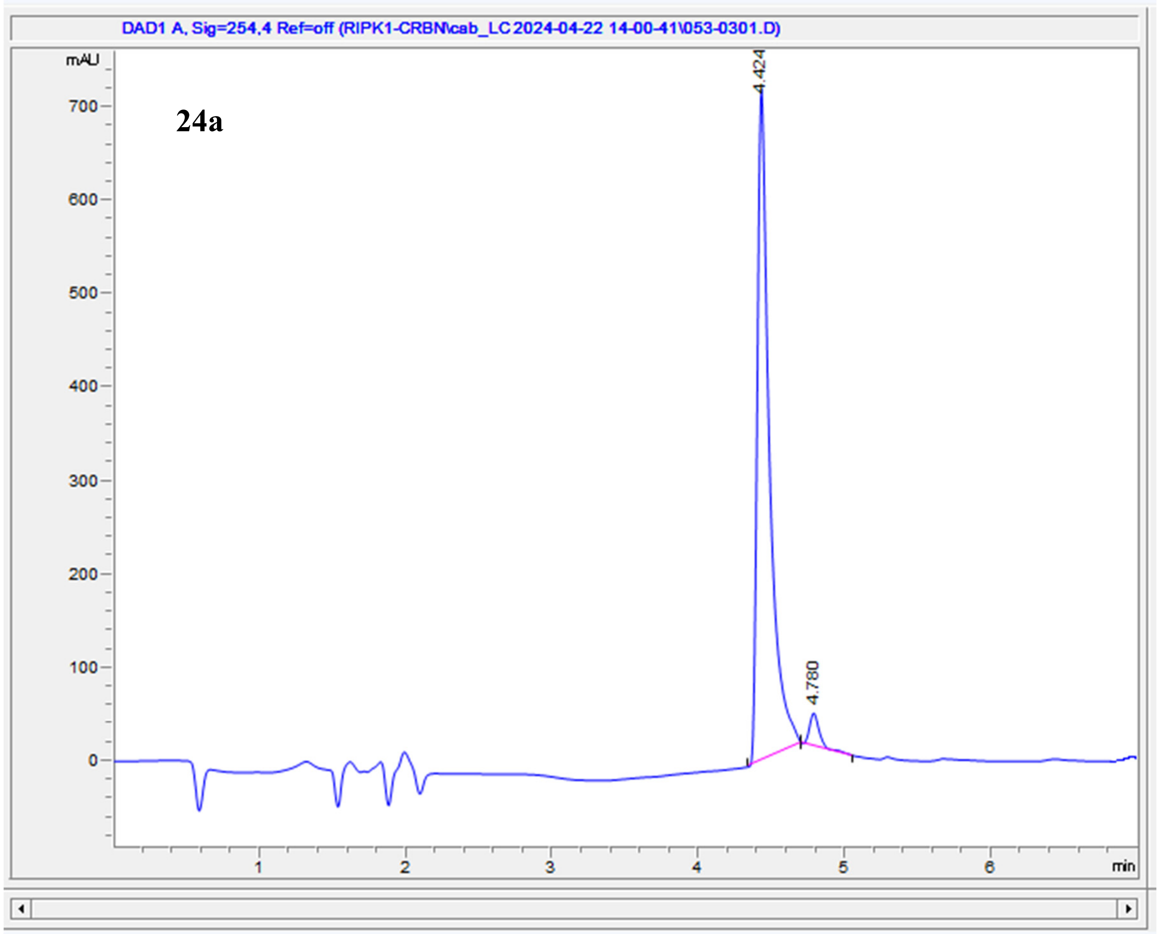

| # | Time | Area | Height | Width | Area% | Symmetry |
|---|---|---|---|---|---|---|
| 1 | 4.424 | 4208 | 722.1 | 0.084 | 96.131 | 0.457 |
| 2 | 4.78 | 169.4 | 36.1 | 0.0707 | 3.869 | 0.654 |

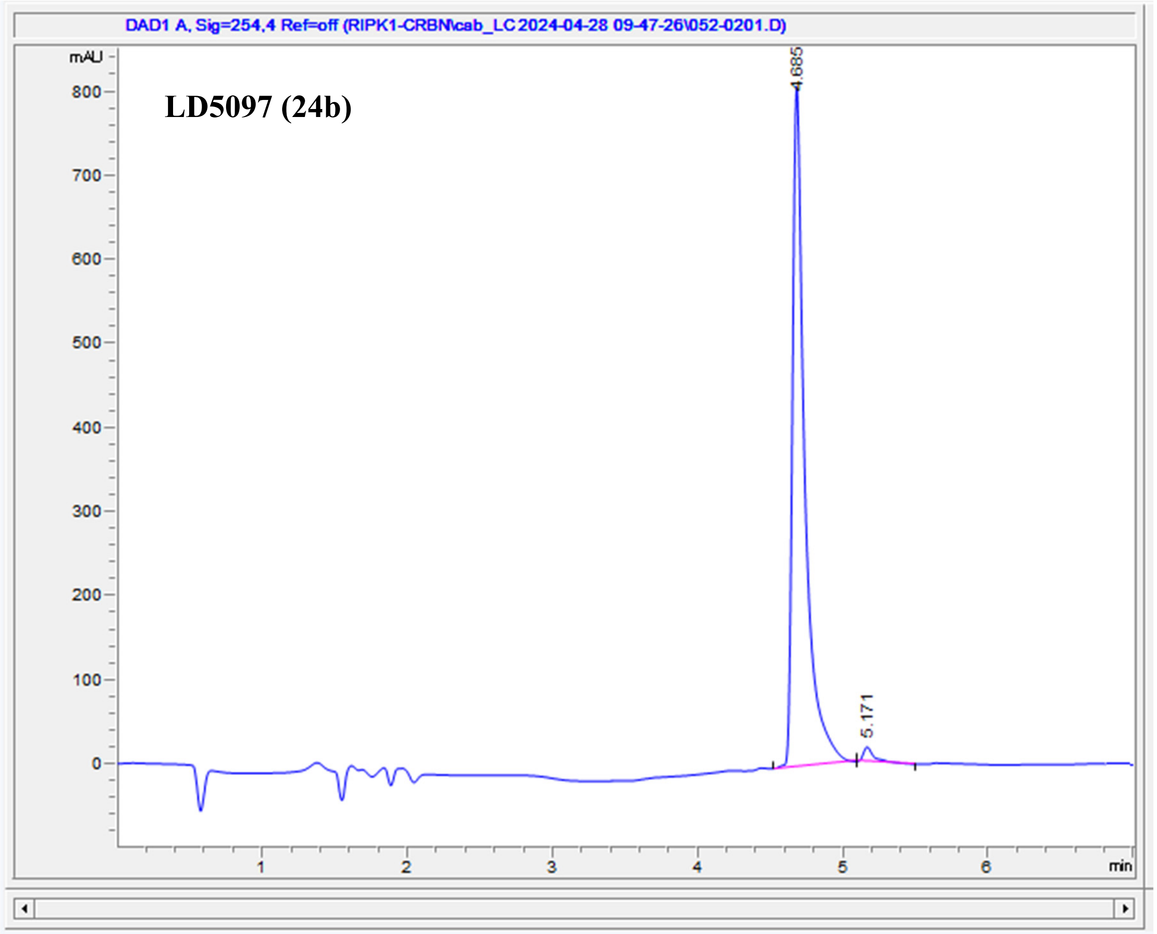

| # | Time | Area | Height | Width | Area% | Symmetry |
|---|---|---|---|---|---|---|
| 1 | 4.685 | 4806.8 | 814.2 | 0.0849 | 98.067 | 0.458 |
| 2 | 5.171 | 94.7 | 17.9 | 0.0756 | 1.933 | 0.466 |

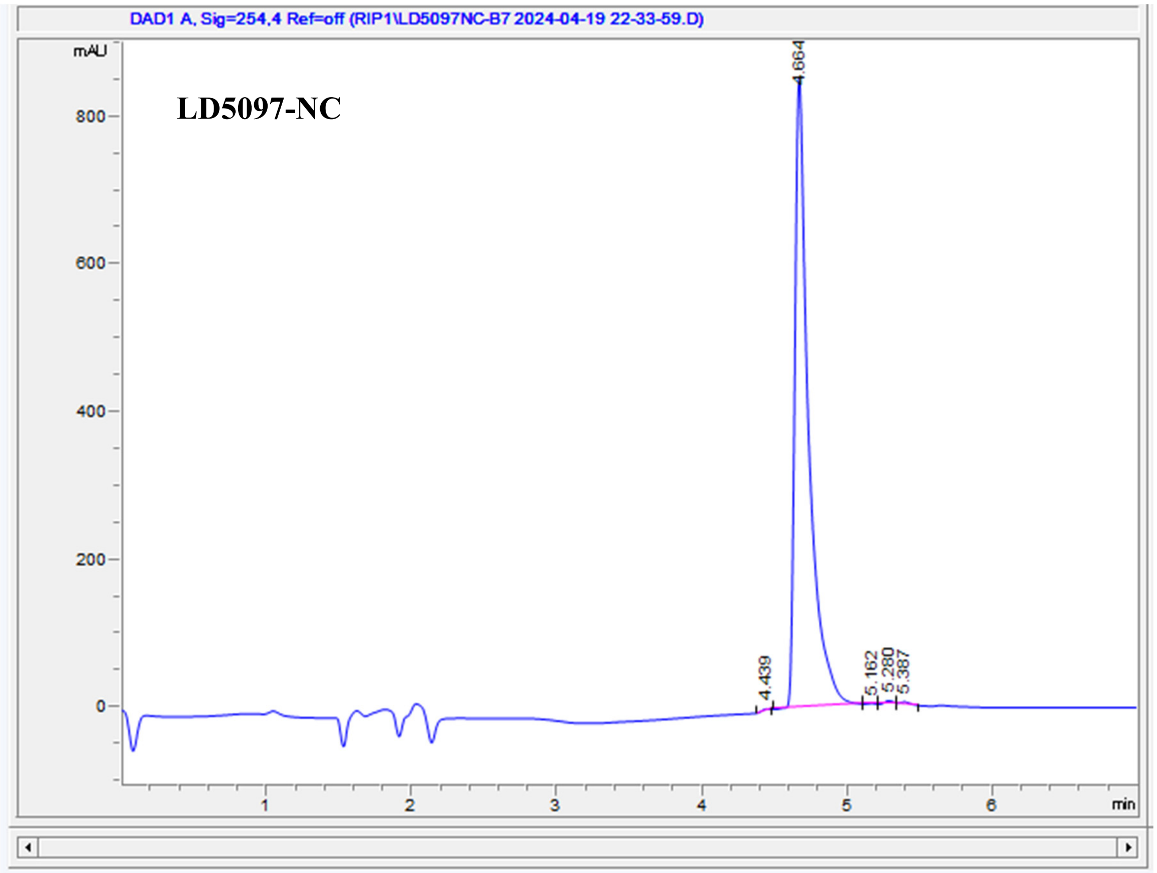

| # | Time | Area | Height | Width | Area% | Symmetry |
|---|---|---|---|---|---|---|
| 1 | 4.439 | 9.6 | 2.6 | 0.0566 | 0.168 | 1.652 |
| 2 | 4.664 | 5653.5 | 855.9 | 0.0931 | 99.351 | 0.425 |
| 3 | 5.162 | 3.2 | 1.1 | 0.0487 | 0.056 | 0.949 |
| 4 | 5.28 | 15 | 4 | 0.0622 | 0.264 | 0.948 |
| 5 | 5.387 | 9.2 | 2.5 | 0.0571 | 0.161 | 0.49 |
